# An alternative framework for transcriptome-wide association studies to detect and decipher gene-trait associations

**DOI:** 10.1101/2025.03.14.643391

**Authors:** Zheng Li, Boran Gao, Xiang Zhou

## Abstract

Transcriptome-wide association studies (TWAS) are widely used to uncover transcriptomic mechanisms underlying disease. Here, we present VINTAGE, an alternative TWAS framework designed to identify and decipher gene-trait associations through two complementary tests: a genetic variance test that generalizes TWAS and unifies it with SKAT, offering a clearer understanding of TWAS false signals when expression is not relevant to SNP-trait associations; and a local genetic correlation test that distinguishes it from TWAS by explicitly quantifying and testing the proportion of genetic effects on trait mediated through gene expression. Applied to eQTLGen and eighteen traits from UK Biobank, VINTAGE improved gene-trait association power by an average of 5% and 87% over SKAT and TWAS, respectively. VINTAGE is also the only method effective in assessing potential gene mediation effects, revealing that most genes lack detectable mediation effects (median = 12%), which explains the power advantage of VINTAGE and SKAT over TWAS. Notably, VINTAGE identified 61 genes with significant mediation, highlighting the role of expression in genetic influences on traits.

## Introduction

Over the past two decades, genome-wide association studies (GWASs) have unveiled many genetic variants associated with various diseases and disease-related complex traits^1^. Nevertheless, the molecular mechanisms underlying the majority of these identified associations remain unclear, hindering the translation of discoveries into clinical practices^2^. One potential molecular mechanism through which a genetic variant influences a complex trait involves gene regulation, where changes in gene expression may act as a mediating factor^3^. Such mechanism is supported by the enrichment of GWAS signals in expression quantitative trait loci (eQTLs) for multiple traits^3-6^ and has been investigated through transcriptome-wide association study (TWAS)^7,8^. TWAS integrates gene expression mapping studies with GWASs and aims to identify genes whose genetically regulated expression (GReX) is associated with a trait of interest. Conventional TWAS methods involve a two-stage analytic process. In the first stage, TWAS builds expression prediction models in the expression study where SNPs are used to predict gene expression. In the second stage, TWAS uses the estimated SNP prediction weights to construct GReX in the GWAS and tests the association of GReX with the trait. By integrating the gene expression mapping study with the GWAS, TWAS effectively aggregates SNP associations in the GWAS into interpretable functional units of genes, thereby facilitating the regulatory interpretation of the results. Numerous TWASs have been carried out, revealing important regulatory mechanisms underlying many complex traits^9-12^.

Despite the popularity of TWAS, there has been limited investigation into the validity of its modeling assumptions and, consequently, the proper interpretation of its association results. Originally, TWAS was designed with the primary intention of assessing the mediation effect of gene expression on the trait by testing for the association between GReX and the trait^7,8^. However, as shown here, the TWAS test statistic emerges as part of a score test statistic used to assess SNP effects on the trait. Consequently, TWAS effectively only tests the association between a set of SNPs with the trait without directly and explicitly testing for the SNP effects on the trait mediated through gene expression. Indeed, as shown here, TWAS can display significant p-values even when gene expression does not mediate the SNP effects on the trait. Moreover, TWAS makes a key modeling assumption: that the SNP effects on the trait are scalar multiplication of the SNP effects on gene expression. This assumption is highly restrictive, implying that the SNP effects inferred from the expression mapping study are fully informative for testing the SNP effects on the trait, indicating that, from a mediation analysis perspective, the SNP effects on the trait are entirely mediated through gene expression. Unfortunately, as shown here, this assumption does not hold for most genes in real world applications.

To overcome the aforementioned limitations of TWAS, we have developed an alternative framework, VINTAGE (**V**ariant-set test **IN**tegrative **T**W**A**S for **GE**ne-based analysis), for integrating GWAS with gene expression study. A unique feature of VINTAGE is its ability to assess the complexity of the gene-trait relationship from two distinct angles, using two complementary statistical tests: a genetic variance test that examines the combined effects of multiple SNPs in a gene region on the trait; and a genetic correlation test that examines the potential mediation effects of gene expression. Through these two distinct tests, VINTAGE provides complementary information to comprehensively characterize gene-trait relationship.

Importantly, VINTAGE’s genetic variance test relaxes the restricted TWAS modeling assumption by assuming that the SNP effects on trait are correlated with the SNP effects on gene expression, rather simply being a direct multiplication of the latter. As a result, VINTAGE transforms into TWAS only when the correlation parameter approaches either one or negative one. Surprisingly, when the correlation parameter approaches zero, VINTAGE transforms into SKAT^13^ (Sequence Kernel association Test), a widely used yet seemingly unrelated method, for common variant analysis. Consequently, VINTAGE’s genetic variance test unifies SKAT and TWAS into a coherent analytic framework, including both methods as special cases, thus achieving robust power on detecting gene-trait association across diverse scenarios, accommodating settings that favor either SKAT or TWAS, or fall somewhere in between. The unifying nature of VINTAGE’s genetic variance test framework reframes TWAS as a genetic variance test, similar to SKAT, rather than assessing the role of gene expression in associations. This perspective not only explains the false signals observed in TWAS, where the p-value is significant – and increasingly so with strong SNP effects on the trait – even when gene expression does not mediate any SNP effects on the trait, but also accounts for the surprising power advantage of SKAT over TWAS in real data applications, as most genes do not exhibit strong mediation effects in real datasets. On the other hand, VINTAGE’s genetic correlation test distinguishes it from TWAS by explicitly assessing the potential mechanism of gene expression through testing the correlation parameter, which quantifies the degree to which the genetic effects on trait are potentially mediated through gene expression. As a result, unlike TWAS, the significance of VINTAGE’s genetic correlation test indicates a potential mediation mechanism involving gene expression. We illustrate the benefits of VINTAGE through extensive simulations and real data applications to eighteen complex traits from UK Biobank.

## Results

### VINTAGE overview and proof-of-concept simulations

VINTAGE is described in the Materials and methods section, with technical details provided in the Supplementary Text. In brief, VINTAGE is a general framework for integrating a gene expression mapping study with a genome-wide association study (GWAS) to identify and decipher gene-trait associations (Figure 1). VINTAGE examines one gene at a time and performs two distinct tests: a genetic variance test aimed at identifying gene-trait associations, and a genetic correlation test aimed at identifying genes whose expression may mediate genetic effects on the trait. VINTAGE relies on a local genetic correlation parameter to both characterize the proportion of genetic effects on trait that is potentially mediated through gene expression and leverage the gene expression study to enhance gene-trait association analysis in GWAS. Through the local genetic correlation parameter, the genetic variance test in VINTAGE unifies two seemingly unrelated methods, SKAT and TWAS, into the same analytic framework, incorporating both as special cases.

**Figure 1.**
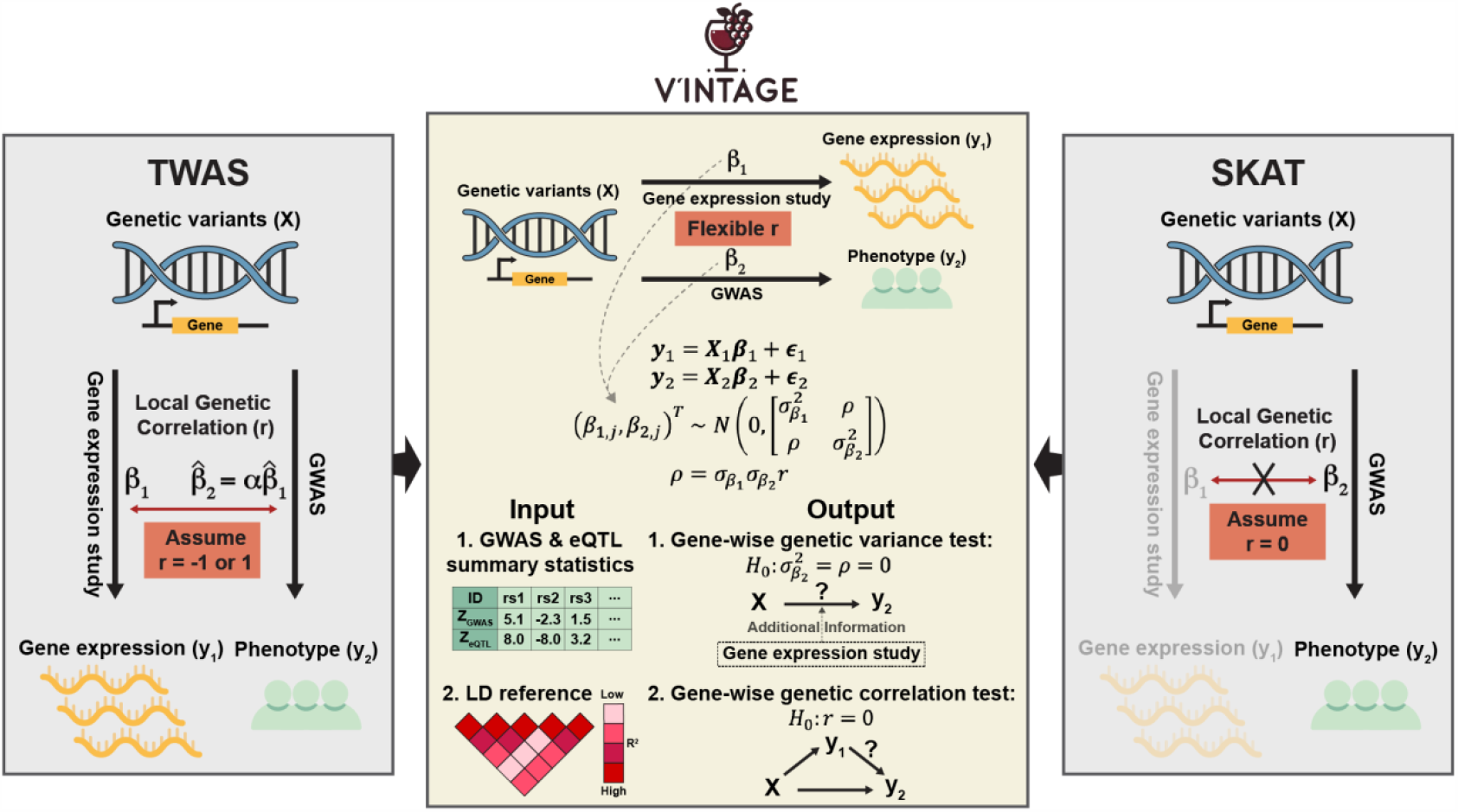
Schematic overview of VINTAGE. VINTAGE is a versatile analytic framework designed for integrating a gene expression mapping study with GWAS to identify genes associated with a trait of interest. VINTAGE can utilize either individual-level data or summary statistics. The summary statistics from GWAS and gene expression study are in the form of marginal Z scores, their sample sizes, and SNP-SNP correlation matrices constructed from a reference panel. VINTAGE evaluates the gene-trait relationship from two distinct angles, using two separate statistical tests: a genetic variance test that examines the combined effects of multiple SNPs in a gene region on the trait; and a genetic correlation test that examines the mediation effects of gene expression. The genetic variance test in VINTAGE effectively bridges SKAT and TWAS by explicitly modeling and inferring the local genetic correlation between gene expression and the trait, allowing it to achieve robust performance across diverse scenarios. The genetic correlation test in VINTAGE enables it to examine the extent to which gene expression mediates genetic effects on the trait. By combining insights from these two tests, VINTAGE provides a comprehensive characterization of the gene-trait relationship, enhancing our understanding of the molecular mechanisms underlying trait associations.

We conducted proof-of-concept simulations to illustrate the unifying nature of VINTAGE’s genetic variance test. To do so, we randomly selected 110,000 White British (WB) individuals from UKBB and divided them into an eQTL mapping study of 10,000 individuals and a GWAS of 100,000 individuals. We obtained genotype data from cis-SNPs of 10,000 genes from these individuals to simulate gene expression in the eQTL mapping study and a complex trait in the GWAS. In the proof-of-concept simulations, we examined type I error control and power of the individual genetic variance tests inside VINTAGE. Each individual test, denoted as *T*_*w*_, is constructed under a particular weight choice of *w* that controls the relative contribution of the variance score, which equals a key component of SKAT, versus the covariance score, which equals a key component of TWAS. In the simulations, the individual tests produced well-calibrated p-values regardless of the weights (Figures 2A and S1), and so are the combined p-values from these individual tests (Figure 2B). In addition, the power of the individual tests depends on whether the assigned weight matches the underlying local genetic correlation (Figures 3A and S2). Specifically, when *w* = 0, the test (*T*_0_) reduces to SKAT. As expected, the power of *T*_0_ remained largely similar across different local genetic correlation values, since the gene expression study is not informative for gene-trait association analysis in GWAS when using *T*_0_ as the test statistic. In addition, *T*_0_ achieved the highest power among the other individual tests when the underlying local genetic correlation *r* = 0. For example, in the baseline simulation settings with moderate *PVE* parameters, the power of *T*_0_ was 0.33, 0.32, 0.34, 0.33, 0.33, 0.31, and 0.30 when *r* = -1.0, -0.8, -0.5, 0.0, 0.5, 0.8, and 1.0, respectively. In addition, when *r* = 0, the power of *T*_*w*_ was 0.33, 0.33, 0.33, 0.33, 0.33, 0.32, 0.31, 0.28, 0.23, 0.17, and 0.087 when *w* = 0.0, 0.1, 0.2, 0.3, 0.4, 0.5, 0.6, 0.7, 0.8, 0.9, and 1.0, respectively, where the highest power was achieved by using small *w*. When *w* = 1, the test (*T*_1_) reduces to the TWAS test. As expected, the power of *T*_1_ was the lowest when *r* = 0 but kept increasing as the magnitude of *r* increased and achieved the highest power among all the individual tests when *r* = ±1. For example, in the baseline simulation settings with moderate PVE parameters, the power of *T*_1_ was 0.63, 0.43, 0.21, 0.087, 0.19, 0.41, and 0.61 when *r* = -1.0, -0.8, -0.5, 0.0, 0.5, 0.8, and 1.0, respectively. In addition, when *r* = 1, the power of *T*_*w*_ was 0.30, 0.33, 0.35, 0.38, 0.41, 0.45, 0.49, 0.54, 0.57, 0.61, and 0.61 when *w* = 0.0, 0.1, 0.2, 0.3, 0.4, 0.5, 0.6, 0.7, 0.8, 0.9, and 1.0, respectively, where the highest power was achieved by using *w* = 0.9 and 1. Finally, by aggregating p-values from all individual tests, VINTAGE achieved robust power performance across a wide range of local genetic correlations (Figure 3B).

**Figure 2.**
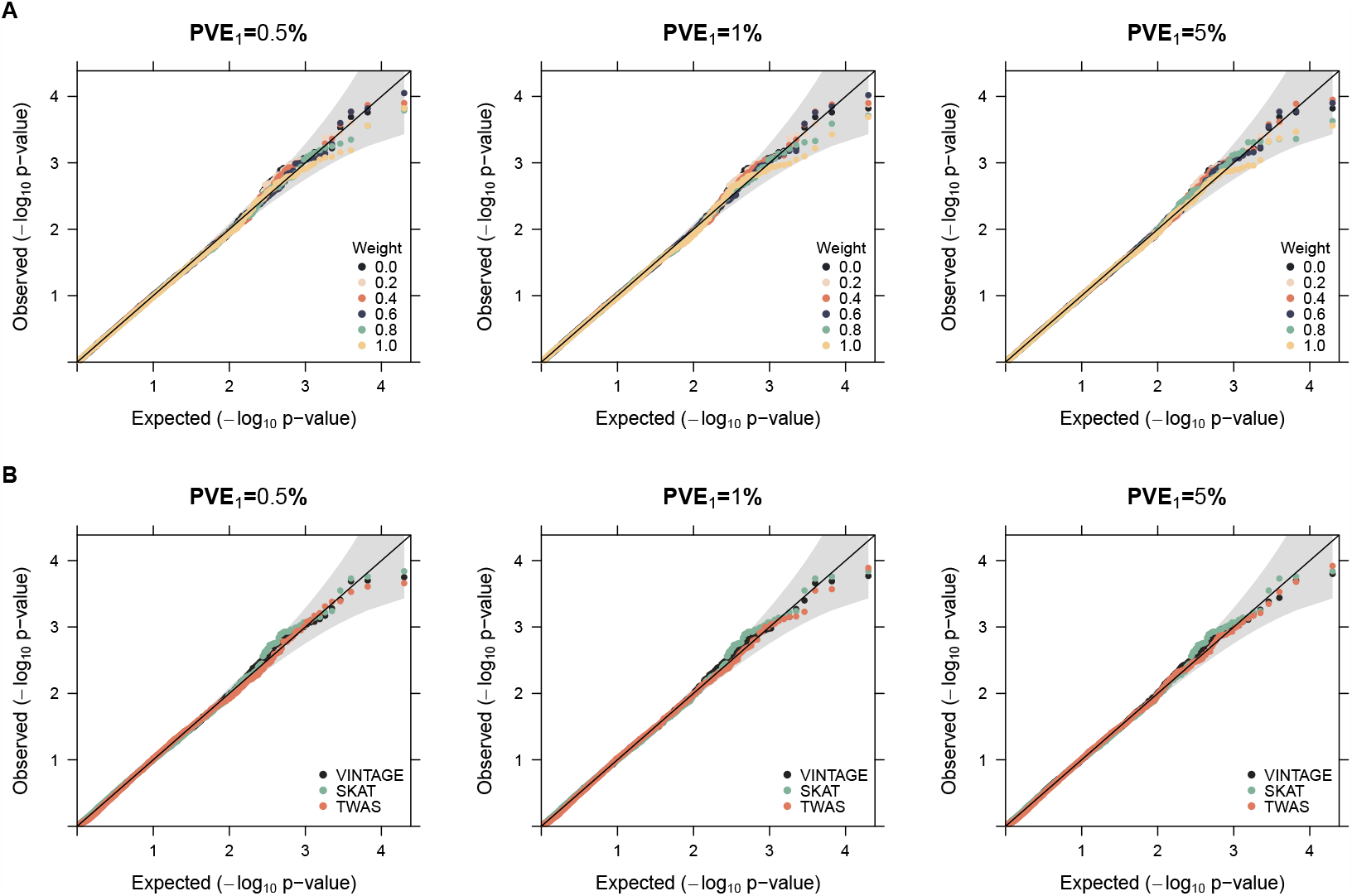
Quantile-quantile plots of -log_10_ p-values from different methods for gene-wise genetic variance test under the null simulations. Evaluated methods include: (**A**) individual tests from VINTAGE, *T*_*w*_, each constructed under a different weight value of *w* that ranges from 0 to 1 with an increment of 0.2, where the weight controls the relative contributions from the variance and covariance component scores; (**B**) the gene-wise genetic variance test of VINTAGE, which aggregates p-values from individual tests into a single p-value using the Cauchy combination approach; SKAT with an unweighted linear kernel; and TWAS that adapts multiple PRS methods to estimate gene expression prediction weights and performs an omnibus test. Null simulations were conducted under different *PVE*_1_ values (0.5%, 1%, and 5%) with in-sample LD matrices and a polygenic genetic architecture (*q* = *p*).

**Figure 3.**
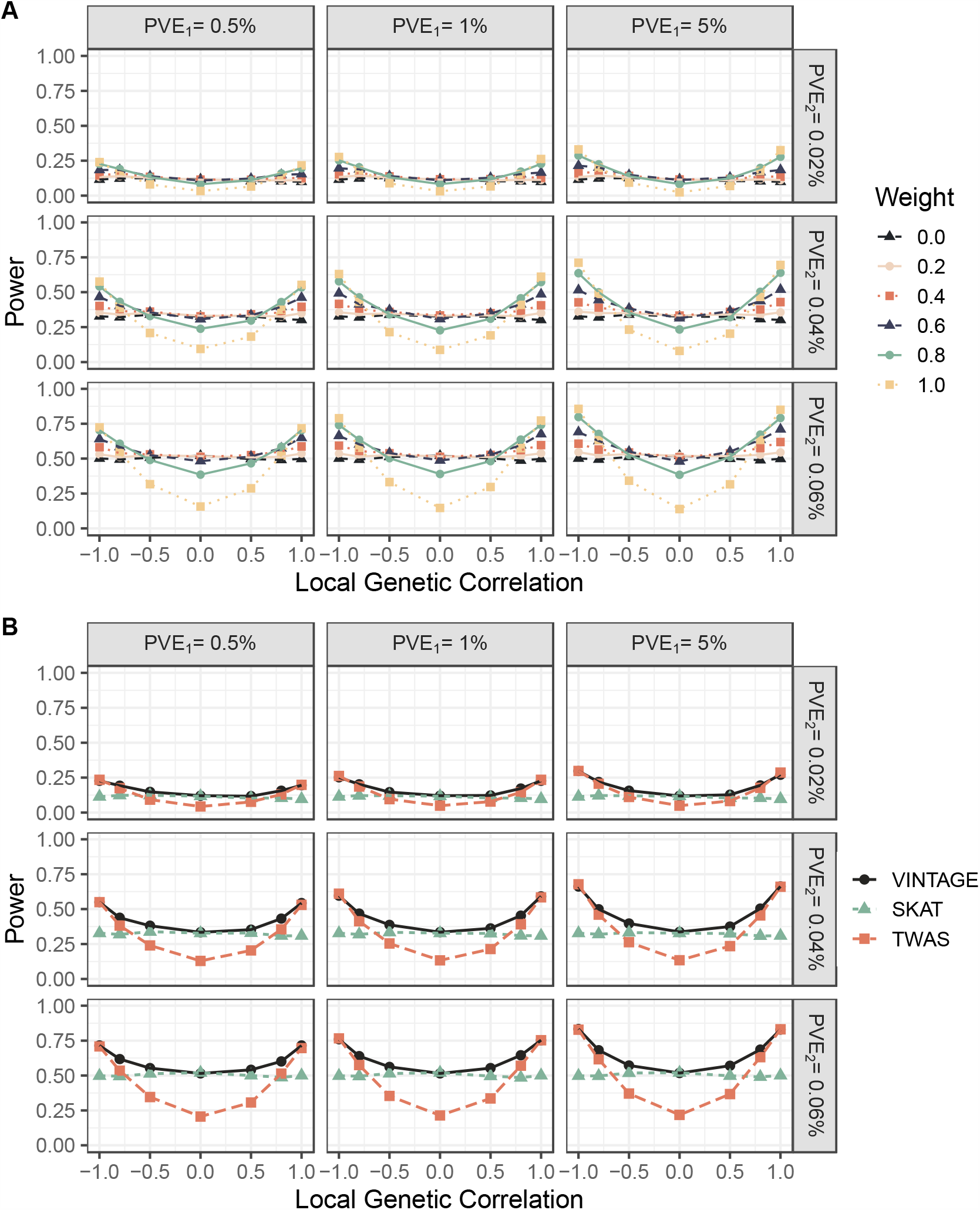
Power of different methods for gene-wise genetic variance test in the simulations. Compared methods include: (**A**) individual tests from VINTAGE, *T*_*w*_, each constructed under a different weight value of *w* that ranges from 0 to 1 with an increment of 0.2, where the weight controls the relative contributions from the variance and covariance component scores; (**B**) the gene-wise genetic variance test of VINTAGE, which aggregates p-values from all individual tests into a single p-value using the Cauchy combination approach; SKAT with an unweighted linear kernel; and TWAS that adapts multiple PRS methods to estimate gene expression prediction weights and performs an omnibus test. Simulations were conducted under different *PVE*_1_ values (0.5%, 1%, and 5%), different *PVE*_2_ values (0.02%, 0.04%, and 0.06%), and different local genetic correlations (-1.0, -0.8, -0.5, 0.0, 0.5, 0.8, 1.0) with in-sample LD matrices and a polygenic genetic architecture (*q* = *p*).

### Main simulation studies

We conducted extensive and realistic simulations to evaluate the performance of VINTAGE and compared it with existing approaches. Simulation details are provided in the Materials and methods. For the gene-wise genetic variance test of VINTAGE, which aims to detect gene-trait associations, we compared it with two representative methods, SKAT^13^ and TWAS^8^, that can also make use of summary statistics as input. For the gene-wise genetic correlation test of VINTAGE, which aims to identify genes whose expression may potentially mediate genetic effects on the trait, we compared it with SUPERGNOVA^14^ and LAVA^15^. Detailed descriptions of the compared methods are provided in the Materials and methods. In total, we explored 375 (60 nulls and 315 alternatives) simulation settings by varying the number of SNPs with non-zero effects on gene expression and the trait (*q*), their effect sizes on gene expression (*PVE*_1_) and the trait (*PVE*_2_), the local genetic correlation between gene expression and the trait (*r*), and the used linkage disequilibrium (LD) reference panels.

For the gene-wise genetic variance test, we first examined the type I error control of the three methods (VINTAGE, SKAT, and TWAS) under the null. We found that all three methods consistently produced well-calibrated p-values, irrespective of the SNP effects underlying gene expression (Figure 2B) or the nature of the genetic architecture underlying gene expression, whether sparse or polygenic (Figure S3). When external LD matrices were used, TWAS produced slightly inflated p-values, especially when the cis-SNP heritability underlying gene expression was relatively high (*PVE*_1_ = 5%; Figure S4). In contrast, the p-values from VINTAGE and SKAT remained largely calibrated (Figure S4).

Next, we evaluated the power of the three methods on detecting gene-trait associations and examined individual factors that influence their power. As expected, the power of SKAT and TWAS were consistent with the power of the two individual tests in VINTAGE, *T*_0_ and *T*_1_, constructed with the weights *w* = 0 and 1, respectively (Figure 3B). By aggregating p-values from all individual tests, VINTAGE exhibited superior power compared to SKAT and TWAS across the majority of local genetic correlation values. Only in the extreme settings of *r* = 0 or ±1, which perfectly aligned with the modeling assumptions made in SKAT and TWAS, respectively, did VINTAGE show a slight reduction in power compared to the two methods (Figure 3B). Also as expected, the power of all methods increased with increasing genetic effects on the trait (Figure 3B). The power of VINTAGE and TWAS also increased with increasing genetic effects on gene expression, while the power of SKAT remained largely similar as SKAT did not use information from the gene expression data (Figure 3B).

In addition, the power of different methods was influenced by the genetic architecture underlying gene expression and the trait, represented by the number of SNPs with non-zero effects on both gene expression and the trait, *q* (Figures S5-S7). For these simulations under sparse genetic architecture, an important complication arises from the discrepancy between the simulation parameter 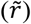 and the true local genetic correlation (*r*) (details discussed in the Supplementary Text). Consequently, regardless of the value of 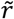, *r* is always equal to ±1 when *q* = 1 or enriched around these two values when *q* = 2 or 10. Indeed, the local genetic correlation was estimated close to either -1 or +1 under sparse settings (Figures S8-S10). As a result, the power of both VINTAGE and TWAS remained largely similar regardless of the value of 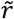 used in the simulations. For example, in the scenario with *q* = 1 and with moderate *PVE* parameters, the power of VINTAGE was 0.39, 0.37, 0.35, 0.37, 0.37, 0.37, and 0.40, and the power of TWAS was 0.43, 0.40, 0.36, 0.36, 0.37, 0.40, and 0.43 when 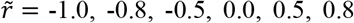, and 1.0, respectively (Figure S6). On the other hand, the power of SKAT increased as *q* increased owing to its polygenic assumption made about the SNP effects on the trait.

Importantly, we found that the power of TWAS exceeded the expected type I error rate of 0.00025% under the scenario with no local genetic correlations (*r* = 0; Figure 3B). This result is undesirable because TWAS is specifically designed to assess whether there is an association between gene expression and the trait. Consequently, TWAS is not expected to show power exceeding the type I error rate when the local genetic correlation is zero, which represents a null scenario for TWAS where gene expression is not mediating the genetic effects on the trait. Moreover, in the absence of local genetic correlation, the power of TWAS unfortunately continued to rise as the genetic effect size on the trait (*PVE*_2_) increased (Figure 3B). For example, in the scenario with a moderate cis-SNP heritability of gene expression (*PVE*_1_ = 1%) and no local genetic correlation (*r* = 0), the power of TWAS was 0.049, 0.13, and 0.21 when *PVE*_2_ = 0.02%, 0.04%, and 0.06%, respectively. This observation aligns with recent discussions regarding the “inflation “ of TWAS^16-18^, where TWAS has been reported to display inflated p-values when exploring the genetic relationship between gene expression and trait. Such “inflation “ is resolved and no longer observed in the genetic correlation test of VINTAGE. This observation thus highlights the importance of segregating the test that assesses gene-trait association from the test that assesses expression mediation, as is done in VINTAGE.

For the gene-wise genetic correlation test, we compared VINTAGE with SUPERGNOVA and LAVA. First, we examined the type I error control of the three methods under the null (Figure S11). We found that VINTAGE effectively controlled for type I error and produced well-calibrated p-values when both *PVE*_1_ and *PVE*_2_ were large. The VINTAGE p-values were on the conservative side when either *PVE*_1_ or *PVE*_2_ was small, as the local genetic correlation became hard to estimate. In contrast, SUPERGNOVA consistently produced deflated p-values across all *PVE*_1_ or *PVE*_2_ values. This issue likely arises because SUPERGNOVA cannot accurately estimate the phenotypic correlation parameter between gene expression and the trait in a local region with a limited number of SNPs, as opposed to the original application scenario where the parameter is estimated between two traits across the entire genome and shared by each local region. On the other hand, LAVA consistently produced highly inflated p-values across all *PVE*_1_ or *PVE*_2_ values, resulting in type I error rates ranging from 20%-45% across all simulation settings, substantially exceeding the expected rate of 5%. The observations on type I error control of LAVA are consistent with a previous benchmarking study on local genetic correlation methods^19^.

Next, we evaluated the power of the three methods on detecting non-zero local genetic correlations (Figure 4). Because LAVA was highly inflated under the null, we first focused on power comparison of VINTAGE with SUPERGNOVA at the nominal p-value threshold (Figure 4A). As expected, the power of both methods increased as the local genetic correlation increased. Additionally, VINTAGE exhibited greater power than SUPERGNOVA across a range of local genetic correlations. Besides the issue with estimating the phenotypic correlation parameter, the lower power of SUPERGNOVA is also likely attributed to its reliance on the relatively inefficient method of moments (MoM) inference algorithm, which was initially developed for testing broader genomic regions with a large number of SNPs. For example, in the baseline simulation settings with moderate *PVE* parameters, the power of VINTAGE was 0.79, 0.54, 0.23, 0.030, 0.22, 0.55, and 0.78, while the power of SUPERGNOVA was 0.14, 0.098, 0.044, 0.029, 0.046, 0.085, and 0.13, when *r* = -1.0, -0.8, -0.5, 0.0, 0.5, 0.8, and 1.0, respectively. As expected, the power of VINTAGE increased with higher values of either *PVE*_1_ or *PVE*_2_ . However, the power of SUPERGNOVA increased with higher values of *PVE*_2_ but decreased with higher values of *PVE*_1_. This observation is likely due to an inaccurate approximation in calculating the empirical variance of the local genetic covariance, which is a key component in constructing the test statistic of SUPERGNOVA. Because LAVA was highly inflated under the null, we also compared the power of the three methods across different false discovery rates (FDRs; Figure 4B). We found that VINTAGE was more powerful than the other two methods across a range of local genetic correlations and FDRs, with its power performance followed by LAVA and SUPERGNOVA. For example, in the same simulation setting described above, the power of VINTAGE was 0.88, 0.62, 0.081, 0.086, 0.63, and 0.89, while the power of LAVA was 0.65, 0.16, 0.0095, 0.0010, 0.17, and 0.62, and the power of SUPERGNOVA was 0.045, 0.0021, 0.0010, 0.012, 0.015, and 0.062 at an FDR of 50%, when *r* = -1.0, -0.8, -0.5, 0.5, 0.8, and 1.0, respectively. In addition, we found that the behavior of all three methods remained largely similar when external LD matrices were used (Figures S15-S19).

**Figure 4.**
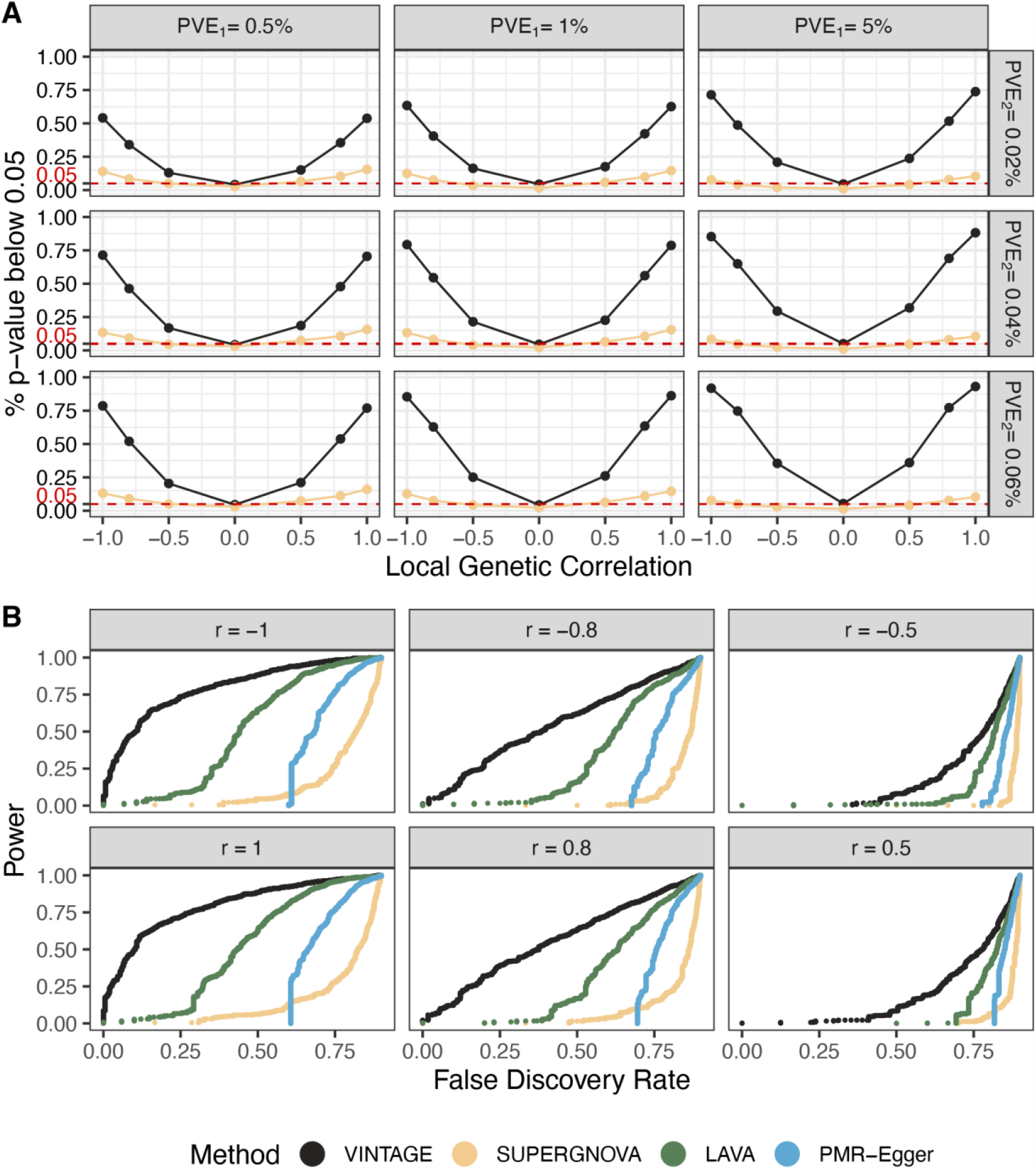
Evaluating the gene-wise genetic correlation test in the simulations. Compared methods include VINTAGE, SUPERGNOVA, LAVA, and PMR-Egger. (**A**) We used the proportion of simulation replicates that passed the p-value threshold of 0.05 to evaluate the type I error rate (*r* = 0) or power (*r* ≠ 0). Simulations were conducted under different *PVE*_1_ values (0.5%, 1%, and 5%), different *PVE*_2_ values (0.02%, 0.04%, and 0.06%), and different local genetic correlations (-1.0, -0.8, -0.5, 0.0, 0.5, 0.8, 1.0) with in-sample LD matrices and a polygenic genetic architecture (*q* = *p*). Horizontal dashed lines (red) indicate the expected type I error rate of 0.05. Simulation replicates with negative heritability estimates or negative variance estimates from SUPERGNOVA or LAVA were excluded from the evaluation. (**B**) We evaluated the power across different false discovery rates in settings with moderate *PVE* parameters (i.e., *PVE*_1_ = 1% and *PVE*_2_ = 0.04%), different local genetic correlations (-1.0, -0.8, -0.5, 0.5, 0.8, 1.0), in-sample LD matrices, and a polygenic genetic architecture. Power and false discovery rates were calculated by pairing simulation replicates under the alternative (1,000 replicates) with those under the null (9,000 replicates).

On top of the baseline simulation settings, we varied the number of cis-SNPs that have non-zero effects on both gene expression and the trait to be either 1, 2, or 10 to evaluate the performance of the gene-wise genetic correlation test under sparse genetic architectures. We evaluated type I error and power against the true genetic correlation calculated as the Pearson correlation coefficient between the simulated genetic effects on gene expression and on trait (details in Materials and methods; Figures S20-S22). In the analysis, we found that VINTAGE produced well-controlled type I error in simulation replicates where the true correlation was close to zero. In addition, the power of VINTAGE’s gene-wise genetic correlation test decreased as *q* decreased (Figures S20-S22). For example, in the scenario with moderate *PVE* parameters and with *r* in the range of 0.8 to 1.0, the power of VINTAGE was 0.70, 0.60, and 0.44 when *q* = 10, 2, and 1, respectively (Figure S21). However, despite the power decrease due to the sparse genetic architecture, VINTAGE still retained a non-negligible level of power for detecting non-zero local genetic correlations. In addition, we examined another null sparse genetic setting where two distinct sets of SNPs, each consisted of 1, 2, or 10 SNPs, displayed non-zero effects: one on gene expression and the other on the trait (details in Materials and methods). In this setting, VINTAGE also produced calibrated type I error control (Figures S12-S14).

Finally, both VINTAGE and LAVA produced approximately unbiased estimates for local genetic correlations while the estimates from SUPERGNOVA were biased towards zero (Figure S23). Among these three methods, VINTAGE produced the most accurate local genetic correlation estimates with much smaller variance compared to the other two methods. Importantly, the accuracy of the local genetic correlation estimates from different methods depends on the number of SNPs, *q*, that has non-zero effects on both gene expression and trait (Figures S8-S10). As *q* decreased, the local genetic correlation was estimated close to either -1 or +1 regardless of the value of the simulation parameter 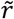 sdue the discrepancy between the simulation parameter and the true local genetic correlation (details discussed in the Supplementary Text). Lastly, the estimation results look largely similar when external LD matrices were used (Figure S24).

### Additional simulation studies

#### Additional method comparisons

In addition to the methods in the main simulation studies, we compared three SKAT and TWAS extensions – ACAT-O^20^, VC-TWAS^21^, and CoMM^22^ – for the gene-wise genetic variance test, and compared PMR-Egger^18^ for the gene-wise genetic correlation test. Detailed descriptions of the compared methods are provided in the Materials and methods section. For these comparisons, we focused on baseline simulation settings.

For the gene-wise genetic variance test, we first examined the type I error control of the three additional methods under the null. We found that all three methods consistently produced well-calibrated p-values, regardless of the SNP effects underlying gene expression (Figure S25). Next, we evaluated the power of these methods in detecting gene-trait associations (Figure S26). As expected, we found that the power of ACAT-O was largely similar to SKAT. Similar to TWAS, the power of VC-TWAS was the lowest when *r* = 0 but increased with the magnitude of *r*, achieving the highest power when *r* = ±1. However, the power of VC-TWAS was consistently lower than that of VINTAGE across different local genetic correlation values. In addition, the power of VC-TWAS fell between that of TWAS and SKAT. Compared to TWAS, VC-TWAS assumes random SNP effects on the trait, allowing it to achieve higher power when *r* is close to 0 but lower power when *r* is large (e.g., greater than 0.8). Compared to SKAT, VC-TWAS was more powerful when *r* was large, as it leverages additional information from the gene expression data; however, it was less powerful when *r* = 0 due to the extra parameter needed to model SNP effect variance on the trait. Finally, we found that the power of CoMM was comparable to VINTAGE when *PVE*_1_ was small and equaled 0.5%. However, as *PVE*_1_ increased to 1% and 5%, the power of CoMM became more similar to TWAS and less powerful than VINTAGE across the majority of local genetic correlation values. Only in the extreme settings when *r* = ±1, which aligned with the modeling assumption made by CoMM, did CoMM show comparable or slightly higher power than VINTAGE. This observation arises from the joint modeling nature of CoMM. Specifically, when *PVE*_1_ is large, CoMM estimates SNP effects on gene expression using expression data alone, resulting in performance similar to TWAS. However, as *PVE*_1_ decreases, CoMM borrows information from the GWAS data to estimate the SNP effects on gene expression. In the extreme case where *PVE*_1_ = 0, it directly uses SNP effect estimates on the trait as effect estimates on gene expression, yielding comparable power to SKAT and VINTAGE.

For the gene-wise genetic correlation test, consistent with the findings in the PMR-Egger paper, we observed that PMR-Egger produced highly inflated p-values across all *PVE*_1_ and *PVE*_2_ values examined under the null (Figures S11). For example, given a fixed p-value threshold of 0.05, the type I error rates of PMR-Egger ranged from 26%-73% across all simulation settings, substantially exceeding the expected rate of 5%. The p-value inflation is expected, as PMR-Egger relies on the Egger assumption to model horizontal pleiotropy, and it is thus only effective when the horizontal pleiotropy effects are relatively small or exhibit consistent signs, which is not the case examined here. Due to the inflation of PMR-Egger p-values under the null, to ensure fair comparison, we examined the power of all four methods across different FDRs (Figure 4B). We found that PMR-Egger consistently yielded lower power than VINTAGE and LAVA across all FDRs. Similar to SUPERGNOVA, PMR-Egger began to exhibit non-negligible power only at high FDR levels. For example, in the settings with moderate *PVE* parameters, the FDR of PMR-Egger was 76% and 66% to achieve the power of 0.5 when *r* = 0.8, and 1.0, respectively. In contrast, the FDR was 38% and 10% for VINTAGE, 63% and 44% for LAVA, and 86% and 82% for SUPERGNOVA to achieve the power of 0.5 when *r* = 0.8, and 1.0, respectively. These observations indicate that PMR-Egger was not effective in distinguishing the null cases from alternatives, again likely due to the Egger assumption being effective only when the horizontal pleiotropy effects are relatively small or exhibit consistent signs. Indeed, PMR-Egger becomes inadequate when horizontal pleiotropy effects are larger and signs vary across SNPs, as demonstrated in the original PMR paper – a scenario that aligns with our simulation settings.

#### Evaluating the influence of the sample size of the gene expression study

We conducted additional simulations to evaluate the performance of the gene-wise genetic correlation test for data with a sample size comparable to the Genotype-Tissue Expression (GTEx) study^3^ (n = 838). Simulation details are provided in the Materials and methods. Consistent with our main simulations, we found that VINTAGE p-values were on the conservative side, more so when the sample size of the gene expression study was small, as the local genetic correlation became harder to estimate (Figure S27A). On the other hand, SUPERGNOVA and LAVA failed frequently when the sample size of the gene expression study was small (Figure S27B). For example, SUPERGNOVA failed in 47% and 34% of the simulation replicates when the sample size of the gene expression study was 500 and 1,000, respectively, while LAVA failed 31% and 18%. The high failure rate for the two methods is likely due to the inherent limitation of MoM-based algorithms used by both methods, which can produce negative heritability estimates when the sample size is small or when the true heritability is close to zero. As expected, the power of all three methods decreased as the sample size of the gene expression study decreased (Figure S27C-D). For example, at the nominal p-value threshold of 0.05 and a local genetic correlation of 0.8, the power of VINTAGE was 0.56, 0.21, and 0.11 when the sample size of the gene expression study was 10,000, 1,000, and 500, respectively (Figure S27C). In contrast, the corresponding power of SUPERGNOVA was 0.11, 0.10, and 0.08 (Figure S27C). Note that we did not evaluate the power of LAVA at a fixed p-value threshold because it was highly inflated under the null. Instead, we further evaluated the power of different methods across a range of FDRs to ensure fair comparison. At an FDR of 50% and a local genetic correlation of 0.8, the corresponding power of VINTAGE was 0.63, 0.14, and 0.021, while the power of LAVA was 0.17, 0.10, and 0.053, and the power of SUPERGNOVA was 0.015, 0.0098, and 0.0030 (Figure S27D). These observations highlight the importance of having a large sample size in the gene expression study to achieve a powerful gene-wise genetic correlation test.

#### Evaluating the influence of inconsistent genetic architectures

We conducted additional simulations to evaluate the performance of both the gene-wise genetic variance and correlation tests under settings where the genetic architecture is sparse in one study, while it is polygenic in the other. Simulation details are provided in the Materials and methods.

For the gene-wise genetic variance test, we first evaluated the type I error control of VINTAGE, SKAT, and TWAS under the null. For this evaluation, we did not further explore the null settings where the genetic architecture underlying gene expression is sparse, as those are essentially the same as the previous simulations with a sparse genetic architecture (Figure S3). Instead, we focused on settings where the underlying genetic architecture is polygenic and a sparse set of *q* cis-SNPs explained a large proportion of variance in gene expression as quantified by 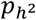, among all cis-SNPs in the gene region. In the simulations, we found that all three methods produced well-calibrated p-values regardless of the value of *q* and the value of 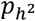 (Figure S28). Next, we evaluated the power of these three methods. In addition to our previous findings that the power of both VINTAGE and TWAS was less influenced by the local genetic correlation as *q* decreased when the genetic architecture underlying both gene expression and the trait was sparse (Figures S5-S7), we also found that the power of VINTAGE and TWAS was less influenced by the local genetic correlation as 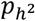 decreased (Figure S29). For example, in scenario I with *q* = 10, the power of VINTAGE ranged from 0.36 to 0.52 when 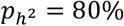, while it ranged from 0.35 to 0.38 when 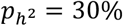 across all local genetic correlation values (Figure S29A). Similarly, the power of TWAS ranged from 0.19 to 0.52 when 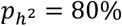, while it ranged from 0.15 to 0.27 when 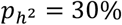 across all local genetic correlation values (Figure S29A). This is expected because as 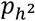 decreases, the local genetic correlation across the entire gene region approaches zero regardless of how strong the correlation is among the sparse set of *q* SNPs. In contrast, the power of SKAT remained largely similar in scenario I where the genetic architecture underlying the trait is polygenic (Figure S29A) and increased as *q* increased in scenario II where the genetic architecture underlying the trait is sparse (Figure S29B). This is consistent with our observations in the main simulation studies as SKAT does not use information from the gene expression data (Figures 3B and S5-S7).

For the gene-wise genetic correlation test, as in the main simulations, we evaluated type I error and power against the true genetic correlation calculated as the Pearson correlation coefficient between the simulated genetic effects on gene expression and on trait (details in Materials and methods; Figures S30). We did not evaluate LAVA due to its inflated type I errors (Figures S11-S14) and did not evaluate SUPERGNOVA as it was underpowered (Figure 4). In the simulations, we found that VINTAGE produced well-controlled type I error in simulation replicates where the true correlation was close to zero. In addition, the power of VINTAGE decreased as 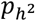 increased (Figure S30). For example, in scenario I with *q* = 2 and with *r* in the range of 0.6 to 0.8, the power of VINTAGE was 0.57, 0.46, and 0.31 when 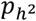 was 30%, 50%, and 80%, respectively. Presumably, this is because after accounting for the true local genetic correlation, the power of the gene-wise genetic correlation test increases when either gene expression or trait has a genetic architecture that is more polygenic (i.e., smaller 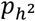).

#### Evaluating genetic correlation test accounting for the influence of the genetic variance test

We conducted additional simulations where we took into account the influence of the genetic variance test to evaluate the performance of the genetic correlation test. Simulation details are provided in the Materials and methods. We first examined the type I error control of both procedures. We found that, after accounting for the genetic variance test, the genetic correlation test produced more calibrated p-values across all combinations of *PVE*_1_ and *PVE*_2_ parameters (Figure S31). This is expected because, when coupled with the genetic variance test, simulation replicates with genetic variances close to zero would be excluded from further testing for local genetic correlations – these are the cases where the local genetic correlation parameter is less identifiable and including them would lead to conservative/deflated p-values for the local genetic correlation test. Next, we evaluated the power of both procedures. Under the nominal p-value threshold, we found that the procedure ignoring the genetic variance test was more powerful than the procedure that took it into account (Figure S32). This is expected, as bypassing the genetic variance test allows for a larger set of genes to be tested for genetic correlation, which, when paired with a nominal p-value threshold that does not account for multiple testing, would lead to higher power (along with increased family-wise error rate). On the other hand, under the Bonferroni corrected p-value threshold, although the power of both procedures tended to be low due to the lack of power in local genetic correlation analysis, the procedure that accounted for the genetic variance test was slightly more powerful than the procedure that did not (Figure S33). This is also expected, as the procedure that accounts for the genetic variance test narrows the focus to a smaller set of genes that are more likely to exhibit appreciable local genetic correlation, thereby improving power at the same family-wise error rate. The Bonferroni corrected threshold provides a fairer comparison between the two procedures, as it accounts for the different number of genes tested for local genetic correlation in each procedure.

#### Evaluating settings where the two studies have distinct sets of non-zero effect SNPs

We conducted similar simulations where we used two sets of SNPs, with 1, 2, or 10 SNPs in each set: one set with non-zero effects on gene expression and the other set with non-zero effects on the trait, to evaluate the performance of the gene-wise genetic variance test. The two sets of SNPs were distinct and did not exhibit extremely high LD (R^2^ < 0.95) with each other. Simulation details are provided in the Materials and methods. The results for the gene-wise genetic variance test closely mirrored those for the gene-wise genetic correlation test described in the main simulations. We found that the power of all three methods -- VINTAGE, SKAT, and TWAS -- remained largely similar to the baseline simulations, where all cis-SNPs had non-zero effects on both gene expression and the trait and where the local genetic correlation was zero (Figure S35). This is because in both scenarios, including the baseline scenario and sparse scenario, the gene expression data do not provide any additional information to inform the gene-trait associations. The power of all methods was slightly lower when *q* = 1, likely due to the polygenic assumption inherent in these methods (Figure S35).

### Real data applications

We applied VINTAGE and other approaches for gene-based analysis on 18 quantitative traits in the UKBB. We analyzed 13,725 protein-coding genes using summary statistics from UKBB and eQTLGen phase I study^25^, along with LD matrices estimated from the reference 1000G^26^ (details in the Materials and methods).

We first performed gene-wise genetic variance tests using VINTAGE, SKAT, and TWAS. Consistent with its high power in simulations, VINTAGE identified a larger number of associated genes across traits compared to SKAT and TWAS (Figure 5A and Table 1). Specifically, VINTAGE identified a total of 25,166 associated genes across all traits, which is 5% and 87% more than those identified by SKAT (23,987) and TWAS (13,435), respectively. Because the local genetic correlations between gene expression and the trait were close to zero for most identified genes (Figure S36), TWAS was not well positioned to incorporate information from the gene expression study for gene association analysis. Consequently, TWAS identified substantially fewer genes than VINTAGE and SKAT, consistent with the simulation settings where the local genetic correlation was low (Figures 3B, 5A, and Table 1). In addition, because the modeling framework of VINTAGE includes both SKAT and TWAS as special cases, the majority of genes identified by SKAT (96%) and TWAS (92%) were also detected by VINTAGE (Figure 5B), with small variations across traits (Figure S37A). In contrast, 91% and 49% of the genes identified by VINTAGE were detected by SKAT and TWAS, respectively (Figure 5C and S37B). The number of identified genes varied substantially across traits and was correlated with trait heritability (correlation R^2^ = 0.69; Figure 5D, S38, and Table 1). Finally, consistent with the use of blood eQTL data, we found that the extra number of genes identified from VINTAGE relative to SKAT was higher in blood-related traits than in the other traits (Figure S39A-B).

**Table 1.** Summary of the number of genes identified in the analysis of 18 traits from UKBB. For gene-wise genetic variance tests, compared methods include VINTAGE, SKAT with an unweighted linear kernel, and TWAS that adapts multiple PRS methods to estimate gene expression prediction weights and performs an omnibus test. For gene-wise genetic correlation tests, we only included results from VINTAGE as SUPERGNOVA was unable to identify any genes and LAVA failed to control for type I error.

| Trait | Abbreviation | Gene-wise Genetic Variance Test |  |  | Gene-wise Genetic Correlation Test |
| --- | --- | --- | --- | --- | --- |
|  |  | VINTAGE | SKAT | TWAS | VINTAGE |
| Age at menarche | AM | 410 | 389 | 208 | 0 |
| Bone mineral density | BMD | 829 | 790 | 394 | 0 |
| Body mass index | BMI | 1,327 | 1,259 | 658 | 2 |
| Basal metabolic rate | BMR | 2,167 | 2,075 | 1,133 | 2 |
| Diastolic blood pressure | DBP | 766 | 708 | 404 | 1 |
| Eosinophils count | EOS | 1,500 | 1,403 | 828 | 6 |
| Forced expiratory volume | FEV1 | 1,096 | 1,021 | 572 | 4 |
| Forced vital capacity | FVC | 1,353 | 1,277 | 674 | 1 |
| Hip circumference | HC | 1,347 | 1,256 | 704 | 1 |
| Neuroticism score | NS | 185 | 185 | 114 | 0 |
| Platelet count | PLT | 2,596 | 2,514 | 1,424 | 7 |
| Red blood cell count | RBC | 1,953 | 1,853 | 1,085 | 5 |
| RBC distribution width | RDW | 1,968 | 1,883 | 1,148 | 11 |
| Systolic blood pressure | SBP | 818 | 768 | 400 | 1 |
| Standing height | SH | 3,957 | 3,850 | 2,104 | 6 |
| White blood cell count | WBC | 1,793 | 1,723 | 992 | 13 |
| Waist circumference | WC | 966 | 914 | 502 | 1 |
| Years of education | YE | 135 | 119 | 91 | 0 |

**Figure 5.**
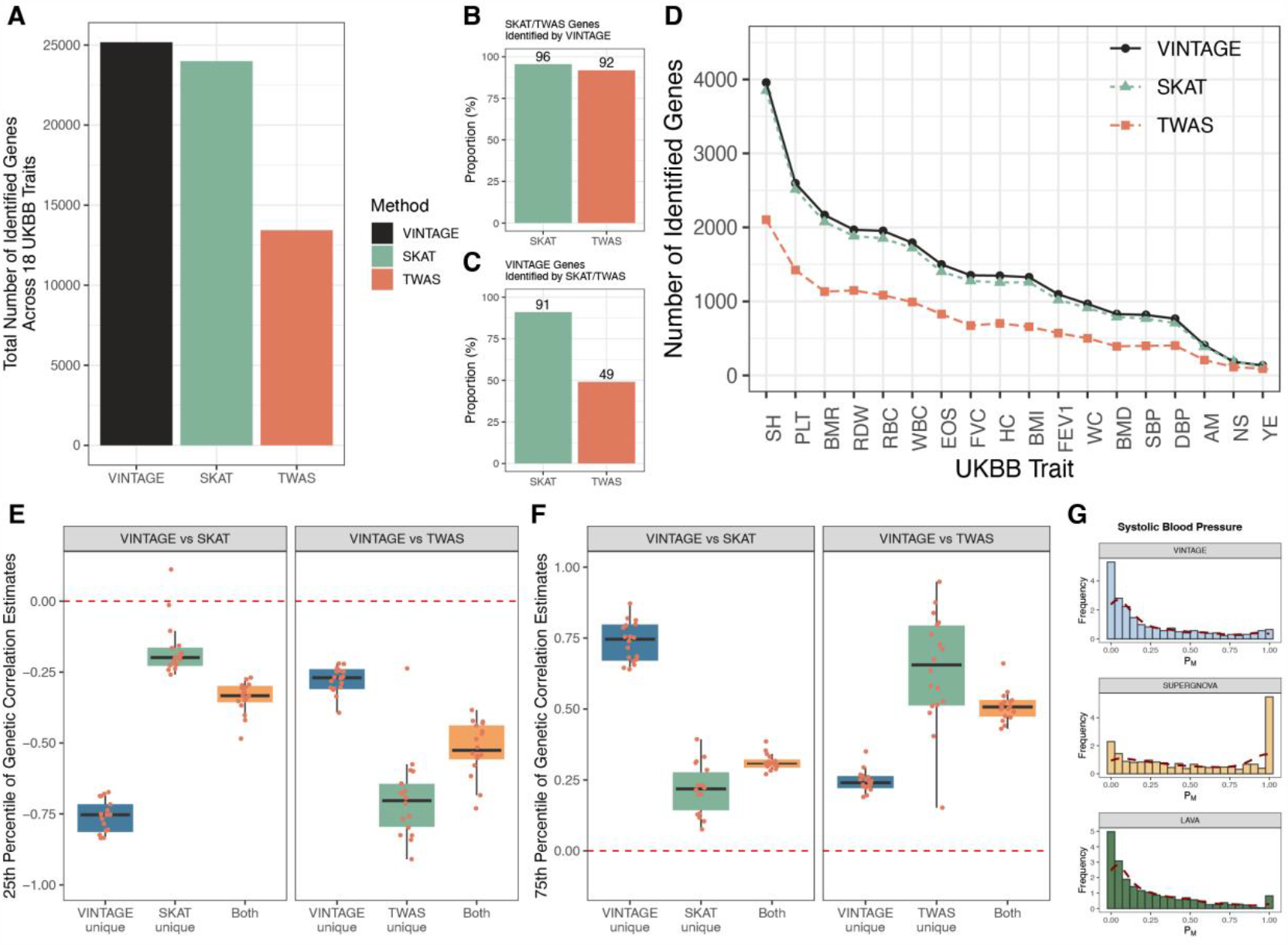
Gene-based analysis of 18 quantitative traits in UKBB by different methods. For gene-wise genetic variance analysis, compared methods include VINTAGE, SKAT, and TWAS. For gene-wise genetic correlation analysis, compared methods include VINTAGE, SUPERGNOVA, and LAVA. (**A**) Total number of genes across all 18 traits identified by different methods in the gene-wise genetic variance tests. (**B**) Proportion of genes identified by SKAT or TWAS across all 18 traits that were also identified by VINTAGE. (**C**) Proportion of genes identified by VINTAGE across all 18 traits that were also identified by SKAT or TWAS. (**D**) Number of genes identified by different genetic variance testing methods per trait. (**E**) 25^th^ and (**F**) 75^th^ percentile of the estimated local genetic correlations among VINTAGE uniquely identified genes comparing to SKAT, SKAT uniquely identified genes comparing to VINTAGE, genes identified by both VINTAGE and SKAT, VINTAGE uniquely identified genes comparing to TWAS, TWAS uniquely identified genes comparing to VINTAGE, or genes identified by both VINTAGE and TWAS. The length of the box corresponds to the interquartile range, with the center line and value corresponding to the median, and the upper and lower whiskers represent the largest or lowest value no further than 1.5 times the interquartile range from the third and first quartiles, respectively. (**G**) Histograms showing the distribution of the squared local genetic correlations estimated by VINTAGE, SUPERGNOVA, or LAVA between genes that passed the gene-wise genetic variance tests and systolic blood pressure. For both SUPERGNOVA and LAVA, the local genetic correlation estimates were constrained to -1 or 1 if they fell below -1 or above 1, respectively. The 18 analyzed quantitative traits include age at menarche (AM), bone mineral density (BMD), body mass index (BMI), basal metabolic rate (BMR), diastolic blood pressure (DBP), eosinophils count (EOS), forced expiratory volume (FEV1), forced vital capacity (FVC), hip circumference (HC), neuroticism score (NS), platelet count (PLT), red blood cell count (RBC), RBC distribution width (RDW), systolic blood pressure (SBP), standing height (SH), white blood cell count (WBC), waist circumference (WC), and years of education (YE).

We carefully compared the genes detected by the three methods by dividing them into six categories, including two categories of genes detected by two methods and four categories of genes detected by only one method (Figures 5E-F and S40-S41). As expected, the estimated local genetic correlation varied across gene categories, and such variation can explain the power of different methods in detecting different categories of genes. Specifically, when comparing VINTAGE with SKAT, the estimated local genetic correlations were closer to 0 in the set of SKAT uniquely identified genes while closer to -1 (Figure 5E) or +1 (Figure 5F) in the set of VINTAGE uniquely identified genes. The genes identified by both VINTAGE and SKAT had intermediate local genetic correlation estimates compared to the other two gene categories (Figures 5E-F). Similarly, when comparing VINTAGE with TWAS, the estimated local genetic correlations were closer to - 1 (Figure 5E) or +1 (Figure 5F) in the set of TWAS uniquely identified genes while closer to 0 in the set of VINTAGE uniquely identified genes. These results reinforce the conclusion that the power of different methods is highly influenced by local genetic correlations.

For genes detected by at least one of the three methods, we further conducted gene-wise genetic correlation tests using VINTAGE and SUPERGNOVA. We did not apply LAVA for the test as it failed to control for type I error. In the analysis, we found that most genes did not exhibit detectable mediation effects, with the median proportion of potential mediation estimated at only 12% and the mean proportion at 24% (Figures 5G and S42). Importantly, VINTAGE stood out as the only effective method for the genetic correlation test and identified a total of 61 genes across all traits that exhibited a significant local genetic correlation between gene expression and the trait (Table S1). Consistent with the use of blood eQTL data, the number of genes identified from VINTAGE with significant local genetic correlation was higher in blood-related traits than in the other traits (Figure S39C-D). In contrast, SUPERGNOVA were unable to identify any genes, consistent with its very low power observed in the simulation studies. The 61 genes identified by VINTAGE in the gene-wise genetic correlation tests represented potential mediators that mediate the SNP effects on the trait through gene expression, with the estimated proportion of mediation ranged from 0.25 to 1.0 (mean = 0.79; median = 0.81). Certainly, some of these genes may be associated with the phenotype due to expression pleiotropy, that the SNP effects on the trait may be mediated by one of the neighboring genes (Figures S43-S44; Supplementary Text). Finally, we compared local genetic correlation estimates from VINTAGE, SUPERGNOVA, and LAVA (Figures S36, S45-S46). We found that the local genetic correlation estimates from VINTAGE and LAVA both distributed in a bell shape around zero and were largely consistent with each other, with an average correlation R^2^ of 0.74 across traits (Figure S36, S46-S47). This aligns with our observation in the simulation studies, where both VINTAGE and LAVA produced approximately unbiased estimates with relatively small variance for local genetic correlations (Figure S23). On the other hand, the estimates from SUPERGNOVA were uniformly distributed with two peaks at the boundaries of -1 and +1, likely reflecting the considerable variance of those estimates produced by SUPERGNOVA (Figure S45).

Importantly, the combination of the gene-wise genetic variance and correlation tests in VINTAGE allowed us to interpret the gene-trait association results. We list two gene examples here. The first example, *CAMK1D* (Calcium/Calmodulin Dependent Protein Kinase 1D), was identified by VINTAGE with a significant association with systolic blood pressure (SBP; VINTAGE pg = 1 × 10^−6^; SKAT p = 0.015; TWAS p = 4 × 10^−10^) and a significant local genetic correlation (VINTAGE p_r_ = 2.8 × 10^−5^; SUPERGNOVA p = 0.63). *CAMK1D* encodes a protein kinase critical for regulating the synthesis of aldosterone, a steroid hormone responsible for maintaining sodium homeostasis, blood volume, and blood pressure^27,28^. The estimated local genetic correlation between *CAMK1D* and SBP was 0.81, corresponding to a relatively strong potential mediation effect of gene expression, with a proportion mediated *p*_*M*_ = 65%. The top SBP associated SNPs (e.g., rs61848342: p = 8.4 × 10^−7^) were also top eQTLs (e.g., rs61848342: p = 1.1 × 10^−308^) located upstream of *CAMK1D* TSS (Figure 6A). Analysis of a promoter capture Hi-C dataset revealed an interaction between this region and the promoter of *CAMK1D*^*29*^. All this evidence suggests that a large proportion of the SNP effects on SBP may be mediated through *CAMK1D* expression. Some SBP associated SNPs located in the genic region of *CAMK1D* (e.g. rs76987186: p = 2.1 × 10^−6^) were not eQTLs (e.g. rs76987186: p = 0.038; Figure 6A), suggesting that a fraction of the SNP effects on SBP may also be mediated through other pathways. As another example, *FURIN* (Furin, Paired Basic Amino Acid Cleaving Enzyme) was identified by VINTAGE with a significant association with SBP (VINTAGE p_g_ = 1.0 × 10^−6^; SKAT p = 2.0 × 10^−13^; TWAS p = 4.9 × 10^−4^) but with no significant local genetic correlation (VINTAGE p_r_ = 0.77; SUPERGNOVA p = 0.87). *FURIN* encodes a proprotein convertase enzyme that activates renal sodium transporter and key components in the renin-angiotensin-aldosterone system (RAAS) to regulate blood pressure^30-32^. Several SNPs in *FURIN* have been shown to be associated with SBP in previous GWASs^33,34^. For example, rs2071410 is located in the genic region of *FURIN* and showed strong association evidence with SBP in both UKBB (p =3.3 × 10^−30^; Figure 6B) and another GWAS (p = 2.9 × 10^−12^)^34^. However, rs2071410 was only weakly associated with *FURIN* expression in the eQTLGen study (p = 9.9 × 10^−7^; Figure 6B). In addition, *FURIN* eQTLs, primarily located in the upstream region of *FURIN*, displayed only weak association evidence with SBP in UKBB (Figure 6B). For example, the p-value for the lead eQTL, rs372098, was 1.7 × 10^−94^ in the eQTLGen study but was only 0.15 for SBP in UKBB. The estimated local genetic correlation between *FURIN* and SBP was only 0.066, corresponding to a very small potential mediation effect of gene expression, with a proportion mediated *p*_*M*_ = 0.43%. All this evidence indicates that SNPs in the genic and adjacent regulatory regions of *FURIN* likely influence SBP through mechanisms other than the expression changes in *FURIN*.

**Figure 6.**
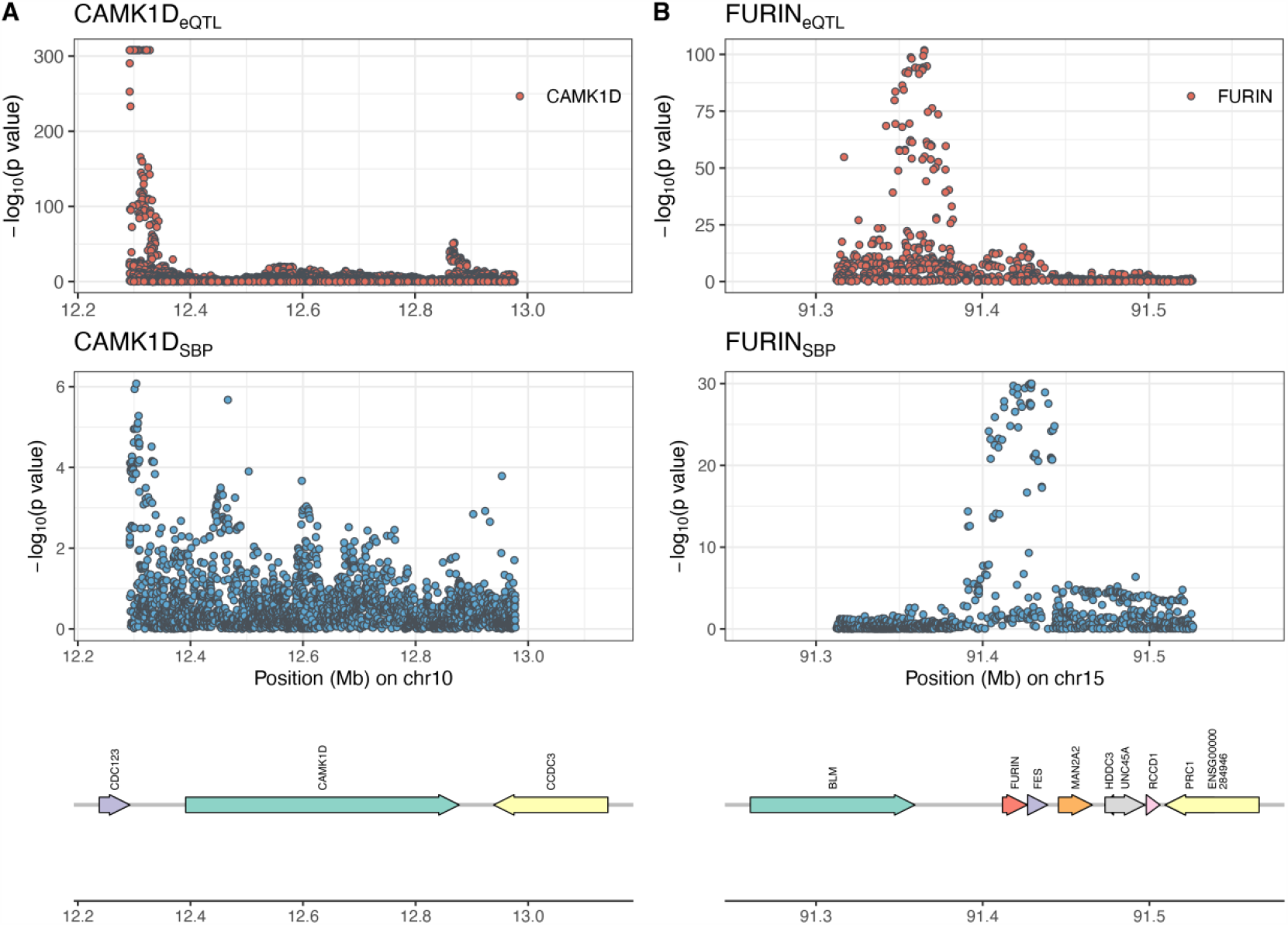
P-values for SNPs from the eQTL study and GWAS are plotted against their physical positions in the susceptibility gene regions identified by VINTAGE. -log_10_ p-values are shown for SNPs in the eQTLGen study (top panels) and in the UKBB GWAS (bottom panels). The two genes include (**A**) *CAMK1D* and (**B**) *FURIN*, which were identified to be significantly associated with systolic blood pressure by VINTAGE.

The results obtained from VINTAGE also allowed us to examine how the increase in the sample size of the gene expression mapping study (*n*_1_), which is expected in the near future, may help improve the power of the gene-wise genetic variance and correlation tests (analysis details in the Materials and methods). We observed that the power of gene-wise genetic variance tests could benefit from an increasing sample size of the gene expression mapping study, especially for genes with large local genetic correlations (Figure S48A). In particular, for genes with effects on trait greater than 0.01% (i.e., 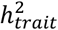 ≥ 0.01%), the power of the gene-wise genetic variance test was 0.58, 0.61, and 0.62 among all genes when *n*_1_ = 100, 20,000, and 361,194, respectively, and became 0.66, 0.74, and 0.77 among genes with |*r*| ≥ 0.8. Therefore, a substantial number of genes could benefit from incorporating a larger gene expression mapping study, as the number of genes with 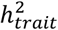 ≥ 0.01% ranged from 1,315 to 7,997 (mean = 3,945; median = 4,047) across the 18 analyzed traits, and among them, the number of genes with |*r*| ≥ 0.8 ranged from 239 to 568 (mean = 415; median = 417; Figure S48B). However, the benefits of increasing the sample size of the expression mapping study are limited for genes with small effects on the trait (i.e., 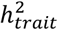∈ [0.0001%, 0.005%)), corresponding to 3,028 to 7,212 genes (mean = 5,572; median = 5,594) across traits regardless of the local genetic correlation (Figures S48A-B). In particular, among genes with 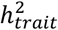 ∈ [0.001%, 0.005%) and with |*r*| ≥ 0.8, the power of the gene-wise genetic variance test increased from 0.03 to 0.06 as *n*_1_ increased from 100 to 361,194. While the relative power increase was 100%, greater than the 17% increase observed among genes with effects on the trait greater than 0.01%, the absolute increase was considerably smaller (3% compared to 11%). The limited power gain for genes with small effect sizes on the trait is likely due to the limited association information GWAS can provide.

In addition to the gene-wise genetic variance tests, we also evaluated how increasing the sample size of the gene expression mapping study may help improve the power of the gene-wise genetic correlation tests. We observed that, while the power of the gene-wise genetic correlation test increased with larger sample size of the gene expression mapping study, it remained largely under-powered (Figure S48C). For example, for genes with cis-SNP heritability of gene expression (i.e., 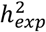) ≥ 1% and 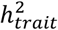≥ 0.01%, the power of the gene-wise genetic correlation test to identify genes with |*r*| ≥ 0.8 was only 6% and 7% when *n*_1_ = 20,000 and 361,194, respectively, under a Bonferroni corrected p-value threshold. The observation of an under-powered gene-wise genetic correlation test in the present real data application is consistent with the relatively small number of identified signals in other local genetic correlation studies^14,15,35^. Indeed, we identified a much smaller number of only six significant genes from the gene-wise genetic correlation test using the GTEx data (Table S2). These findings suggest that a much more powerful statistical approach, either within the framework of local genetic correlation test or based on another statistical framework, is likely needed for a comprehensive investigation of the mediation role of gene expression in a trait of interest.

Additionally, we applied MESuSiE to estimate the number of SNPs with non-zero effects on gene expression and the trait for the same 18 quantitative traits, focusing on genes that passed the gene-wise genetic variance test of VINTAGE. In the analysis, we found that the number of SNPs with non-zero effects on either expression or the trait is much higher for genes that passed the gene-wise genetic correlation test compared to those did not pass the test (Figure S49). Specifically, for genes that did not pass the gene-wise genetic correlation test, the average number of non-zero effect SNPs was estimated to be 5.3 (median = 4.0) for gene expression and 3.1 (median = 3) for traits. In contrast, for genes that passed the gene-wise genetic correlation test, the estimate was 12.1 (median = 12.0) for gene expression and 6.5 (median = 6.0) for traits. In addition, the average number of shared SNPs with non-zero effects on both gene expression and trait was estimated to be 2.5 (median = 2.0) and 6.2 (median = 6.0) for genes that did not pass and passed the gene-wise genetic correlation test, respectively. These results suggest that the small number of non-zero effect SNPs in the genic region contributes to the low power of the gene-wise genetic correlation test, which is consistent with our simulation results (Figures S20-S22).

Finally, VINTAGE is computationally efficient and requires at most 20 seconds to analyze a gene (Figure S50).

## Discussion

We have presented VINTAGE, a unified and powerful method for integrating GWAS with gene expression study to identify and decipher genes associated with a trait of interest. VINTAGE bridges SKAT and TWAS with the local genetic correlation parameter and includes both methods as special cases. By inferring the local genetic correlation parameter, VINTAGE is able to both test and quantify the proportion of genetic effects on the trait potentially mediated through gene expression and leverage such information to guide the integration of the gene expression mapping study towards gene association mapping in GWAS. We have illustrated the benefits of VINTAGE through both simulations and integrative analyses of a large-scale eQTL mapping study, eQTLGen, with eighteen complex traits from UKBB.

VINTAGE provides the statistical foundation that bridges SKAT and TWAS by introducing the local genetic correlation parameter. This foundation justifies the combination of the two methods towards the common analytic goal of identifying gene-trait associations. Given that the VINTAGE framework opens door for the integration of SKAT and TWAS, the natural follow-up question is to determine the best approach for this integration. We explored a total of five distinct approaches, including one successful approach that is presented in the main text and implemented in the default version of VINTAGE’s variance component test, along with four unsuccessful approaches that are presented in the Supplementary Text. Besides these explored approaches, one additional approach is to run SKAT and TWAS separately and then aggregates their p-values into a single p-value using the Cauchy combination approach. This procedure is closely related to a simplified version of VINTAGE, a version that uses only two weight values: 0, which corresponds to SKAT, and 1, which corresponds to TWAS. We have implemented this simple version into VINTAGE (termed as SKAT+TWAS) and compared it with standard VINTAGE in both simulations and real data applications. In the analysis, we found that VINTAGE’s genetic variance test remained more powerful than SKAT+TWAS, outperforming it by 1.4% to 15.4% (mean = 6.8%; median = 6.8%; Figure S51A) in simulations and 1.5% in real data applications, with 372 additional gene discoveries (Figure S51B). In particular, when *r* = 0 or ±1, the average power gain from VINTAGE relative to SKAT+TWAS was 4.4% across all settings, while for *r* = ±0.5 or ±0.8, the average power gain increased to 9.2% (Figure S51A). These observations highlight that a generalized construction of the genetic variance test is especially important for identifying genes with moderate local genetic correlations, which may play crucial roles in mediating the genetic effects on traits through gene expression regulation in real settings. Indeed, consistent with simulations, in the real data applications, we found that the advantage of VINTAGE over SKAT+TWAS was especially apparent when the estimated local genetic correlations were moderate (|*r*| ∈ [0.3,0,7)), leading to a 5.0% increase in the identified gene-trait associations (Figure S51C). In contrast, when |*r*| ∈ [0,0.3) or |*r*| ∈ [0.7,1.0], both methods had comparable performance (Figure S51C).

The inference algorithm in VINTAGE is versatile, capable of handling either individual-level data or summary statistics as input. In scenarios involving summary statistics, the inference algorithm in VINTAGE makes an important assumption: that the two LD matrices, one from the gene expression mapping study and the other from the GWAS, share the same set of eigenvectors. Consequently, only a single LD matrix computed from a reference panel is needed to represent the two LD matrices in the model. Such assumption appears to hold reasonably well even when the two studies contain slightly different LD structures (Figures S52-S53). Specifically, for the gene-wise genetic variance test, regardless of whether the LD matrices were computed using 10,000 randomly selected WB individuals from UKBB or using 503 European individuals from 1000G (details in the Materials and methods), the p-values from VINTAGE remained well calibrated under the null (Figures S52A-B). For the gene-wise genetic correlation test, similar to what we have observed in the main simulations, the p-values from VINTAGE were on the conservative side, especially in the settings where *PVE*_1_ and *PVE*_2_ were small (Figures S52C-D). In addition, we found that the p-values from VINTAGE were less conservative when the LD matrices were computed using 1000G reference panel as compared to using UKBB (Figures S52C-D), highlighting the importance of using an LD matrix from a reference panel that is similar to the gene expression study (i.e., 1000G). Consistent with the simulations under the null, the power of VINTAGE’s genetic variance test was highly similar regardless of which LD matrices was used; while the power of VINTAGE’s genetic correlation test was slightly lower when the LD matrix was obtained from UKBB as compared to 1000G (Figure S53).

Several future extensions for VINTAGE are possible. First, while we have focused on modelling quantitative traits with VINTAGE, it would be desirable to extend VINTAGE to accommodate binary, count, or ordinal data types in a principled way, by, for example, extending VINTAGE into the generalized linear model framework. For example, we could use a probit or a logistic link to extend VINTAGE to directly model binary data type in GWAS. Extending VINTAGE to model different data types would broad the scope of VINTAGE’s applications and is thus an important avenue for future research. Second, we have focused on modelling gene expression data from a single tissue. However, determining the disease relevant tissue is often challenging, and the effect of gene expression on trait may act in a tissue-specific fashion^36^. In addition, restricting the analysis to a single tissue might also overlook the shared transcriptomic regulation across tissues, limiting the effective use of sample sizes of some expression mapping studies^37,38^. Therefore, extending VINTAGE to jointly analyze multiple tissue types from expression mapping studies such as GTEx has the potential to not only enhance the power of gene-based analysis but also pinpoint the relevant tissues to the trait of interest. Third, just like marginal/conventional TWAS, VINTAGE may be prone to horizontal pleiotropy. Recent strategies to control for horizontal pleiotropy include Liang et al.^17^, which uses a noncentral chi-squared distribution to account for polygenicity, and PMR^18^, which explicitly employs an Egger term or a variance component term to control for horizontal pleiotropy. Incorporating these strategies into VINTAGE is an important future direction. Lastly, we emphasize that VINTAGE is not limited to the integrative analysis of a GWAS and gene expression mapping study. Other molecular phenotypes such as DNA methylation, protein expression, and chromatin accessibility can also be integrated to attain a more comprehensive understanding of the molecular mechanisms underlying disease associations.

Finally, we explored two recent TWAS fine-mapping methods – cTWAS^39^ and GIFT^40^ – in relation to our marginal gene-wise genetic variance and correlation tests in real data applications. Detailed descriptions of the two methods are provided in Materials and methods. Note that these two methods are TWAS fine-mapping methods, which differ from the TWAS methods and SKAT-type methods that are the primary focus of the present study. TWAS fine-mapping aims to refine the identified gene-trait associations from TWAS within LD blocks to pinpoint genes that are more likely to be biologically relevant and thus are often considered as a secondary analysis following TWAS. For cTWAS, we integrated eQTLGen with UKBB and analyzed the same 18 quantitative traits, focusing on LD blocks containing genes identified by the genetic variance tests of VINTAGE. For GIFT, we focused our analysis on systolic blood pressure due to the computational burden of the GIFT software. Note that we did not evaluate cTWAS and GIFT in simulations because the simulation framework is targeted for TWAS but not TWAS fine-mapping. In the analysis, we found that most genes identified by cTWAS were also identified by the genetic variance test of VINTAGE (1610 out of 1846 genes across all traits; Figure S54A). After fine-mapping, the average number of genes identified per block reduced from 2.5 – 5.6 (mean = 4.1; median = 4.1) to 0.13 – 0.48 (mean = 0.29; median = 0.27) due to the nature of fine-mapping. However, cTWAS failed to identify any genes in a substantial proportion of regions (74%) that contain genes identified by the genetic variance tests. This is likely due to cTWAS being built upon the marginal/conventional TWAS framework, indicating that a fine-mapping method specifically tailored towards our framework is needed to optimize the fine-mapping performance. In addition, we found that 34 out of the 61 genes identified by the gene-wise genetic correlation tests were also identified by cTWAS, while 27 genes were missed, suggesting that cTWAS may have left out many genes with potential mediating effects, again consistent with its methodological connection to traditional TWAS (Figure S54B). Similarly, we found that most genes identified by GIFT were also identified by the genetic variance test of VINTAGE (52 out of 66 genes), while it failed to detect *CAMK1D* that was identified by the gene-wise genetic correlation test of VINTAGE (Figure S55). The results support the importance of developing fine-mapping methods based on the VINTAGE framework in the future.

### Materials and methods VINTAGE model

VINTAGE is a general analytic framework for integrating a gene expression mapping study with a genome-wide association study (GWAS) to identify genes that are associated with a trait of interest. VINTAGE unifies two seemingly unrelated methods, SKAT and TWAS, into the same analytic framework, and includes both methods as special cases. A key feature of VINTAGE is its ability to model jointly the SNP effects on gene expression in the expression study and the SNP effects on the trait of interest in the GWAS, thus facilitating the identification and interpretation of gene associations. VINTAGE requires either individual level data (described in this section) or summary statistics (described in a following section) from both the expression study and GWAS, which are assumed to contain non-overlapping individuals with the same genetic ancestry. With these two studies, VINTAGE examines one gene at a time. For the gene of focus, we denote ***y***_1_ as the *n*_1_-vector of gene expression measurements on *n*_1_ individuals in the expression study and ***X***_1_ as the corresponding *n*_1_ × *p* genotype matrix for its *p* cis-SNPs that reside in the genic and adjacent regulatory regions. We denote ***y***_2_ as the *n*_2_ -vector of trait measurements in the GWAS and ***X***_2_ as the corresponding *n*_2_ × *p* genotype matrix for the same set of *p* cis-SNPs. We center and standardize the gene expression vector ***y***_1_, the trait vector ***y***_2_, as well as each column of the genotype matrices ***X***_1_ and ***X***_2_ to have a mean of zero and variance of one. We consider the following two equations to model the relationship among SNPs, gene expression, and the trait:

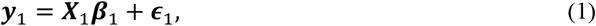

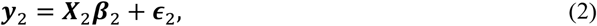

where we can ignore the intercepts in both equations due to data centering. Above, equation (1) describes the relationship between gene expression and cis-SNP genotypes in the gene expression data, where ***β***_1_ is a *p*-vector of SNP effects on the gene expression and ***ϵ***_1_ is an *n*_1_-vector of residual errors with each element independently following a normal distribution 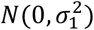. Similarly, equation (2) describes the relationship between the trait and the same set of SNPs in the GWAS data, where ***β***_2_ is a *p*-vector of SNP effects on the trait and ***ϵ***_2_ is an *n*-vector of residual errors with each element independently following a normal distribution 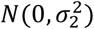. Our goal is to incorporate the SNP effects on gene expression in equation (1) to facilitate the inference of SNP effects on the trait in equation (2). To do so, we assume that the genetic effects of the *j*th SNP on the gene expression and the trait may be correlated with each other and follow a bivariate normal distribution *a priori*:

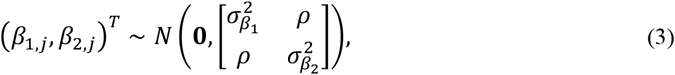

where 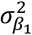 represents the proportion of variance in the gene expression explained by a cis-SNP, or the per-SNP heritability of the gene expression; 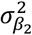 represents the proportion of variance in the trait explained by a cis-SNP, or the per-SNP heritability of the trait; and *ρ* represents the per-SNP local genetic covariance between the gene expression and the trait in the genomic region. We can further obtain the local heritability as 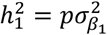 and 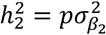 for the gene expression and trait, respectively, obtain the local genetic covariance as *ρ*_*g*_ = *pρ*, and obtain the local genetic correlation as 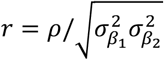.

Our key parameters of interest in the above model are 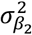 and *r*. Specifically, the variance component parameter 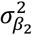 characterizes the effects of the cis-SNPs in the genic and adjacent regulatory regions of the gene on the trait and can be used to test the association of the gene with the trait. The local genetic correlation parameter *r* characterizes the similarity in the SNP effects on gene expression and trait, and thus can be used to quantify the amount of information that we can borrow from the gene expression mapping study to facilitate the mapping of gene associations in the GWAS. Importantly, the parameter *r* also quantifies the genetic effects on the trait mediated through gene expression, as the above model in equations (1)-(3) can be re-parameterized to form a mediation analysis model. Specifically, with the assumption in equation (3), *β*_1,*j*_ and *β*_2,*j*_ can be equivalently expressed as:

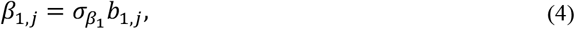

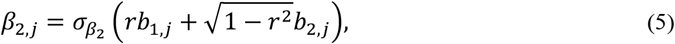

where *b*_1,*j*_ and *b*_2,*j*_ are independent latent variables that follow a standard normal distribution. Consequently, the equation (2) can be re-written as:

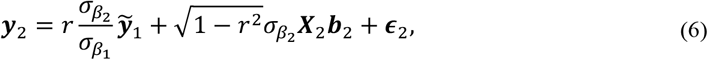

where 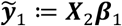 represents the *n*_2_-vector of genetically regulated gene expression for the *n*_2_ individuals in the GWAS. Pairing equations (1) and (6) leads to a mediation analysis model, where the trait is the outcome, gene expression is the mediator, and SNPs are the exposures^41^. In the mediation model, 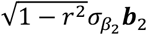 is the vector of direct effects (DE) of the SNPs on the trait while the product term 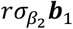 is the vector of indirect effects (IE) of the SNPs on the trait mediated through gene expression. We define the proportion of variance attributed to mediation effects as 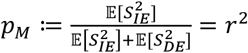, where *S*^2^ denotes sample variance. When *r* = 0, we have *p*_*M*_ = 0, indicating that gene expression does not mediate any of the genetic effects on the trait. When *r* = ±1, we have *p*_*M*_ = 1, indicating that the genetic effects on the trait are entirely mediated through gene expression.

To identify gene association with the trait, we test the null hypothesis *H*_0_: ***β***_2_ = 0, which represents the situation where none of the SNPs in the genic or adjacent regulatory regions of the gene have any effect on the trait. Such null hypothesis is equivalent to testing 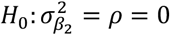, that both the variance parameter 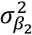 and the covariance parameter *ρ* are zero. Note that we test *ρ* instead of *r* under the null because *r* is no longer defined when 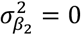. The alternative hypothesis that corresponds to the null is 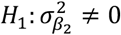, which consists of three scenarios depending on the specific value of *r*. Each scenario corresponds to a distinct role of gene expression in mediating the effects of SNPs on the trait and thus characterizes how informative the gene expression study is for testing the gene-trait association. The three scenarios are listed below:

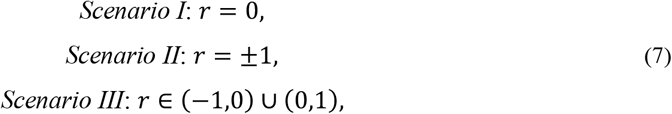

In scenario I, there is no local genetic correlation between the gene expression and the trait. The effects of the genetic variants on the trait may be mediated through other molecular mechanisms (e.g., protein regulatory changes, DNA methylation etc.) or in other tissue/cell types not examined in the gene expression study. Therefore, the gene expression mapping study is not informative for the gene association test in the GWAS. Consequently, the gene association test should be carried out using GWAS data only. This is a scenario that seamlessly aligns with the SKAT modeling assumption, which effectively tests the following hypothesis using GWAS data alone: 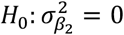 and *ρ* = 0 versus 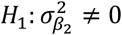 and *ρ* = 0.

In scenario II, the genetic effects on gene expression and the trait are perfectly correlated, indicating that the genetic effects on the trait are entirely mediated through gene expression. Therefore, the gene expression mapping study is fully informative for the gene association test in the GWAS. Consequently, the gene association test should incorporate both gene expression mapping data and GWAS data. This is a scenario that seamlessly aligns with the TWAS modeling assumption, which constrains the genetic effects on the trait to be a scalar multiplication of the genetic effects on the gene expression and further examines if this scalar is zero or not. Using our notation, the two-stage inference framework of TWAS can be expressed by the equation 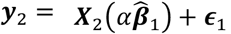 where 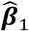 is a *p*-vector of genetic effects on the gene expression estimated using the gene expression data; and *α* is a scaling factor of the estimated effects. TWAS tests the null hypothesis *H*_0_: *α* = 0, which is equivalent to testing 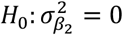 assuming *r* = ±1 in our modeling framework.

By including the above two scenarios as part of the alternative hypothesis, our modeling framework effectively includes SKAT and TWAS as two special cases. Importantly, different from SKAT or TWAS, our modeling framework also considers scenario III, which represents an intermediate scenario where gene expression mediates a portion of the genetic effects on the trait. By including scenario III as part of the alternative hypothesis, our modeling framework relaxes the strong assumptions on the local genetic correlation made in either SKAT or TWAS and effectively allows us to infer the local genetic correlation based on the data at hand to improve the statistical power for the gene-trait association analysis.

Besides gene-trait association test, we also estimate the local genetic correlation parameter *r* and test the null hypothesis *H*_0_: *r* = 0. Such local genetic correlation test allows us to examine whether the gene expression mapping study is informative for the gene association test in GWAS and whether the gene expression mediates the genetic effects on the trait. Because the two tests focused on the variance and covariance parameters, respectively, we refer to the hypothesis test for gene association (i.e., 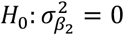 and *ρ* = 0) as the “gene-wise genetic variance test “ and the hypothesis test for local genetic correlation (i.e., *H*_0_: *r* = 0) as the “gene-wise genetic correlation test “.

Finally, we would like to make two comments on the model. First, due to the observational nature of the data, the statistical model itself is not well-equipped to distinguish the directionality of the mediation effects. In particular, a non-zero local genetic correlation can be caused by either mediation effect or reverse mediation effects, the latter of which occurs when the trait mediates the genetic effects on gene expression. While such reverse mediation effects are more biologically likely to occur in *trans*-^42^ rather than *cis*-region of a gene, at least for the majority of genes, we still caution against over-interpreting the mediation effects inferred from our model. Second, the definition and estimation of the genetic correlation parameter requires polygenicity, meaning that all or at least a large number of SNPs display non-zero effects on both gene expression and the trait. Caution needs to be taken under sparse genetic architectures, where only a small number of SNPs display non-zero effects on gene expression or the trait, as the genetic correlation parameter becomes less defined or identifiable. In the extreme case when *q* = 1, the local genetic correlation parameter is no longer well-defined and becomes unidentifiable. This is because the data lack sufficient information to distinguish between independent SNP effects with a correlation of zero and fully correlated SNP effects with a correlation of either +1 or -1 depending on the sign consistency of SNP effects on gene expression and the trait, or anywhere in between. In this case, as we will show in the simulations, the local genetic correlation is indeed estimated close to either +1 or -1. Consequently, the genetic variance test of VINTAGE is expected to behave similarly as TWAS. Certainly, as we will show in the real data applications, the genetic variance test of VINTAGE does not behave similarly as TWAS but instead resembles that of SKAT, suggesting that the polygenic assumption likely holds for many genes in practice.

### An optimal and unified gene-wise genetic variance test

We have developed an optimal and unified variance component score test to carry out the aforementioned gene-wise genetic variance test. The test is optimal in the sense that it maximizes power within the class of tests that is a linear combination of SKAT and TWAS statistics and achieves robust performance to a wide range of local genetic correlation values between the gene expression and the trait. The test is unified as it includes the SKAT and TWAS test statistics as two special cases. Note that devising such effective gene-wise genetic variance test posed significant technical challenges due to the concurrent presence of variance and covariance parameters under the null hypothesis, compounded by the compositional nature of the alternative hypothesis. Indeed, the test presented here stands out as the sole functional one among the five different tests we have carefully explored. The remaining four tests were unsuccessful for various reasons, and we provide a detailed summary of these failed attempts in the Supplementary Text for the benefit of the community.

To derive the test statistic, we first obtain the scores for the variance component 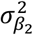 and the covariance component *ρ* under the null hypothesis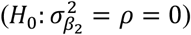:

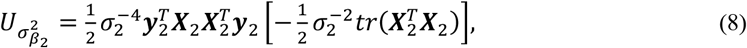

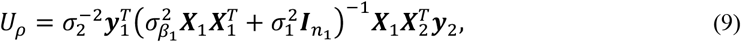

where 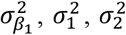 are treated as nuisance parameters; *tr* denotes matrix trace; and 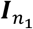 is an *n* × *n* identity matrix. Note that the second term of 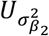 (i.e., 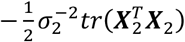)) can be omitted because it is not random.

The above two scores for the variance component parameters serve as backbone for constructing our test statistic. Importantly, these two scores are closely related to the test statistics of SKAT and TWAS (detailed proofs are in the Supplementary Text):

#### Remark 1.

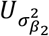, *the score for the variance component* 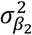, *is equivalent to the SKAT statistic with an unweighted and linear kernel function under the null hypothesis*.

#### Remark 2.

*U*_*ρ*_, *the score for the covariance component ρ, is equivalent to the two-stage TWAS test statistic under the null hypothesis, when the following two conditions are satisfied: (1) A BLUP prediction model is used in the expression study to obtain the SNP weights on gene expression; (2) the uncertainty associated with the SNP weights on gene expression is taken into account when constructing the TWAS test statistic*.

Because of the close relationship of the two scores to the SKAT and TWAS test statistics and because SKAT and TWAS represent two extreme cases of the alternative hypothesis in our modeling framework, we obtain the final test statistic in the form of a weighted combination of the two scores as follows:

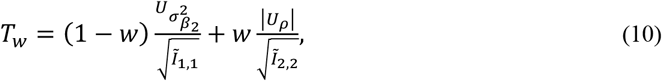

where Ĩ _1,1_ and Ĩ _2,2_ are the elements in the information matrix that correspond to 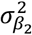 and *ρ*, respectively. Normalizing 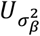 and *U*_*ρ*_ with Ĩ _1,1_ and Ĩ _2,2_ ensures that the two terms are on the same scale. Note that the second term uses the absolute value of *U*_*ρ*_ to keep the sign consistent with 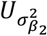 in the first term, so that both terms in *T*_*w*_ are positive and their large values reflect a deviation from the null hypothesis. The weight *w* is a scalar that controls the relative contribution of the two terms to the test statistic. Specifically, when *w* = 0, *T*_*w*_ (= *T*_0_) reduces to the SKAT statistic and is expected to achieve high power when the local genetic correlation equals zero. When *w* = 1, *T*_*w*_ (= *T*_1_) reduces to the TWAS test statistic and is expected to achieve high power when the local genetic correlation equals one or negative one. When 0 < *w* < 1, *T*_*w*_ is expected to achieve high power when the local genetic correlation values are in the range of (-1, 0) or (0, 1). In practical applications, since the true local genetic correlation is unknown, we conduct grid search on a set of *K* prespecified weight values ranging from 0 to 1 with equal increments (*K* = 11 by default). For each weight value, we calculate the corresponding test statistic *T*_*w*_ and p-value *p*_*w*_. We then use the Cauchy combination rule^20,43^ to combine the p-values obtained from different weights into a single p-value, denoted as *p*_*g*_. In the last step, the combined Cauchy test statistic is given by:

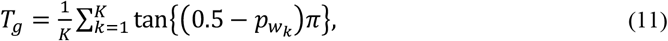

where 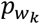 is the p-value that corresponds to 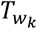 and tan denotes the tangent function. We convert the combined test statistic *T*_*g*_ into the final p-value based on the Cauchy distribution.

While the above procedure appears to be straightforward, the step of obtaining calibrated p-value 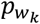 from each 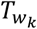 is challenging, as the exact distribution of 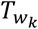 is difficult to obtain. Therefore, we developed a highly scalable simulation-based approach to simulate the test statistics under the null, with which we obtain 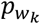 from 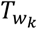 . Detailed procedures of the simulation-based testing approach are described in the Supplementary Text.

### A powerful gene-wise genetic correlation test

In addition to the gene-wise genetic variance test assessing the gene association with the trait, we have developed a gene-wise genetic correlation test to evaluate the role of gene expression in mediating the genetic variant and trait association. To do so, we obtain the standard Rao’s score test statistic with respect to the local genetic correlation *r* under the null hypothesis (*H*_0_: *r* = 0):

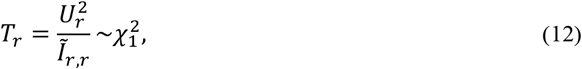

where 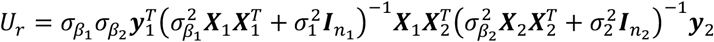 is the score for *r* with nuisance parameters 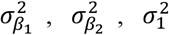 and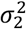; and 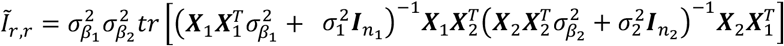 is the element in the information matrix that corresponds to *r*. We denote the p-value corresponding to the test statistic in equation (12) as *p*_*r*_ and we calculate it based on a chi-squared distribution with the degrees of freedom equal to one. Detailed derivations are provided in the Supplementary Text. Note that the local genetic correlation is defined only for genes with non-zero genetic variance. Therefore, we carry out the gene-wise genetic correlation test only for genes that exhibit significance in the gene-wise genetic variance test.

### Scalable inference algorithm

We have developed a parameter-expanded expectation maximization (PX-EM) algorithm for inference. To enable scalable computation, instead of directly working on equations (1)-(2), we transformed the genotype matrices and their corresponding coefficients into the principal component (PC) space. Specifically, we reparametrize the models defined in equations (1)-(2) into the following transformed models:

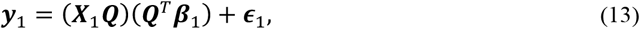

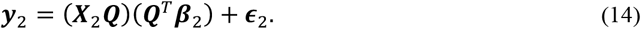

Above, ***Q*** is a *p* × *p* orthogonal matrix of eigenvectors derived from the eigenvalue decomposition of the LD matrix ***R***_*l*_, where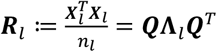 . Here, we have assumed that the two LD matrices, one from the eQTL study (*l* = 1) and the other from the GWAS (*l* = 2), share the same eigenvectors, an assumption that approximately holds when both studies contain the same genetic ancestry. We note that the above transformation from equations (1)-(2) to (13)-(14) does not change the likelihood for all parameters because of the orthogonality of ***Q*** (i.e., ***Q***^*T*^***Q*** = ***QQ***^*T*^ = *I*_*p*_). However, such transformation decorrelates the columns of the transformed design matrices 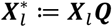 and thus substantially improves the computational speed of the PX-EM algorithm. In particular, after the initial eigenvalue decomposition that incurs a cubic computational complexity in *p*, the computational complexity of the PX-EM algorithm becomes linear in *p* in each iteration.

### Extension towards summary statistics

Although we have presented VINTAGE based on individual-level genotype, gene expression, and phenotype data, we note that VINTAGE can be easily extended to utilize only summary statistics from the gene expression study and GWAS as input for inference. These summary statistics take the forms of marginal z-scores and a SNP-SNP correlation matrix. The SNP-SNP correlation matrix is commonly referred to as the LD matrix and can be estimated from a reference panel containing individuals with the same genetic ancestry as those in the gene expression study and GWAS. To address potential biases caused by an inaccurate estimation of the LD matrix, we focus on the top k eigenvalues and eigenvectors for inference, where *k* is determined in a way such that 99% of variance in the LD matrix is explained. Specifically, instead of 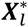, we work on the reduced design matrix 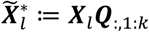 and its corresponding coefficients 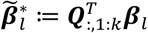. We then express the sufficient statistics in our model using summary statistics following Zou et al.^44^:

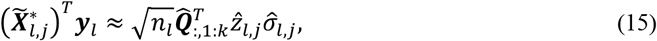

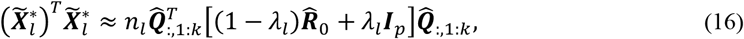

where the subscript *l* denotes the gene expression (*l* = 1) or GWAS data (*l* = 2); the subscript *j* = 1, … *k* denotes the *j*th genotype PC; 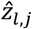 is the marginal z-score obtained from marginal regression analysis; 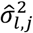 is the residual error variance estimate obtained from marginal regression analysis and can be calculated from 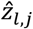 sin the form of 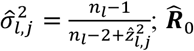 is a *p* × *p* LD matrix that can be estimated from a reference panel; and 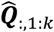 is the *p* × *k* matrix of top *k* eigenvectors obtained from the eigenvalue decomposition of 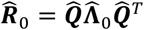.

We make two comments regarding equations (15)-(16). First, the right-hand side of equation (15) includes a product term 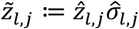, representing the PVE (Proportion of phenotypic Variance Explained)-adjusted marginal z-score. This inclusion distinguishes equation (15) from the typical approximation equation commonly found in the GWAS literature^45-47^, where the parameter 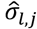 is omitted and only 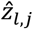 remains. We included the parameter 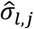 in equation (15) to make the equality 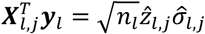 exact, allowing it to handle not only the conventional GWAS scenario, where the SNP effect size on the outcome trait is small and negligible (i.e.,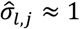), but also the gene expression study scenario, where the SNP effect size on the gene expression is non-negligible. Second, the right-hand side of equation (16) includes a regularized form of the LD matrix 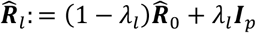, where *λ*_*l*_ ∈ [0,1] serves as the degree of regularization. We estimate the regularization parameter *λ*_*l*_ by maximizing the likelihood under the null (***β***_*l*_ = 0), in the form of 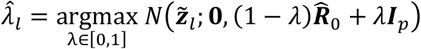, to improve consistency between the regularized LD matrix and PVE-adjusted z-scores following Zou et al^44^.

### Guideline on using VINTAGE in practice

VINTAGE integrates GWAS with gene expression data, requiring both datasets as input. For each dataset, VINTAGE can utilize either individual-level data or summary statistics. The summary statistics are in the form of marginal z-scores and SNP-SNP correlation matrices, which can be estimated from a reference panel. With the input data, VINTAGE offers two distinct statistical tests: a gene-wise genetic variance test to assess gene-trait association and a gene-wise genetic correlation test to assess expression mediation. The first genetic variance test unifies SKAT and TWAS and is applied for a genome-wide scan to assess whether SNPs within each gene region are associated with the trait. The output from this test is a set of p-values, one for each gene, reflecting the association significance of SNPs in the gene region with the trait. For genes with significant associations identified in the first test, VINTAGE provides the second genetic correlation test to assess further whether these associations are mediated by gene expression. The output from this test includes a p-value and an estimate of the local genetic correlation for each gene, the latter is further processed to calculate the proportion of SNP effects on the trait mediated by gene expression. Together, these two tests provide a comprehensive characterization of the gene-trait relationships. We provide VINTAGE as an open-source R package with detailed tutorials guiding the users to analyze their own datasets (https://zhengli09.github.io/VINTAGE-analysis/).

### Proof-of-concept and main simulations

We conducted extensive and realistic simulations to evaluate the performance of our method and compared it with existing approaches (details in the Compared methods). To do so, we first obtained all 487,409 individuals in UKBB with imputed BGEN genotype files. We excluded among them those who are not of White British (WB) descent (*n* = 78,437), who are not included in the genotype PC computation (*n* = 71,427 related samples), who have sex chromosome aneuploidy (*n* = 337), and who are redacted (*n* = 10). A total of 337,198 individuals were retained after these QC steps. Among them, we randomly selected 10,000 and 100,000 non-overlapping individuals to serve as the gene expression and GWAS data, respectively. For the selected individuals, we extracted 92,693,895 autosome variants in the UKBB 2015 release and performed SNP quality control (QC) by filtering out variants with a minor allele frequency (MAF) < 5%, with an imputation information score ≤ 0.8, with a Hardy-Weinberg equilibrium (HWE) test p-value < 10^-10^, with a genotype call rate < 0.95, are indels and are not present in the Haplotype Reference Consortium Release 1.1 (HRC r1.1) imputation file. After these QC steps, we retained 5,119,766 SNPs that are common in both gene expression and GWAS data. Afterwards, we extracted cis-SNPs that reside within 100kb upstream of the transcription start site (TSS) and 100kb downstream of the transcription end site (TES) of each gene and randomly selected 10,000 genes that have at least 10 cis-SNP for simulations. Note that we followed previous studies^18,48,49^ and used a region size of 100kb instead of the size of 1Mb that were used in some earliest TWASs^7,8^ to mitigate the LD-hitchhiking effects^50^. The size of the genomic regions ranges from 200.4kb to 2,372.9kb (mean = 268.2kb; median = 232.2kb) with the number of cis-SNPs per gene ranging from 11 to 5,301 (mean = 475; median = 425). Afterwards, we centered and standardized each column of the genotype matrices to have a mean of zero and a variance of one. We used all 10,000 genes for null simulations. We used a random subset of 1,000 genes for power simulations due to the large number of simulation settings explored there.

With the extracted genotype data, we simulated gene expression and trait based on equations (1) and (2). Specifically, for each gene in turn, we randomly selected *q* cis-SNPs to have non-zero effects on both gene expression and trait, where *q* is set to be either 1, 2, 10, or *p*. Note that *q* = *p* represents a polygenic genetic architecture where all cis-SNPs have a non-zero effect while the other values of *q* represent sparse genetic architectures. For each SNP with non-zero effects, we simulated its effect sizes on expression and trait from a bivariate normal distribution with mean 0 and a variance-covariance matrix 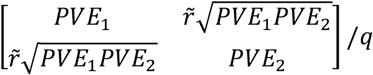. We set the local genetic correlation parameter 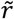 in the bivariate normal distribution to be either -1, -0.8, -0.5, 0, 0.5, 0.8, or 1. In addition to the genetic effects, we simulated the residual errors for either the gene expression or the trait from a normal distribution, with mean zero and a variance set to be either 1 − *PVE*_1_ (for gene expression) or 1 − *PVE*_2_ (for the trait). We then summed the genetic effects and the residual errors to form the simulated gene expression and trait. The simulation setup ensures that the average proportion of phenotypic variance in gene expression explained by cis-SNPs (i.e., cis-SNP heritability) is *PVE*_1_ while the average proportion of phenotypic variance in the trait explained by cis-SNPs is *PVE*_2_. We varied *PVE*_1_ to be 0.5%, 1%, or 5% and varied *PVE*_2_ to be 0%, 0.02%, 0.04%, or 0.06% in our simulations. Finally, we centered and standardized the gene expression and trait separately to have zero mean and unit variance.

With the simulated gene expression and trait data, along with the corresponding genotype matrices, we fitted linear regression models in the two datasets to obtain marginal z-scores. In addition, we obtained an LD matrix from a reference panel, which includes either the same 10,000 UKBB WB individuals in the gene expression data (i.e., in-sample LD matrix) or 503 individuals of European ancestry from the 1000 Genomes Project (1000G) (i.e., external LD matrix; data details in the Real data applications). With the marginal z-scores and LD matrix as input, we applied our method and other approaches for gene-based analysis.

We established a set of baseline simulation settings with a polygenic genetic architecture (*q* = *p*) that are fitted using in-sample LD matrices. In the baseline settings, we examined all combinations of *PVE*_1_, *PVE*_2_, and 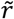 parameters, where *PVE*_1_ = 1% represents moderate cis-SNP heritability of gene expression and *PVE*_2_ = 0.04% represents moderate genetic effects on the trait. On top of the baseline simulation settings, we varied the parameter *q* to evaluate the performance of different methods under a sparse genetic architecture and varied the LD reference panel to evaluate the influence of using an external LD matrix. In the simulations, we set up the null simulations to have *PVE*_1_ ≠ 0, *PVE*_2_ = 0 and 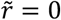 for the gene-wise genetic variance test. For the gene-wise genetic correlation test, we set up the null simulations to have *PVE*_1_ ≠ 0, *PVE*_2_ ≠ 0 and 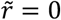 under a polygenic genetic architecture. However, for simulations under a sparse genetic architecture, there is an important complication for evaluating the type I error and power. To understand this complication, let us first consider a simple example with two distinct sparse scenarios (*q* = 1 for simplicity). In the first sparse scenario, a single SNP in the genomic region is causal, affecting both gene expression and the trait. This scenario clearly represents an alternative setting of non-zero local genetic correlation with true *r* = ±1, indicating that the SNP’s effect on the trait is potentially mediated through gene expression. The complexity, however, arises from the simulation of this setting. When a bivariate normal distribution is used to simulate causal SNP effects on both gene expression and the trait, the correlation parameter in the bivariate normal distribution, denoted as 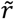, can take any value within [-1, 1]. Regardless of the value of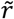, the resulting scenario remains unchanged: the causal SNP affects both gene expression and the trait, leading to the true *r* = ±1. Therefore, in this sparse scenario, 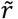 becomes non-informative, as any value of 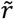 produces the same outcome of *r* = ±1. Consequently, simply setting 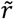 to be zero does not represent null settings for the local genetic correlation test. To see a proper null setting, we consider the second sparse scenario. In the second sparse scenario, a single SNP in the genomic region is causal for gene expression, while a different SNP is causal for the trait. This scenario clearly represents a null setting of zero local genetic correlation, where true *r* = 0 and the SNP’s effect on the trait is not mediated through gene expression. As you can see, the complexity here arises from the discrepancy between the simulation parameter 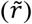 and the true underlying local genetic correlation (*r*), a situation that does not occur in non-sparse settings. In non-sparse settings, 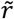 is always consistent with *r*. Due to this complication, instead of showing power versus 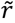, we show power versus *r* to evaluate the performance of the gene-wise genetic correlation test under sparse settings, with *r* calculated as the Pearson correlation coefficient between the simulated genetic effects on gene expression and on trait. In addition, we examine the null settings described in the second sparse scenario above, where two distinct sets of SNPs, each consists of *q* (= 1, 2, or 10) SNPs, display non-zero effects: one on gene expression and the other on the trait. In these settings, we first randomly select *q* SNPs to have non-zero effects on gene expression. Among the remaining SNPs in the region that are not in extreme high LD with the selected SNPs (R^2^ < 0.95), we randomly select another *q* SNPs to have non-zero effects on the trait. The corresponding effect sizes of these SNPs are simulated in the same way described earlier. Because the two sets of SNPs do not overlap with each other, they represent null settings for the genetic correlation test.

With the above setup, we first carried out a set of proof-of-concept simulations for the gene-wise genetic variance test. In these simulations, we examined the performance of the individual test, *T*_*w*_, each constructed under a different weight value of *w*, serving as a component for constructing the final gene-wise genetic variance test. The weights range from 0 to 1 with increments of 0.1, resulting in a total of 11 individual tests. Here, we focused on the baseline simulations and explored a total of 66 settings that included 3 nulls and 63 alternatives. This set of simulations allows us to examine how the behavior of individual tests constructed under different weights varies with respect to different local genetic correlations.

Besides the proof-of-concept simulations, we also carried out a set of main simulations to evaluate the performance of our method and compare it with other approaches. We evaluated two types of tests including the gene-wise genetic variance test and the gene-wise genetic correlation test. In the main simulations, we explored a total of 375 simulation settings that included 15 nulls and 315 alternatives for the genetic variance test and a total of 315 simulation settings that included 45 nulls and 315 alternatives for the local genetic correlation test.

In both proof-of-concept and main simulations, we performed 10,000 simulation replicates for each null simulation setting to evaluate the performance of different methods on type I error control. For each alternative simulation setting, we performed 1,000 simulation replicates to evaluate power. For the gene-wise genetic variance test, we calculated the power of each method as the proportion of replicates passing the p-value threshold of 2.5 × 10^−6^ (i.e., 0.05/20000). For the gene-wise genetic correlation test, we followed Zhang et al.^14,19^ and calculated the power of each method as the proportion of replicates passing the p-value threshold of 0.05 due to the very small sample size in the gene expression study and consequently the lack of power in local genetic correlation analyses. In all simulation scenarios, we also examined the estimation accuracy of the local genetic correlation parameter.

### Additional simulations

Besides the proof-of-concept and main simulations described above, we conducted additional simulations to evaluate the influence of various other factors, including the sample size of the gene expression study, unequal LD structures, inconsistent genetic architectures, and settings where the two studies have distinct sets of non-zero effect SNPs, on the performance of the two tests. We also examined the performance of the genetic correlation test after accounting for the influence of the genetic variance test and evaluated two existing methods, COJO^23^ and MESuSiE^24^, for their ability to accurately estimate the number of SNPs with non-zero effects on gene expression and the trait. Details of these simulations are provided below.

#### Evaluating the influence of the sample size of the gene expression study

We conducted additional simulations to evaluate the power of the gene-wise genetic correlation test for data with a sample size comparable to the Genotype-Tissue Expression (GTEx) study (n = 838)^3^. In the simulations, we focused on baseline settings with moderate *PVE* parameters and varied the sample size of the gene expression study to be 500 or 1,000. We evaluated the performance of VINTAGE, SUPERGNOVA, and LAVA for gene-wise genetic correlation test and compared it with our main simulations, where the sample size was set to be 10,000 that mimics eQTLGen study.

Besides the above simulations, we also conducted real data-informed simulations, where we set the simulation parameters fully based on real data estimates and varied the sample size of the gene expression mapping study. These additional simulations in the real data analysis allowed us to evaluate how sample size of the gene expression mapping study contributes to the power of both gene-wise genetic variance tests and gene-wise genetic correlation tests in realistic applications. Specifically, we focused on the same set of 13,725 genes analyzed in the real data applications (data details in the Real data applications). We obtained the cis-SNP heritability estimates of gene expression (i.e., 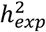), cis-SNP heritability estimates of 18 analyzed traits (i.e., 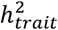), and the local genetic correlation estimates between gene expression and the trait (i.e., *r*) in each genomic region, all from VINTAGE. We extracted the parameter estimates for the main trait of focus, SBP, and obtained the LD matrix using the same 10,000 UKBB WB individuals in the main simulations. In these additional simulations, we directly simulated marginal z-scores for the gene expression mapping study with a sample size of *n*_1_ and GWAS with a sample size of *n*_2_. To do so, we first simulated the SNP effect sizes on gene expression and on trait following the same procedure described in the main simulations. There, we set *PVE*_1_ to be 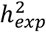, *PVE* to be 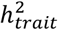, *local genetic correlation* to be *r*, and focused on a polygenic genetic architecture. Afterwards, we simulated marginal z-scores for each study from a multivariate normal distribution with mean 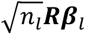 and variance-covariance matrix ***R***, where ***β***_*l*_ are simulated SNP effect sizes on gene expression (*l* = 1) or on trait (*l* = 2), and ***R*** is the gene-wise LD matrix. We varied the sample size of the gene expression mapping study to be 100, 500, 1,000, 2,000, 5,000, 10,000, 20,000, 50,000, 100,000, 200,000, and 361,194, while fixing the sample size of the GWAS to be 361,194. Finally, with the marginal z-scores and LD matrix as input, we applied VINTAGE for the gene-wise genetic variance test and gene-wise genetic correlation test across the 13,725 genes. For power calculation, we evaluated the power of the two types of tests in different parameter ranges and calculated the power as the proportion of the genes that passed the p-value significance threshold in each test. Specifically, for the gene-wise genetic variance test, we set the parameter range for 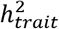 to be [1×10^-6^,1×10^-5^), [1×10^-5^,5×10^-5^), [5×10^-5^,1×10^-4^), or [1×10^-4^,1]; set the parameter range for the absolute value of *r* (i.e., |*r*|) to be [0,1], [0.5,1], or [0.8,1]; and set the p-value threshold to be the same value of 2.5 × 10^−6^ as in the main simulations. For the gene-wise genetic correlation test, we set the parameter range for 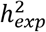 to be [0.01,1] or [0.05,1]; set the parameter range for 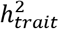 to be [1×10^-5^,5×10^-5^), [5×10^-5^,1×10^-4^), or [1×10^-4^,1]; set the parameter range for |*r*| to be [0.1,1], [0.5,1], or [0.8,1]; and set the p-value threshold to be 0.05 divided by the number of genes that passed the gene-wise genetic variance test.

#### Evaluating the influence of unequal LD structures

We explored the scenario where the two LD matrices, one from the gene expression mapping study and the other from the GWAS, deviate slightly from each other. Specifically, instead of simulating both gene expression and trait based on individuals from UKBB, we simulated gene expression based on the genotypes of 503 European individuals from 1000G, while simulated the trait based on 100,000 WB individuals from UKBB. We computed the LD matrix using either the same set of 10,000 UKBB WB individuals for simulating the gene expression data in the main simulations or 503 European individuals from 1000G. To account for the influence of the much smaller sample size of the gene expression mapping study on power, we increased the *PVE*_1_ parameter to 10%, 20%, and 50% accordingly, ensuring that *n*_1_*PVE*_1_ remains approximately the same. The other parameters were set to be consistent with the baseline settings in the main simulations.

#### Evaluating the influence of inconsistent genetic architectures

We conducted additional simulations to evaluate the performance of both the gene-wise genetic variance and correlation tests under settings where the genetic architecture is sparse in one study, while it is polygenic in the other. Specifically, we examined two scenarios: In Scenario I, we set the genetic architecture for gene expression in the expression study to be sparse and set the genetic architecture for the trait in GWAS to be polygenic; in Scenario II, we reversed these conditions. In either scenario, we randomly selected *q* cis-SNPs to have non-zero effects on both gene expression and the trait, where *q* was set to be 1, 2, or 10. For sparse architecture, we set the proportion of variance in gene expression or trait explained by these *q* SNPs to be *PVE*_*l*_ . For polygenic architecture, we set the proportion of variance in gene expression or trait explained by these *q* SNPs to be 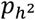· *PVE*_*l*_, while set the proportion explained by the remaining cis-SNPs to be 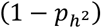s · *PVE*_*l*_. In the simulations, we specified 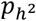 to be 0.3, 0.5, or 0.8 and fixed *PVE*_*l*_ to be 1% for the gene expression study (*l* = 1) and to be 0.04% for GWAS (*l* = 2). For each of the *q* SNPs with non-zero effects, we simulated its effect sizes from a bivariate normal distribution with mean 0 and a variance-covariance matrix being either 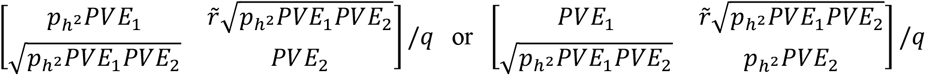 when either the gene expression or trait has a polygenic genetic architecture. For the remaining cis-SNPs in the polygenic phenotype, we simulated their effect sizes from a normal distribution with a mean of 0 and a variance of 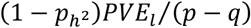. Similarly, we set the local genetic correlation parameter 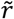 to be either -1, -0.8, -0.5, 0, 0.5, 0.8, or 1 in the bivariate normal distribution.

#### Evaluating genetic correlation test accounting for the influence of the genetic variance test

We conducted additional simulations where we took into account the influence of the genetic variance test to evaluate the performance of the genetic correlation test. For this evaluation, we focused on the baseline simulation settings and considered two procedures for evaluating the genetic correlation test: one ignores the genetic variance test, as described in the main simulation studies, while the other takes it into account by first conducting the genetic variance test and only performing the genetic correlation test if it passed the significance threshold of 2.5 × 10^−6^ in the genetic variance test. We calculated the power of the genetic correlation test under each procedure based on two p-value thresholds: a nominal threshold of 0.05 and a Bonferroni corrected threshold.

#### Evaluating COJO and MESuSiE for estimating the number of non-zero effect SNPs

We conducted additional analyses to infer the number of SNPs with non-zero effects on gene expression, denoted as *q*_1_, and the number of SNPs with non-zero effects on trait, denoted as *q*_2_, within each gene region. To account for the potential overlap, we further introduced *q*_*o*_ to denote the number of SNPs with non-zero effects on both gene expression and the trait. We termed the SNPs with non-zero effects on both gene expression and the trait as SNPs with shared effects, and the SNPs with non-zero effects on either gene expression or the trait as SNPs with study-specific effects. We evaluated two existing methods, COJO (version 1.94.1) and MESuSiE (version 1.0), for their ability to accurately estimate these numbers through simulations. To do so, for each gene of focus, we randomly selected one shared cis-SNP to have a non-zero effect on both gene expression and the trait (i.e. *q*_*o*_ = 1), and selected two additional cis-SNPs -- one with a non-zero effect on gene expression and the other with a non-zero effect on the trait (i.e. *q*_1_ = 2 and *q*_2_ = 2). In these simulations, we further set *PVE*_1_ = 1%, *PVE*_2_ = 0.04%, and varied *r* to be either -1, -0.8, -0.5, 0, 0.5, 0.8, and 1. For the shared cis-SNP, we simulated its effect sizes on gene expression and trait from the same bivariate normal distribution as in the main simulations. For the study-specific cis-SNPs, we simulated their effect sizes for gene expression from a normal distribution with a mean of 0 and a variance of *PVE*_1_*⁄q*_1_, and their effect sizes for the trait from a normal distribution with a mean of 0 and a variance of *PVE*_2_*⁄q*_2_. We then applied COJO and MESuSiE to infer the number of shared and study-specific SNPs with non-zero effects on gene expression and the trait. For COJO, we followed its online tutorial and performed a stepwise selection, selecting independently associated SNPs for gene expression and the trait separately. We further processed the outputs from COJO to classify SNPs as either shared or study-specific. For MESuSiE, we adapted it to jointly analyze the gene expression and trait data, estimating the number of shared and study-specific SNPs based on those included in the credible sets for each category. In the process, we followed the MESuSiE online tutorial and set the maximum number of non-zero effects within the region to 20.

#### Evaluating settings where the two studies have distinct sets of non-zero effect SNPs

We conducted similar simulations where we used two sets of SNPs, with 1, 2, or 10 SNPs in each set: one set with non-zero effects on gene expression and the other set with non-zero effects on the trait, to evaluate the performance of the gene-wise genetic variance test. The two sets of SNPs were distinct and did not exhibit extreme high LD (R^2^ < 0.95) with each other. In these simulations, we focused on alternative settings with moderate *PVE* parameters. We did not further explore the null settings as those are essentially the same as the previous simulations with a sparse genetic architecture, where only one set of SNPs has non-zero effects on gene expression and the effects of all SNPs on the trait are zero.

### Real data applications

We applied our method and the other approaches to analyze 18 quantitative traits in the UKBB using GWAS summary statistics. These traits include standing height (SH), red blood cell count (RBC), RBC distribution width (RDW), white blood cell count (WBC), platelet count (PLT), systolic blood pressure (SBP), diastolic blood pressure (DBP), eosinophils count (EOS), neuroticism score (NS), bone mineral density (BMD), body mass index (BMI), basal metabolic rate (BMR), hip circumference (HC), waist circumference (WC), forced expiratory volume (FEV1), forced vital capacity (FVC), age at menarche (AM), and years of education (YE). Among the 18 analyzed traits, we consider RBC, RDW, WBC, PLT, SBP, DBP, and EOS as blood-related traits. These traits exhibit an observed SNP heritability estimated to be above 0.1, as reported by Neales’s lab. Their summary data (imputed-v3) was downloaded from the online resource of Neale’s lab (https://github.com/Nealelab/UK_Biobank_GWAS), which contains marginal GWAS z-scores for 13.7 million SNPs on 361,194 individuals of European ancestry. We focused our analysis on autosomal SNPs with a MAF ≥ 1%, resulting in an analyzed set of 9,376,157-9,377,442 SNPs across the 18 traits.

In addition to the GWAS data, we also obtained cis-eQTL mapping summary statistics from the eQTLGen phase I study (https://eqtlgen.org/phase1.html)^25^. The eQTLGen data consists of blood (80.4%) and peripheral blood mononuclear cell (PBMC; 19.6%) samples from 31,684 individuals across 37 cohorts, with the majority of individuals being of European ancestry. The summary statistics data includes marginal z-scores for 11 million cis-eQTLs that have a MAF ≥ 1% and that are within 1Mb around the center of each gene. We focused our analysis on protein-coding genes annotated in GENCODE release 40^51^.

Besides the GWAS and eQTL data, we obtained individual-level genotype data from the 1000 Genomes Project phase 3 to serve as the LD reference panel. Specifically, we obtained genotype data from 503 European individuals in the 1000G, retained 6,042,565 SNPs with a MAF ≥ 5%, and calculated gene-wise LD matrices for these SNPs.

In the analysis, we extracted cis-SNPs that reside within 100kb upstream of the TSS and 100kb downstream of the TES of each gene. We obtained a common set of SNPs across the UKBB data, eQTLGen data, and the 1000G reference panel. We filtered out strand ambiguous SNPs that have reference and alternative alleles A/T, T/A, C/G, and G/C. In addition, we applied the diagnostic tool implemented in the susieR package (version 0.12.41)^44^ to filter out SNPs that have a potential mismatch with the LD matrix in either the UKBB or eQTLGen data. In total, we analyzed 13,725 protein-coding genes, with the corresponding genomic region size ranging from 200.1kb to 2,505.0kb (mean = 269.6kb; median = 232.1kb) and the number of cis-SNPs per gene ranging from 1 to 8,367 (mean = 404; median = 361). Across all LD matrices estimated from the 1000G reference panel, the number of top k eigenvalues and eigenvectors that explained 99% of the variance in each LD matrix ranged from 1 to 392 (mean = 52; median = 45). For the gene-wise genetic variance test, we applied Bonferroni correction^52^ and declared a gene to be significantly associated with the trait if its p-value passed the transcriptome-wide significance threshold of 2.5×10^-6^ (i.e., 0.05/20000). For significant genes identified in the gene-wise genetic variance test, we further conducted a gene-wise genetic correlation test and declared significance based on a Bonferroni corrected p-value threshold of 0.05 divided by the number of tested genes.

Besides analyses with eQTLGen, we applied VINTAGE to integrate GTEx data with UKBB and analyzed the same 18 quantitative traits. Specifically, we obtained genotype and gene expression data for 558 individuals of European ancestry from GTEx v8. We retained 8,592,339 autosomal variants with a MAF ≥ 1% for the genotype data and focused our analysis on the gene expression measurements from whole blood samples. For each gene, we performed single-variant regression analyses for variants that reside within 100kb upstream of the gene TSS and 100kb downstream of the gene TES, controlling for covariates that include the top five genotype PCs, 60 PEER factors, sequencing platform, sequencing protocol, and sex. Marginal z-scores of variants from these regression analyses were used for the subsequent analyses. Finally, we applied VINTAGE for gene-based analyses following a procedure similar to that with the eQTLGen data.

Finally, we assessed the computational time of VINTAGE in the real data and examined its variation in relation to the number of top eigenvalues and eigenvectors, k, used for the analysis of each gene region. For this evaluation, we grouped genes into bins based on the value of k, with a bin width of 10 eigenvalues and eigenvectors. Subsequently, we obtained the average computation time taken by VINTAGE to analyze the genes in each bin.

### Compared methods

We compared VINTAGE with existing methods for two types of tests: the gene-wise genetic variance test and the gene-wise genetic correlation test. In the case of the gene-wise genetic variance test, we compared VINTAGE with five existing methods, including two representative methods that can make use of summary statistics as input: SKAT^13^ and TWAS^8^, and their extensions: ACAT-O^20^, VC-TWAS^21^, and CoMM^22^. For SKAT, we focused on the unweighted linear kernel version and extended it to using GWAS summary statistics as input. The resulting test statistic can be expressed as 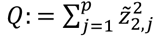 and its p-value can be evaluated using the Davies method^53^ with eigenvalues extracted from the LD matrix 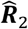. This version of SKAT is also equivalent to the method fastBAT^54^. For TWAS, we followed OTTERS (release v1.0.0)^55^ to first construct gene expression prediction models using various polygenic risk score (PRS) methods without LD clumping. These PRS methods use summary-level gene expression data as input and include P+T, lassosum, SDPR, and PRS-CS. The TWAS test statistic based on each PRS method can be expressed in the form of GWAS summary statistics as 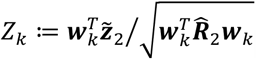, where ***w***_*k*_ are the expression prediction weights estimated by the *k*th PRS method. OTTERS converted the TWAS test statistics to p-values based on a standard normal distribution and then aggregated the p-values across different PRS methods into a single p-value using the Cauchy combination approach. For ACAT-O, we applied the STAAR package (version 0.9.7.1)^56^ for model fitting. ACAT-O is an omnibus test that aggregates p-values from six set-based tests, including ACAT-V, SKAT, and the burden test, each using two sets of SNP weights. For VC-TWAS, we applied SDPR to estimate the SNP effects on gene expression and constructed the test statistic in the form of a SKAT statistic, with SNP weights equal to the square of its effect on gene expression. VC-TWAS relaxes the assumption made in a conventional TWAS that SNP effects on a trait are scalar multiplication of SNP effects on gene expression, instead allowing for random SNP effects on the trait, with variance proportional to the corresponding SNP effects on gene expression. For CoMM, we used the CoMM package (version 0.1.0) and applied the particular version of CoMM (i.e., CoMM-S^4^) that can use summary statistics from both the gene expression study and GWAS. CoMM shares the same assumptions as TWAS but jointly fits the gene expression prediction model and the GReX-trait association model to account for uncertainly in gene expression prediction.

In the case of gene-wise genetic correlation test, we compared VINTAGE with SUPERGNOVA (release v1.0)^14^, LAVA (release v0.1.0)^15^, and PMR-Egger (version 1.0)^18^. For SUPERGNOVA, we followed its online tutorial to prepare the necessary input summary statistics files, LD reference files, and genome partition files. The genome partition files contain, for each gene, the boundary positions of the analyzed genomic regions defined in the Real data applications. For LAVA, we followed its online tutorial to process the input files, perform univariate tests of local heritability for both gene expression and the trait, and analyze the local genetic correlation between gene expression and the trait. In LAVA, we set the parameter “sample.overlap.file “ to NULL, as there is no sample overlap between the expression study and GWAS. For PMR-Egger, we followed its online tutorial to detect potentially mediating genes of traits after controlling for horizontal pleiotropy. PMR-Egger employs the Egger assumption to control for horizontal pleiotropy, effectively assuming that the horizontal pleiotropy effects are the same across all SNPs, similar to the burden test assumption for rare variant tests.

Additionally, we explored two recent TWAS fine-mapping methods – cTWAS (version 0.4.15) ^39^ and GIFT (version 1.0) ^40^ – in relation to our marginal gene-wise genetic variance and correlation tests. cTWAS carries out fine-mapping by including GReXs and genetic variants residing in a focal region into a Bayesian variable selection model. GIFT, on the other hand, builds upon a frequentist framework and carries out fine-mapping through conditional analyses. For both methods, we focused on LD blocks containing genes identified by the genetic variance tests of VINTAGE. For cTWAS, we followed its online tutorial to declare genes with a posterior inclusion probability (PIP) > 0.8 and is also included in credible sets to be potentially biologically relevant. For GIFT, we declared a gene to be potentially biologically relevant based on a Bonferroni-corrected p-value threshold.

## Data availability

The individual-level genotype data of UK Biobank is available at http://www.ukbiobank.ac.uk under the application number 98971. The GWAS summary statistics (V3) for eighteen complex traits from UK Biobank are publicly available at http://www.nealelab.is/uk-biobank. The cis-eQTL summary statistics from eQTLGen phase I study are available at https://www.eqtlgen.org/phase1.html. The individual-level genotype data of 1000 Genomes Project (1000G) phase 3 is available at https://www.internationalgenome.org. The Genotype-Tissue Expression (GTEx) Project data (v8) is available at the GTEx Portal (https://gtexportal.org/home/) and dbGaP under the accession number phs000424.v8. SNP RSID is reported under the build 156 according to dbSNP at https://www.ncbi.nlm.nih.gov/snp/. GENCODE (release 40) is available at https://www.gencodegenes.org/human/release_40lift37.html.

## Code availability

We used R (version 4.4.0) for statistical analysis. VINTAGE (version 0.1.005) is implemented as an R package with underlying efficient C++ codes interfaced through Rcpp and is freely available at https://github.com/zhengli09/VINTAGE. The unweighted linear kernel version of SKAT is implemented within VINTAGE available at https://github.com/zhengli09/VINTAGE. OTTERS (version 1.0.0) is available at https://github.com/daiqile96/OTTERS. SUPERGNOVA (version 1.0) is available at https://github.com/qlu-lab/SUPERGNOVA. LAVA (version 0.1.0) is available at https://github.com/josefin-werme/LAVA. susieR (version 0.12.41) is available at https://github.com/stephenslab/susieR. Processing of plink bed/fam/bim files was conducted using PLINK (version 2.00 alpha) available at https://www.cog-genomics.org/plink/2.0/. STAAR (version 0.9.7.1) is available at https://github.com/xihaoli/STAAR. VC-TWAS is available at https://github.com/yanglab-emory/TIGAR. CoMM (version 0.1.0) is available at https://github.com/gordonliu810822/CoMM. PMR-Egger (version 1.0) is available at https://github.com/yuanzhongshang/PMR. cTWAS (version 0.4.15) is available at https://github.com/xinhe-lab/multigroup_ctwas. GIFT (version 1.0) is available at https://github.com/yuanzhongshang/GIFT.

## Acknowledgements

This study was supported by the National Institutes of Health (NIH) Grants R01HG009124 and R01GM144960 (to X.Z.). The funders had no role in study design, data collection and analysis, decision to publish or preparation of the manuscript. This study has been conducted using UK Biobank resource under Application Number 98971. UK Biobank was established by the Wellcome Trust medical charity, Medical Research Council, Department of Health, Scottish Government and the Northwest Regional Development Agency. It has also had funding from the Welsh Assembly Government, British Heart Foundation and Diabetes UK. The UK Biobank resource was approved by the UK Biobank Research Ethics Committee and all participants provided written informed consent to participate. The Genotype-Tissue Expression (GTEx) Project was supported by the Common Fund of the Office of the Director of the National Institutes of Health, and by NCI, NHGRI, NHLBI, NIDA, NIMH, and NINDS.

## Ethics statement

This study was approved by the ethics committee of the University of Michigan Institutional Review Board (IRB) HUM00156494.

## Author contributions

X.Z. conceived the idea and provided funding support. Z.L. and X.Z. designed the experiments. Z.L. developed the method and implemented the software. Z.L. performed simulations and analyzed real data. Z.L. and X.Z. wrote the manuscript with input from B.G. All authors read and approved the final manuscript.

