## Supplemental Figures for "An alternative framework for transcriptome-wide association studies to detect and decipher gene-trait associations"

### Supplementary Figures

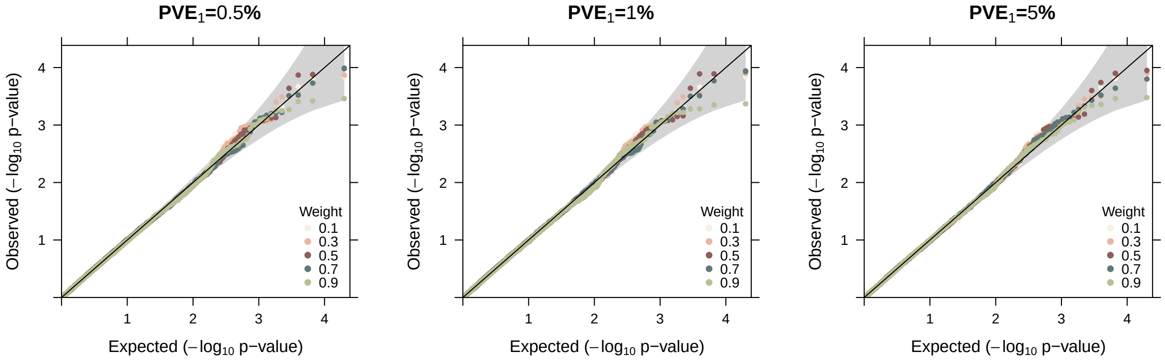


**Figure S1. Quantile-quantile plots of -log$_{10}$ p-values from the remaining individual tests for gene-wise genetic variance test under the null simulations.** Evaluated tests, $T_w$, is each constructed under a different weight value of $w$ that ranges from 0.1 to 0.9 with an increment of 0.2, where the weight controls the relative contributions from the variance and covariance component scores. Null simulations were conducted under different $PVE_1$ values (0.5%, 1%, and 5%) with in-sample LD matrices and a polygenic genetic architecture ($q = p$).

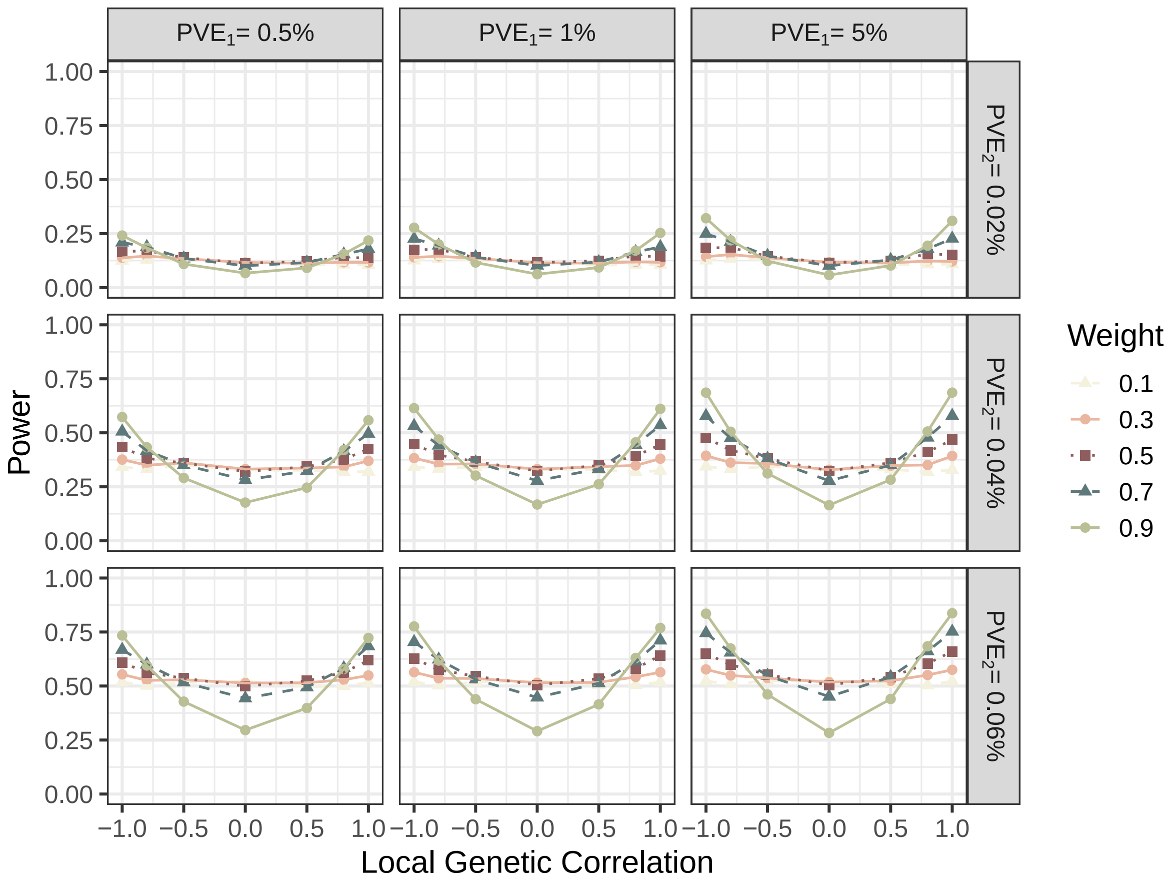


**Figure S2. Power of the remaining individual tests for gene-wise genetic variance test in the simulations.** Evaluated tests, $T_w$, is each constructed under a different weight value of $w$ that ranges from 0.1 to 0.9 with an increment of 0.2, where the weight controls the relative contributions from the variance and covariance component scores. Simulations were conducted under different $PVE_1$ values (0.5%, 1%, and 5%), different $PVE_2$ values (0.02%, 0.04%, and 0.06%), and different local genetic correlations (-1.0, -0.8, -0.5, 0.0, 0.5, 0.8, 1.0) with in-sample LD matrices and a polygenic genetic architecture ($q = p$).

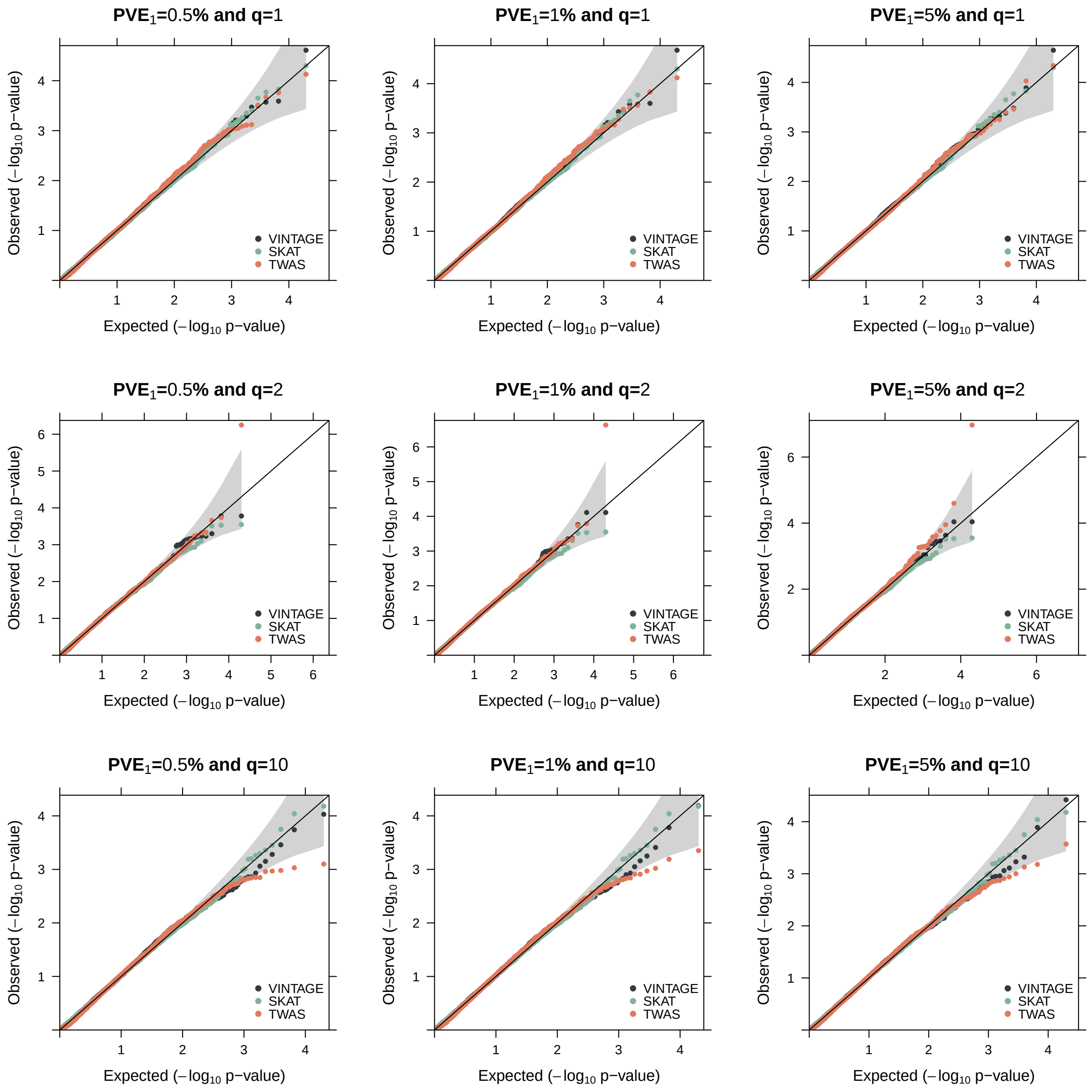


**Figure S3. Quantile-quantile plots of -$\log_{10}$ p-values from different methods for gene-wise genetic variance test in the null simulations.** The null simulations were conducted under sparse genetic architectures with (**Top**) $q = 1$, (**Middle**) $q = 2$, or (**Bottom**) $q = 10$ SNPs that have non-zero genetic effects on both gene expression and trait. Evaluated methods include VINTAGE, SKAT with an unweighted linear kernel, and TWAS that adapts multiple PRS methods to estimate gene expression prediction weights and performs an omnibus test. The null simulations were also conducted under different $PVE_1$ values (Left: 0.5%, Middle: 1%, and Right: 5%) with in-sample LD matrices.

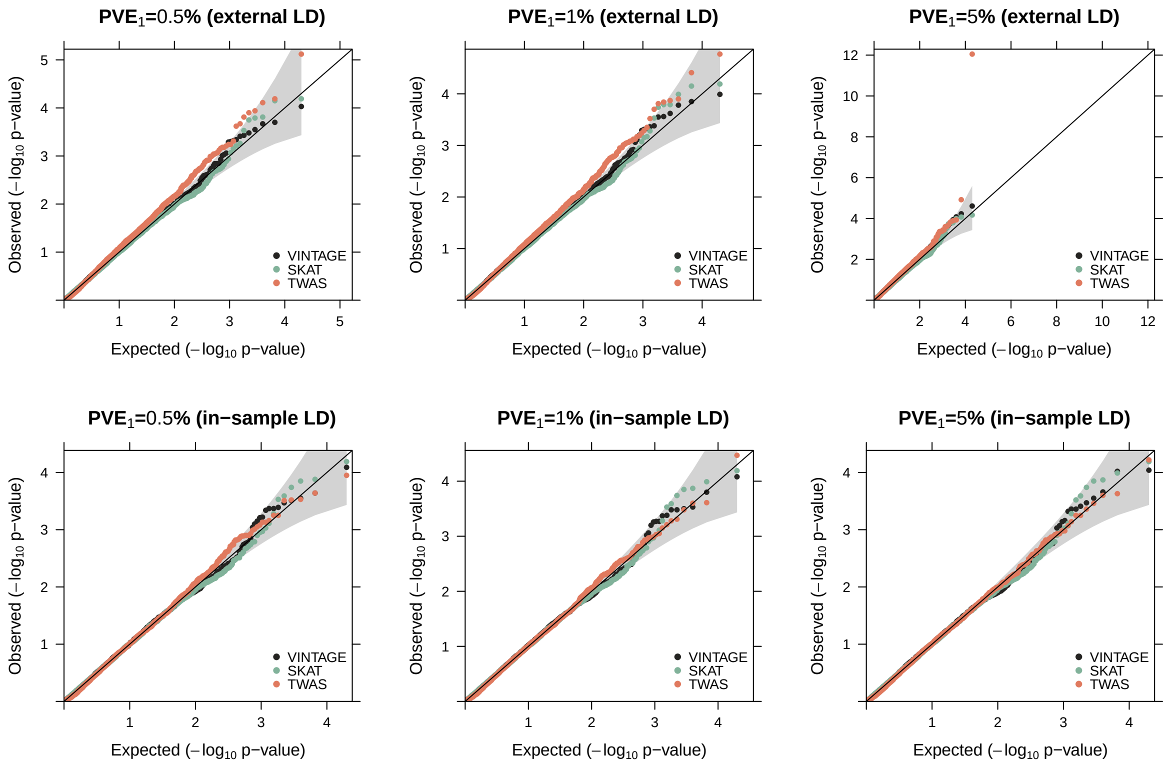


**Figure S4. Quantile-quantile plots of -$\log_{10}$ p-values from different methods for gene-wise genetic variance test in the null simulations.** The null simulations were conducted with (**Top**) external LD matrices or (**Bottom**) in-sample LD matrices. The in-sample LD matrices were computed using the same set of 10,000 UKBB WB individuals for simulating the gene expression data and the external LD matrices were computed using the 503 European individuals from the 1000 Genomes Project. Compared methods include VINTAGE, SKAT with an unweighted linear kernel, and TWAS that adapts multiple PRS methods to estimate gene expression prediction weights and performs an omnibus test. The null simulations were also conducted under different $PVE_1$ values (Left: 0.5%, Middle: 1%, and Right: 5%) and a polygenic genetic architecture ($q = p$).

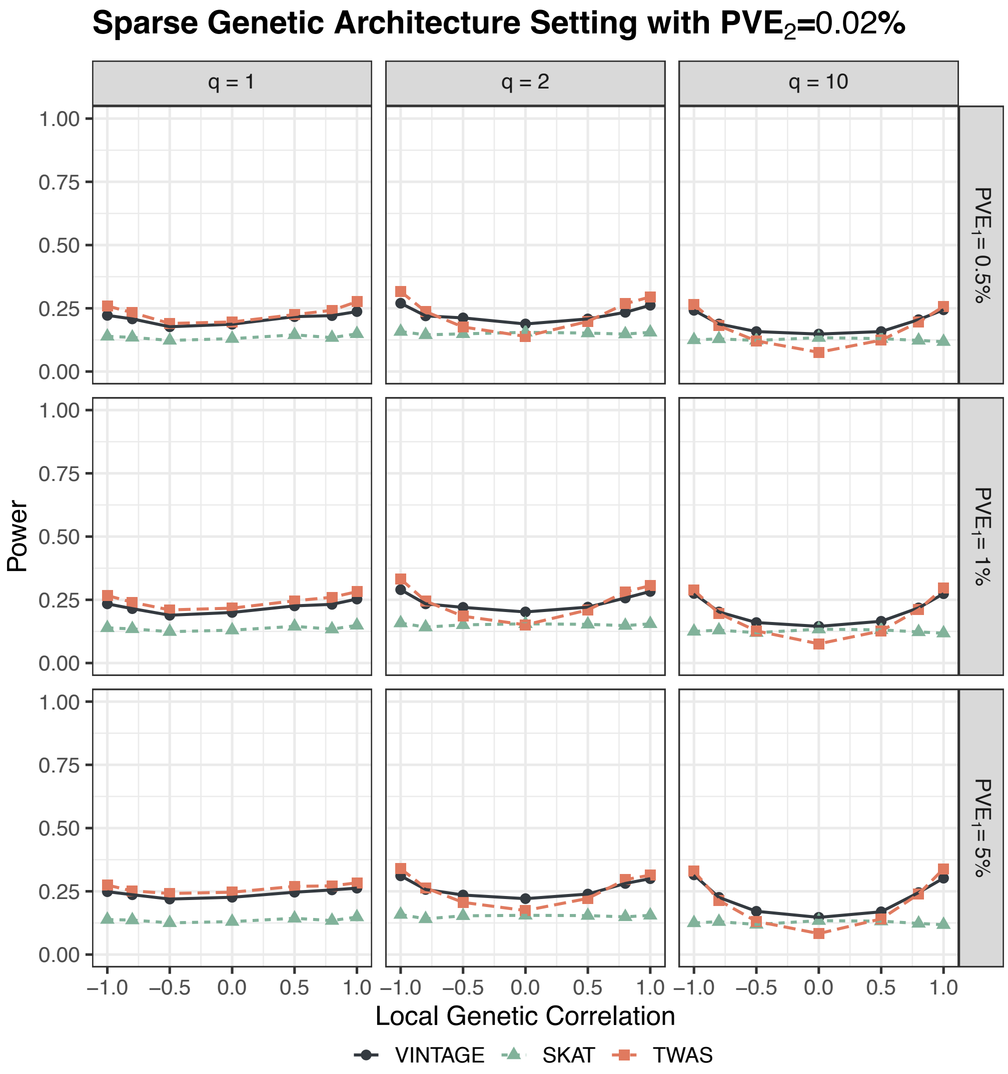


**Figure S5. Power of different methods for gene-wise genetic variance test in the simulations.** The simulations were conducted under sparse genetic architectures with (**Left**) $q = 1$, (**Middle**) $q = 2$, or (**Right**) $q = 10$ SNPs that have non-zero genetic effects on both gene expression and trait. Compared methods include VINTAGE, SKAT with an unweighted linear kernel, and TWAS that adapts multiple PRS methods to estimate gene expression prediction weights and performs an omnibus test. The simulations were also conducted under different $PVE_1$ values (Top: 0.5%, Middle: 1%, and Bottom: 5%) and different local genetic correlations (-1.0, -0.8, -0.5, 0.0, 0.5, 0.8, 1.0) with in-sample LD matrices and a **relatively low** $PVE_2$ **(0.02%)**.

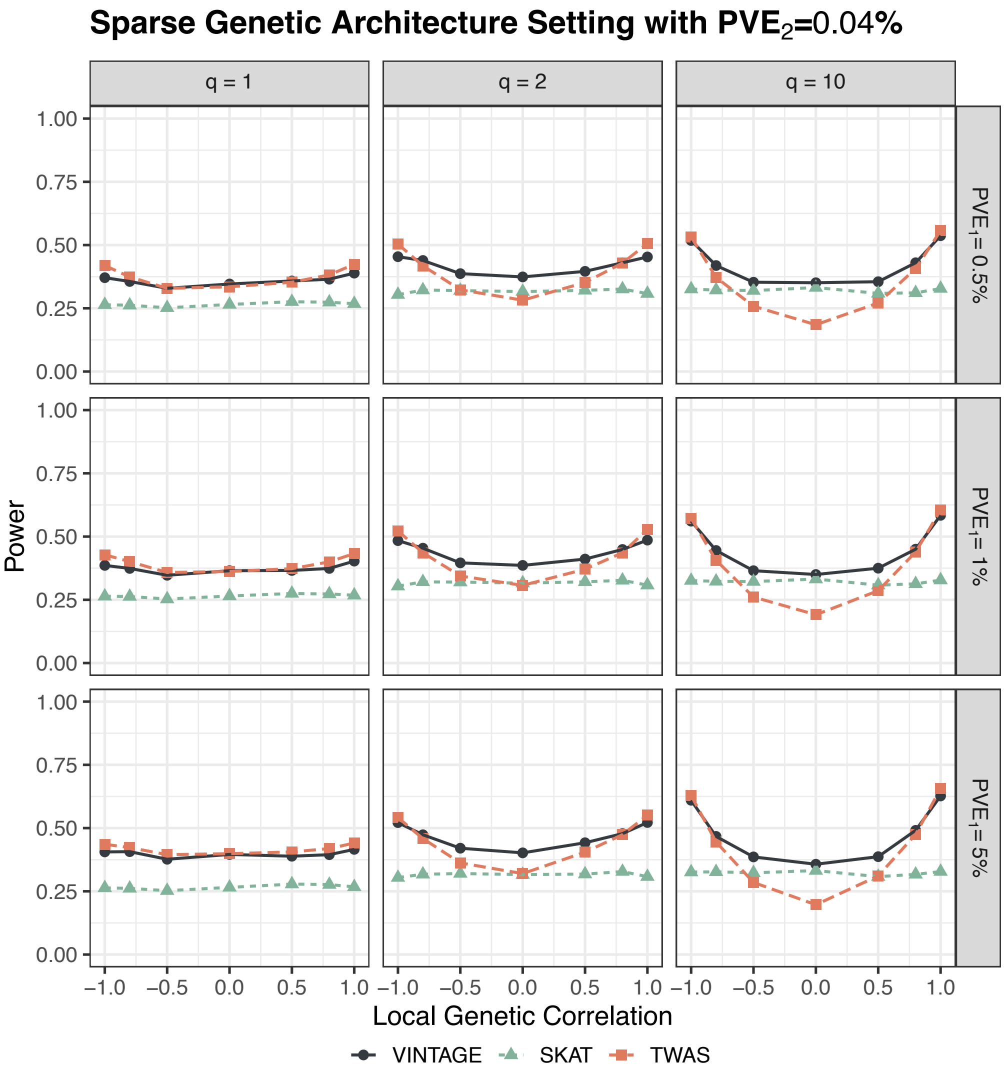


**Figure S6. Power of different methods for gene-wise genetic variance test in the simulations.** The simulations were conducted under sparse genetic architectures with (**Left**) $q = 1$, (**Middle**) $q = 2$, or (**Right**) $q = 10$ SNPs that have non-zero genetic effects on both gene expression and trait. Compared methods include VINTAGE, SKAT with an unweighted linear kernel, and TWAS that adapts multiple PRS methods to estimate gene expression prediction weights and performs an omnibus test. The simulations were also conducted under different $PVE_1$ values (Top: 0.5%, Middle: 1%, and Bottom: 5%) and different local genetic correlations (-1.0, -0.8, -0.5, 0.0, 0.5, 0.8, 1.0) with in-sample LD matrices and a **relatively moderate** $PVE_2$ **(0.04%)**.

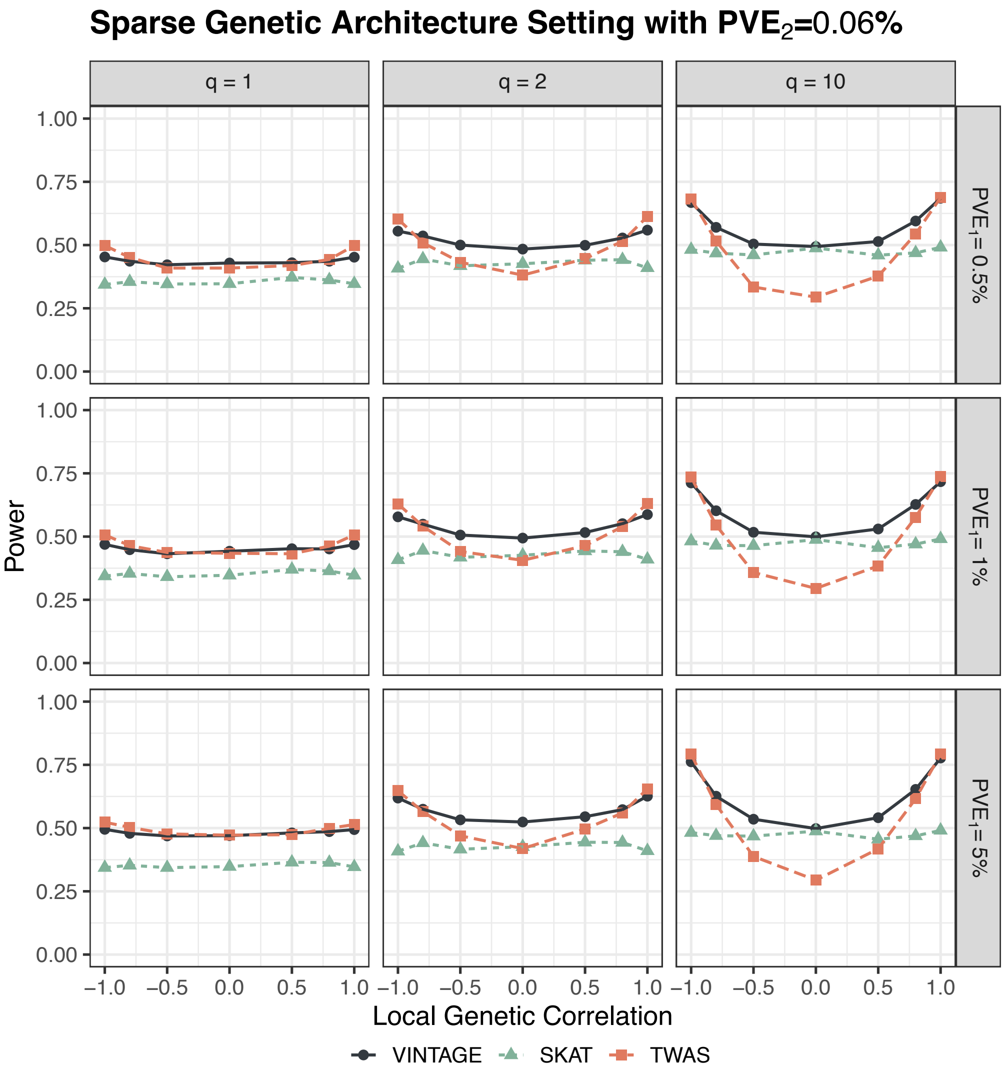


**Figure S7. Power of different methods for gene-wise genetic variance test in the simulations.** The simulations were conducted under sparse genetic architectures with (**Left**) $q = 1$, (**Middle**) $q = 2$, or (**Right**) $q = 10$ SNPs that have non-zero genetic effects on both gene expression and trait. Compared methods include VINTAGE, SKAT with an unweighted linear kernel, and TWAS that adapts multiple PRS methods to estimate gene expression prediction weights and performs an omnibus test. The simulations were also conducted under different $PVE_1$ values (Top: 0.5%, Middle: 1%, and Bottom: 5%) and different local genetic correlations (-1.0, -0.8, -0.5, 0.0, 0.5, 0.8, 1.0) with in-sample LD matrices and a **relatively high** $PVE_2$ **(0.06%)**.

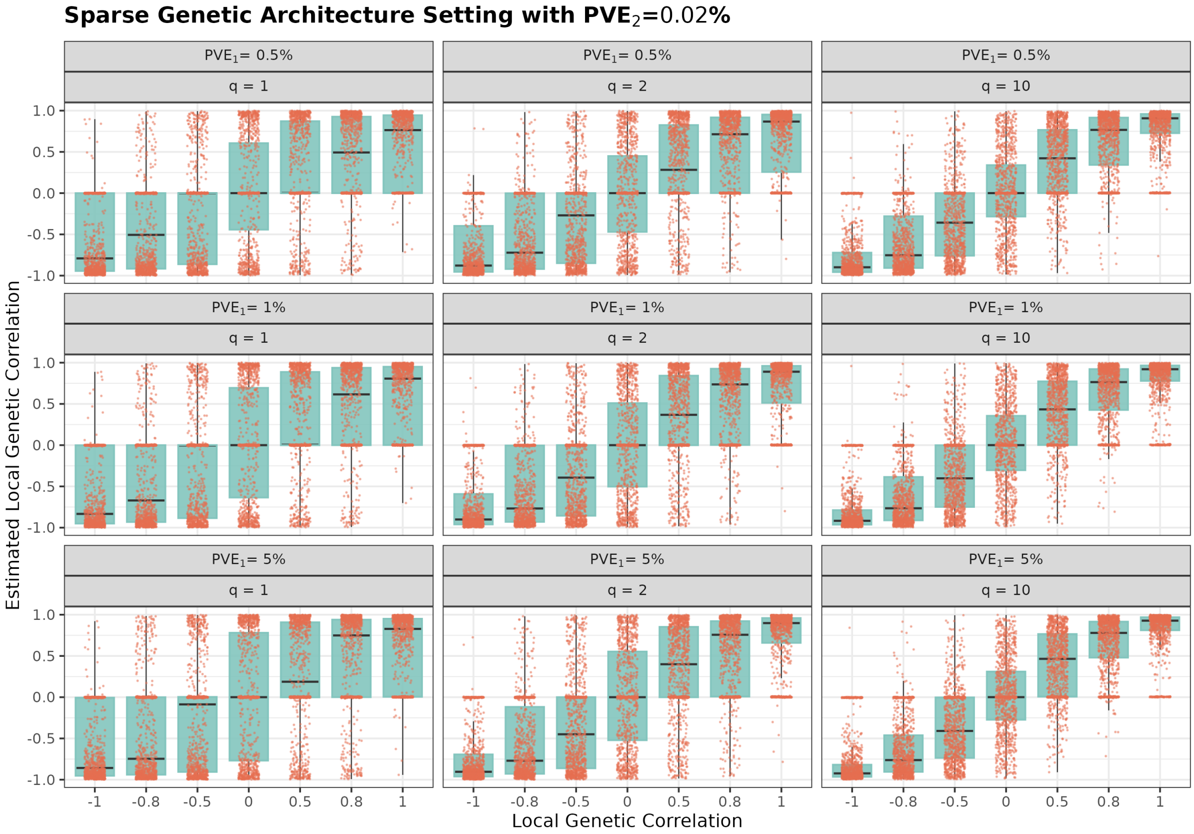


**Figure S8. Estimation accuracy of local genetic correlations under sparse genetic architectures.** Boxplots indicate the estimated local genetic correlations from VINTAGE in the simulations. These simulations were conducted under sparse genetic architectures with (**Left**) $q = 1$, (**Middle**) $q = 2$, or (**Right**) $q = 10$ SNPs that have non-zero genetic effects on both gene expression and trait. The simulations were also conducted under different $PVE_1$ values (Top: 0.5%, Middle: 1%, and Bottom: 5%) and different local genetic correlations (-1.0, -0.8, -0.5, 0.0, 0.5, 0.8, 1.0) with in-sample LD matrices and a **relatively low** $PVE_2$ **(0.02%)**.

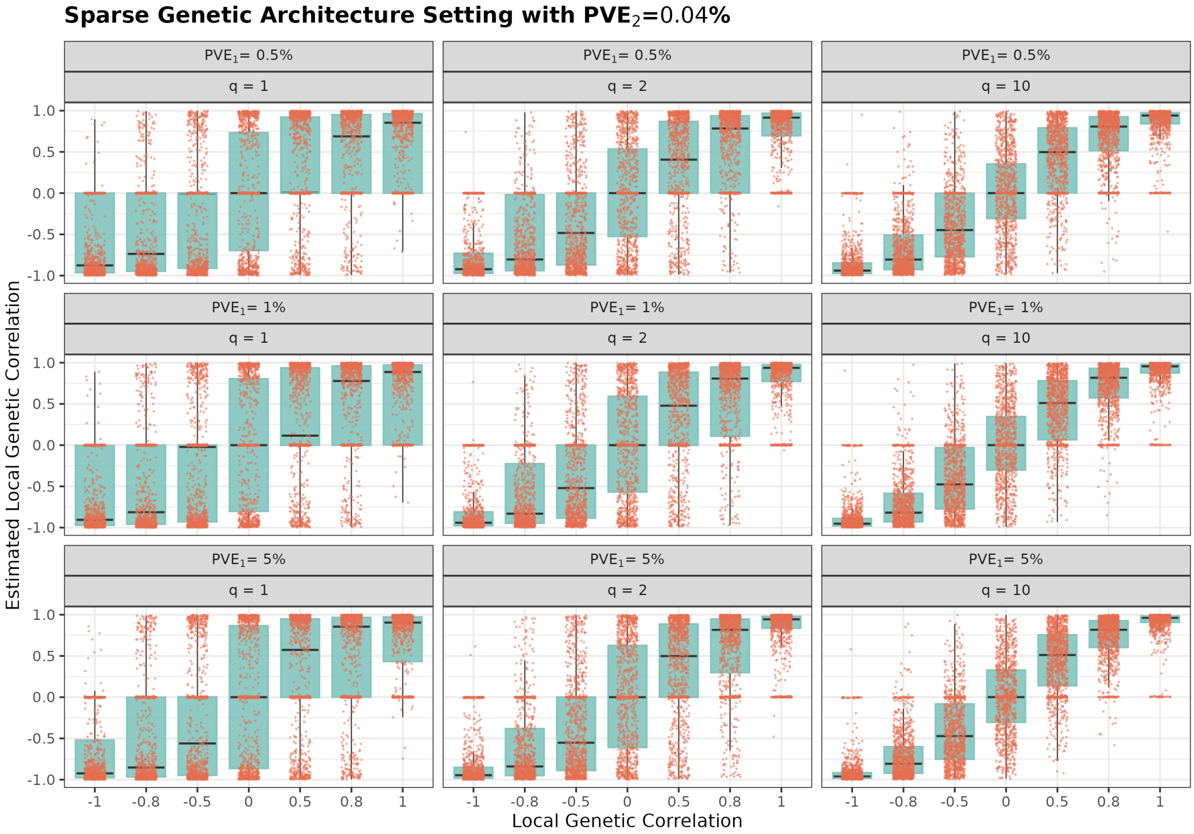


**Figure S9. Estimation accuracy of local genetic correlations under sparse genetic architectures.** Boxplots indicate the estimated local genetic correlations from VINTAGE in the simulations. These simulations were conducted under sparse genetic architectures with (**Left**) $q = 1$, (**Middle**) $q = 2$, or (**Right**) $q = 10$ SNPs that have non-zero genetic effects on both gene expression and trait. The simulations were also conducted under different $PVE_1$ values (Top: 0.5%, Middle: 1%, and Bottom: 5%) and different local genetic correlations (-1.0, -0.8, -0.5, 0.0, 0.5, 0.8, 1.0) with in-sample LD matrices and a **relatively moderate** $PVE_2$ **(0.04%)**.

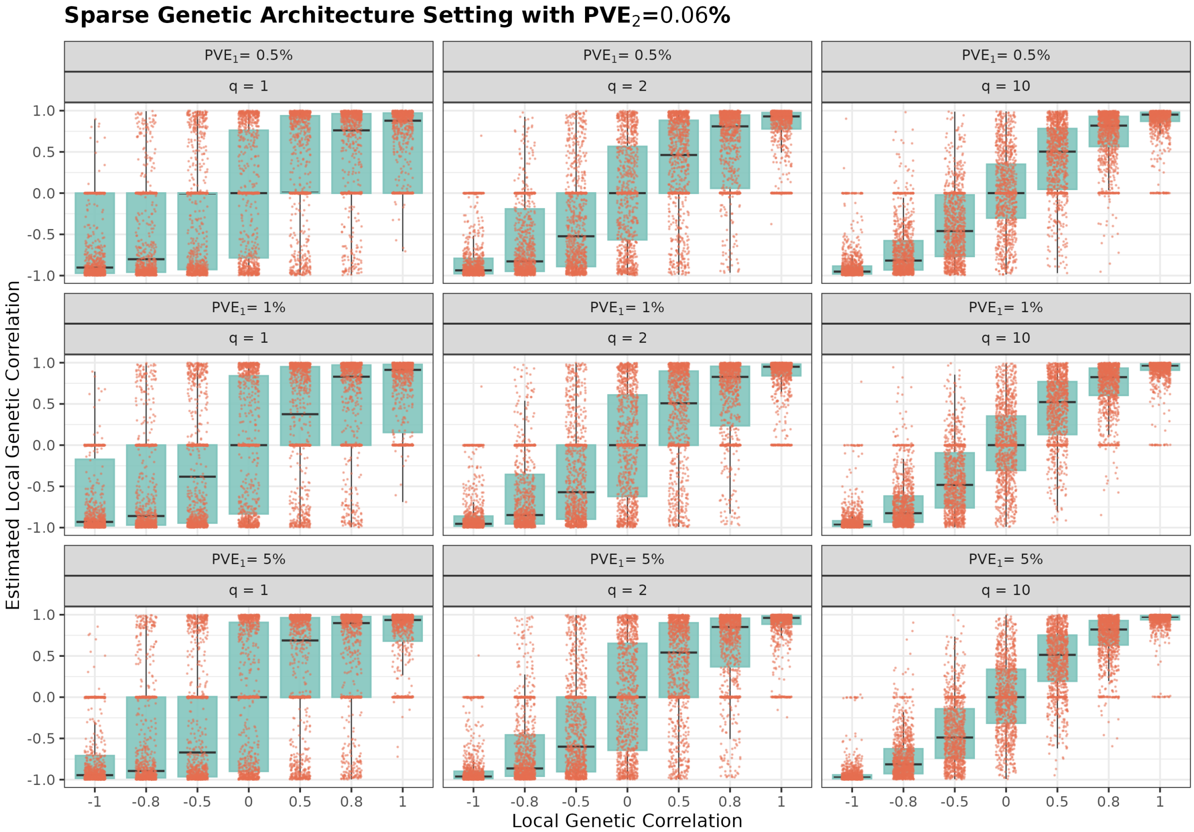


**Figure S10. Estimation accuracy of local genetic correlations under sparse genetic architectures.** Boxplots indicate the estimated local genetic correlations from VINTAGE in the simulations. These simulations were conducted under sparse genetic architectures with (**Left**) $q = 1$, (**Middle**) $q = 2$, or (**Right**) $q = 10$ SNPs that have non-zero genetic effects on both gene expression and trait. The simulations were also conducted under different $PVE_1$ values (Top: 0.5%, Middle: 1%, and Bottom: 5%) and different local genetic correlations (-1.0, -0.8, -0.5, 0.0, 0.5, 0.8, 1.0) with in-sample LD matrices and a **relatively high** $PVE_2$ **(0.06%)**.

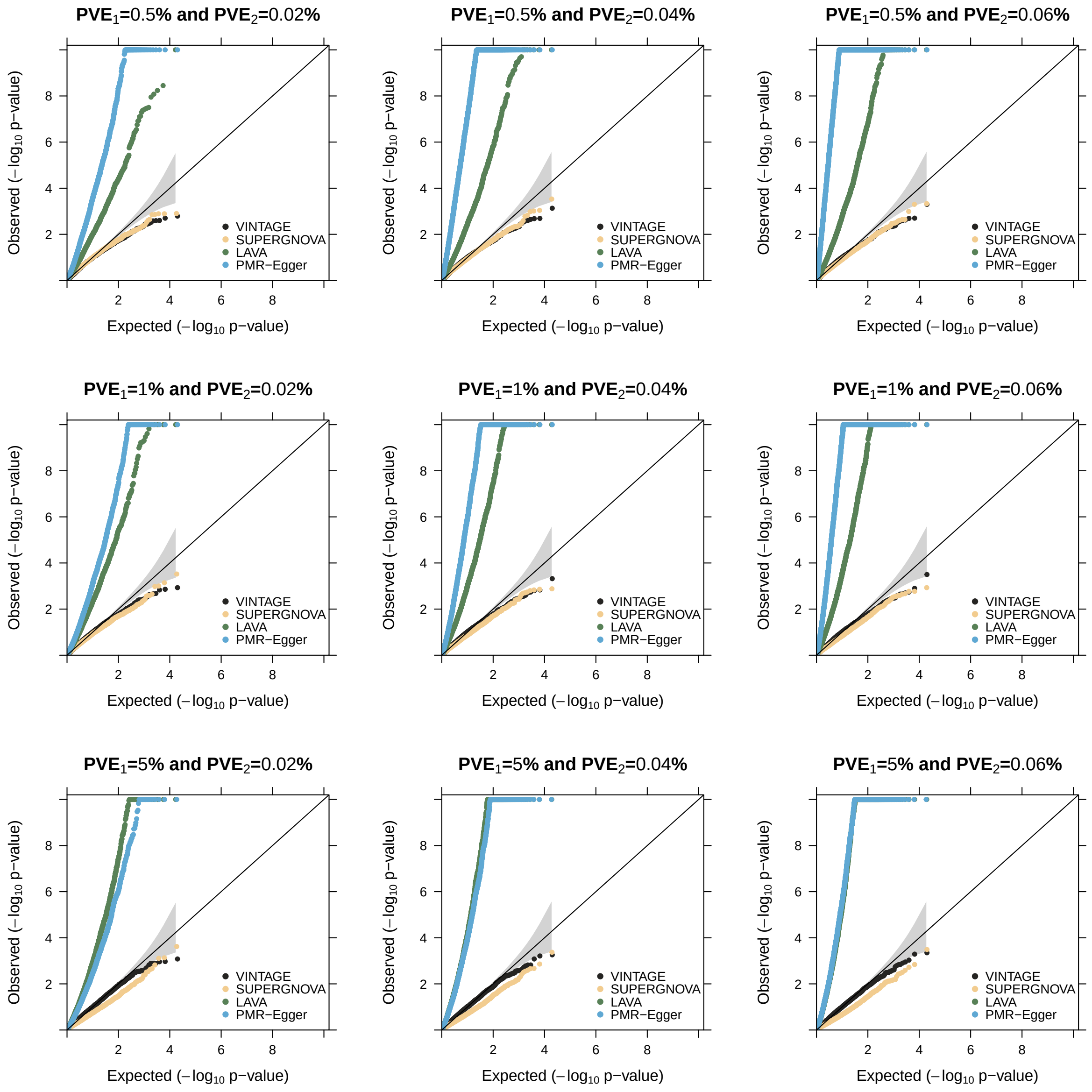


**Figure S11. Quantile-quantile plots of -log$_{10}$ p-values from different methods for gene-wise genetic correlation test in the null simulations.** Evaluated methods include VINTAGE, SUPERGNOVA, LAVA, and PMR-Egger. The null simulations were conducted under different $PVE_1$ values (Top: 0.5%, Middle: 1%, and Bottom: 5%) and different $PVE_2$ values (Left: 0.02%, Middle: 0.04%, and Right: 0.06%) with in-sample LD matrices and a polygenic genetic architecture ($q = p$). Simulation replicates with negative heritability estimates or negative variance estimates from SUPERGNOVA or LAVA were excluded from the evaluation.

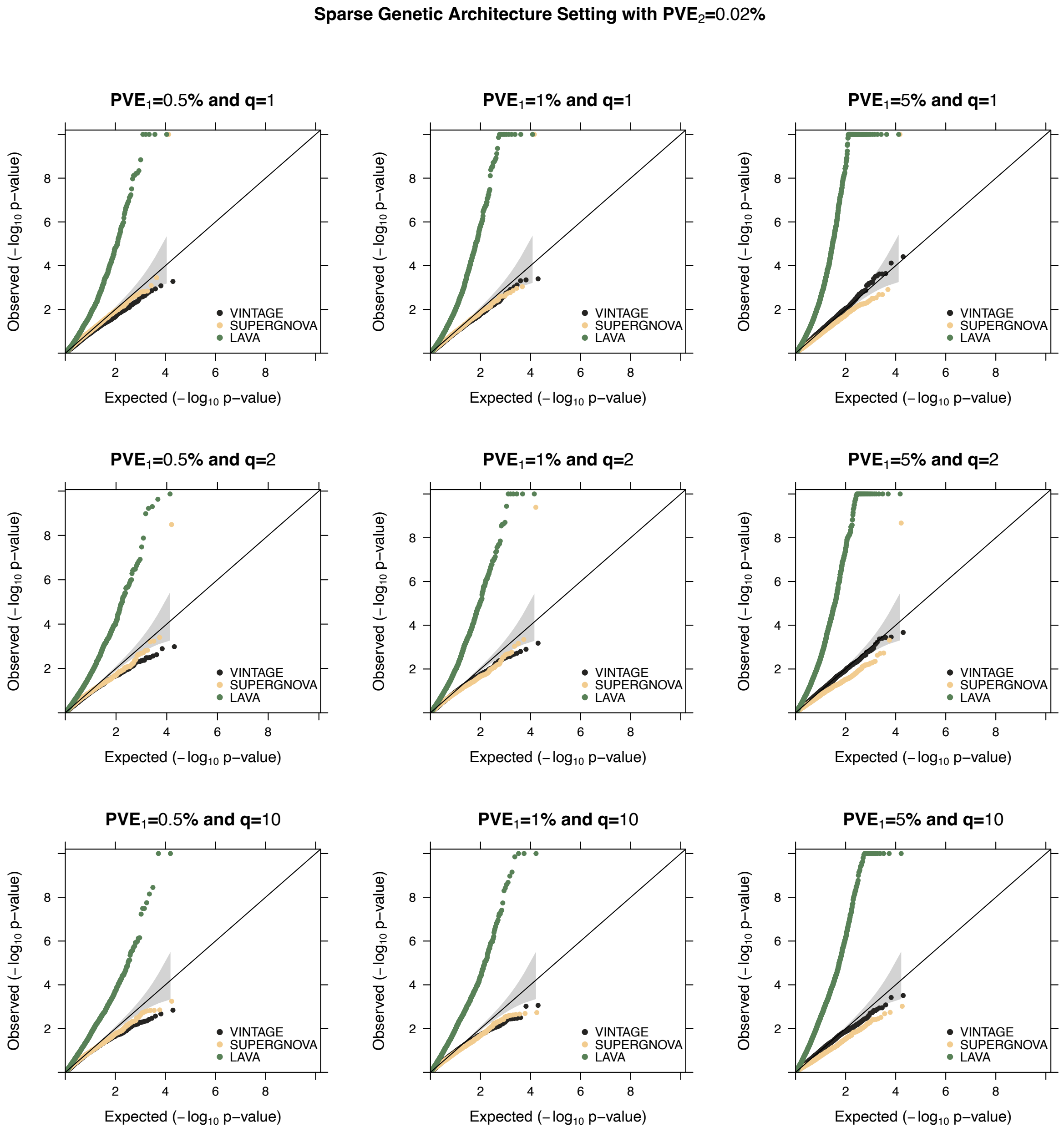


**Figure S12. Quantile-quantile plots of -log$_{10}$ p-values from different methods for gene-wise genetic correlation test in the null simulations.** The null simulations were conducted under sparse genetic architectures with (**Top**) $q = 1$, (**Middle**) $q = 2$, or (**Bottom**) $q = 10$ SNPs that have non-zero genetic effects on gene expression and trait. To create proper null sparse settings for the genetic correlation test, we first randomly selected $q$ SNPs to have non-zero effects on gene expression. Among the remaining SNPs in the region that are not in extreme high LD with the selected SNPs ($R^2 < 0.95$), we randomly selected another $q$ SNPs to have non-zero effects on the trait. Evaluated methods include VINTAGE, SUPERGNOVA, and LAVA. The null simulations were also conducted under different $PVE_1$ values (Left: 0.5%, Middle: 1%, and Right: 5%) with in-sample LD matrices and a **relatively low** $PVE_2$ **(0.02%)**.

### Sparse Genetic Architecture Setting with $PVE_2$=0.04%

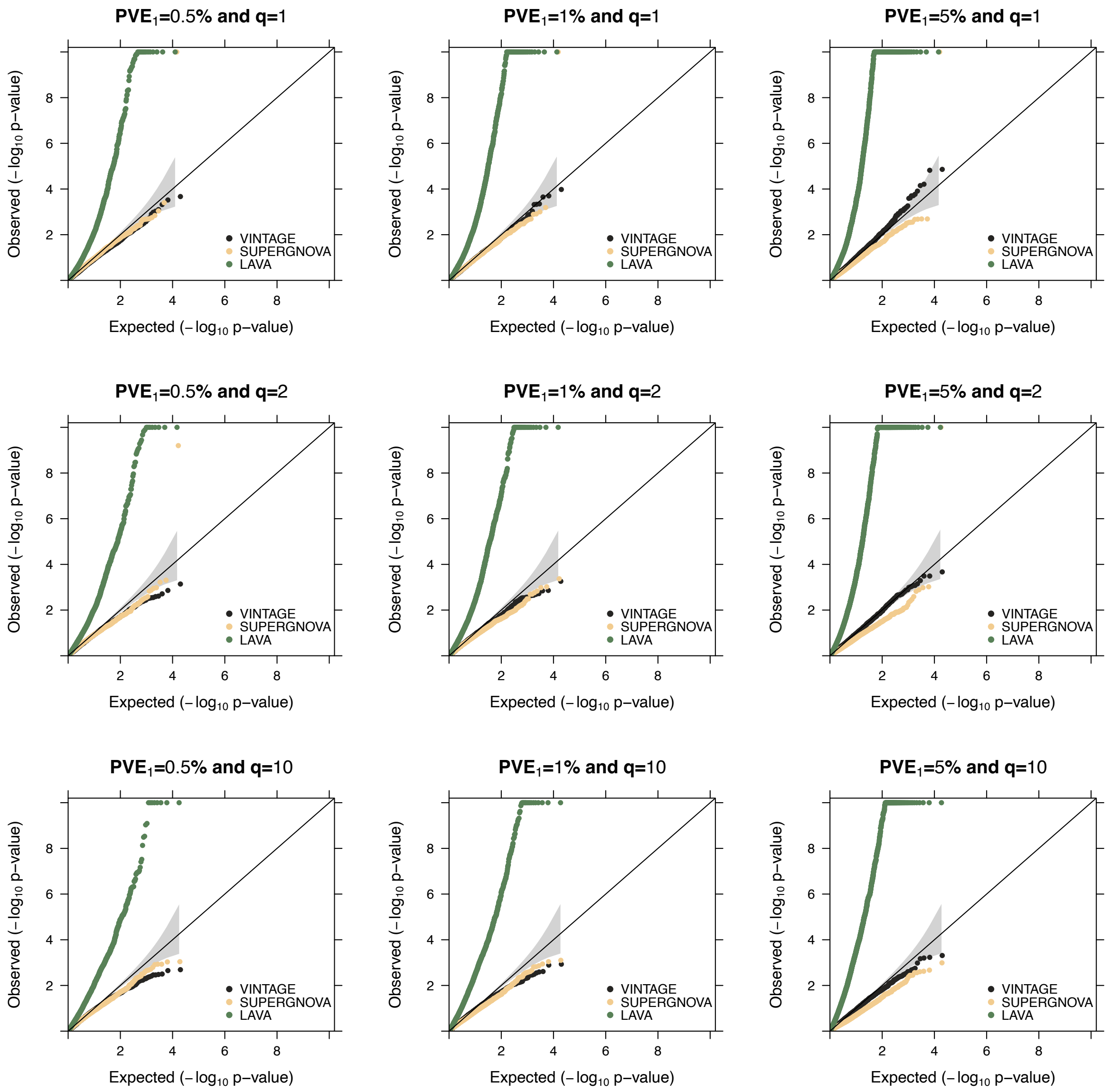


**Figure S13. Quantile-quantile plots of -log$_{10}$ p-values from different methods for gene-wise genetic correlation test in the null simulations.** The null simulations were conducted under sparse genetic architectures with (**Top**) $q = 1$, (**Middle**) $q = 2$, or (**Bottom**) $q = 10$ SNPs that have non-zero genetic effects on gene expression and trait. To create proper null sparse settings for the genetic correlation test, we first randomly selected $q$ SNPs to have non-zero effects on gene expression. Among the remaining SNPs in the region that are not in extreme high LD with the selected SNPs ($R^2 < 0.95$), we randomly selected another $q$ SNPs to have non-zero effects on the trait. Evaluated methods include VINTAGE, SUPERGNOVA, and LAVA. The null simulations were also conducted under different $PVE_1$ values (Left: 0.5%, Middle: 1%, and Right: 5%) with in-sample LD matrices and a **relatively moderate** $PVE_2$ **(0.04%)**.

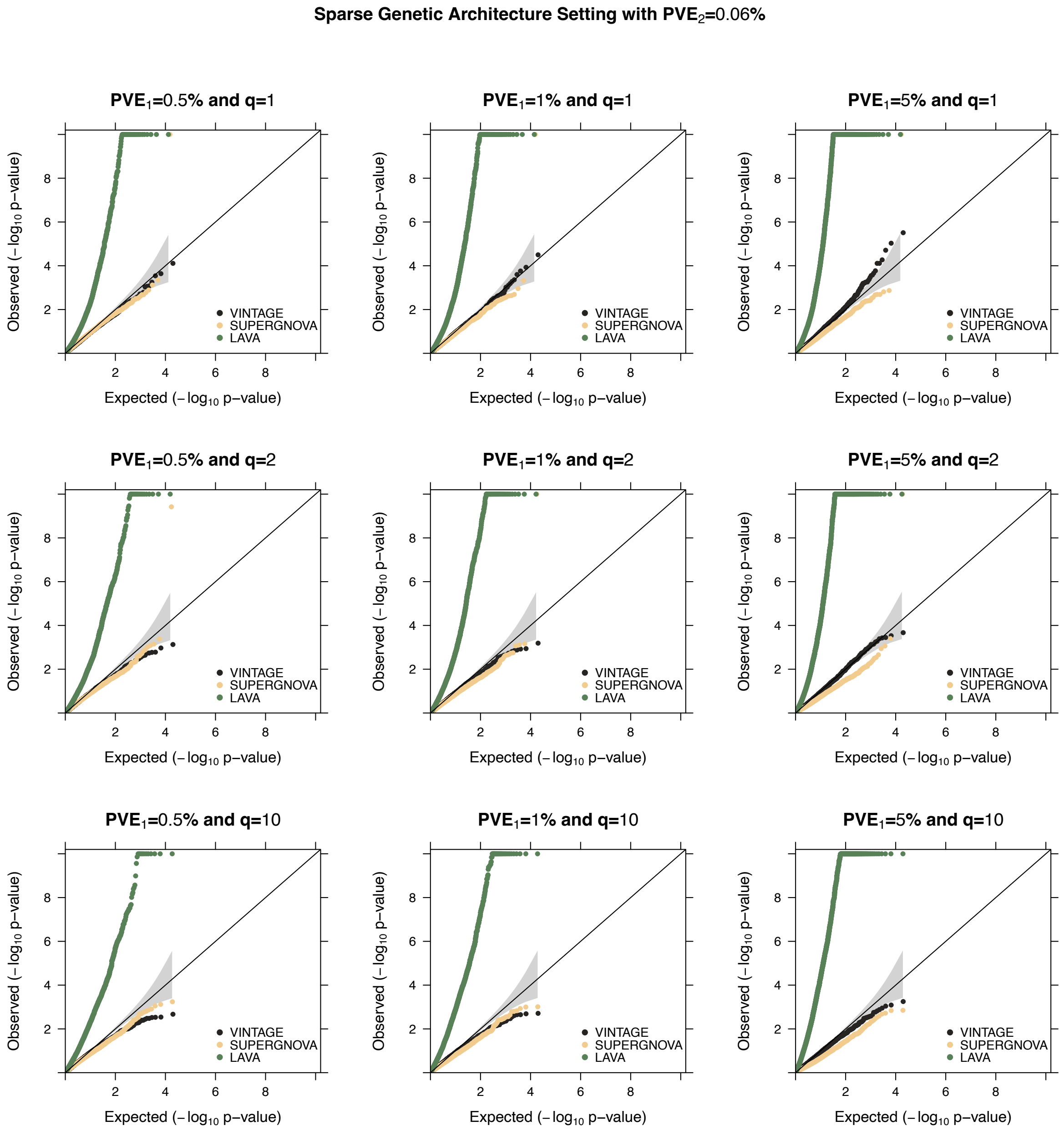


**Figure S14. Quantile-quantile plots of -log$_{10}$ p-values from different methods for gene-wise genetic correlation test in the null simulations.** The null simulations were conducted under sparse genetic architectures with (**Top**) $q = 1$, (**Middle**) $q = 2$, or (**Bottom**) $q = 10$ SNPs that have non-zero genetic effects on gene expression and trait. To create proper null sparse settings for the genetic correlation test, we first randomly selected $q$ SNPs to have non-zero effects on gene expression. Among the remaining SNPs in the region that are not in extreme high LD with the selected SNPs ($R^2 < 0.95$), we randomly selected another $q$ SNPs to have non-zero effects on the trait. Evaluated methods include VINTAGE, SUPERGNOVA, and LAVA. The null simulations were also conducted under different $PVE_1$ values (Left: 0.5%, Middle: 1%, and Right: 5%) with in-sample LD matrices and a **relatively high** $PVE_2$ **(0.06%)**.

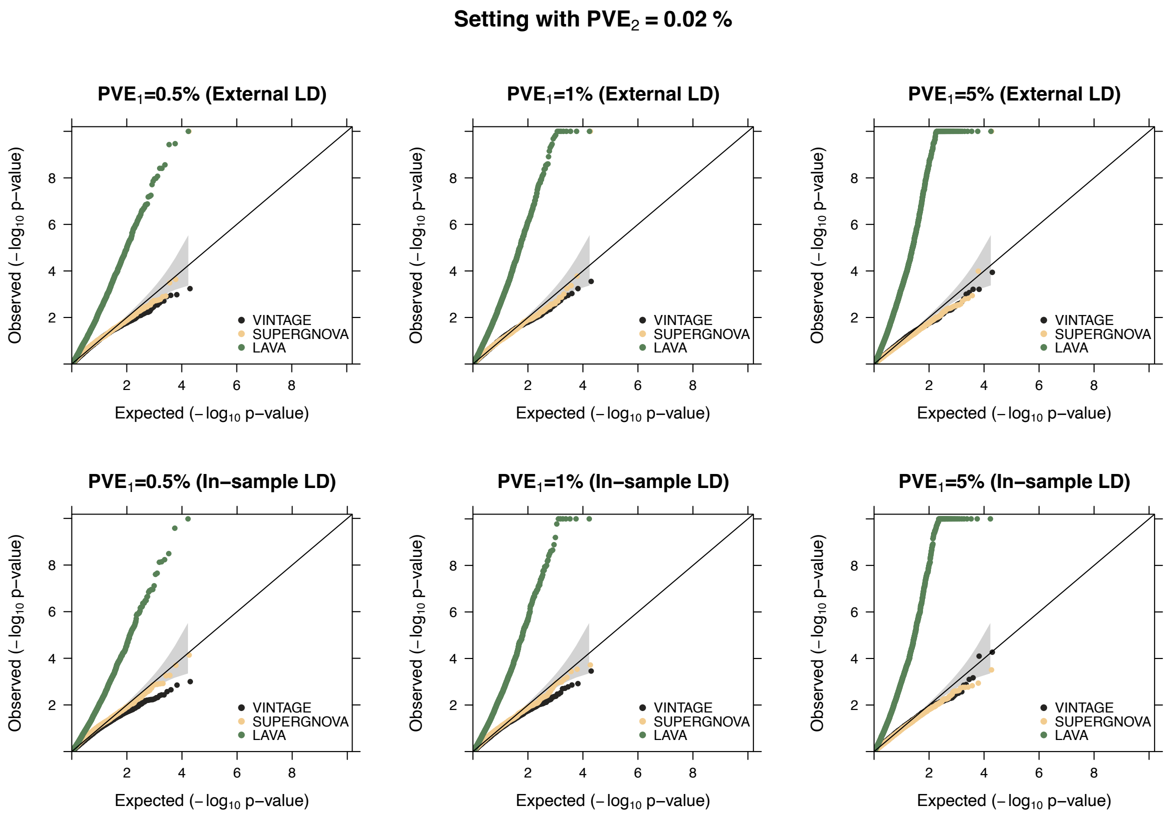


**Figure S15. Quantile-quantile plots of -$\log_{10}$ p-values from different methods for gene-wise genetic correlation test in the null simulations.** The null simulations were conducted with (**Top**) external LD matrices or (**Bottom**) in-sample LD matrices. The in-sample LD matrices were computed using the same set of 10,000 UKBB WB individuals for simulating the gene expression data while the external LD matrices were computed using the 503 European individuals from the 1000 Genomes Project. Compared methods include VINTAGE, SUPERGNOVA, and LAVA. The null simulations were also conducted under different $PVE_1$ values (Left: 0.5%, Middle: 1%, and Right: 5%), a polygenic genetic architecture ($q = p$), and a **relatively low** $PVE_2$ **(0.02%)**.

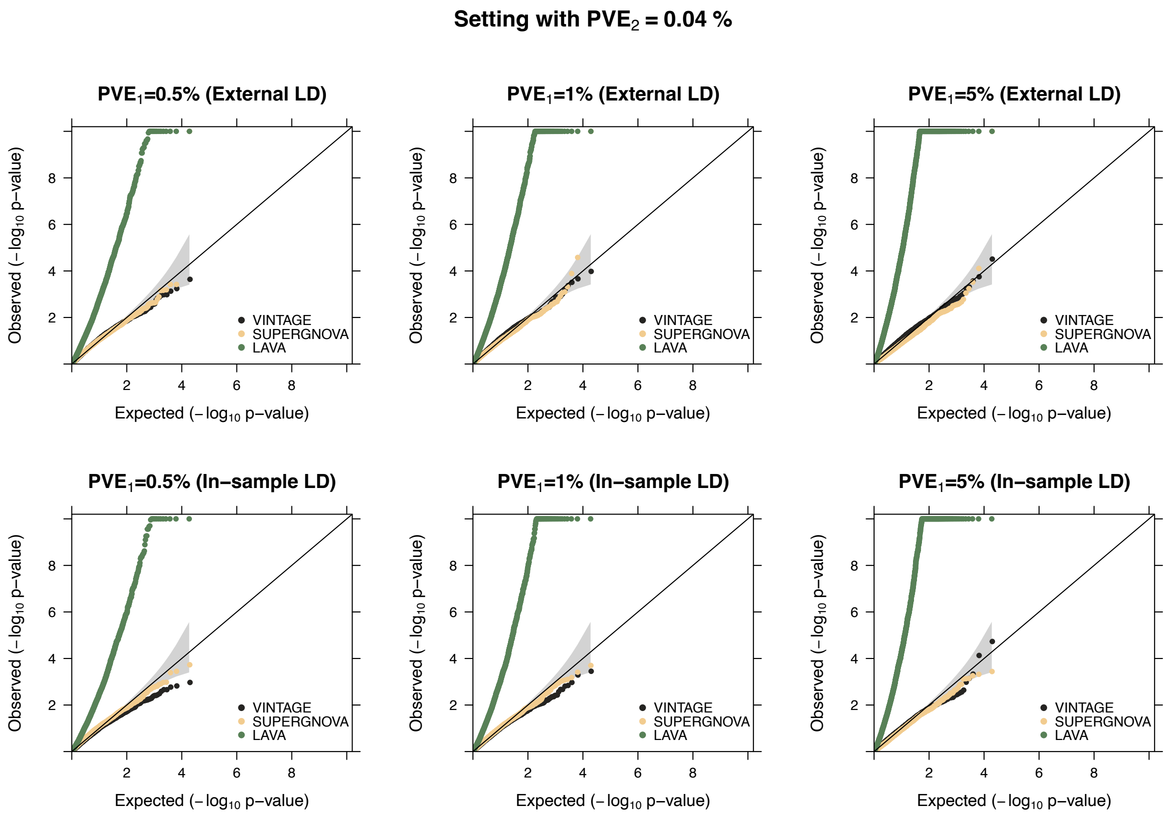


**Figure S16. Quantile-quantile plots of -log$_{10}$ p-values from different methods for gene-wise genetic correlation test in the null simulations.** The null simulations were conducted with (**Top**) external LD matrices or (**Bottom**) in-sample LD matrices. The in-sample LD matrices were computed using the same set of 10,000 UKBB WB individuals for simulating the gene expression data while the external LD matrices were computed using the 503 European individuals from the 1000 Genomes Project. Compared methods include VINTAGE, SUPERGNOVA, and LAVA. The null simulations were also conducted under different $PVE_1$ values (Left: 0.5%, Middle: 1%, and Right: 5%), a polygenic genetic architecture ($q = p$), and a **relatively moderate** $PVE_2$ **(0.04%)**.

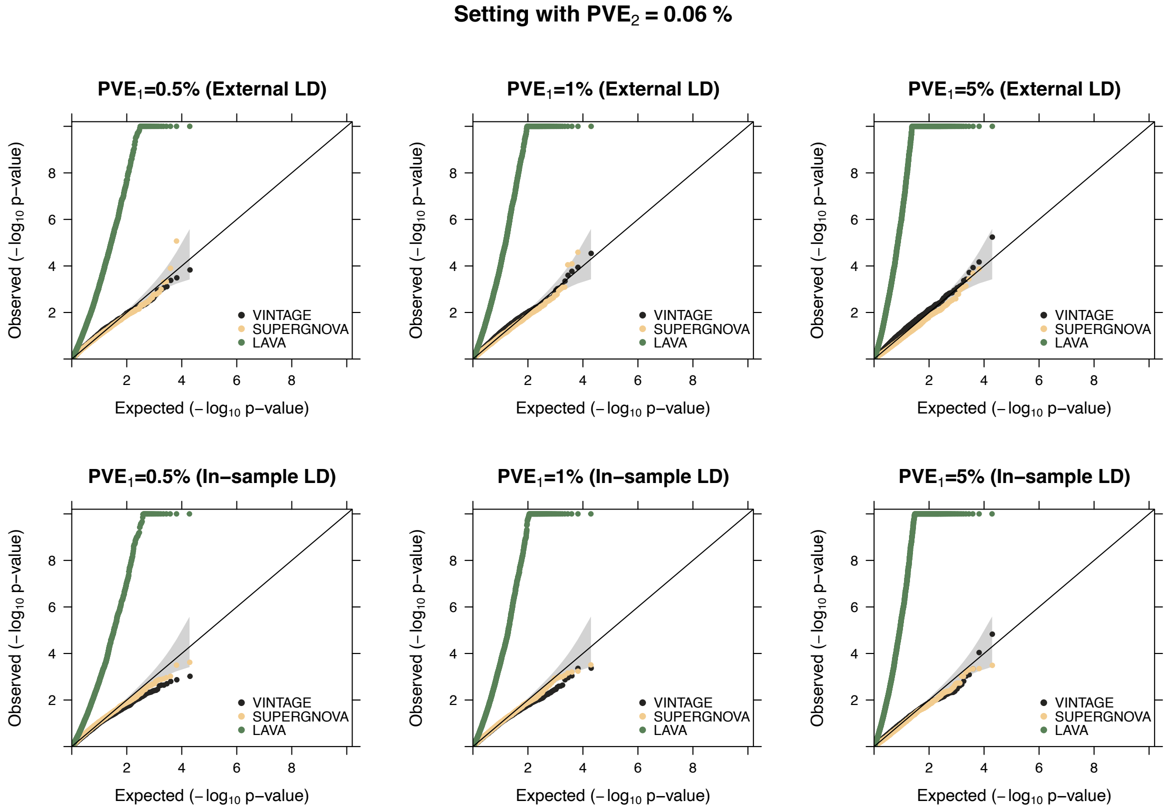


**Figure S17. Quantile-quantile plots of -log$_{10}$ p-values from different methods for gene-wise genetic correlation test in the null simulations.** The null simulations were conducted with (**Top**) external LD matrices or (**Bottom**) in-sample LD matrices. The in-sample LD matrices were computed using the same set of 10,000 UKBB WB individuals for simulating the gene expression data while the external LD matrices were computed using the 503 European individuals from the 1000 Genomes Project. Compared methods include VINTAGE, SUPERGNOVA, and LAVA. The null simulations were also conducted under different $PVE_1$ values (Left: 0.5%, Middle: 1%, and Right: 5%), a polygenic genetic architecture ($q = p$), and a **relatively high** $PVE_2$ **(0.06%)**.

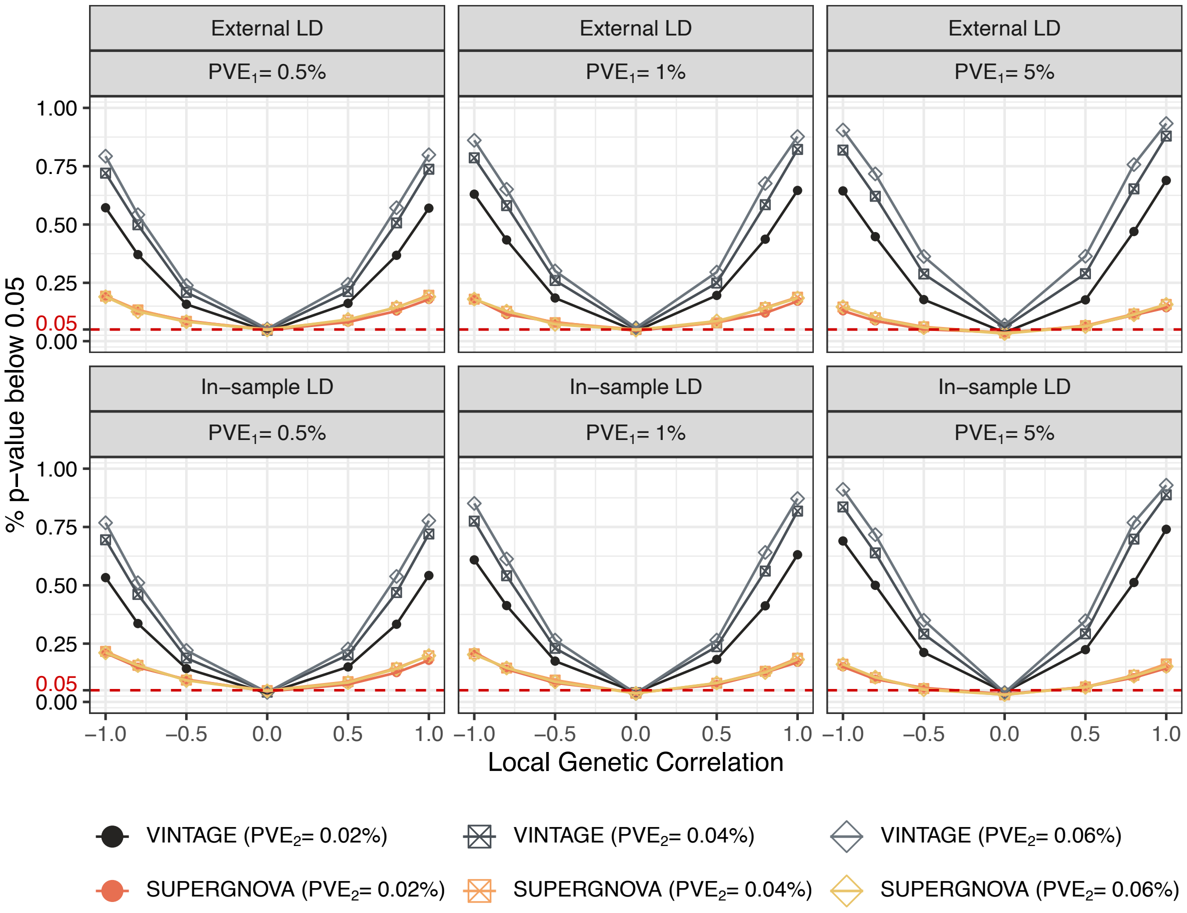


**Figure S18. Power of different methods for gene-wise genetic correlation test in the simulations.** The simulations were conducted with (**Top**) external LD matrices or (**Bottom**) in-sample LD matrices. The in-sample LD matrices were computed using the same set of 10,000 UKBB WB individuals for simulating the gene expression data while the external LD matrices were computed using the 503 European individuals from the 1000 Genomes Project. Compared methods include VINTAGE and SUPERGNOVA. The simulations were also conducted under different $PVE_1$ values (0.5%, 1%, and 5%), different $PVE_2$ values (0.02%, 0.04%, and 0.06%), and different local genetic correlations (-1.0, -0.8, -0.5, 0.0, 0.5, 0.8, 1.0) under a polygenic genetic architecture ($q = p$).

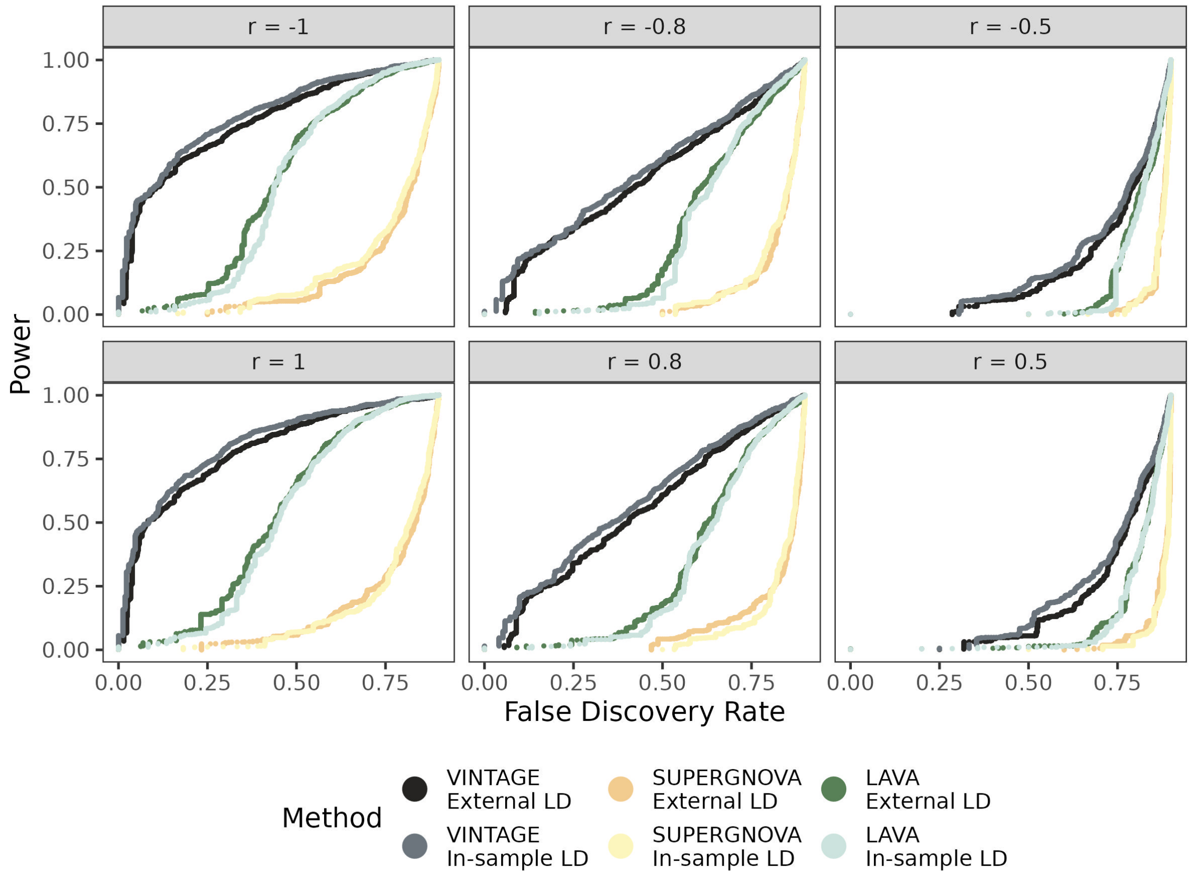


**Figure S19. Power as function of false discovery rate for gene-wise genetic correlation test in the simulations.** Power and false discovery rates were calculated by pairing simulation replicates under the alternative (1,000 replicates) with those under the null (9,000 replicates). The simulations were conducted with (**Top**) external LD matrices or (**Bottom**) in-sample LD matrices. The in-sample LD matrices were computed using the same set of 10,000 UKBB WB individuals for simulating the gene expression data while the external LD matrices were computed using the 503 European individuals from the 1000 Genomes Project. Evaluated methods include VINTAGE, SUPERGNOVA, and LAVA. The simulations were conducted under moderate $PVE$ parameters ($PVE_1 = 1\%$ and $PVE_2 = 0.04\%$), different local genetic correlations, and a polygenic genetic architecture ($q = p$). Simulation replicates with negative heritability estimates or negative variance estimates from SUPERGNOVA or LAVA were excluded from the evaluation.

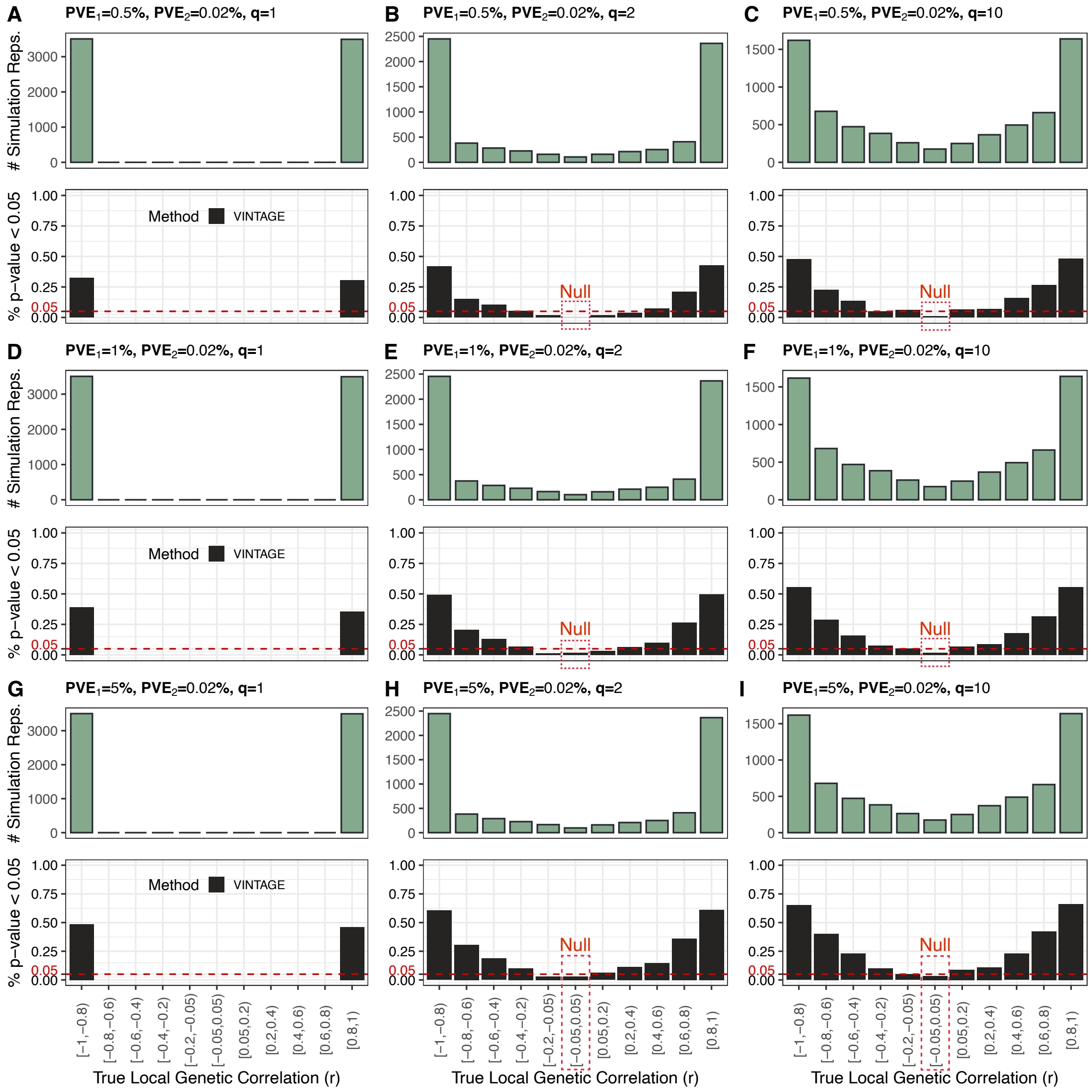


**Figure S20. Power of VINTAGE for gene-wise genetic correlation test under sparse simulation settings and a relatively low $PVE_2$ (0.02%).** In each setting indicated in the title, 7,000 simulation replicates – 1,000 for each value of $\tilde{r}$ (= -1.0, -0.8, -0.5, 0.0, 0.5, 0.8, and 1.0) for simulating the SNP effect sizes – were grouped into 11 categories based on the range of their true underlying local genetic correlations, calculated as the Pearson correlation coefficient between the simulated genetic effect sizes on gene expression and on trait. Top figures show the number of simulation replicates in each category and the bottom figures show the proportion of simulation replicates that passed the p-value threshold of 0.05 in each category. The simulations were conducted under sparse genetic architecture with $q$ = 1, 2, or 10 SNPs that have non-zero effects on both gene expression and the trait, different $PVE_1$ values (0.5%, 1%, and 5%), and in-sample LD matrices.

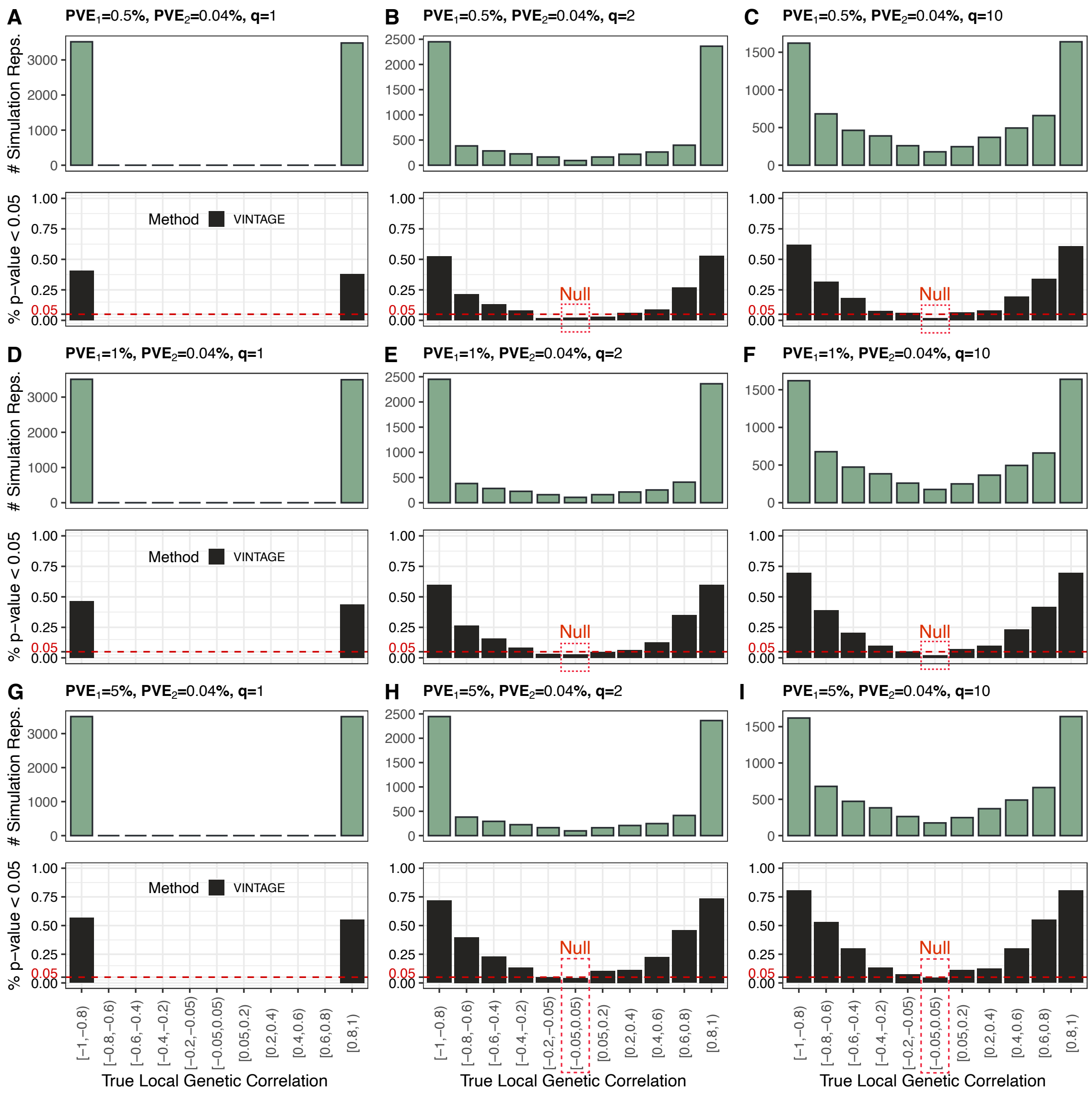


**Figure S21. Power of VINTAGE for gene-wise genetic correlation test under sparse simulation settings and a relatively moderate $PVE_2$ (0.04%).** In each setting indicated in the title, 7,000 simulation replicates – 1,000 for each value of $\tilde{r}$ (= -1.0, -0.8, -0.5, 0.0, 0.5, 0.8, and 1.0) for simulating the SNP effect sizes – were grouped into 11 categories based on the range of their true underlying local genetic correlations, calculated as the Pearson correlation coefficient between the simulated genetic effect sizes on gene expression and on trait. Top figures show the number of simulation replicates in each category and the bottom figures show the proportion of simulation replicates that passed the p-value threshold of 0.05 in each category. The simulations were conducted under sparse genetic architecture with $q$ = 1, 2, or 10 SNPs that have non-zero effects on both gene expression and the trait, different $PVE_1$ values (0.5%, 1%, and 5%), and in-sample LD matrices.

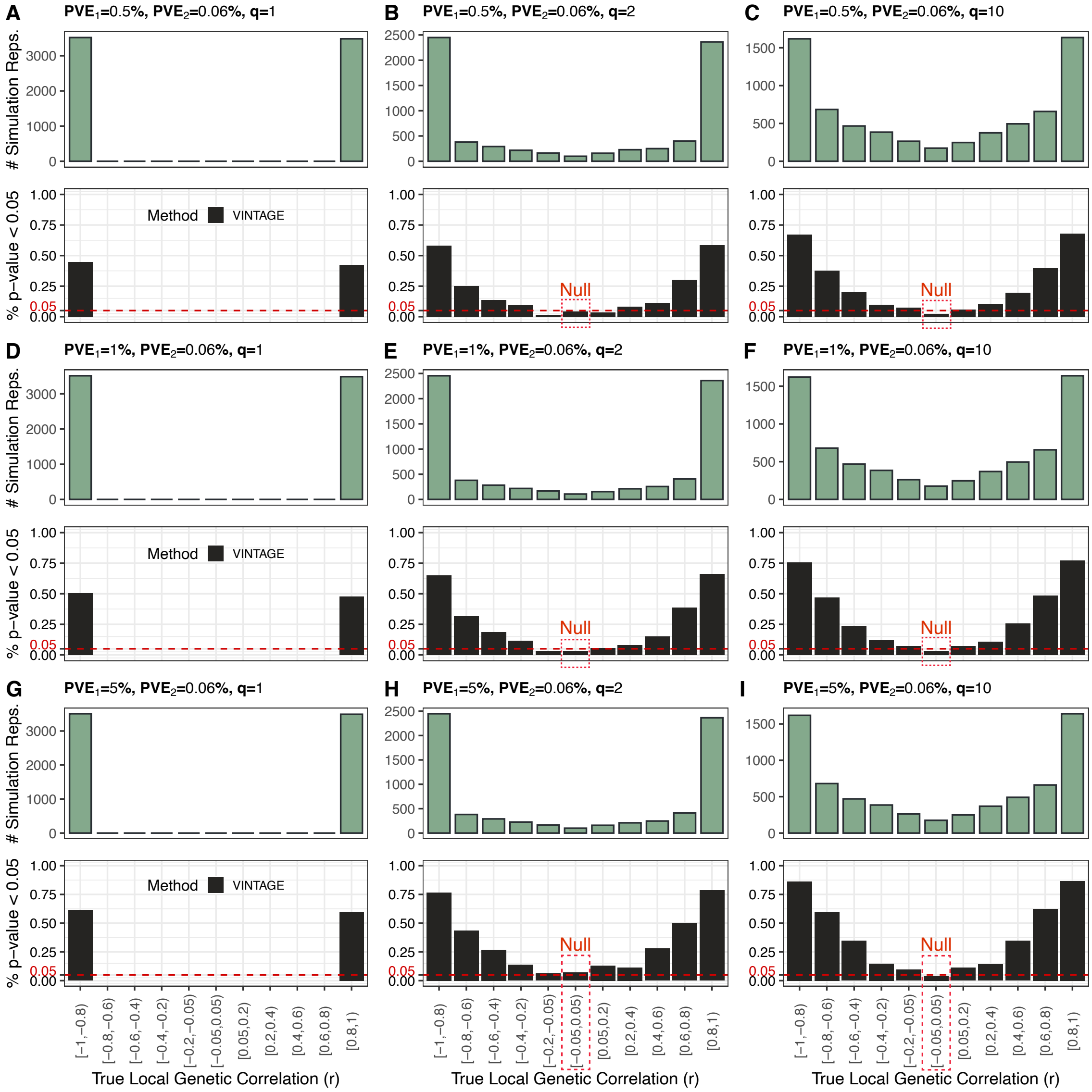


**Figure S22. Power of VINTAGE for gene-wise genetic correlation test under sparse simulation settings and a relatively high $PVE_2$ (0.06%).** In each setting indicated in the title, 7,000 simulation replicates – 1,000 for each value of $\tilde{r}$ (= -1.0, -0.8, -0.5, 0.0, 0.5, 0.8, and 1.0) for simulating the SNP effect sizes – were grouped into 11 categories based on the range of their true underlying local genetic correlations, calculated as the Pearson correlation coefficient between the simulated genetic effect sizes on gene expression and on trait. Top figures show the number of simulation replicates in each category and the bottom figures show the proportion of simulation replicates that passed the p-value threshold of 0.05 in each category. The simulations were conducted under sparse genetic architecture with $q$ = 1, 2, or 10 SNPs that have non-zero effects on both gene expression and the trait, different $PVE_1$ values (0.5%, 1%, and 5%), and in-sample LD matrices.

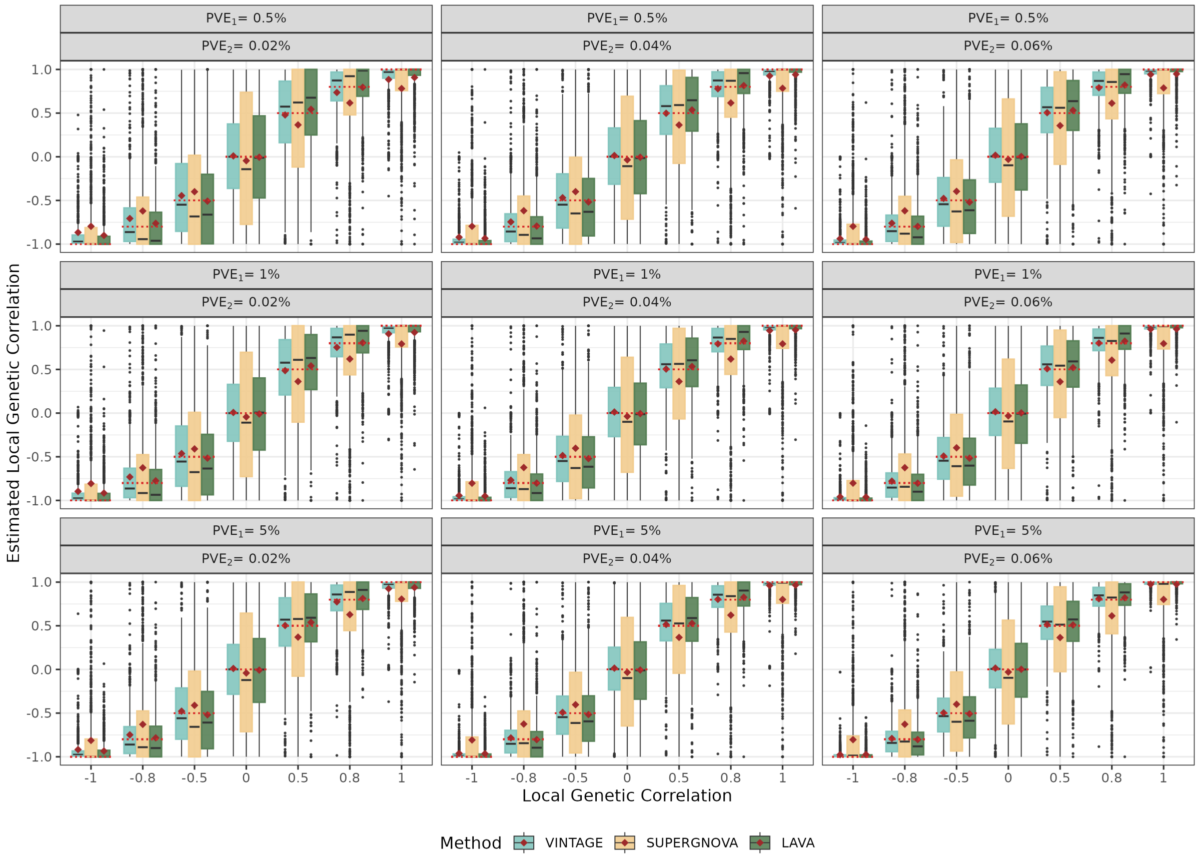


**Figure S23. Estimation accuracy of local genetic correlations in the simulations.** Boxplots indicate the estimated local genetic correlations from different methods. Compared methods include VINTAGE, SUPERGNOVA, and LAVA. The simulations were also conducted under different $PVE_1$ values (Top: 0.5%, Middle: 1%, and Bottom: 5%), different $PVE_2$ values (Left: 0.02%, Middle: 0.04%, and Right: 0.06%), and different local genetic correlations (-1.0, -0.8, -0.5, 0.0, 0.5, 0.8, 1.0) with in-sample LD matrices and a polygenic genetic architecture ($q = p$). Horizontal dotted lines (red) indicate the true local genetic correlations and the diamonds (red) represent the average local genetic correlation estimates across simulation replicates. For a better visualization, local genetic correlation estimates from SUPERGNOVA or LAVA were set to be -1 or 1 if they fell below -1 or above 1, respectively.

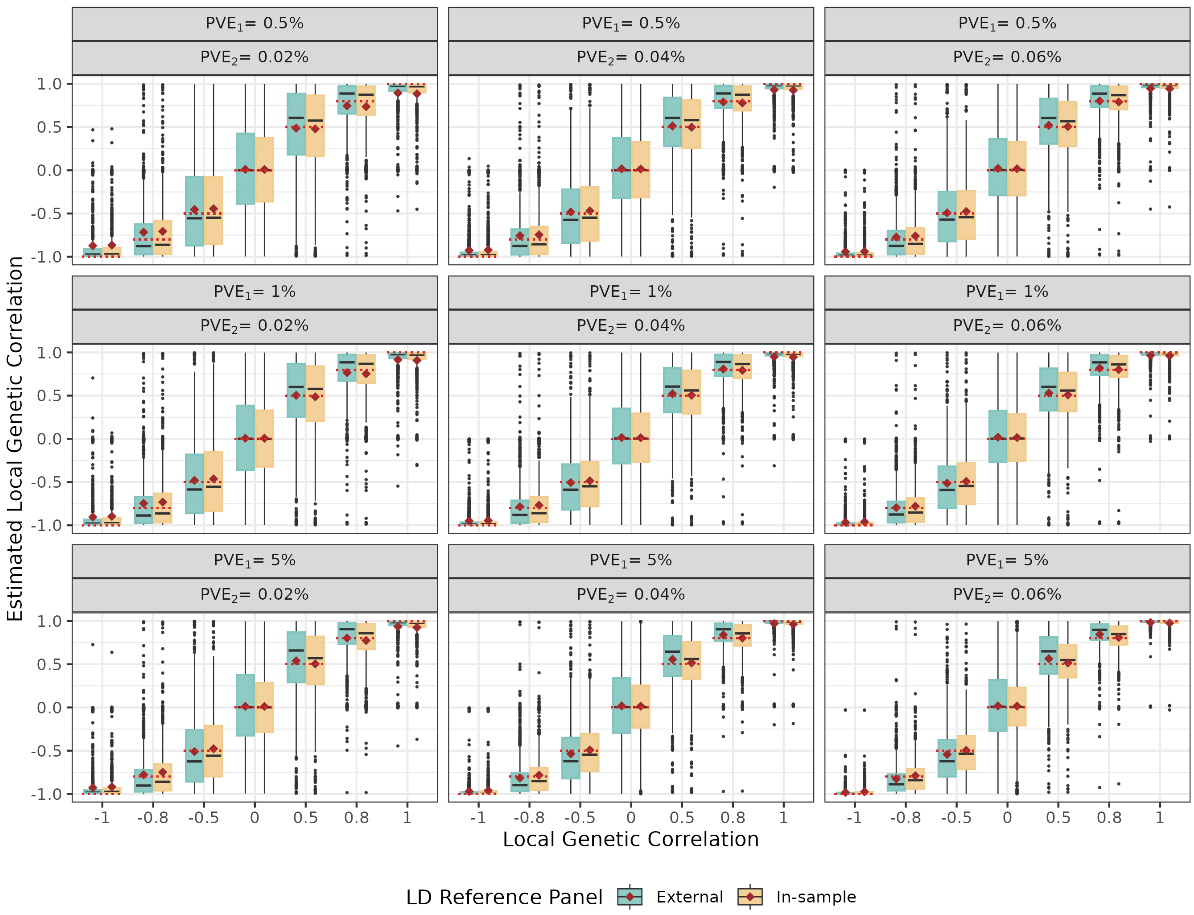


**Figure S24. Estimation accuracy of local genetic correlations in the simulation.** Boxplots indicate the estimated local genetic correlations from VINTAGE using in-sample or external LD matrices. The in-sample LD matrices were computed using the same set of 10,000 UKBB WB individuals for simulating the gene expression data and the external LD matrices were computed using the 503 European individuals from the 1000 Genomes Project. The simulations were also conducted under different $PVE_1$ values (Top: 0.5%, Middle: 1%, and Bottom: 5%), different $PVE_2$ values (Left: 0.02%, Middle: 0.04%, and Right: 0.06%), and different local genetic correlations (-1.0, -0.8, -0.5, 0.0, 0.5, 0.8, 1.0) under a polygenic genetic architecture ($q = p$). Horizontal dotted lines (red) indicate the true local genetic correlations and the diamonds (red) represent the average local genetic correlation estimates across simulation replicates.

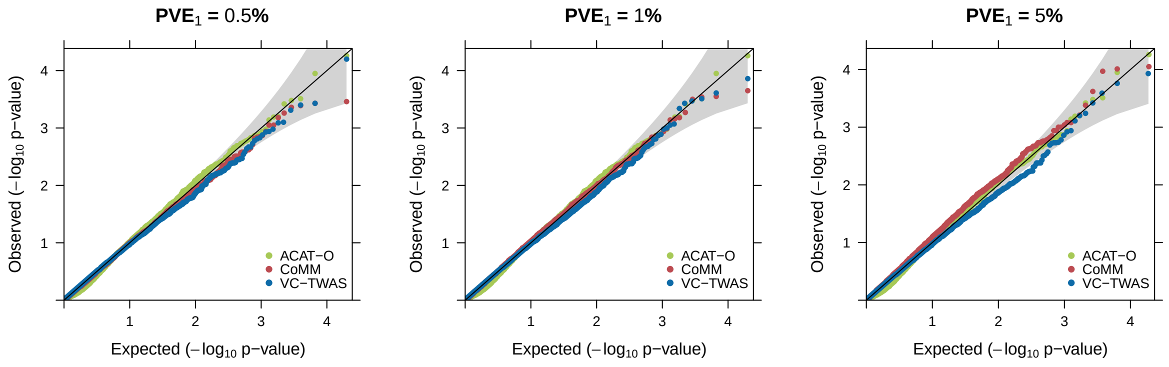


**Figure S25. Quantile-quantile plots of -log$_{10}$ p-values from three additional methods for gene-wise genetic variance test under the null simulations.** Evaluated methods include ACAT-O, which is a SKAT extension, CoMM, and VC-TWAS, which are TWAS extensions. Null simulations were conducted under different $PVE_1$ values (0.5%, 1%, and 5%) with in-sample LD matrices and a polygenic genetic architecture ($q = p$).

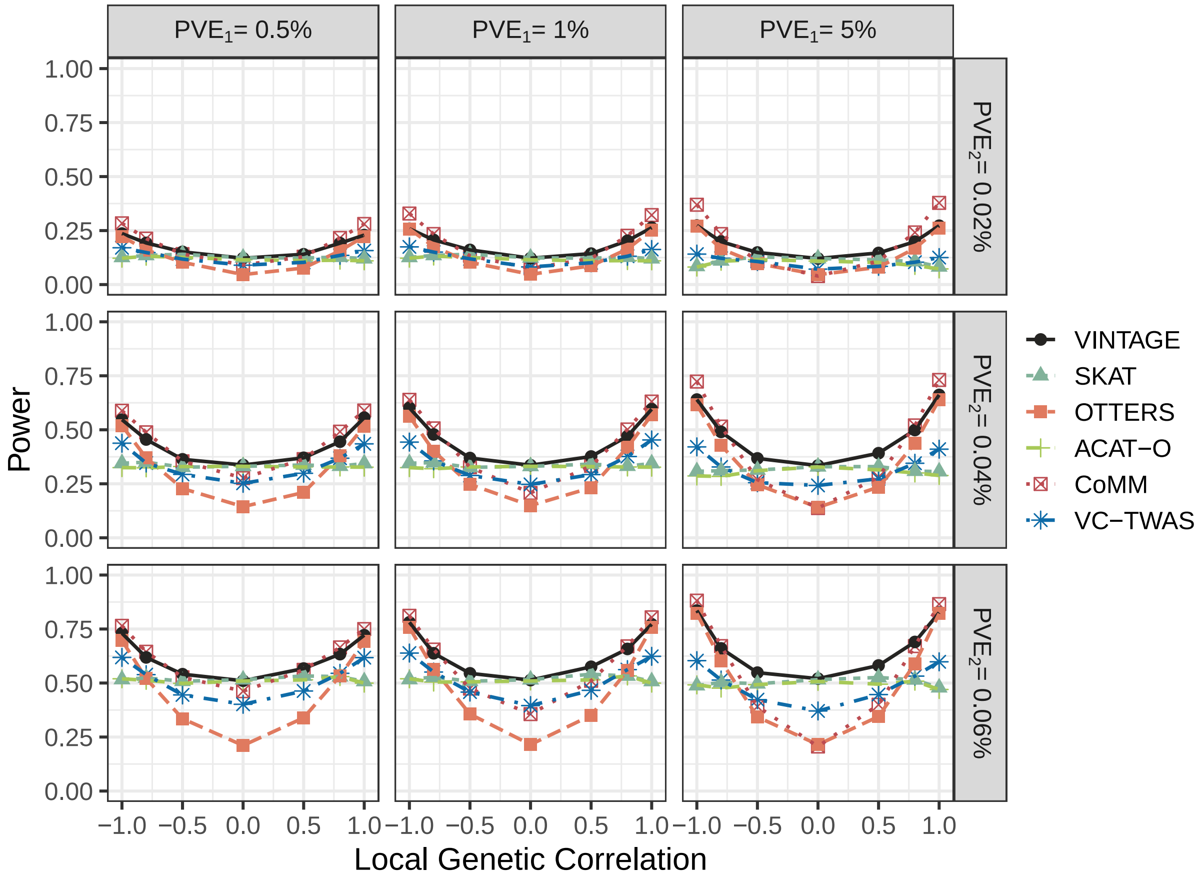


**Figure S26. Power of additional methods for gene-wise genetic variance test in the simulations.** Compared methods include VINTAGE, SKAT with an unweighted linear kernel, TWAS that adapts multiple PRS methods to estimate gene expression prediction weights and performs an omnibus test, ACAT-O, which is a SKAT extension, CoMM, and VC-TWAS, which are TWAS extensions. The simulations were also conducted under different $PVE_1$ values (0.5%, 1%, and 5%), different $PVE_2$ values (0.02%, 0.04%, and 0.06%), and different local genetic correlations (-1.0, -0.8, -0.5, 0.0, 0.5, 0.8, 1.0) with in-sample LD matrices and a polygenic genetic architecture ($q = p$).

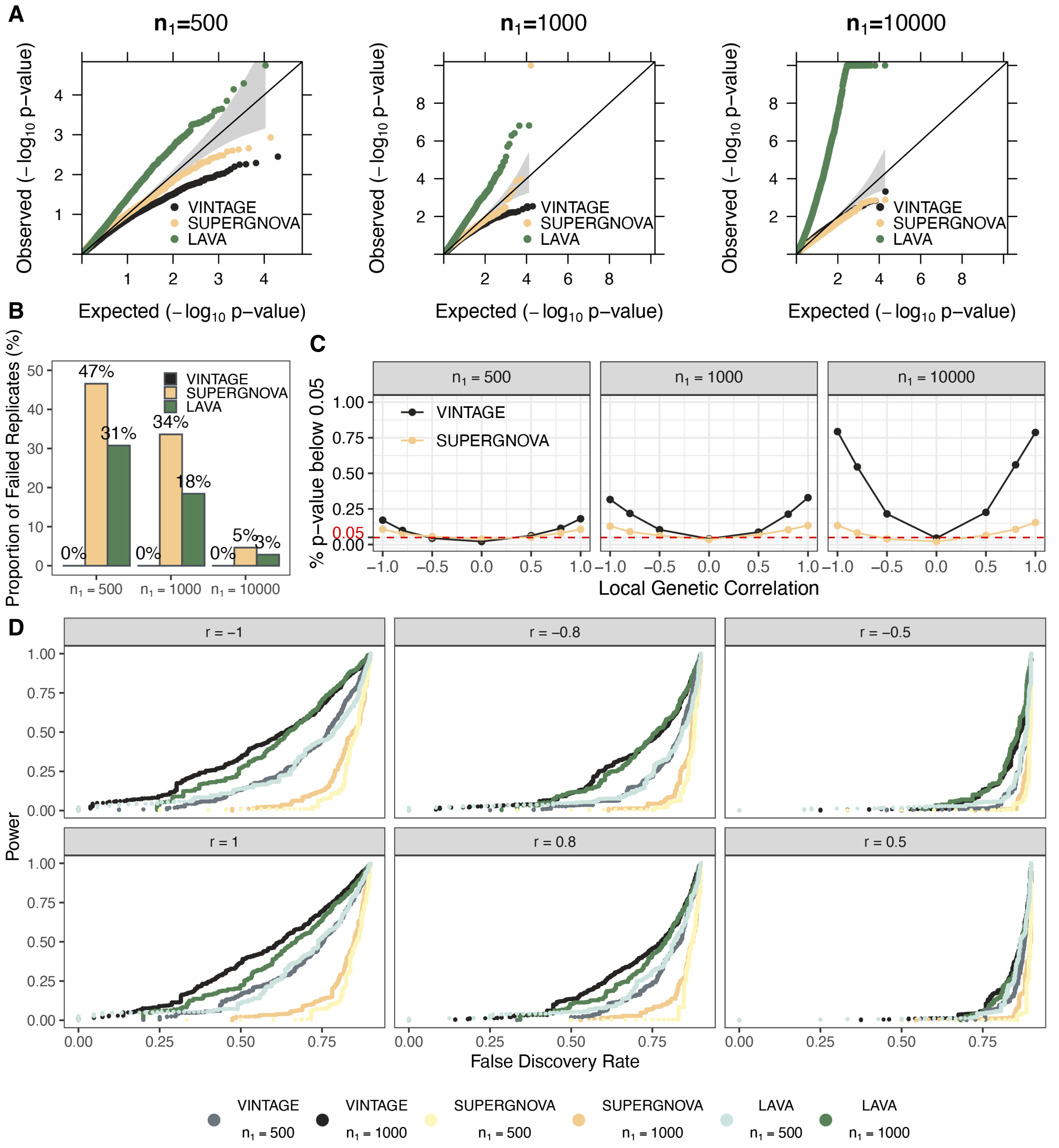


**Figure S27. Evaluating the influence of the sample size of the gene expression study in the simulations.** The performance of VINTAGE, SUPERGNOVA, and LAVA was evaluated for the gene-wise genetic correlation test across different sample sizes of the gene expression study. Evaluated sample sizes include $n_1 = 500$, 1,000, and 10,000. (**A**) Quantile-quantile plots of -$\log_{10}$ p-values in the null simulations. (**B**) Proportion of failed simulation replicates in the null simulations. (**C**) Type I error rate ($r = 0$) or power ($r \neq 0$) was evaluated based on the proportion of simulation replicates that passed the p-value threshold of 0.05. (**D**) Power was evaluated across different false discovery rates with power and false discovery rates calculated by pairing simulation replicates under the alternative (1,000 replicates) with those under the null (9,000 replicates). The simulations were conducted under settings with moderate $PVE$ parameters (i.e., $PVE_1 = 1\%$ and $PVE_2 = 0.04\%$) and different local genetic correlations (-1.0, -0.8, -0.5, 0.0, 0.5, 0.8, 1.0) with in-sample LD matrices and a polygenic genetic architecture ($q = p$).

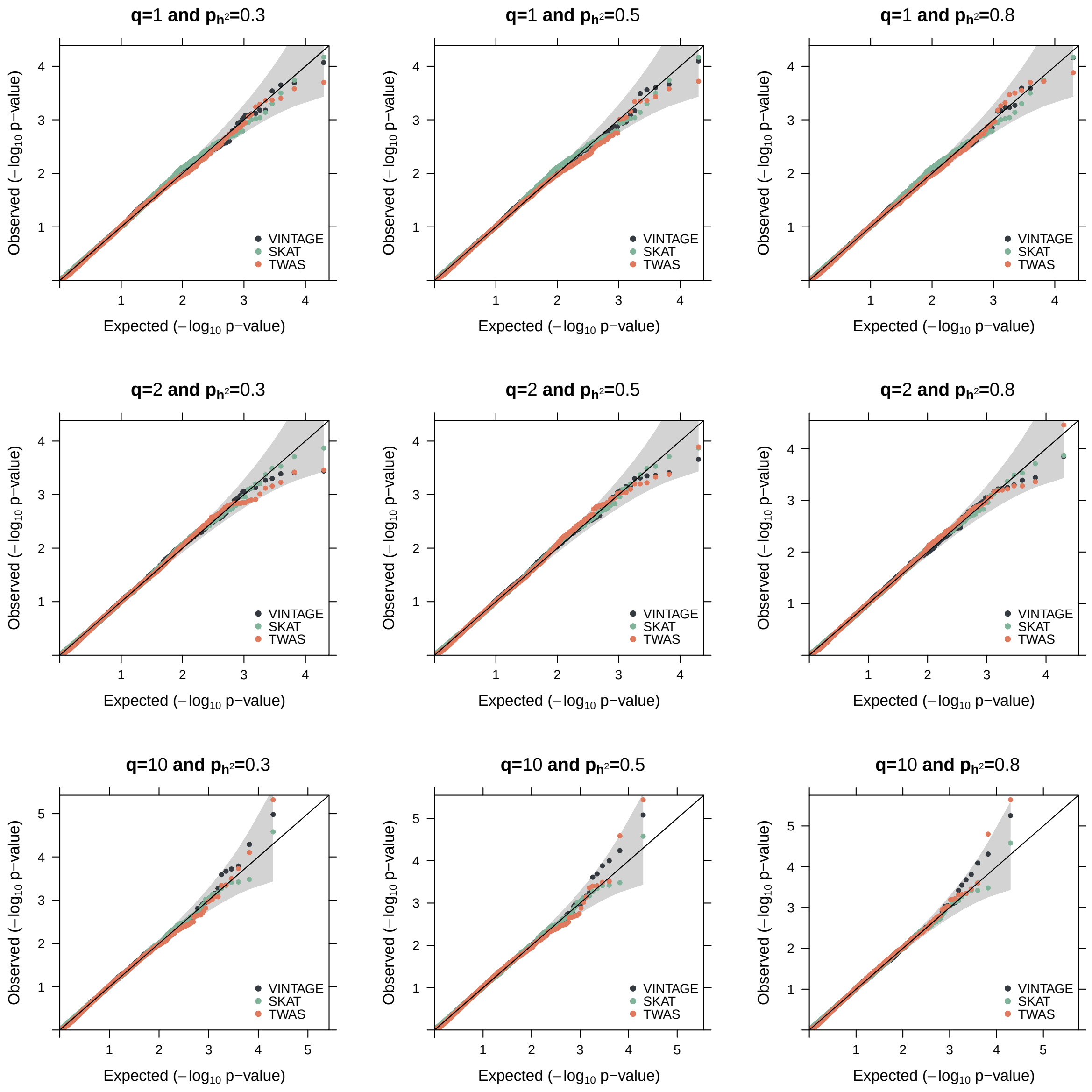


**Figure S28. Quantile-quantile plots of -$\log_{10}$ p-values from different methods for gene-wise genetic variance test in the null simulations.** The null simulations were conducted under settings where the genetic architecture underlying gene expression was polygenic and a sparse set of $q$ cis-SNPs explained a large proportion of variance in gene expression as quantified by $p_{h^2}$, among all cis-SNPs in the gene region. Evaluated methods include VINTAGE, SKAT with an unweighted linear kernel, and TWAS that adapts multiple PRS methods to estimate gene expression prediction weights and performs an omnibus test. In the simulations, $q$ was set to be 1 (**Top**), 2 (**Middle**), or 10 (**Bottom**) and $p_{h^2}$ was set to be 0.3 (**Left**), 0.5 (**Middle**), or 0.8 (**Right**) with moderate $PVE$ parameters (i.e., $PVE_1 = 1\%$ and $PVE_2 = 0.04\%$) and in-sample LD matrices.

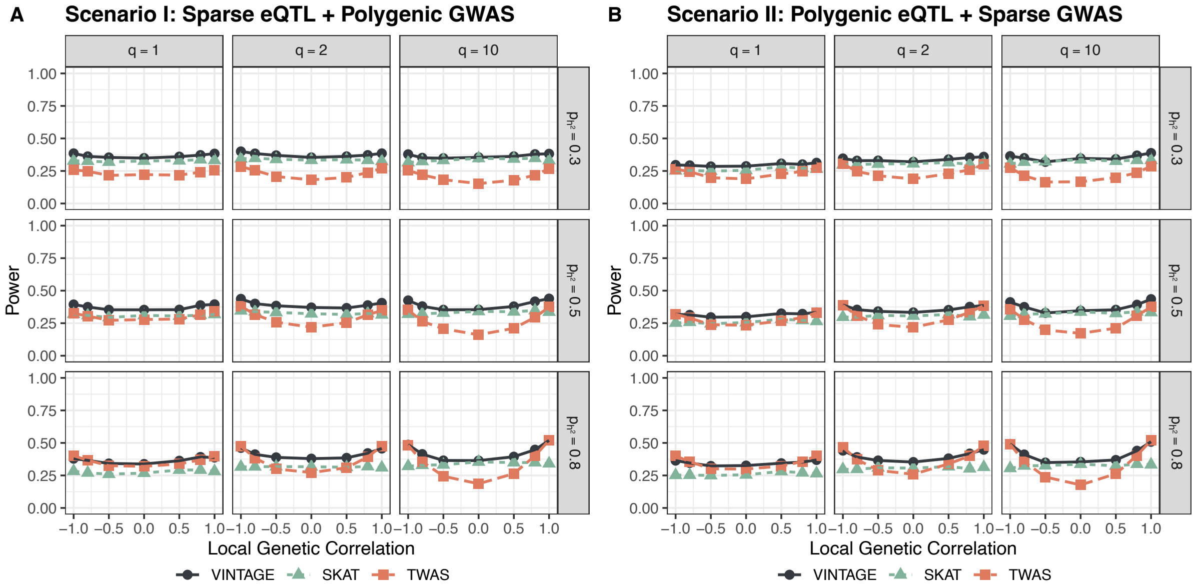


**Figure S29. Power of different methods for gene-wise genetic variance test in the simulations.** Evaluated methods include VINTAGE, SKAT with an unweighted linear kernel, and TWAS that adapts multiple PRS methods to estimate gene expression prediction weights and performs an omnibus test. The simulations were conducted under settings where the genetic architecture is sparse in one study, while it is polygenic in the other. Two scenarios were examined. (**A**) In scenario I, the genetic architecture for gene expression in the expression study was set to be sparse and the genetic architecture for the trait in GWAS was set to be polygenic. (**B**) In scenario II, conditions in scenario I were reversed. For sparse architecture, the proportion of variance was fully explained by a sparse set of $q$ cis-SNPs. For polygenic architecture, the sparse set of $q$ cis-SNPs explained a large proportion of variance as quantified by $p_{h^2}$, among all cis-SNPs in the gene region. In the simulations, $q$ was set to be 1, 2, or 10 and $p_{h^2}$ was set to be 0.3, 0.5, or 0.8 with moderate $PVE$ parameters (i.e., $PVE_1 = 1\%$ and $PVE_2 = 0.04\%$) and in-sample LD matrices.

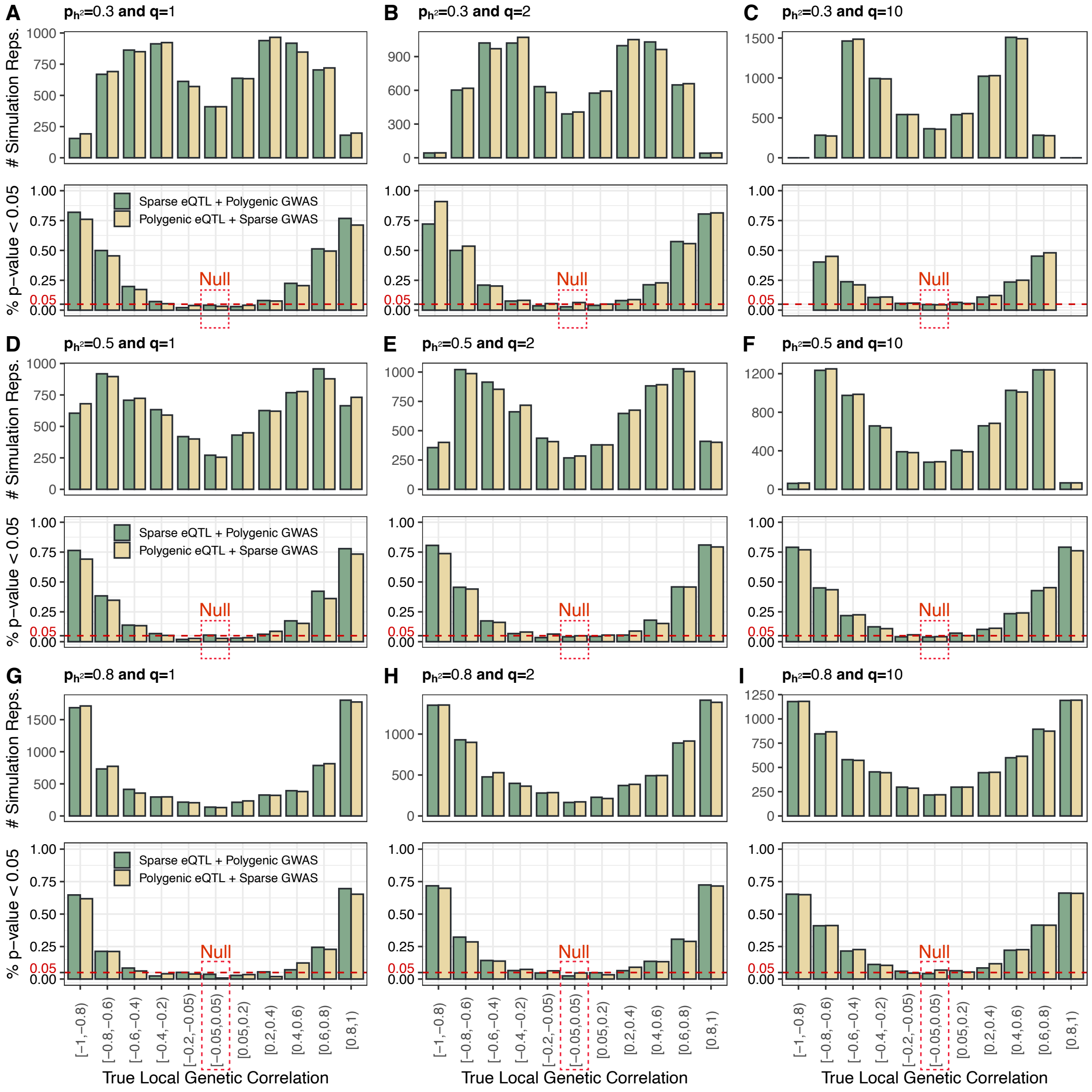


**Figure S30. Power of VINTAGE for gene-wise genetic correlation test under simulation settings where the genetic architecture is sparse in one study, while it is polygenic in the other.** In each setting indicated in the title, 7,000 simulation replicates – 1,000 for each value of $\tilde{r}$ (= -1.0, -0.8, -0.5, 0.0, 0.5, 0.8, and 1.0) for simulating the SNP effect sizes – were grouped into 11 categories based on the range of their true underlying local genetic correlations, calculated as the Pearson correlation coefficient between the simulated genetic effect sizes on gene expression and on trait. Top figures show the number of simulation replicates in each category and the bottom figures show the proportion of simulation replicates that passed the p-value threshold of 0.05 in each category. Two scenarios were examined: In scenario I, the genetic architecture for gene expression in the expression study was set to be sparse and the genetic architecture for the trait in the GWAS was set to be polygenic; In scenario II, conditions in scenario I were reversed. For sparse genetic architecture, the proportion of variance was fully explained by a sparse set of $q$ cis-SNPs. For polygenic genetic architecture, the sparse set of $q$ cis-SNPs explained a large proportion of variance as quantified by $p_{h^2}$, among all cis-SNPs in the gene region. In the simulations, $q$ was set to be 1, 2, or 10, $p_{h^2}$ was set to be 0.3, 0.5, or 0.8, $PVE_1 = 1\%$, $PVE_2 = 0.04\%$, and in-sample LD matrices were used.

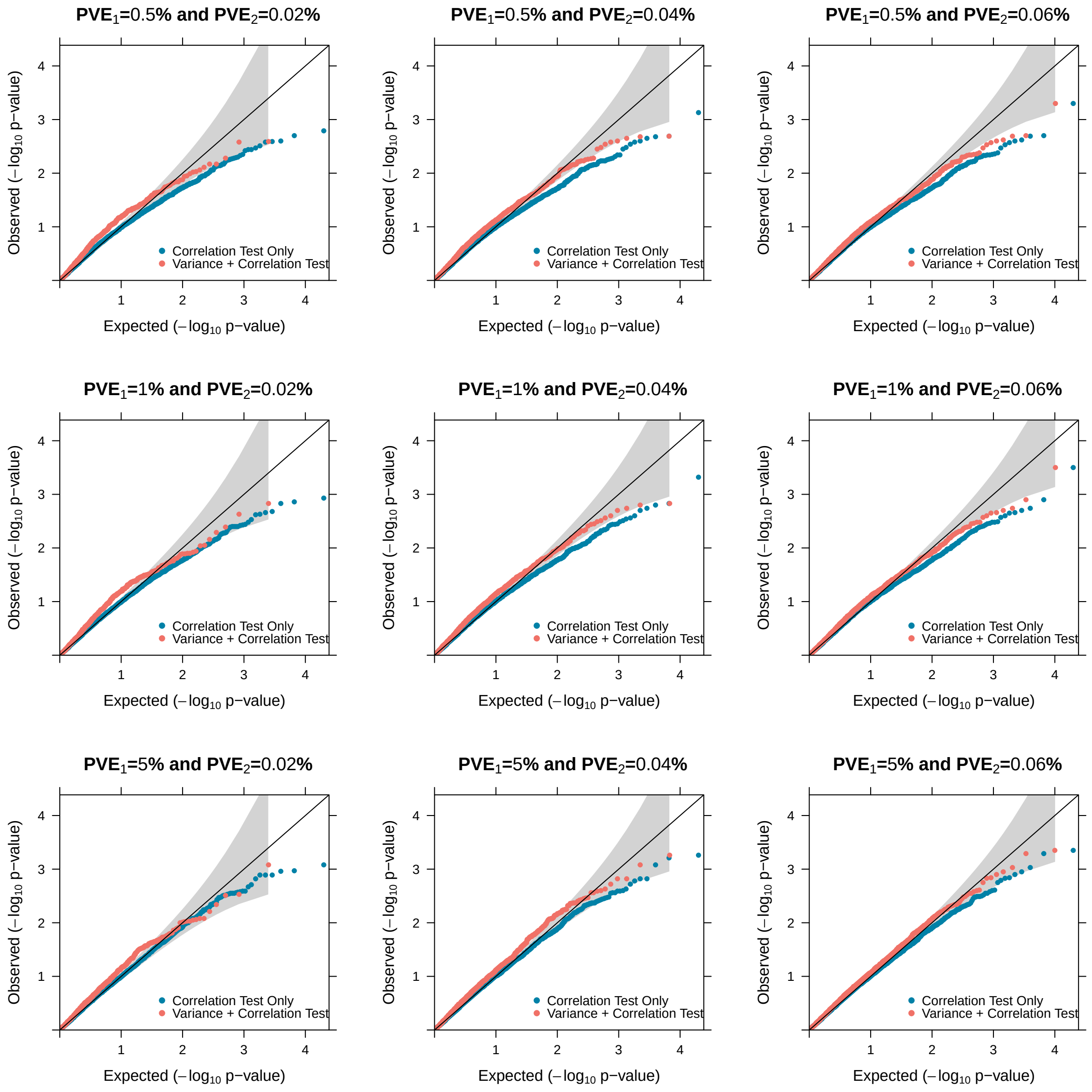


**Figure S31. Quantile-quantile plots of -log$_{10}$ p-values from two procedures for evaluating the gene-wise genetic correlation test in the null simulations.** The two procedures include **"Correlation Test"** that ignores the genetic variance test, as described in the main simulation studies, while **"Variance+Correlation Test"** takes it into account by first conducting the genetic variance test and only performing the genetic correlation test if it passed the significance threshold of $2.5 \times 10^{-6}$ in the genetic variance test. The simulations were also conducted under different $PVE_1$ values (0.5%, 1%, and 5%), different $PVE_2$ values (0.02%, 0.04%, and 0.06%), and different local genetic correlations (-1.0, -0.8, -0.5, 0.0, 0.5, 0.8, 1.0) with in-sample LD matrices and a polygenic genetic architecture ($q = p$).

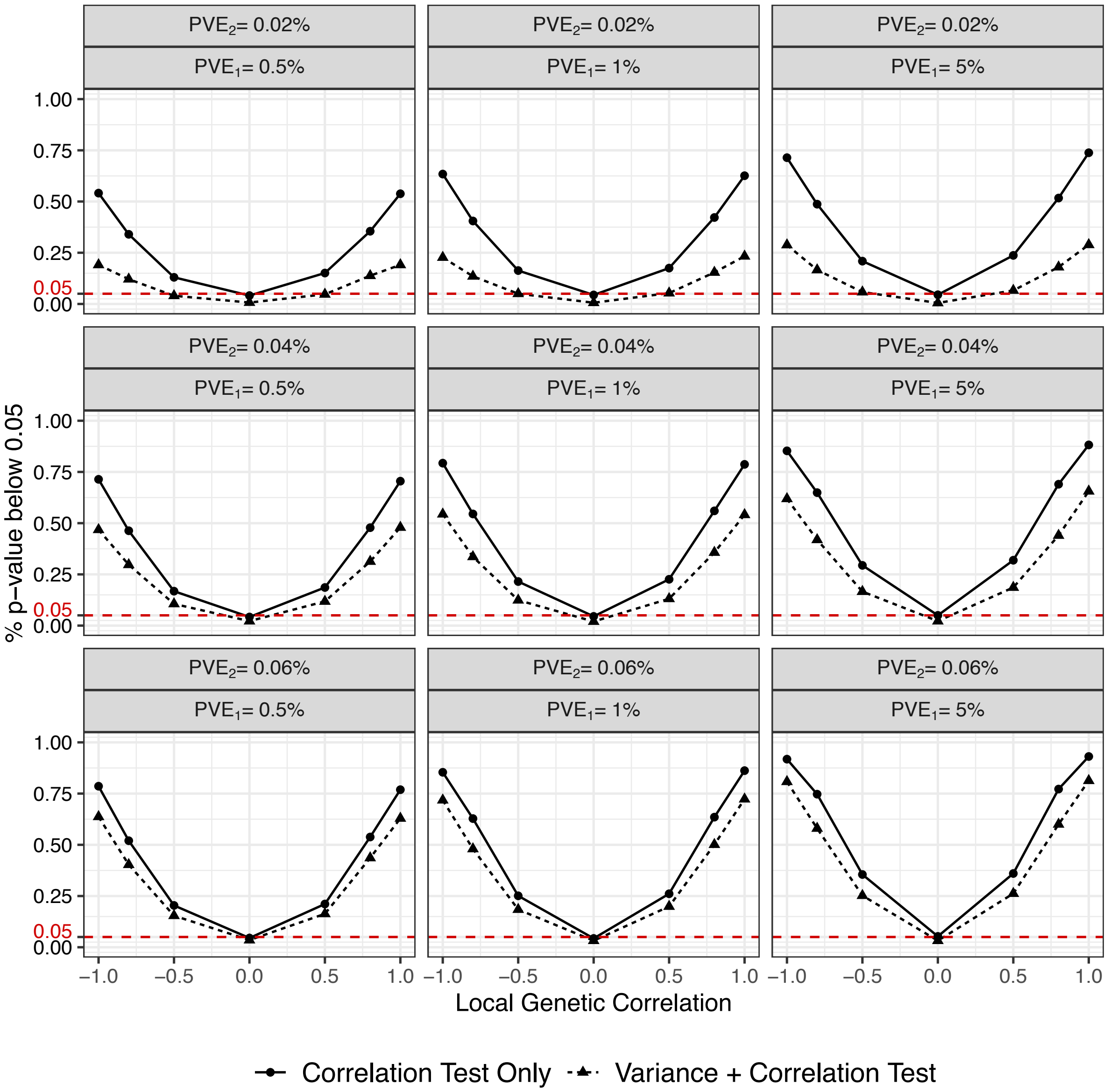


**Figure S32. Power of two procedures for gene-wise genetic correlation test in the simulations.** The two procedures include **"Correlation Test"** that ignores the genetic variance test, as described in the main simulation studies, while **"Variance+Correlation Test"** takes it into account by first conducting the genetic variance test and only performing the genetic correlation test if it passed the significance threshold of $2.5 \times 10^{-6}$ in the genetic variance test. Power of the genetic correlation test under each procedure was evaluated based on a nominal p-value threshold of 0.05. The simulations were also conducted under different $PVE_1$ values (0.5%, 1%, and 5%), different $PVE_2$ values (0.02%, 0.04%, and 0.06%), and different local genetic correlations (-1.0, -0.8, -0.5, 0.0, 0.5, 0.8, 1.0) with in-sample LD matrices and a polygenic genetic architecture ($q = p$).

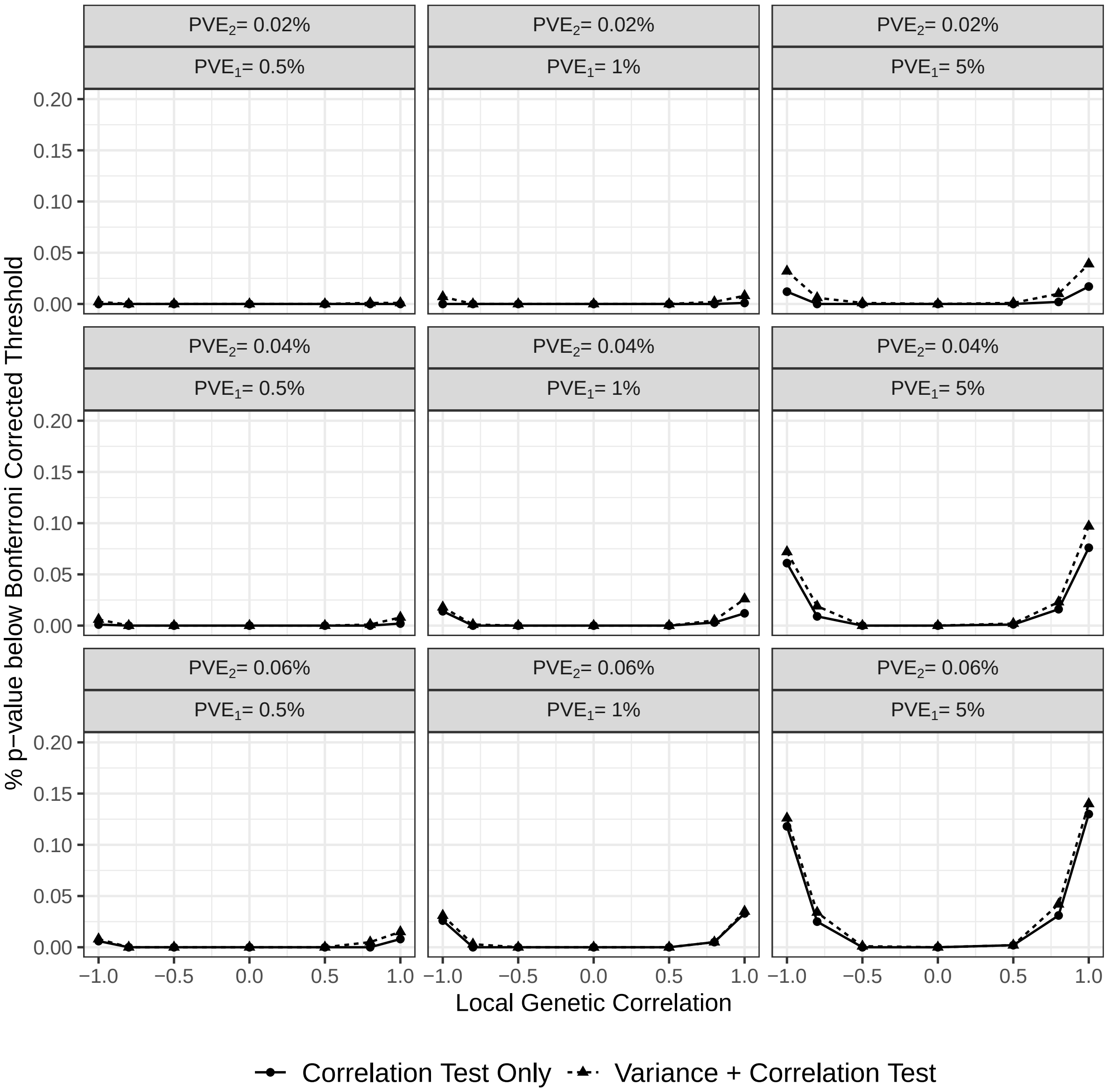


**Figure S33. Power of two procedures for gene-wise genetic correlation test in the simulations.** The two procedures include **"Correlation Test"** that ignores the genetic variance test, as described in the main simulation studies, while **"Variance+Correlation Test"** takes it into account by first conducting the genetic variance test and only performing the genetic correlation test if it passed the significance threshold of $2.5 \times 10^{-6}$ in the genetic variance test. Power of the genetic correlation test under each procedure was evaluated based on a Bonferroni corrected p-value threshold of 0.05, adjusting for the number of genes that passed the genetic variance test. The simulations were also conducted under different $PVE_1$ values (0.5%, 1%, and 5%), different $PVE_2$ values (0.02%, 0.04%, and 0.06%), and different local genetic correlations (-1.0, -0.8, -0.5, 0.0, 0.5, 0.8, 1.0) with in-sample LD matrices and a polygenic genetic architecture ($q = p$).

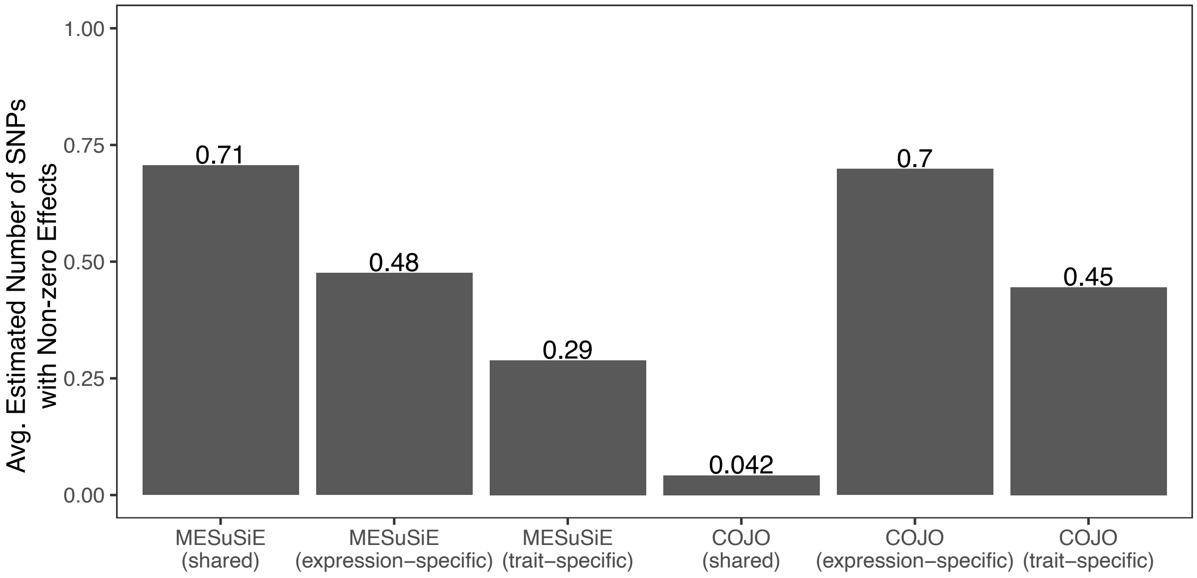


**Figure S34. Evaluating MESuSiE and COJO for estimating the number of non-zero effect SNPs in the simulations.** Barplots showing the average number of SNPs with non-zero effects on both gene expression and the trait (shared), only the gene expression (expression-specific), and only the trait (trait-specific), as estimated by MESuSiE and COJO. Simulations were conducted under settings where one shared cis-SNP was randomly selected to have a non-zero effect on both gene expression and the trait, and two additional cis-SNPs were randomly selected with one having a non-zero effect on gene expression and the other having a non-zero effect on the trait. Simulations were conducted under moderate $PVE$ parameters (i.e., $PVE_1 = 1\%$ and $PVE_2 = 0.04\%$) and different local genetic correlations (-1.0, -0.8, -0.5, 0.0, 0.5, 0.8, 1.0) with in-sample LD matrices and a polygenic genetic architecture ($q = p$).

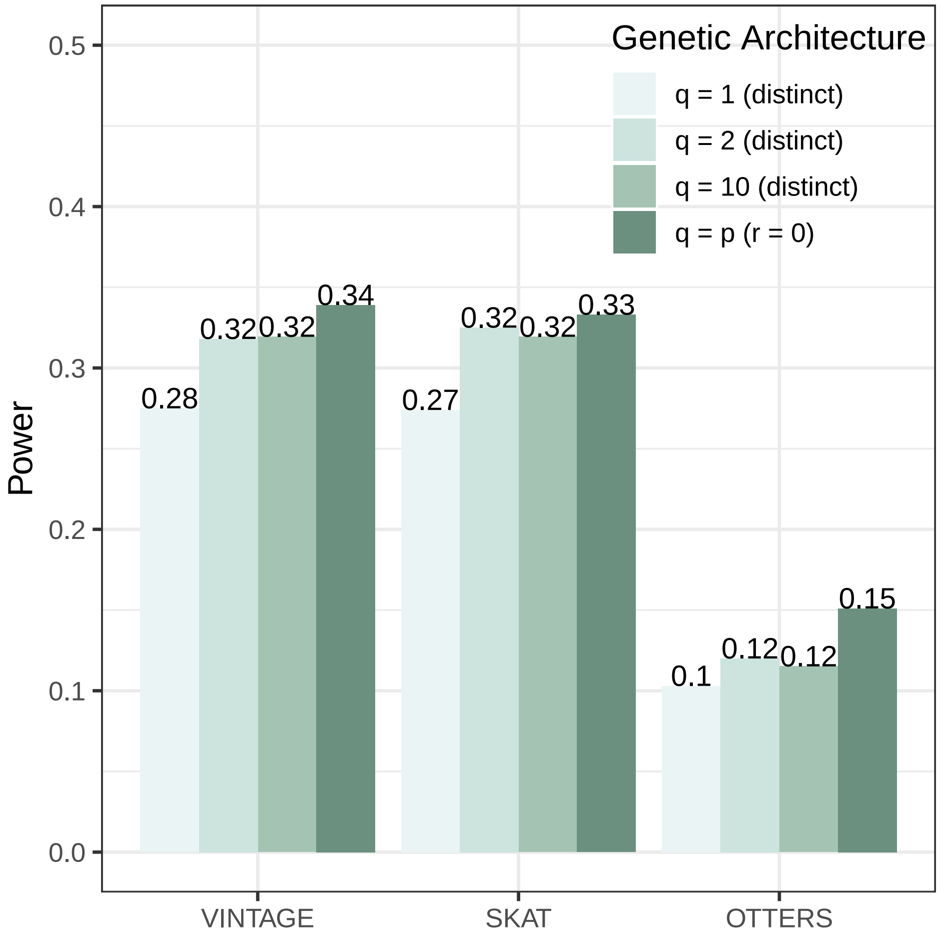


**Figure S35. Power of different methods for gene-wise genetic variance test in simulations where the gene expression study and GWAS have distinct sets of non-zero effect SNPs.** Simulations were conducted under settings where two sets of SNPs were used, with 1, 2, or 10 SNPs in each set: one set with non-zero effects on gene expression and the other set with non-zero effects on the trait, to evaluate the power of the gene-wise genetic variance test. The two sets of SNPs were distinct and did not exhibit extreme high LD ($R^2 < 0.95$) with each other. Power under settings with a polygenic genetic architecture and a local genetic correlation being zero was also included for comparison. Evaluated methods include VINTAGE, SKAT with an unweighted linear kernel, and TWAS that adapts multiple PRS methods to estimate gene expression prediction weights and performs an omnibus test. Simulations were conducted under moderate $PVE$ parameters (i.e., $PVE_1 = 1\%$ and $PVE_2 = 0.04\%$) with in-sample LD matrices.

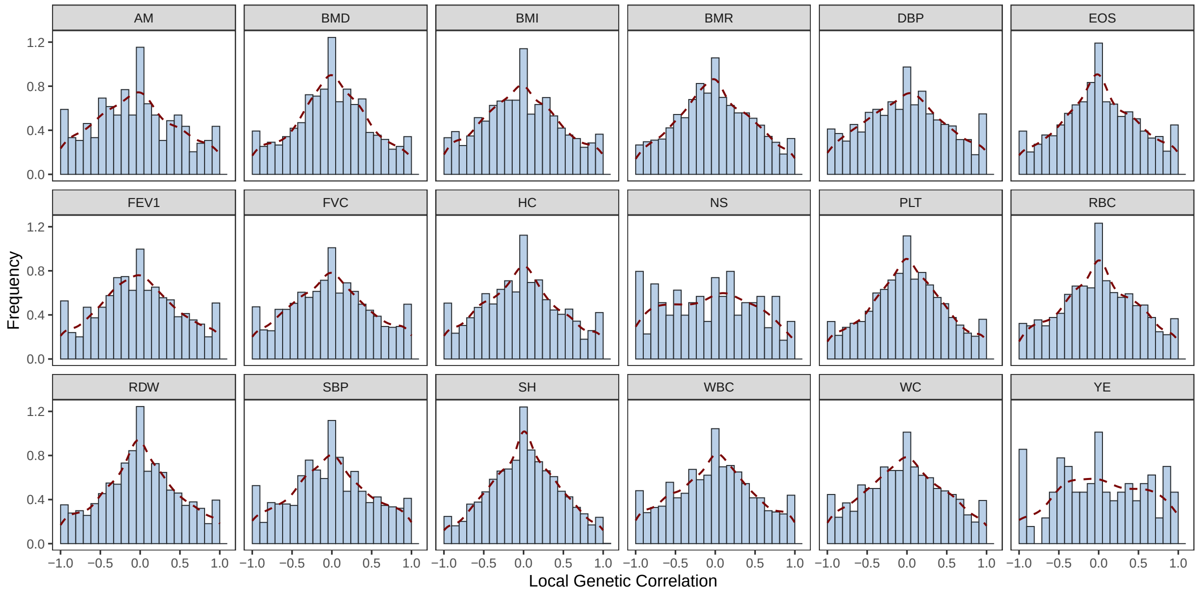


**Figure S36. Histograms of the estimated local genetic correlations by VINTAGE across genes that passed the genetic variance tests for 18 UKBB quantitative traits.**

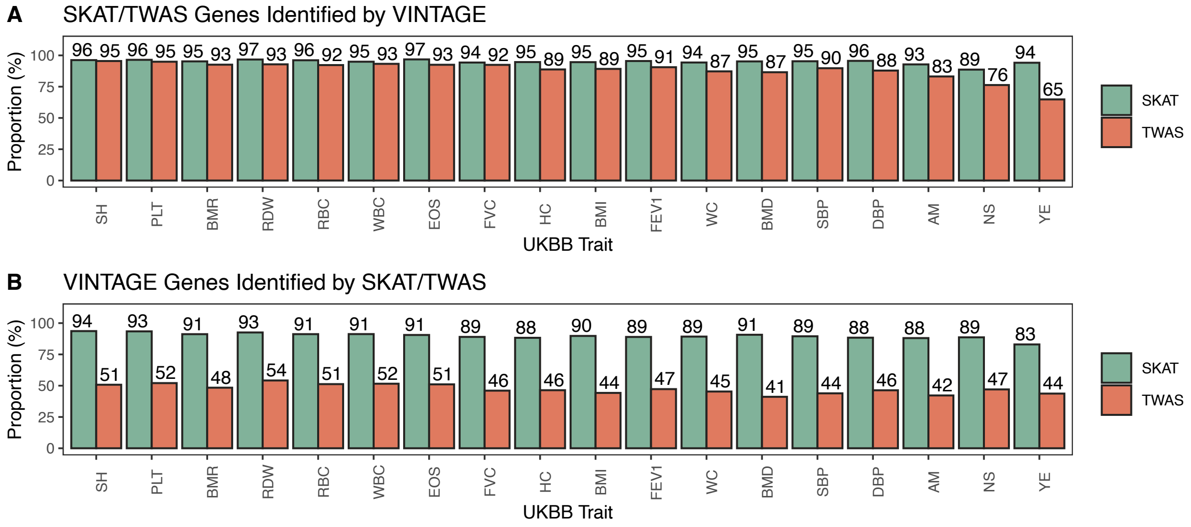


**Figure S37. Barplots comparing the overlapping of the identified genes of different methods in the gene-wise genetic variance analysis.** Compared methods include VINTAGE, SKAT, and TWAS. (**A**) Proportion of genes identified by SKAT or TWAS per trait that were also identified by VINTAGE. (**B**) Proportion of genes identified by VINTAGE per trait that were also identified by SKAT or TWAS.

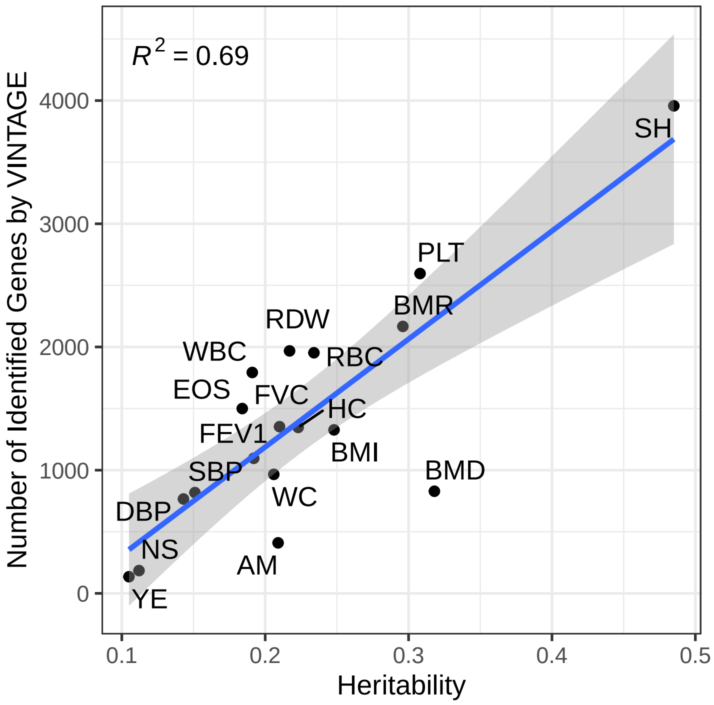


**Figure S38. Scatter plot showing the relationship between trait heritability and the number of identified genes by VINTAGE in the gene-wise genetic variance tests.** Eighteen quantitative traits from UKBB were examined and their heritability was retrieved from the UKB SNP-Heritability Browser.

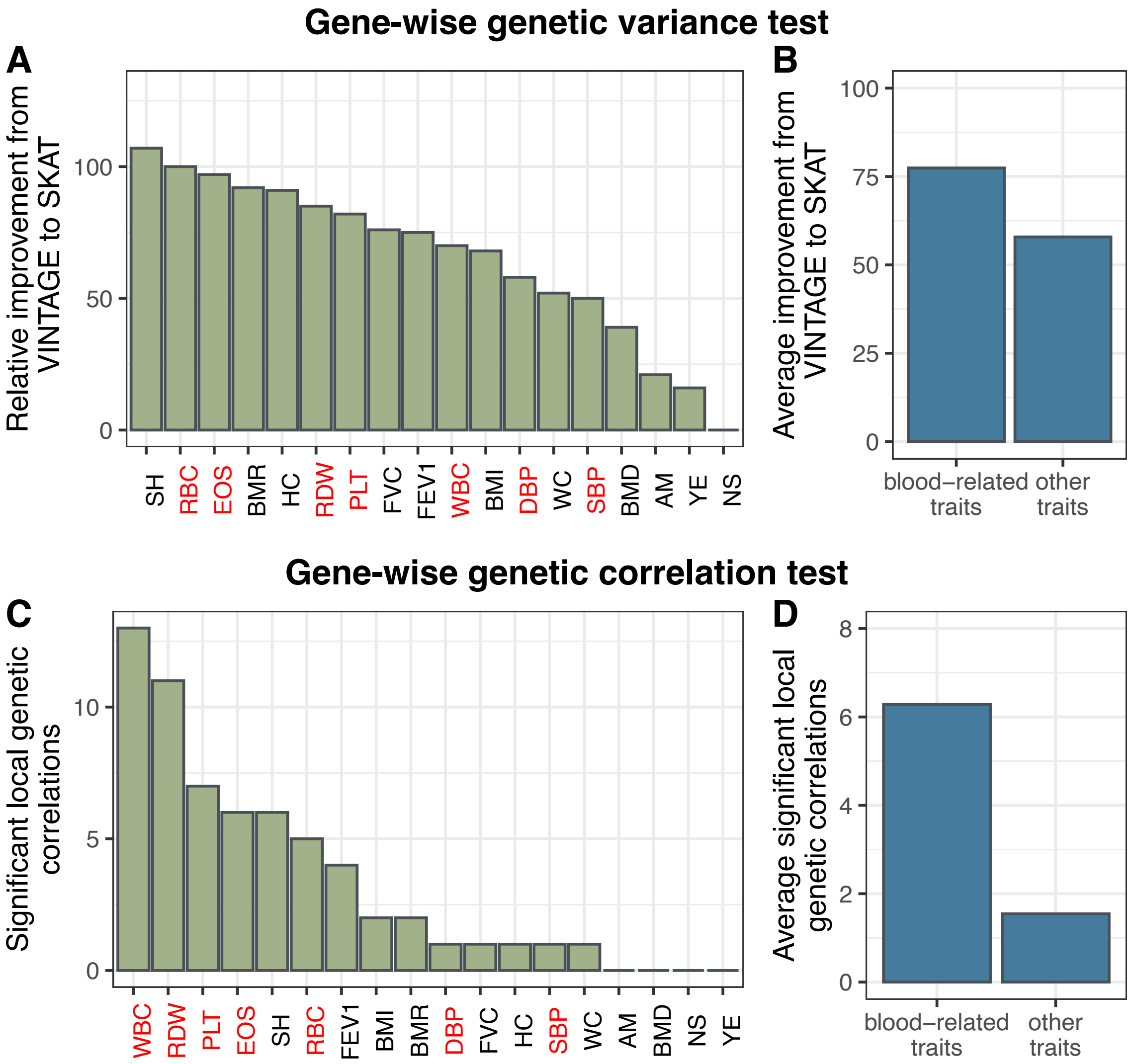


**Figure S39. The performance of VINTAGE is consistent with the use of blood gene expression data.** (**A**) Number of extra identified genes from VINTAGE's genetic variance test relative to SKAT where blood-related traits are colored in red. (**B**) Average number of extra identified genes in blood-related traits and other traits. (**C**) Number of identified genes from VINTAGE's genetic correlation test where blood-related traits are colored in red. (**D**) Average number of identified genes with significant local genetic correlation in blood-related traits and other traits.

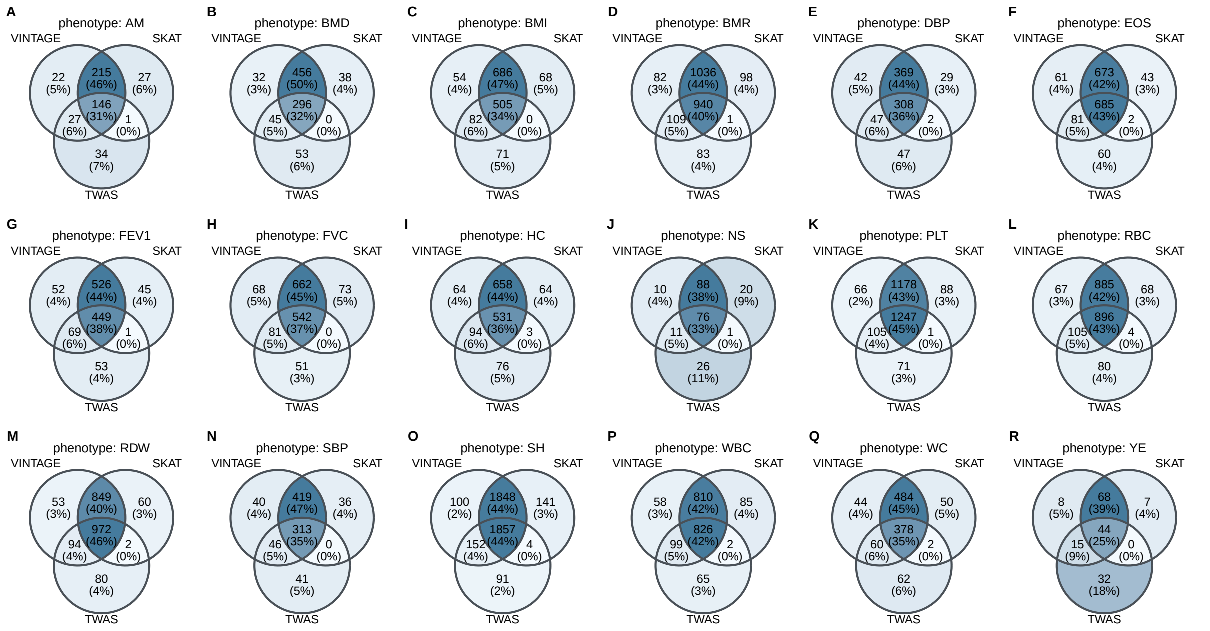


**Figure S40. Venn diagrams comparing the number and relation between the identified genes by different methods in 18 quantitative traits from UKBB.** Compared methods include VINTAGE, SKAT with an unweighted linear kernel, and TWAS that adapts multiple PRS methods to estimate gene expression prediction weights and performs an omnibus test.

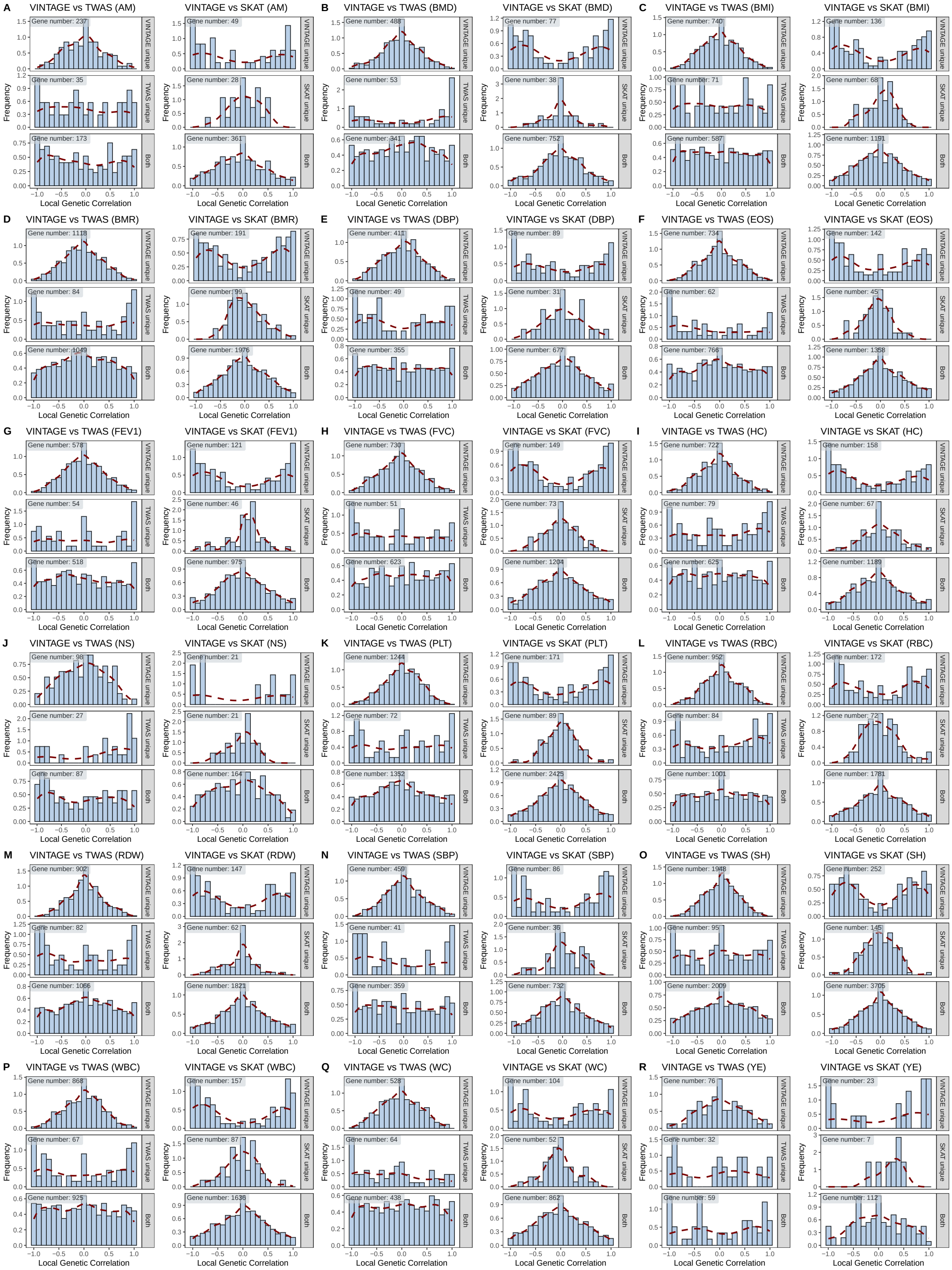


**Figure S41. Histograms of the estimated local genetic correlations by VINTAGE within different sets of identified genes.** For each of the 18 quantitative traits from UKBB, the gene sets include the VINTAGE uniquely identified genes comparing to SKAT, SKAT uniquely identified genes comparing to VINTAGE, genes identified by both VINTAGE and SKAT, VINTAGE uniquely identified genes comparing to TWAS, TWAS uniquely identified genes comparing to VINTAGE, and genes identified by both VINTAGE and TWAS.

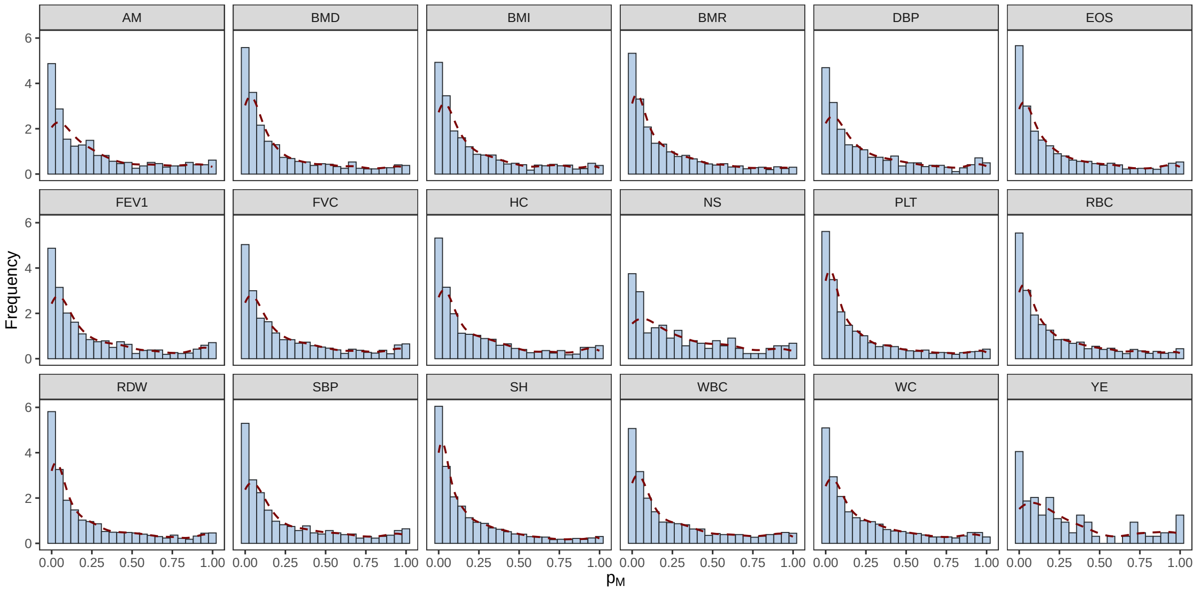


**Figure S42. Histograms of the squared local genetic correlations estimated by VINTAGE across genes that passed the genetic variance tests for 18 UKBB quantitative traits.**

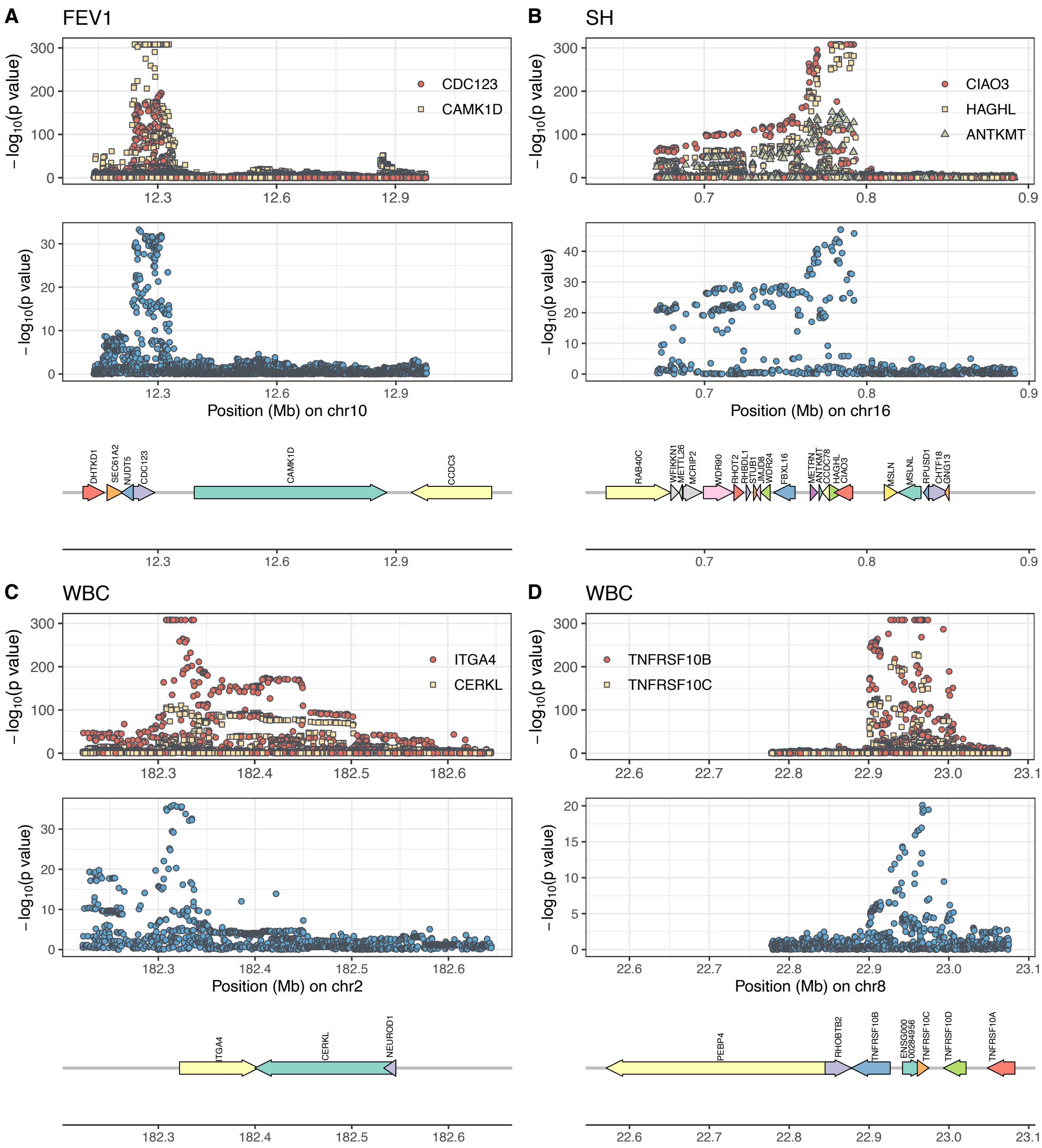


**Figure S43. P-values for SNPs from the eQTL study and GWAS are plotted against their physical positions in four gene regions.** Four sets of genes were identified by VINTAGE in these regions that might be associated with the phenotype due to expression pleiotropy. -$\log_{10}$ p-values are shown for SNPs in the eQTLGen study (top panels) and in the UKBB GWAS (bottom panels). The four gene sets and their associated phenotypes include (**A**) *CDC123* and *CAMK1D* with forced expiratory volume, (**B**) *CIAO3*, *HAGHL*, and *ANTKMT* with standing height, (**C**) *ITGA4* and *CERKL* with white blood cell count, and (**D**) *TNFRSF10B* and *TNFRSF10C* with white blood cell count.

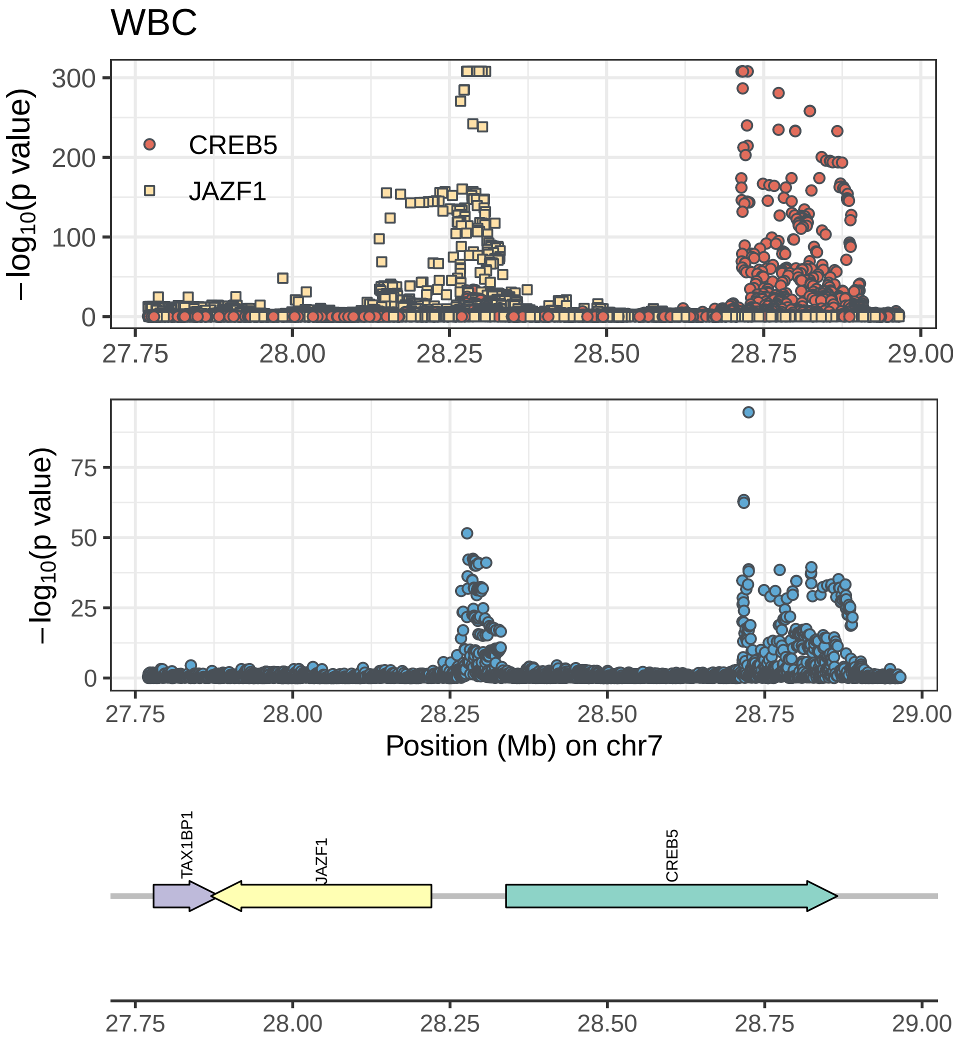


**Figure S44. P-values for SNPs from the eQTL study and GWAS are plotted against their physical positions in the *CREB5/JAZF1* gene region.** *CREB5* and *JAZF1* were identified by VINTAGE to be associated with white blood cell count and the associations do not appear to be driven by expression pleiotropy. -$\log_{10}$ p-values are shown for SNPs in the eQTLGen study (top panels) and in the UKBB GWAS (bottom panels).

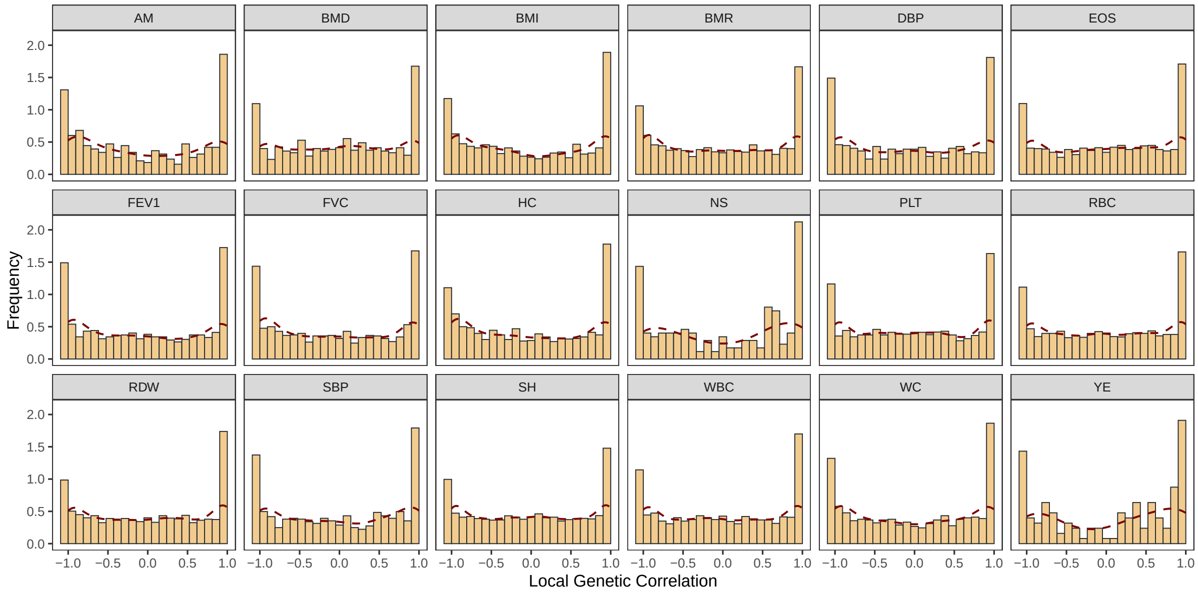


**Figure S45. Histograms of the estimated local genetic correlations by SUPERGNOVA across genes that passed the genetic variance tests for 18 UKBB quantitative traits.** The local genetic correlation estimates from SUPERGNOVA were constrained to -1 or 1 if they fell below -1 or above 1, respectively.

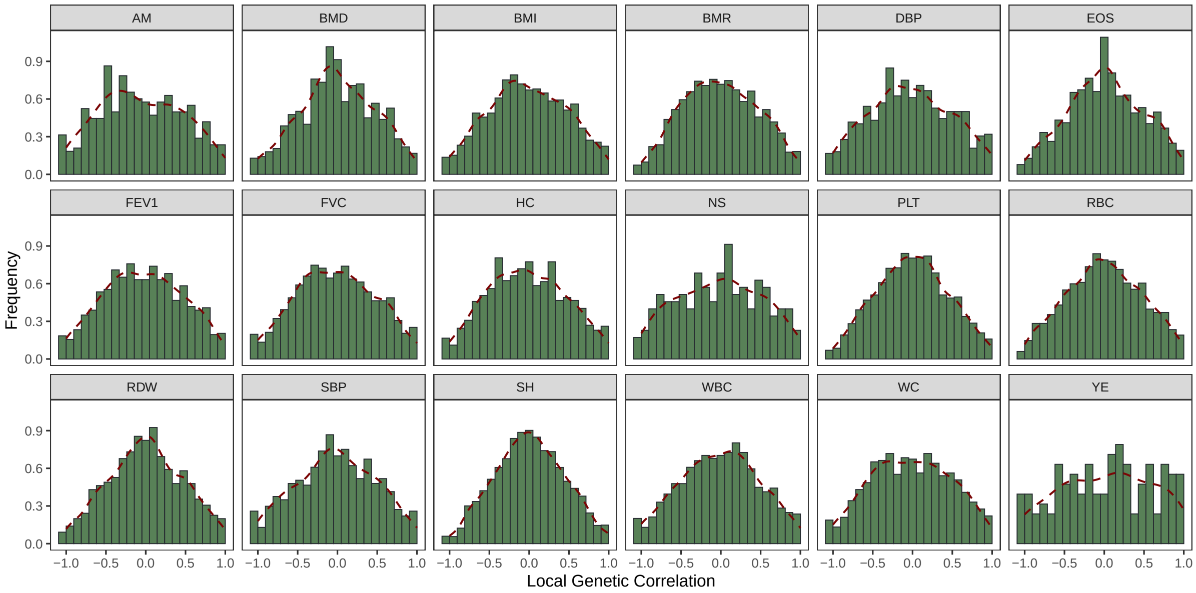


**Figure S46. Histograms of the estimated local genetic correlations by LAVA across genes that passed the genetic variance tests for 18 UKBB quantitative traits.** The local genetic correlation estimates from LAVA were constrained to -1 or 1 if they fell below -1 or above 1, respectively.

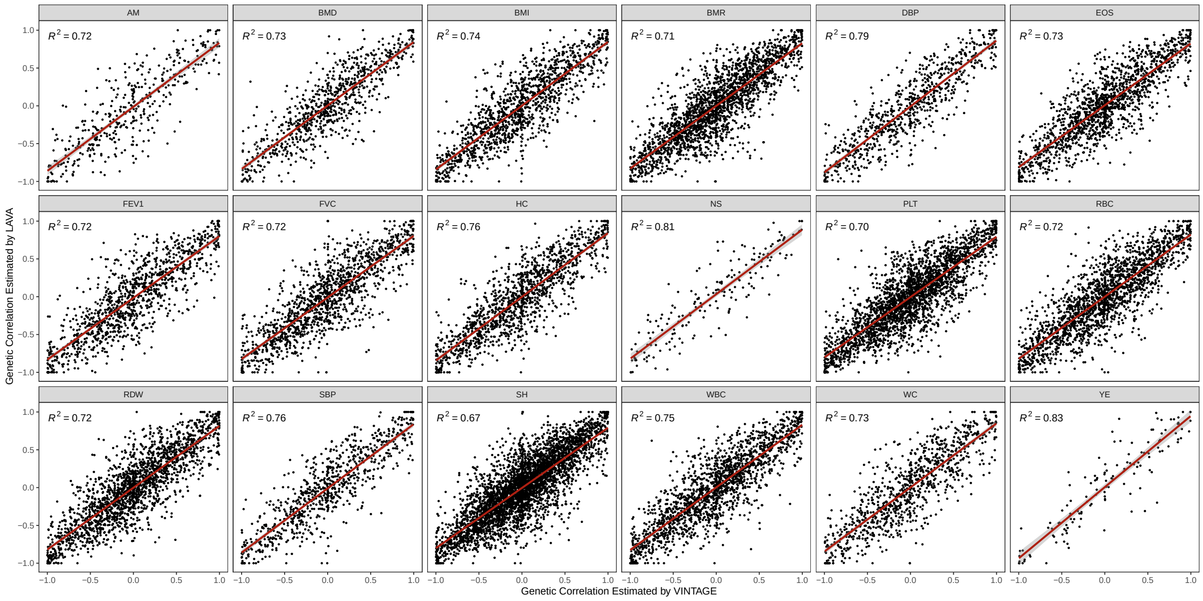


**Figure S47. Scatter plots showing the consistency of the local genetic correlation estimates from VINTAGE and LAVA for 18 UKBB quantitative traits.** Estimates for genes that passed the local genetic variance tests are included. $R^2$ represents the squared correlation between the local genetic correlation estimate from the two methods.

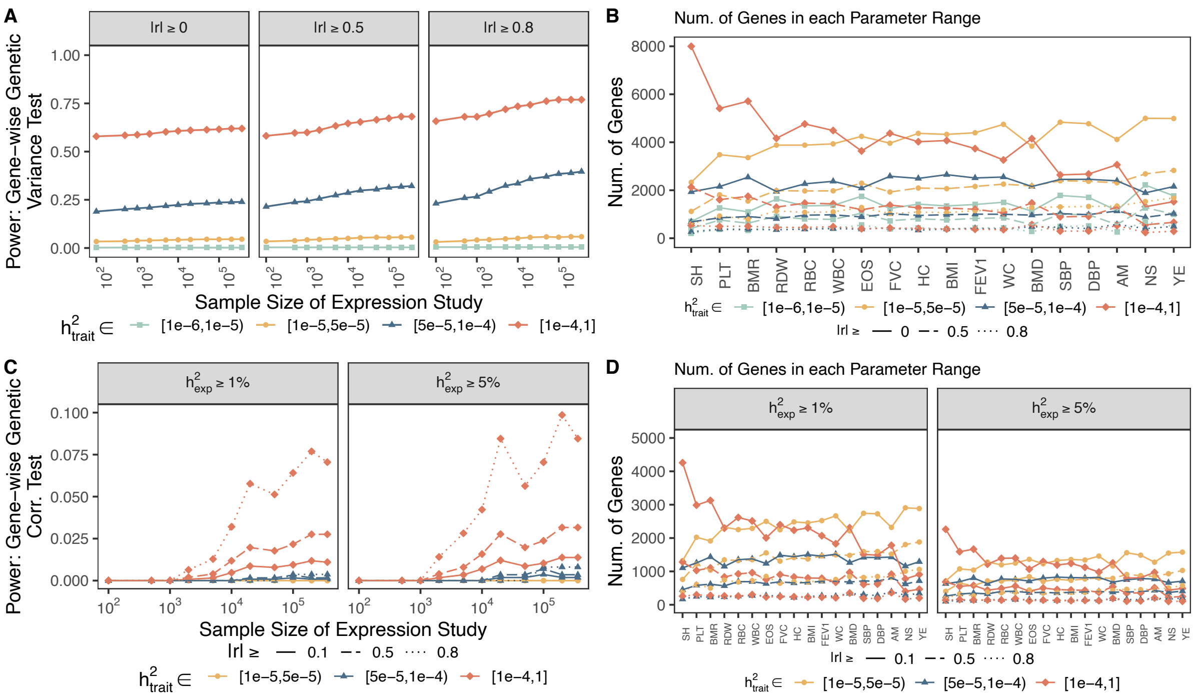


**Figure S48. Evaluating the contribution of gene expression mapping study to gene-wise genetic variance and correlation tests.** Power of (**A**) gene-wise genetic variance tests and (**C**) correlation tests were evaluated in the simulations under different sample sizes of the gene expression mapping study and a fixed sample size of the GWAS. Simulations were conducted for the 13,725 genes examined in the real data analysis with simulation parameters $PVE_1$, $PVE_2$, and $r$ set to be their real data estimates from VINTAGE. In particular, $PVE_1$, $PVE_2$, and $r$ were set to be the cis-SNP heritability estimates of gene expression ($h^2_{exp}$), trait ($h^2_{trait}$), and local genetic correlation between gene expression and the trait in the analysis of eQTLGen data and GWAS data of systolic blood pressure (SBP) from UKBB. Power was evaluated within different parameter ranges. The number of genes in each range of parameters is shown for (**B**) the gene-wise genetic variance tests and (**D**) correlation tests across all 18 traits analyzed in real data applications. The evaluated sample size of the gene expression study was 100, 500, 1,000, 2,000, 5,000, 10,000, 20,000, 50,000, 100,000, 200,000, and 361,194 while the sample size of GWAS was fixed to be 361,194.

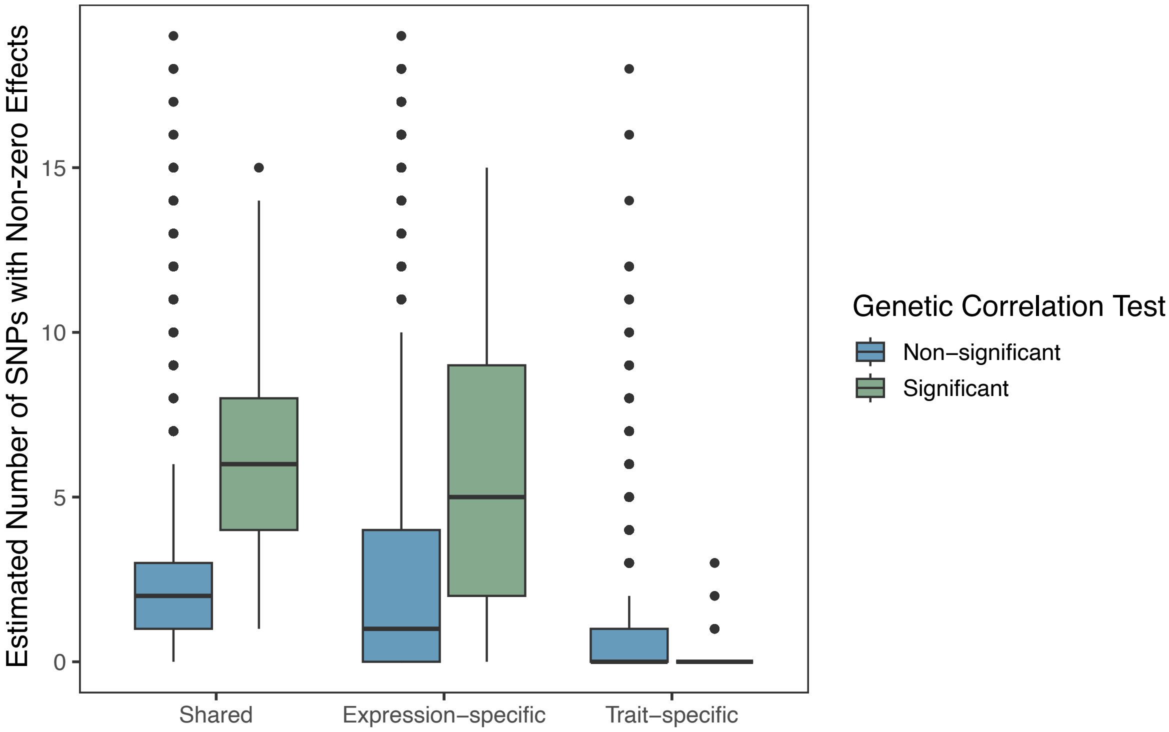


**Figure S49. Estimated number of SNPs with non-zero effects on gene expression and trait by MESuSiE.** Boxplots showing the number of SNPs of each gene with non-zero effects on both gene expression and the trait (shared), only the gene expression (expression-specific), and only the trait (trait-specific), as estimated by MESuSiE. The analysis was conducted for the same 18 quantitative traits in the real data applications, focusing on genes that passed the gene-wise genetic variance test of VINTAGE. The number of SNPs was plotted separately for genes that passed or did not pass the gene-wise genetic correlation test of VINTAGE.

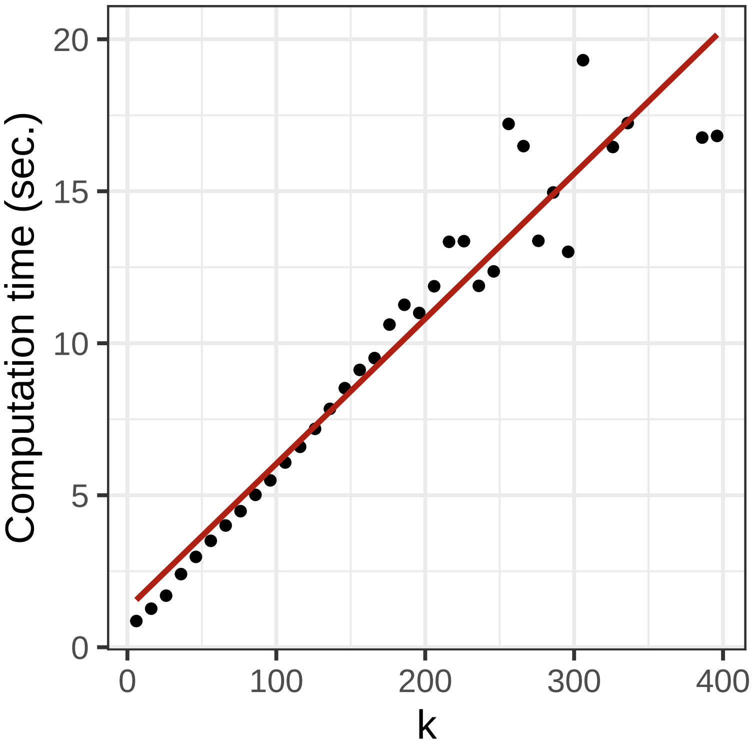


**Figure S50. Computational time of VINTAGE.** Computation was carried out on a single thread of an Intel(R) Xeon(R) Gold 6138 CPU @ 2.00GHz. Genes were binned based on the number of cis-SNPs with the bin width equaling to 100 SNPs. The computational time was calculated as the average time by VINTAGE on analyzing the genes in each bin.

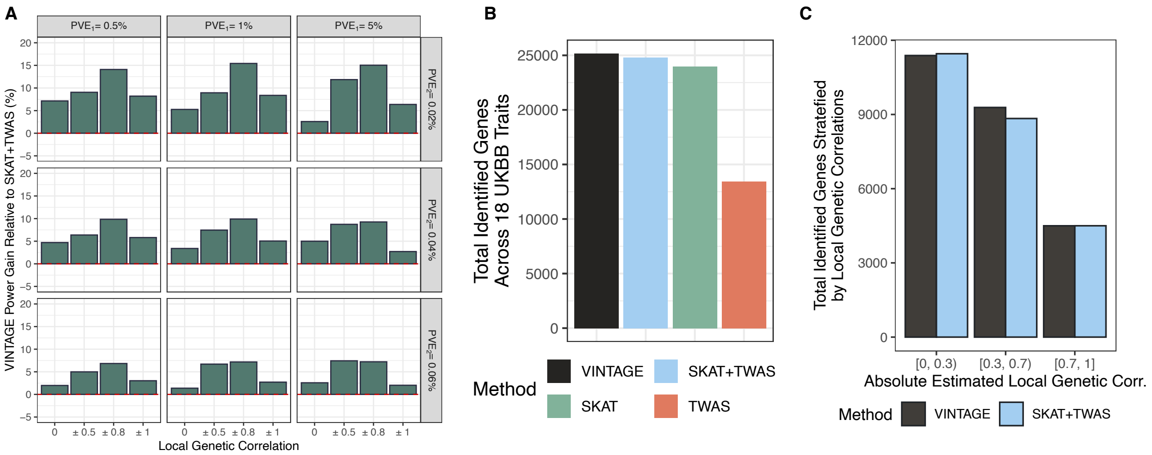


**Figure S51. Evaluating the naive integration approach for the gene-wise genetic variance test.** (**A**) Power of a naive integration approach that first runs SKAT and TWAS separately and then aggregates their p-values into a single p-value using the Cauchy combination approach. This approach is referred to as SKAT+TWAS. Simulations were conducted under different $PVE_1$ values (0.5%, 1%, and 5%), different $PVE_2$ values (0.02%, 0.04%, and 0.06%), and different local genetic correlations (-1.0, -0.8, -0.5, 0.0, 0.5, 0.8, and 1.0) with in-sample LD matrices and a polygenic genetic architecture ($q = p$). (**B**) Total number of identified genes across all 18 traits from UK Biobank identified by different methods in the gene-wise genetic variance tests. Compared methods include VINTAGE, SKAT, TWAS, and SKAT+TWAS. (**C**) Total number of identified genes stratified based on the absolute value of the estimated local genetic correlations. Compared methods include VINTAGE and SKAT+TWAS.

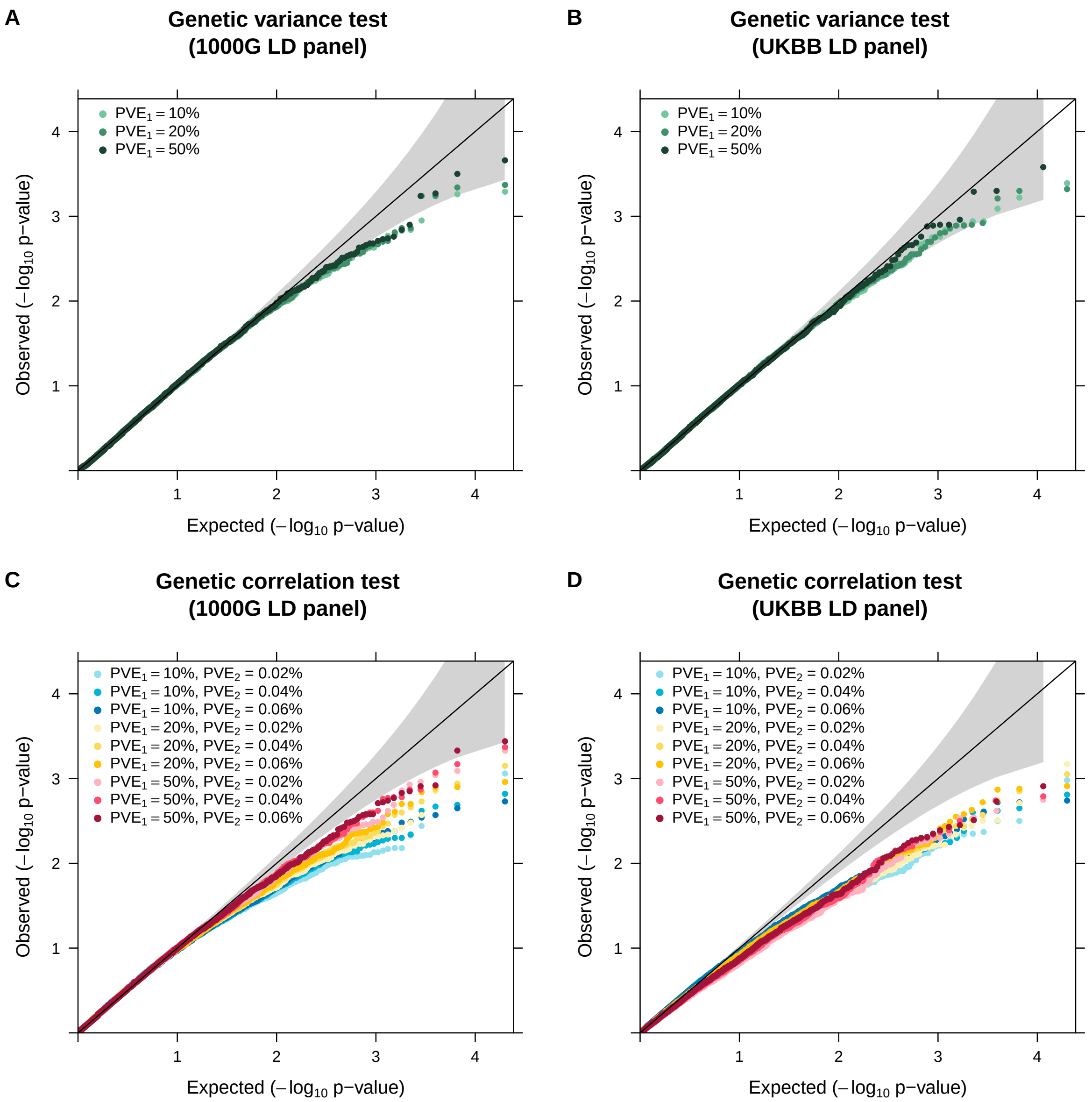


**Figure S52. Influence of unequal LD structures on VINTAGE in the null simulations.** The influence was evaluated for both the (**A-B**) gene-wise genetic variance test and the (**C-D**) gene-wise genetic correlation test. Null simulations were conducted under different $PVE_1$ values (10%, 20%, and 50%) and $PVE_2$ values (0.02%, 0.04%, and 0.06%) with LD matrices computed using (**A**,**C**) 503 European individuals from 1000G or (**B**,**D**) 10,000 WB individuals from UKBB. Null simulations were conducted under a polygenic genetic architecture (q=p).

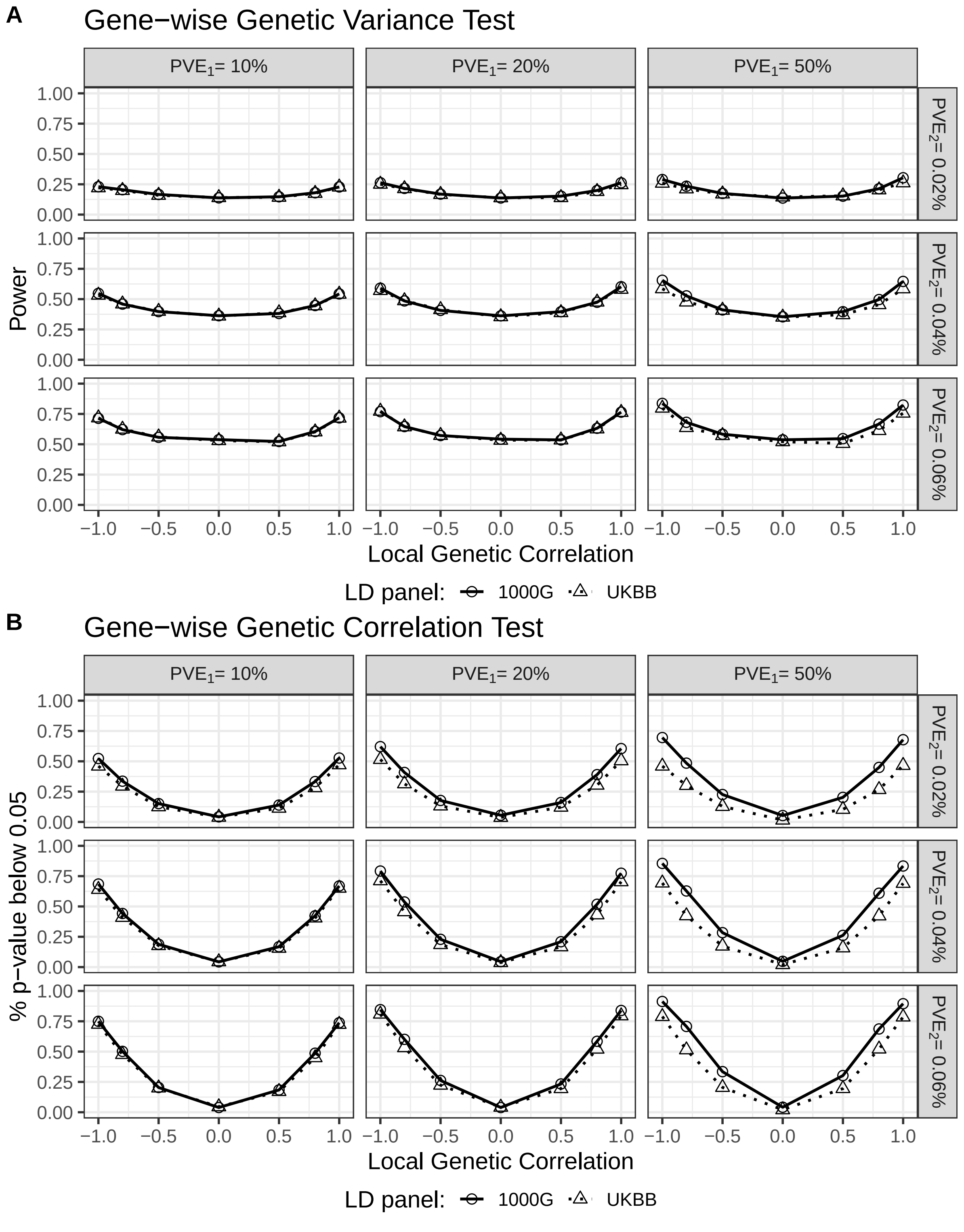


**Figure S53. Influence of unequal LD structures on the power of VINTAGE.** The influence was evaluated for both the (**A**) gene-wise genetic variance test and the (**B**) gene-wise genetic correlation test, with LD matrices computed using either 503 European individuals from 1000G or 10,000 WB individuals from UKBB. Simulations were conducted under different $PVE_1$ values (10%, 20%, and 50%), different $PVE_2$ values (0.02%, 0.04%, and 0.06%), and different local genetic correlations (-1.0, -0.8, -0.5, 0.0, 0.5, 0.8, 1.0) under a polygenic genetic architecture (q=p).

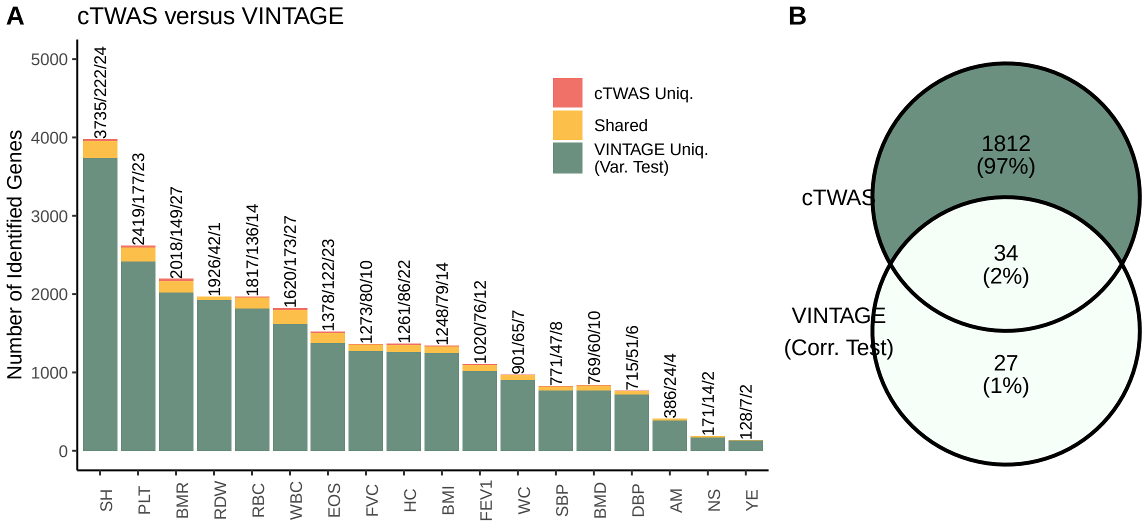


**Figure S54. Exploration of cTWAS in relation to the gene-wise genetic variance and correlation tests in real data applications.** (**A**) Stacked barplots showing the number of genes per trait uniquely identified by the genetic variance test of VINTAGE, uniquely identified by cTWAS, and identified by both methods. The specific numbers are indicated on top of each bar. (**B**) Venn diagram showing the relationship of the genes identified by the gene-wise genetic correlation test of VINTAGE and cTWAS across all 18 traits from UKBB.

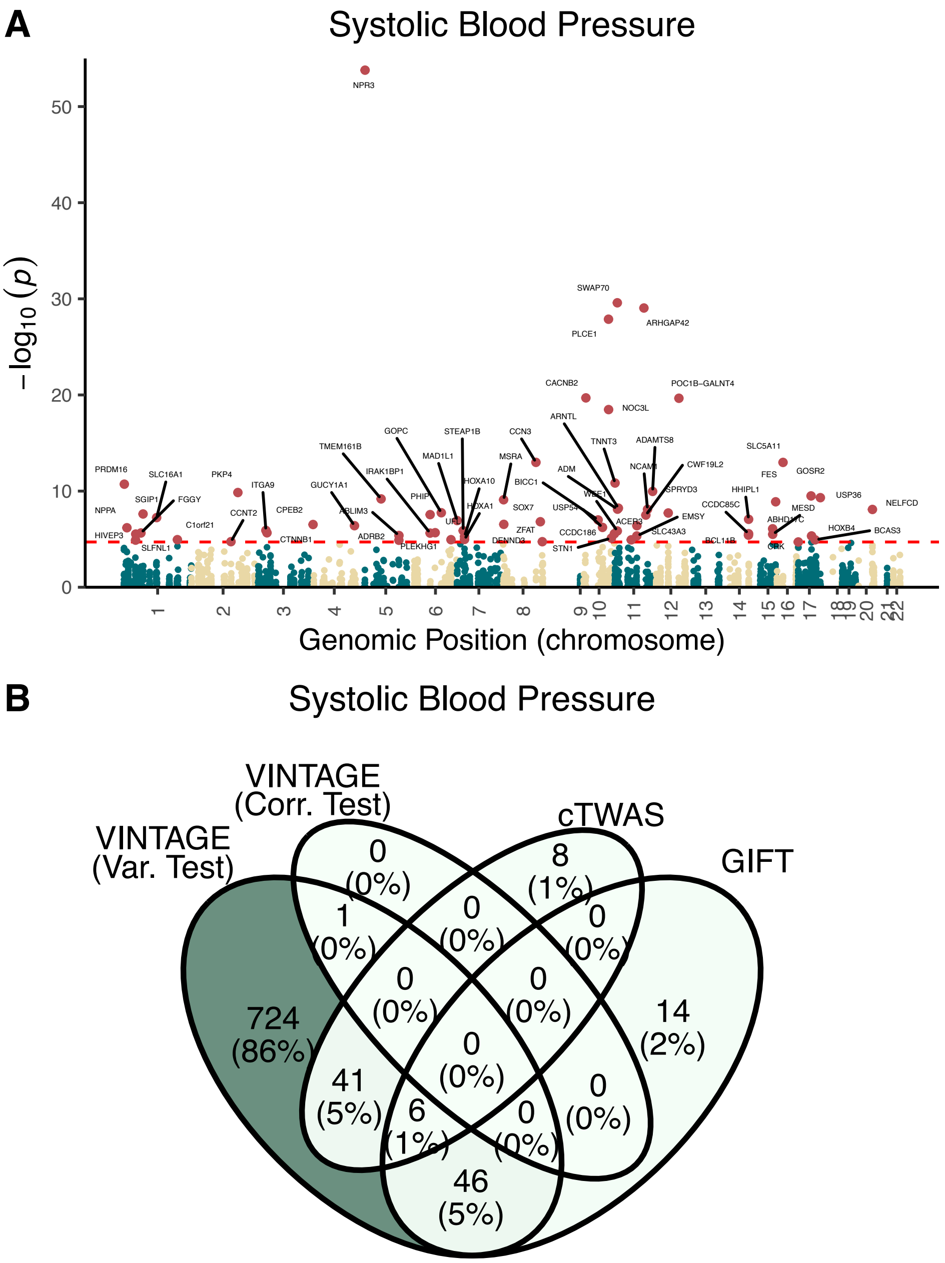


**Figure S55. Exploration of GIFT in relation to the gene-wise genetic variance and correlation tests in the analysis of systolic blood pressure in real data applications.** (**A**) Manhattan plot showing the -$\log_{10}$ p-values of genes from GIFT against their physical positions on the genome. Genes that passed the transcriptome-wide significance threshold of $2.5 \times 10^{-6}$ are colored in red. (**B**) Venn diagram showing the relationship of the genes identified by the gene-wise genetic variance test of VINTAGE, gene-wise genetic correlation test of VINTAGE, cTWAS, and GIFT in the analysis of systolic blood pressure.

| Trait | Gene | Locus | $h_1^2$ | $h_2^2$ | $r$ | p value |
|---|---|---|---|---|---|---|
| BMI | ETV5 | chr3:185764103-185828107 | 0.013 | 0.00023 | -0.99 | $3.3 \times 10^{-5}$ |
| BMI | ADARB1 | chr21:46493768-46646475 | 0.18 | 0.00089 | 0.94 | $4.4 \times 10^{-8}$ |
| BMR | LRMDA | chr10:77191382-78319926 | 0.68 | 0.0012 | -0.83 | $5.0 \times 10^{-6}$ |
| BMR | AXIN2 | chr17:63524681-63557766 | 0.15 | 0.00035 | -0.94 | $7.0 \times 10^{-7}$ |
| DBP | NSG2 | chr5:173472709-173670504 | 0.0025 | 0.00040 | -0.99 | $5.6 \times 10^{-5}$ |
| EOS | BCL2L11 | chr2:111876955-111926022 | 0.018 | 0.0011 | 0.97 | $1.7 \times 10^{-5}$ |
| EOS | GATA2 | chr3:128198270-128212044 | 0.018 | 0.0077 | 0.97 | $5.1 \times 10^{-6}$ |
| EOS | TRERF1 | chr6:42192669-42419789 | 0.10 | 0.0018 | -0.8 | $1.6 \times 10^{-5}$ |
| EOS | ELMO1 | chr7:36892511-37488826 | 0.36 | 0.00073 | -0.89 | $2.0 \times 10^{-6}$ |
| EOS | ICOSLG | chr21:45636897-45660849 | 0.078 | 0.00068 | -0.94 | $1.1 \times 10^{-5}$ |
| EOS | OLIG2 | chr21:34398243-34401504 | 0.014 | 0.00033 | 0.99 | $1.8 \times 10^{-6}$ |
| FEV1 | GM2A | chr5:150591711-150650001 | 0.97 | 0.00029 | -0.99 | $3.6 \times 10^{-5}$ |
| FEV1 | CDC123 | chr10:12237964-12292588 | 0.19 | 0.0010 | -0.94 | $7.3 \times 10^{-6}$ |
| FEV1 | CAMK1D | chr10:12391546-12877545 | 0.62 | 0.0020 | -0.83 | $1.4 \times 10^{-10}$ |
| FEV1 | AXIN2 | chr17:63524681-63557766 | 0.13 | 0.00024 | -0.92 | $1.7 \times 10^{-5}$ |
| FVC | CAMK1D | chr10:12391546-12877545 | 0.62 | 0.00087 | -0.77 | $1.4 \times 10^{-6}$ |
| HC | ADARB1 | chr21:46493768-46646475 | 0.17 | 0.00045 | 0.93 | $3.0 \times 10^{-5}$ |
| PLT | EHD3 | chr2:31457018-31492317 | 0.036 | 0.0031 | -0.9 | $7.8 \times 10^{-6}$ |
| PLT | BAK1 | chr6:33540324-33548070 | 0.61 | 0.021 | -0.83 | $1.8 \times 10^{-6}$ |
| PLT | VLDLR | chr9:2621182-2660056 | 0.21 | 0.00047 | 0.99 | $1.9 \times 10^{-6}$ |
| PLT | RSU1 | chr10:16632610-16859462 | 0.79 | 0.00032 | -1 | $1.2 \times 10^{-6}$ |
| PLT | GRTP1 | chr13:113978478-114018463 | 0.032 | 0.0014 | -0.98 | $2.5 \times 10^{-7}$ |
| PLT | CD33 | chr19:51728320-51747115 | 0.094 | 0.00054 | 0.84 | $1.6 \times 10^{-5}$ |
| PLT | TRABD | chr22:50624342-50638027 | 0.26 | 0.0084 | -0.77 | $3.6 \times 10^{-6}$ |
| RBC | SLC12A7 | chr5:1050499-1112178 | 0.36 | 0.0013 | 0.69 | $2.4 \times 10^{-5}$ |
| RBC | CD36 | chr7:79998891-80308593 | 0.47 | 0.00056 | 0.86 | $3.9 \times 10^{-6}$ |
| RBC | ABO | chr9:136108665-136151440 | 0.70 | 0.0065 | 0.96 | $2.7 \times 10^{-10}$ |
| RBC | LSM4 | chr19:18417046-18433922 | 0.12 | 0.0011 | -0.79 | $8.9 \times 10^{-6}$ |
| RBC | TYMP | chr22:50964181-50968461 | 0.36 | 0.0024 | -0.93 | $1.5 \times 10^{-11}$ |
| RDW | SLC12A7 | chr5:1050499-1112178 | 0.35 | 0.014 | 0.5 | $9.5 \times 10^{-6}$ |
| RDW | KIAA0319 | chr6:24544332-24646419 | 0.19 | 0.0032 | -0.71 | $1.9 \times 10^{-5}$ |
| RDW | CD36 | chr7:79998891-80308593 | 0.41 | 0.00088 | -0.85 | $2.3 \times 10^{-6}$ |
| RDW | ABO | chr9:136108665-136151440 | 0.65 | 0.0023 | -0.87 | $8.6 \times 10^{-6}$ |
| RDW | PSMB11 | chr14:23511421-23513269 | 0.056 | 0.0058 | 0.87 | $5.1 \times 10^{-6}$ |
| RDW | DNAJA4 | chr15:78556428-78574538 | 0.11 | 0.0020 | 0.77 | $1.3 \times 10^{-5}$ |
| RDW | RCCD1 | chr15:91498111-91506355 | 0.15 | 0.0054 | 0.87 | $2.5 \times 10^{-8}$ |
| RDW | TMEM98 | chr17:31254928-31272124 | 0.028 | 0.00098 | -0.98 | $2.1 \times 10^{-5}$ |
| RDW | AFMID | chr17:76183439-76203783 | 0.044 | 0.0048 | -0.73 | $4.2 \times 10^{-6}$ |
| RDW | SIGLEC14 | chr19:52142731-52150078 | 0.80 | 0.00033 | -0.89 | $1.2 \times 10^{-5}$ |
| RDW | TYMP | chr22:50964181-50968461 | 0.38 | 0.00066 | -0.96 | $2.7 \times 10^{-7}$ |
| SBP | CAMK1D | chr10:12391546-12877545 | 0.60 | 0.00051 | 0.81 | $2.8 \times 10^{-5}$ |
| SH | LTBP1 | chr2:33172020-33624576 | 0.010 | 0.0067 | 0.89 | $5.2 \times 10^{-6}$ |
| SH | STK32B | chr4:5053207-5502721 | 0.030 | 0.00043 | 0.98 | $2.6 \times 10^{-6}$ |

Continued on next page

**Table S1 – continued from previous page**

| Trait | Gene | Locus | $h_1^2$ | $h_2^2$ | $r$ | p value |
|---|---|---|---|---|---|---|
| SH | CIAO3 | chr16:779760-791329 | 0.24 | 0.0015 | 0.96 | $2.2 \times 10^{-8}$ |
| SH | HAGHL | chr16:776936-785525 | 0.17 | 0.0010 | 0.85 | $2.8 \times 10^{-6}$ |
| SH | ANTKMT | chr16:770581-772590 | 0.042 | 0.00079 | 0.93 | $6.2 \times 10^{-7}$ |
| SH | AXIN2 | chr17:63524681-63557766 | 0.14 | 0.0016 | -0.82 | $8.0 \times 10^{-6}$ |
| WBC | LAPTM5 | chr1:31205316-31230621 | 0.13 | 0.00070 | 0.97 | $5.6 \times 10^{-6}$ |
| WBC | NLRP3 | chr1:247579458-247612410 | 0.27 | 0.0015 | 0.77 | $9.3 \times 10^{-6}$ |
| WBC | ITGA4 | chr2:182321929-182403667 | 0.26 | 0.0014 | -0.91 | $4.8 \times 10^{-7}$ |
| WBC | CERKL | chr2:182399768-182545392 | 0.097 | 0.0017 | -0.84 | $1.3 \times 10^{-5}$ |
| WBC | SLC12A7 | chr5:1050499-1112178 | 0.41 | 0.00037 | 0.91 | $2.6 \times 10^{-6}$ |
| WBC | CREB5 | chr7:28338940-28865511 | 0.53 | 0.0056 | -0.94 | $2.6 \times 10^{-21}$ |
| WBC | JAZF1 | chr7:27870192-28220414 | 0.36 | 0.0015 | -0.94 | $3.7 \times 10^{-7}$ |
| WBC | TNFRSF10B | chr8:22877646-22926544 | 0.14 | 0.00041 | 0.93 | $1.3 \times 10^{-7}$ |
| WBC | TNFRSF10C | chr8:22960434-22974958 | 0.050 | 0.00052 | -0.9 | $7.6 \times 10^{-6}$ |
| WBC | IFITM3 | chr11:319676-329475 | 0.43 | 0.0024 | 0.7 | $2.9 \times 10^{-7}$ |
| WBC | CHSY1 | chr15:101715932-101792253 | 0.20 | 0.00039 | -0.89 | $1.4 \times 10^{-6}$ |
| WBC | KCTD11 | chr17:7255208-7258263 | 0.054 | 0.00066 | -0.94 | $3.0 \times 10^{-7}$ |
| WBC | TTC39C | chr18:21572737-21715574 | 0.36 | 0.0012 | 0.87 | $8.6 \times 10^{-6}$ |
| WC | ADARB1 | chr21:46493768-46646475 | 0.16 | 0.00042 | 0.96 | $9.7 \times 10^{-6}$ |

**Table S1. Summary of genes displaying significant local genetic correlations as identified by VINTAGE.** The analysis used cis-eQTL summary statistics from the eQTLGen phase I study and GWAS summary statistics of 18 complex traits from UK Biobank. The columns represent the abbreviation of the trait, name of the gene, location of the gene in the format of chromosome, TSS, and TES, estimated heritability of the gene expression ($h_1^2$), estimated heritability of the trait ($h_2^2$), local genetic correlation ($r$), and the p-value of the gene-wise genetic correlation test from VINTAGE.

| Trait | Gene | Locus | $h_1^2$ | $h_2^2$ | $r$ | p value |
|---|---|---|---|---|---|---|
| EOS | CERKL | chr2:182399768-182545392 | 0.62 | 0.00039 | -1.0 | $3.4 \times 10^{-5}$ |
| FVC | CIAO3 | chr16:779760-791329 | 0.36 | 0.00020 | 1.0 | $2.6 \times 10^{-5}$ |
| RDW | HNRNPA1 | chr12:54673977-54680872 | 0.78 | 0.0082 | -0.90 | $1.7 \times 10^{-5}$ |
| SH | ZBTB38 | chr3:141043055-141168634 | 0.11 | 0.0039 | 1.0 | $1.2 \times 10^{-5}$ |
| WBC | JAZF1 | chr7:27870192-28220414 | 0.39 | 0.0012 | -0.95 | $1.1 \times 10^{-5}$ |
| WBC | LPAR1 | chr9:113635543-113800750 | 0.83 | 0.00099 | 0.96 | $2.7 \times 10^{-5}$ |

**Table S2. Summary of genes displaying significant local genetic correlations as identified by VINTAGE.** The analysis used cis-eQTL summary statistics in whole blood samples from the GTEx v8 and GWAS summary statistics of 18 complex traits from UK Biobank. The columns represent the abbreviation of the trait, name of the gene, location of the gene in the format of chromosome, TSS, and TES, estimated heritability of the gene expression ($h_1^2$), estimated heritability of the trait ($h_2^2$), local genetic correlation ($r$), and the p-value of the gene-wise genetic correlation test from VINTAGE.
