## Supplemental Texts for "An alternative framework for transcriptome-wide association studies to detect and decipher gene-trait associations"

**4 Gene-wise genetic correlation test** **22**

**5 Additional simulation and real data analyses** **24**

5.1 Evaluation of COJO and MESuSiE for estimating the number of non-zero effect SNPs 24

5.2 A complication for evaluating type I error and power under a sparse genetic architecture 24

5.3 Exploration of expression pleiotropy . . . . . 25

### 1 VINTAGE model description

#### 1.1 Model setup

We consider the integrative analysis of two genetic studies: a gene expression mapping study and a genome-wide association study (GWAS) to both identify genes associated with a trait of interest and decipher such associations. The two studies are assumed to have no sample overlap. We analyze one gene at a time. For the gene of focus, we denote $\boldsymbol{y}_1$ as the $n_1$-vector of gene expression measurements on $n_1$ individuals in the gene expression mapping study and denote $\boldsymbol{X}_1$ as the corresponding $n_1 \times p$ genotype matrix for $p$ cis-SNPs that reside in the genic and adjacent regulatory regions of the gene. We denote $\boldsymbol{y}_2$ as the $n_2$-vector of trait measurements in the GWAS and denote $\boldsymbol{X}_2$ as the corresponding $n_2 \times p$ genotype matrix for the same set of $p$ cis-SNPs. We center and standardize $\boldsymbol{y}_1$, $\boldsymbol{y}_2$, and each column of $\boldsymbol{X}_1$ and $\boldsymbol{X}_2$ to have a mean of zero and variance of one. We model the relationship among SNPs, gene expression, and the trait with the following two equations:

$$\boldsymbol{y}_1 = \boldsymbol{X}_1\boldsymbol{\beta}_1 + \boldsymbol{\epsilon}_1, \quad \boldsymbol{\epsilon}_1 \sim N(\boldsymbol{0}, \sigma_1^2 \boldsymbol{I}_{n_1}), \tag{1}$$

$$\boldsymbol{y}_2 = \boldsymbol{X}_2\boldsymbol{\beta}_2 + \boldsymbol{\epsilon}_2, \quad \boldsymbol{\epsilon}_2 \sim N(\boldsymbol{0}, \sigma_2^2 \boldsymbol{I}_{n_2}), \tag{2}$$

where $\boldsymbol{\beta}_1$ is a $p$-vector of genetic effects on gene expression and $\boldsymbol{\beta}_2$ is a $p$-vector of genetic effects on the trait. We further assume that the genetic effects of the $j$th SNP on the gene expression and on the trait may be correlated and follow a bivariate normal distribution:

$$\begin{aligned} &(\beta_{1,j}, \beta_{2,j})^T \sim N(\boldsymbol{0}, \boldsymbol{D}), \\ &\boldsymbol{D} = \begin{pmatrix} \sigma_{\beta_1}^2 & \rho \\ \rho & \sigma_{\beta_2}^2 \end{pmatrix}, \end{aligned} \tag{3}$$

where $\sigma_{\beta_1}^2$ represents the per-SNP heritability of the gene expression; $\sigma_{\beta_2}^2$ represents the per-SNP heritability of the trait; and $\rho$ represents the per-SNP local genetic covariance between the gene expression and the trait. We can further obtain the local genetic covariance, local genetic correlation, and local heritability for gene expression and trait as:

$$\rho_g = p\rho, \tag{4}$$

$$r = \rho / \sqrt{\sigma_{\beta_1}^2 \sigma_{\beta_2}^2}, \tag{5}$$

$$h_1^2 = p\sigma_{\beta_1}^2, \tag{6}$$

$$h_2^2 = p\sigma_{\beta_2}^2. \tag{7}$$

For mathematical convenience, we further denote $\boldsymbol{y} = (\boldsymbol{y}_1^T, \boldsymbol{y}_2^T)$, $\boldsymbol{X} = \begin{pmatrix} \boldsymbol{X}_1 & \\ & \boldsymbol{X}_2 \end{pmatrix}$, $\boldsymbol{\beta} =$

$(\boldsymbol{\beta}_1^T, \boldsymbol{\beta}_2^T)^T$, and $\boldsymbol{\epsilon} = (\boldsymbol{\epsilon}_1^T, \boldsymbol{\epsilon}_2^T)^T$. Hence, equations (1) and (2) can be consolidated into:

$$\boldsymbol{y} = \boldsymbol{X}\boldsymbol{\beta} + \boldsymbol{\epsilon}, \quad \boldsymbol{\epsilon} \sim N(\boldsymbol{0}, \boldsymbol{\Sigma}), \tag{8}$$

where $\boldsymbol{\Sigma} = \begin{pmatrix} \sigma_1^2 \boldsymbol{I}_{n_1} & \\ & \sigma_2^2 \boldsymbol{I}_{n_2} \end{pmatrix}$ and $\boldsymbol{\beta} \sim N(\boldsymbol{0}, \boldsymbol{D} \otimes \boldsymbol{I}_p)$ *a priori*.

##### 1.2 Mediation analysis model

The model described in section 1.1 can be re-parameterized to form a mediation analysis model. Specifically, the assumption made on $\beta_{1,j}$ and $\beta_{2,j}$ (equation 3) can be equivalently expressed as:

$$\beta_{1,j} = \sigma_{\beta_1} b_{1,j}, \tag{9}$$

$$\beta_{2,j} = \sigma_{\beta_2}(r b_{1,j} + \sqrt{1 - r^2} b_{2,j}), \tag{10}$$

where $b_{1,j}$ and $b_{2,j}$ are independent latent variables that follow a standard normal distribution *a priori*. Plugging equations (9) and (10) into equation (2), we have:

$$\begin{aligned} \boldsymbol{y}_2 &= \boldsymbol{X}_2 \left[ \sigma_{\beta_2}(r\boldsymbol{b}_1 + \sqrt{1 - r^2}\boldsymbol{b}_2) \right] + \boldsymbol{\epsilon}_2 \\ &= r\sigma_{\beta_2}\boldsymbol{X}_2\boldsymbol{b}_1 + \sqrt{1 - r^2}\sigma_{\beta_2}\boldsymbol{X}_2\boldsymbol{b}_2 + \boldsymbol{\epsilon}_2 \\ &= r\frac{\sigma_{\beta_2}}{\sigma_{\beta_1}}\boldsymbol{X}_2\boldsymbol{\beta}_1 + \sqrt{1 - r^2}\sigma_{\beta_2}\boldsymbol{X}_2\boldsymbol{b}_2 + \boldsymbol{\epsilon}_2 \\ &= r\frac{\sigma_{\beta_2}}{\sigma_{\beta_1}}\tilde{\boldsymbol{y}}_1 + \sqrt{1 - r^2}\sigma_{\beta_2}\boldsymbol{X}_2\boldsymbol{b}_2 + \boldsymbol{\epsilon}_2, \end{aligned} \tag{11}$$

where $\tilde{\boldsymbol{y}}_1 = \boldsymbol{X}_2\boldsymbol{\beta}_1$ represents the $n_2$-vector of genetically regulated gene expression for the $n_2$ individuals in the GWAS. Equations (1) and (11) form a mediation analysis model where SNPs are exposures, gene expression is the mediator, and the trait is the outcome [1]. In the mediation analysis model, $\sqrt{1 - r^2}\sigma_{\beta_2}\boldsymbol{b}_2$ is the vector of direct effects (DE) of the SNPs on the trait while the product term $r\frac{\sigma_{\beta_2}}{\sigma_{\beta_1}}\boldsymbol{\beta}_1 = r\sigma_{\beta_2}\boldsymbol{b}_1$ is the vector of indirect effects (IE) of the SNPs on the trait. We define:

$$p_M := \frac{\mathbb{E}[S_{IE}^2]}{\mathbb{E}[S_{IE}^2] + \mathbb{E}[S_{DE}^2]} = \frac{r^2\sigma_{\beta_2}^2\mathbb{E}[S_{\boldsymbol{b}_1}^2]}{r^2\sigma_{\beta_2}^2\mathbb{E}[S_{\boldsymbol{b}_1}^2] + (1 - r^2)\sigma_{\beta_2}^2\mathbb{E}[S_{\boldsymbol{b}_2}^2]} = r^2, \tag{12}$$

as the proportion of variance attributed to mediation effects where $S^2$ represents sample variance. When $r = 0$, we have $p_M = 0$, indicating that the gene expression does not mediate any of the genetic effects on the trait. When $r = \pm 1$, we have $p_M = 1$, indicating that the genetic effects on the trait are entirely mediated through gene expression.

#### 2 Model parameter estimation

In this section, we present a scalable algorithm for estimating model parameters. We will first introduce a transformation of the current model to allow for scalable computation. Then, we provide derivations of an efficient parameter-expanded expectation maximization (PX-EM) algorithm for parameter estimation [2]. Finally, we extend our derivations to use only summary statistics from the gene expression mapping study and GWAS for inference.

##### 2.1 Model transformation

To enable scalable computation, we transform the genotype matrices and their corresponding coefficients into the principal component (PC) space. Specifically, we re-parameterize the models defined in equations (1) and (2) into the following transformed models:

$$\boldsymbol{y}_1 = (\boldsymbol{X}_1\boldsymbol{Q})(\boldsymbol{Q}^T\boldsymbol{\beta}_1) + \boldsymbol{\epsilon}_1, \tag{13}$$

$$\boldsymbol{y}_2 = (\boldsymbol{X}_2\boldsymbol{Q})(\boldsymbol{Q}^T\boldsymbol{\beta}_2) + \boldsymbol{\epsilon}_2, \tag{14}$$

where $\boldsymbol{Q}$ is a $p \times p$ orthogonal matrix of eigenvectors derived from the eigendecomposition of the LD matrices $\boldsymbol{R}_l$:

$$\boldsymbol{R}_l = \boldsymbol{Q}\boldsymbol{\Lambda}_l\boldsymbol{Q}^T, \tag{15}$$

where $\boldsymbol{R}_l = \frac{1}{n_l}\boldsymbol{X}_l^T\boldsymbol{X}_l$, for $l = 1$ and 2. Here, we have assumed that the two LD matrices, one from the gene expression mapping study ($l = 1$) and the other from the GWAS ($l = 2$), share the same eigenvectors, which approximately holds when both studies are conducted in the same genetic ancestry. We denote the transformed design matrices and the corresponding regression coefficients to be $\boldsymbol{X}_l^* = \boldsymbol{X}_l\boldsymbol{Q}$ and $\boldsymbol{\beta}_l^* = \boldsymbol{Q}^T\boldsymbol{\beta}_l$, respectively, and work on the following transformed models hereafter for parameter estimation:

$$\boldsymbol{y}_1 = \boldsymbol{X}_1^*\boldsymbol{\beta}_1^* + \boldsymbol{\epsilon}_1, \tag{16}$$

$$\boldsymbol{y}_2 = \boldsymbol{X}_2^*\boldsymbol{\beta}_2^* + \boldsymbol{\epsilon}_2, \tag{17}$$

where the likelihood about all the parameters remains unchanged because of the orthogonality of $\boldsymbol{Q}$ (i.e., $\boldsymbol{Q}\boldsymbol{Q}^T = \boldsymbol{Q}^T\boldsymbol{Q} = \boldsymbol{I}_p$). In particular, $\boldsymbol{\beta}_l^*$ still follows the same distribution as described in (3) *a priori* because of the following:

$$\mathrm{Var}[\boldsymbol{\beta}_l^*] = \boldsymbol{Q}^T\,\mathrm{Var}[\boldsymbol{\beta}_l]\boldsymbol{Q} = \sigma_{\beta_l}^2\boldsymbol{I}_p, \tag{18}$$

$$\mathrm{Cov}[\boldsymbol{\beta}_1^*, \boldsymbol{\beta}_2^*] = \boldsymbol{Q}^T\mathbb{E}[\boldsymbol{\beta}_1\boldsymbol{\beta}_2^T]\boldsymbol{Q} = \rho\boldsymbol{I}_p. \tag{19}$$

As a result of such transformation, columns of $\boldsymbol{X}_l^*$ become decorrelated, which substantially improves the computational speed of the PX-EM algorithm. After the initial eigendecomposition that incurs a cubic computational complexity in $p$, the computational complexity of the PX-EM algorithm becomes linear in $p$ in each iteration.

Similarly for mathematical convenience, we further denote $\boldsymbol{X}^* = \begin{pmatrix} \boldsymbol{X}_1^* & \\ & \boldsymbol{X}_2^* \end{pmatrix}$ and $\boldsymbol{\beta}^* = ((\boldsymbol{\beta}_1^*)^T, (\boldsymbol{\beta}_2^*)^T)^T$. Hence, equations (16) and (17) can be consolidated into a single equation as follows:

$$\boldsymbol{y} = \boldsymbol{X}^* \boldsymbol{\beta}^* + \boldsymbol{\epsilon}. \tag{20}$$

#### 2.2 PX-EM algorithm

Here, we derive an efficient PX-EM algorithm for parameter estimation. PX-EM algorithm improves the convergence rate of the original EM algorithm by introducing additional expansion parameters, while maintaining its simplicity and stability. These expansion parameters capture the bias induced by the incorrect imputation model in the E-step and are used to adjust for the estimates of the parameters of interest. We denote all the parameters to be estimated by $\boldsymbol{\theta} = (\sigma_{\beta_1}^2, \sigma_{\beta_2}^2, \sigma_1^2, \sigma_2^2, \rho)$.

##### 2.2.1 Estimation under the full model

We consider the parameter-expanded version of our model as follows:

$$\boldsymbol{y}_1 = \alpha_1 \boldsymbol{X}_1^* \boldsymbol{\beta}_1^* + \boldsymbol{\epsilon}_1, \tag{21}$$

$$\boldsymbol{y}_2 = \alpha_2 \boldsymbol{X}_2^* \boldsymbol{\beta}_2^* + \boldsymbol{\epsilon}_2, \tag{22}$$

where $\alpha_1$ and $\alpha_2$ are expansion parameters. The reduction function for mapping parameters in the expanded space to the original space is defined as:

$$R(\tilde{\boldsymbol{\theta}}, \alpha_1, \alpha_2) = (\alpha_1^2 \tilde{\sigma}_{\beta_1}^2, \alpha_2^2 \tilde{\sigma}_{\beta_2}^2, \alpha_1 \alpha_2 \tilde{\rho}, \tilde{\sigma}_1^2, \tilde{\sigma}_2^2), \tag{23}$$

where $\tilde{\boldsymbol{\theta}}$ are parameters of interest in the expanded space. When both $\alpha_1$ and $\alpha_2$ are set to their null value of 1, we have $\tilde{\boldsymbol{\theta}} = \boldsymbol{\theta}$. To simplify the derivation, we consolidate equations (21)-(22) into a single equation as follows:

$$\boldsymbol{y} = \boldsymbol{X}^* (\boldsymbol{\alpha} \otimes \boldsymbol{I}_p) \boldsymbol{\beta}^* + \boldsymbol{\epsilon}, \tag{24}$$

where $\boldsymbol{\alpha} = diag(\alpha_1, \alpha_2)$ is a $2 \times 2$ matrix with $\alpha_1$ and $\alpha_2$ on the diagonal and zeros elsewhere. We then follow procedures of a traditional EM algorithm to maximize the likelihood with respect to the equation (24).

In the E-step, we derive the $Q$ function as the expected value of the complete-data log-likelihood, with respect to the current conditional distribution of $\boldsymbol{\beta}^*$ given the observed data and the current estimates of the parameters $\boldsymbol{\theta}^{(t)}$. These quantities are presented in the following:

1. The complete-data log-likelihood function is given by:

$$
\begin{aligned}
&\log L(\boldsymbol{\alpha}, \tilde{\boldsymbol{\theta}}; \boldsymbol{y}, \boldsymbol{\beta}^*) \\
=&\log f(\boldsymbol{y}|\boldsymbol{\beta}^*; \boldsymbol{\alpha}, \tilde{\boldsymbol{\theta}}) f(\boldsymbol{\beta}^*; \boldsymbol{\alpha}, \tilde{\boldsymbol{\theta}}) \\
=& -\frac{n_1+n_2+2p}{2} - \frac{1}{2}\log|\tilde{\boldsymbol{\Sigma}}| - \frac{1}{2}\log|\tilde{\boldsymbol{D}} \otimes \boldsymbol{I}_p| - \frac{1}{2}\boldsymbol{y}^T\tilde{\boldsymbol{\Sigma}}^{-1}\boldsymbol{y} \\
&- \frac{1}{2}(\boldsymbol{\beta}^*)^T \left[(\boldsymbol{\alpha}^T \otimes \boldsymbol{I}_p)(\boldsymbol{X}^*)^T\tilde{\boldsymbol{\Sigma}}^{-1}\boldsymbol{X}^*(\boldsymbol{\alpha} \otimes \boldsymbol{I}_p) + (\tilde{\boldsymbol{D}} \otimes \boldsymbol{I}_p)^{-1}\right]\boldsymbol{\beta}^* \\
&+ (\boldsymbol{\beta}^*)^T(\boldsymbol{\alpha}^T \otimes \boldsymbol{I}_p)(\boldsymbol{X}^*)^T\tilde{\boldsymbol{\Sigma}}^{-1}\boldsymbol{y}. 
\end{aligned} \tag{25}
$$

2. The conditional distribution of $\boldsymbol{\beta}^*$ is given by:

$$
\boldsymbol{\beta}^*|\boldsymbol{y}, \boldsymbol{\theta}^{(t)} \sim N(\boldsymbol{\mu}_{\beta^*}^{(t)}, \boldsymbol{\Sigma}_{\beta^*}^{(t)}), \tag{26}
$$

where

$$
\boldsymbol{\Sigma}_{\beta^*}^{(t)} = \left[(\boldsymbol{X}^*)^T(\boldsymbol{\Sigma}^{(t)})^{-1}\boldsymbol{X}^* + (\boldsymbol{D}^{(t)} \otimes \boldsymbol{I}_p)^{-1}\right]^{-1}, \tag{27}
$$

$$
\boldsymbol{\mu}_{\beta^*}^{(t)} = \boldsymbol{\Sigma}_{\beta^*}^{(t)}(\boldsymbol{X}^*)^T(\boldsymbol{\Sigma}^{(t)})^{-1}\boldsymbol{y}. \tag{28}
$$

3. The Q function is given by:

$$
\begin{aligned}
&Q(\tilde{\boldsymbol{\theta}}, \boldsymbol{\alpha}|\boldsymbol{\theta}^{(t)}) \\
=&\mathbb{E}\left[\log L(\boldsymbol{\alpha}, \tilde{\boldsymbol{\theta}}; \boldsymbol{y}, \boldsymbol{\beta}^*)|\boldsymbol{y}, \boldsymbol{\theta}^{(t)}\right] \\
=& -\frac{n_1+n_2+2p}{2}\log 2\pi - \frac{1}{2}\log|\tilde{\boldsymbol{\Sigma}}| - \frac{p}{2}\log|\tilde{\boldsymbol{D}}| - \frac{1}{2}\boldsymbol{y}^T\tilde{\boldsymbol{\Sigma}}^{-1}\boldsymbol{y} \\
&- \frac{1}{2}(\boldsymbol{\mu}_{\beta^*}^{(t)})^T \left[(\boldsymbol{\alpha}^T \otimes \boldsymbol{I}_p)(\boldsymbol{X}^*)^T\tilde{\boldsymbol{\Sigma}}^{-1}\boldsymbol{X}^*(\boldsymbol{\alpha} \otimes \boldsymbol{I}_p) + (\tilde{\boldsymbol{D}} \otimes \boldsymbol{I}_p)^{-1}\right]\boldsymbol{\mu}_{\beta^*}^{(t)} \\
&- \frac{1}{2}tr\left\{\boldsymbol{\Sigma}_{\beta^*}^{(t)}\left[(\boldsymbol{\alpha}^T \otimes \boldsymbol{I}_p)(\boldsymbol{X}^*)^T\tilde{\boldsymbol{\Sigma}}^{-1}\boldsymbol{X}^*(\boldsymbol{\alpha} \otimes \boldsymbol{I}_p) + (\tilde{\boldsymbol{D}} \otimes \boldsymbol{I}_p)^{-1}\right]\right\} \\
&+ (\boldsymbol{\mu}_{\beta^*}^{(t)})^T(\boldsymbol{\alpha}^T \otimes \boldsymbol{I}_p)(\boldsymbol{X}^*)^T\tilde{\boldsymbol{\Sigma}}^{-1}\boldsymbol{y}.
\end{aligned} \tag{29}
$$

In the M-step, we find the parameters $\tilde{\boldsymbol{\theta}}^{(t+1)}$ and $\boldsymbol{\alpha}^{(t+1)}$ that maximize the $Q$ function using the block coordinate descent algorithm. Briefly, we maximize the $Q$ function along each set of parameters in $(\tilde{\boldsymbol{\theta}}, \boldsymbol{\alpha})$ by setting their gradients to $\boldsymbol{0}$, while fixing the other parameters. We update the parameters as follows:

1. Update $\tilde{\boldsymbol{D}}$:

$$
\tilde{\boldsymbol{D}}^{(t+1)} = \frac{1}{p}\begin{pmatrix} (\boldsymbol{\mu}_{\beta_1^*}^{(t)})^T\boldsymbol{\mu}_{\beta_1^*}^{(t)} + tr(\boldsymbol{\Sigma}_{\beta_{1,1}^*}^{(t)}) & (\boldsymbol{\mu}_{\beta_1^*}^{(t)})^T\boldsymbol{\mu}_{\beta_2^*}^{(t)} + tr(\boldsymbol{\Sigma}_{\beta_{1,2}^*}^{(t)}) \\ (\boldsymbol{\mu}_{\beta_2^*}^{(t)})^T\boldsymbol{\mu}_{\beta_1^*}^{(t)} + tr(\boldsymbol{\Sigma}_{\beta_{2,1}^*}^{(t)}) & (\boldsymbol{\mu}_{\beta_2^*}^{(t)})^T\boldsymbol{\mu}_{\beta_2^*}^{(t)} + tr(\boldsymbol{\Sigma}_{\beta_{2,2}^*}^{(t)}) \end{pmatrix}. \tag{30}
$$

2. Update $\alpha_l$ for $l = 1$ and 2:

$$\alpha_l^{(t+1)} = \frac{(\boldsymbol{\mu}_{\beta_l^*}^{(t)})^T (\boldsymbol{X}_l^*)^T \boldsymbol{y}_l}{(\boldsymbol{\mu}_{\beta_l^*}^{(t)})^T (\boldsymbol{X}_l^*)^T \boldsymbol{X}_l^* \boldsymbol{\mu}_{\beta_l^*}^{(t)} + tr\left\{\boldsymbol{\Sigma}_{\beta_{l,l}^*}^{(t)} (\boldsymbol{X}_l^*)^T \boldsymbol{X}_l^*\right\}}. \tag{31}$$

3. Update $\tilde{\sigma}_l^2$ for $l = 1$ and 2:

$$\begin{aligned}(\tilde{\sigma}_l^2)^{(t+1)} =& \frac{1}{n_l}\left(\boldsymbol{y}_l - \alpha_l^{(t+1)} \boldsymbol{X}_l^* \boldsymbol{\mu}_{\beta_l^*}^{(t)}\right)^T \left(\boldsymbol{y}_l - \alpha_l^{(t+1)} \boldsymbol{X}_l^* \boldsymbol{\mu}_{\beta_l^*}^{(t)}\right) \\ &+ \frac{1}{n_l}(\alpha_l^{(t+1)})^2 tr\left[(\boldsymbol{X}_l^*)^T \boldsymbol{X}_l^* \boldsymbol{\Sigma}_{\boldsymbol{\beta}_{l,l}^*}^{(t)}\right]. \end{aligned} \tag{32}$$

Above, $\boldsymbol{\mu}_{\beta_l^*}^{(t)}$ are $p \times 1$ block vectors in $\boldsymbol{\mu}_{\beta^*}^{(t)}$ and $\boldsymbol{\Sigma}_{\beta_{l,l'}^*}^{(t)}$ are $p \times p$ block matrices in $\boldsymbol{\Sigma}_{\beta^*}^{(t)}$ with the following relationship:

$$\boldsymbol{\mu}_{\beta^*}^{(t)} = ((\boldsymbol{\mu}_{\beta_1^*}^{(t)})^T, (\boldsymbol{\mu}_{\beta_2^*}^{(t)})^T)^T, \tag{33}$$

$$\boldsymbol{\Sigma}_{\beta^*}^{(t)} = \begin{pmatrix} \boldsymbol{\Sigma}_{\beta_{1,1}^*}^{(t)} & \boldsymbol{\Sigma}_{\beta_{1,2}^*}^{(t)} \\ \boldsymbol{\Sigma}_{\beta_{2,1}^*}^{(t)} & \boldsymbol{\Sigma}_{\beta_{2,2}^*}^{(t)} \end{pmatrix}. \tag{34}$$

Finally, in the reduction step, we map our parameters of interest from the expanded space to the original space and set the expansion parameters to their null values. Specifically, we have:

$$\boldsymbol{D}^{(t+1)} = \boldsymbol{\alpha}^{(t+1)} \tilde{\boldsymbol{D}}^{(t+1)} \boldsymbol{\alpha}^{(t+1)}, \tag{35}$$

$$\boldsymbol{\Sigma}^{(t+1)} = \tilde{\boldsymbol{\Sigma}}^{(t+1)}, \tag{36}$$

$$\alpha_1^{(t+1)} = \alpha_2^{(t+1)} = 1. \tag{37}$$

We iterate the E-step, M-step, and the reduction step of the EM algorithm until convergence, which is achieved when the difference of the observed-data log-likelihood from two consecutive iterations is below a certain threshold. The default threshold is set to be $1 \times 10^{-5}$ and the formula for the observed-data log-likelihood is given by:

$$\begin{aligned}\log L(\boldsymbol{\theta}; \boldsymbol{y}) =& \log \int f(\boldsymbol{y}|\boldsymbol{\beta}^*) f(\boldsymbol{\beta}^*) d\boldsymbol{\beta}^* \\ =& -\frac{n_1 + n_2}{2}\log 2\pi - \frac{1}{2}\log|\boldsymbol{\Sigma}| - \frac{p}{2}\log|\boldsymbol{D}| - \frac{1}{2}\boldsymbol{y}^T \boldsymbol{\Sigma}^{-1} \boldsymbol{y} \\ &+ \frac{1}{2}\log|\boldsymbol{\Sigma}_{\beta^*}| + \frac{1}{2}\boldsymbol{\mu}_{\beta^*}^T \boldsymbol{\Sigma}_{\beta^*}^{-1} \boldsymbol{\mu}_{\beta^*}, \end{aligned} \tag{38}$$

where

$$\boldsymbol{\Sigma}_{\beta^*} = \left[(\boldsymbol{X}^*)^T \boldsymbol{\Sigma}^{-1} \boldsymbol{X}^* + (\boldsymbol{D} \otimes \boldsymbol{I}_p)^{-1}\right]^{-1}, \tag{39}$$

$$\boldsymbol{\mu}_{\beta^*} = \boldsymbol{\Sigma}_{\beta^*} (\boldsymbol{X}^*)^T \boldsymbol{\Sigma}^{-1} \boldsymbol{y}. \tag{40}$$

We set the initial values of all the parameters, $\boldsymbol{\theta}^{(0)}$, as follows:

$$\rho^{(0)} = 0, \tag{41}$$

$$(\sigma_l^2)^{(0)} = 1, \tag{42}$$

$$(\sigma_{\beta_l}^2)^{(0)} = \arg\max_{\sigma_{\beta_l}^2 \in [0,1]} f(\boldsymbol{y}_l; \sigma_l^2 = 1). \tag{43}$$

In particular, the initial values of $\sigma_{\beta_l}^2$ are set to be the maximum likelihood estimates with respect to the equation (21) or (22), with all other parameters fixed at their initial values. We provide the log-likelihood as follows:

$$\begin{aligned}\log f(\boldsymbol{y}_l; \sigma_l^2 = 1, \alpha_l = 1) = &-\frac{n_l}{2}\log 2\pi - \frac{1}{2}\boldsymbol{y}_l^T\boldsymbol{y}_l - \frac{p}{2}\log\sigma_{\beta_l}^2 - \frac{1}{2}\log|(\boldsymbol{X}_l^*)^T\boldsymbol{X}_l^* + \sigma_{\beta_l}^{-2}\boldsymbol{I}_p| \\ &+ \frac{1}{2}\boldsymbol{y}_l^T\boldsymbol{X}_l^*\left[(\boldsymbol{X}_l^*)^T\boldsymbol{X}_l^* + \sigma_{\beta_l}^{-2}\boldsymbol{I}_p\right]^{-1}(\boldsymbol{X}_l^*)^T\boldsymbol{y}_l,\end{aligned} \tag{44}$$

with which we estimate $\sigma_{\beta_1}^2$ and $\sigma_{\beta_2}^2$ based on the Brent's method [3] implemented in the "optim" function in R.

##### 2.2.2 Estimation under $H_0 : \sigma_{\beta_2}^2 = \rho = 0$

In this scenario, we follow similar procedures as described in the section 2.2.1 to estimate all the nuisance parameters $\boldsymbol{\theta}_2 = (\sigma_{\beta_1}^2, \sigma_1^2, \sigma_2^2)$, with $\boldsymbol{\theta}_1 = (\sigma_{\beta_2}^2, \rho)$ fixed at their null values of 0. All the relevant quantities in the PX-EM algorithm under the null model:

$$\begin{aligned}\boldsymbol{y}_1 &= \alpha_1\boldsymbol{X}_1^*\boldsymbol{\beta}_1^* + \boldsymbol{\epsilon}_1, \\ \boldsymbol{y}_2 &= \boldsymbol{\epsilon}_2, \\ \boldsymbol{\epsilon} &\sim N(\mathbf{0}, \boldsymbol{\Sigma}), \\ \boldsymbol{\beta}_1^* &\sim N(\mathbf{0}, \sigma_{\beta_1}^2\boldsymbol{I}_p),\end{aligned} \tag{45}$$

are listed below:

1. (E-step) The complete-data log-likelihood function:

$$\begin{aligned}&\log L(\alpha_1, \tilde{\boldsymbol{\theta}}_2; \boldsymbol{y}, \boldsymbol{\beta}_1^*) \\ =& -\frac{n_1+n_2+p}{2}\log 2\pi - \frac{n_1}{2}\log\tilde{\sigma}_1^2 - \frac{n_2}{2}\log\tilde{\sigma}_2^2 - \frac{p}{2}\log\tilde{\sigma}_{\beta_1}^2 \\ &- \frac{1}{2}\tilde{\sigma}_1^{-2}\boldsymbol{y}_1^T\boldsymbol{y}_1 - \frac{1}{2}\tilde{\sigma}_2^{-2}\boldsymbol{y}_2^T\boldsymbol{y}_2 - \frac{1}{2}(\boldsymbol{\beta}_1^*)^T\left[\alpha_1^2\tilde{\sigma}_1^{-2}(\boldsymbol{X}_1^*)^T\boldsymbol{X}_1^* + \tilde{\sigma}_{\beta_1}^{-2}\boldsymbol{I}_p\right]\boldsymbol{\beta}_1^* \\ &+ \alpha_1\tilde{\sigma}_1^{-2}(\boldsymbol{\beta}_1^*)^T(\boldsymbol{X}_1^*)^T\boldsymbol{y}_1.\end{aligned} \tag{46}$$

2. (E-step) The conditional distribution of $\boldsymbol{\beta}_1^*$:

$$\boldsymbol{\beta}_1^*|\boldsymbol{y}_1, \boldsymbol{\theta}_2^{(t)} \sim N(\boldsymbol{\mu}_{\beta_1^*}^{(t)}, \boldsymbol{\Sigma}_{\beta_1^*}^{(t)}), \tag{47}$$

where

$$\boldsymbol{\Sigma}_{\beta_1^*}^{(t)} = \left[(\tilde{\sigma}_1^{(t)})^{-2}(\boldsymbol{X}_1^*)^T\boldsymbol{X}_1^* + (\tilde{\boldsymbol{\sigma}}_{\beta_1}^{(t)})^{-2}\boldsymbol{I}_p\right]^{-1}, \tag{48}$$

$$\boldsymbol{\mu}_{\beta_1^*}^{(t)} = (\tilde{\sigma}_1^{(t)})^{-2}\boldsymbol{\Sigma}_{\beta_1^*}^{(t)}(\boldsymbol{X}_1^*)^T\boldsymbol{y}_1. \tag{49}$$

3. (E-step) The $Q$ function:

$$\begin{aligned}
&Q(\tilde{\boldsymbol{\theta}}_2, \alpha_1|\boldsymbol{\theta}_2^{(t)}) \\
=& -\frac{n_1+n_2+p}{2}\log 2\pi - \frac{n_1}{2}\log\tilde{\sigma}_1^2 - \frac{n_2}{2}\log\tilde{\sigma}_2^2 - \frac{p}{2}\log\tilde{\sigma}_{\beta_1}^2 \\
&- \frac{1}{2}\tilde{\sigma}_1^{-2}\boldsymbol{y}_1^T\boldsymbol{y}_1 - \frac{1}{2}\tilde{\sigma}_2^{-2}\boldsymbol{y}_2^T\boldsymbol{y}_2 - \frac{1}{2}(\boldsymbol{\mu}_{\beta_1^*}^{(t)})^T\left[\alpha_1^2\tilde{\sigma}_1^{-2}(\boldsymbol{X}_1^*)^T\boldsymbol{X}_1^* + \tilde{\sigma}_{\beta_1}^{-2}\boldsymbol{I}_p\right]\boldsymbol{\mu}_{\beta_1^*}^{(t)} \\
&- \frac{1}{2}tr\left\{\boldsymbol{\Sigma}_{\beta_1^*}^{(t)}\left[\alpha_1^2\tilde{\sigma}_1^{-2}(\boldsymbol{X}_1^*)^T\boldsymbol{X}_1^* + \tilde{\sigma}_{\beta_1}^{-2}\boldsymbol{I}_p\right]\right\} \\
&+ \alpha_1\tilde{\sigma}_1^{-2}(\boldsymbol{\mu}_{\beta_1^*}^{(t)})^T(\boldsymbol{X}_1^*)^T\boldsymbol{y}_1.
\end{aligned} \tag{50}$$

4. (M-step) The update of $\tilde{\sigma}_{\beta_1}^2$:

$$(\tilde{\sigma}_{\beta_1}^2)^{(t+1)} = \frac{1}{p}\left[(\boldsymbol{\mu}_{\beta_1^*}^{(t)})^T\boldsymbol{\mu}_{\beta_1^*}^{(t)} + tr(\boldsymbol{\Sigma}_{\beta_1^*}^{(t)})\right]. \tag{51}$$

5. (M-step) The update of $\alpha_1$:

$$\alpha_1^{(t+1)} = \frac{(\boldsymbol{\mu}_{\beta_1^*}^{(t)})^T(\boldsymbol{X}_1^*)^T\boldsymbol{y}_1}{(\boldsymbol{\mu}_{\beta_1^*}^{(t)})^T(\boldsymbol{X}_1^*)^T\boldsymbol{X}_1^*\boldsymbol{\mu}_{\beta_1^*}^{(t)} + tr\left\{\boldsymbol{\Sigma}_{\beta_1^*}^{(t)}(\boldsymbol{X}_1^*)^T\boldsymbol{X}_1^*\right\}}. \tag{52}$$

6. (M-step) The update of $\tilde{\sigma}_1^2$:

$$\begin{aligned}
(\tilde{\sigma}_1^2)^{(t+1)} =& \frac{1}{n_1}\left(\boldsymbol{y}_1 - \alpha_1^{(t+1)}\boldsymbol{X}_1^*\boldsymbol{\mu}_{\beta_1^*}^{(t)}\right)^T\left(\boldsymbol{y}_1 - \alpha_1^{(t+1)}\boldsymbol{X}_1^*\boldsymbol{\mu}_{\beta_1^*}^{(t)}\right) \\
&+ \frac{1}{n_1}(\alpha_1^{(t+1)})^2 tr\left[(\boldsymbol{X}_1^*)^T\boldsymbol{X}_1^*\boldsymbol{\Sigma}_{\boldsymbol{\beta}_1^*}^{(t)}\right].
\end{aligned} \tag{53}$$

7. (M-step) The update of $\tilde{\sigma}_2^2$:

$$(\tilde{\sigma}_2^2)^{(t+1)} = \frac{1}{n_2}\boldsymbol{y}_2^T\boldsymbol{y}_2. \tag{54}$$

8. Reduction step:

$$(\sigma_{\beta_1}^2)^{(t+1)} = (\alpha_1^{(t+1)})^2(\tilde{\sigma}_{\beta_1}^2)^{(t+1)}, \tag{55}$$

$$(\sigma_1^2)^{(t+1)} = (\tilde{\sigma}_1^2)^{(t+1)}, \tag{56}$$

$$(\alpha_1)^{(t+1)} = 1. \tag{57}$$

9. Observed-data log-likelihood:

$$\begin{aligned}
\log L(\boldsymbol{\theta}_2; \boldsymbol{y}) &= \log \int f(\boldsymbol{y}|\boldsymbol{\beta}_1^*) f(\boldsymbol{\beta}_1^*) d\boldsymbol{\beta}_1^* \\
&= -\frac{n_1+n_2}{2}\log 2\pi - \frac{n_1}{2}\log\sigma_1^2 - \frac{n_2}{2}\log\sigma_2^2 - \frac{k}{2}\log\sigma_{\beta_1}^2 \\
&\quad - \frac{1}{2}\sigma_1^{-2}\boldsymbol{y}_1^T\boldsymbol{y}_1 - \frac{1}{2}\sigma_2^{-2}\boldsymbol{y}_2^T\boldsymbol{y}_2 + \frac{1}{2}\log|\boldsymbol{\Sigma}_{\beta_1^*}| + \frac{1}{2}\boldsymbol{\mu}_{\beta_1^*}^T\boldsymbol{\Sigma}_{\beta_1^*}^{-1}\boldsymbol{\mu}_{\beta_1^*},
\end{aligned} \tag{58}$$

where

$$\boldsymbol{\Sigma}_{\beta_1^*} = \left[\sigma_1^{-2}(\boldsymbol{X}_1^*)^T\boldsymbol{X}_1^* + \sigma_{\beta_1}^{-2}\boldsymbol{I}_p\right]^{-1}, \tag{59}$$

$$\boldsymbol{\mu}_{\beta_1^*} = \sigma_1^{-2}\boldsymbol{\Sigma}_{\beta_1^*}(\boldsymbol{X}_1^*)^T\boldsymbol{y}_1. \tag{60}$$

##### 2.2.3 Estimation under $H_0 : r = 0$

In this scenario, we estimate all the nuisance parameters $\boldsymbol{\theta}_3 = (\sigma_{\beta_1}^2, \sigma_{\beta_2}^2, \sigma_1^2, \sigma_2^2)$, with $\rho$ fixed at its null value of 0. All the relevant quantities in the PX-EM algorithm can be easily obtained by setting the off-diagonal elements in $\boldsymbol{D}$ to be zeros for equations (25)-(40).

#### 2.3 Extension toward summary statistics

VINTAGE can be easily extended to use only summary statistics from the gene expression mapping study and GWAS for inference. These summary statistics take the forms of marginal z-scores and a SNP-SNP correlation matrix. The SNP-SNP correlation matrix is commonly referred to as the LD matrix and can be estimated from a reference panel containing individuals with the same genetic ancestry as those in the gene expression study and GWAS. To address potential biases caused by an inaccurate estimation of the LD matrix, we focus on the top $k$ eigenvalues and eigenvectors for inference, where $k$ is determined in a way such that 99% of variance in the LD matrix is explained. Specifically, we work on the reduced design matrix $\tilde{\boldsymbol{X}}_l^* = \boldsymbol{X}_l\boldsymbol{Q}_{:,1:k}$ and its corresponding regression coefficients $\tilde{\boldsymbol{\beta}}_l^* = \boldsymbol{Q}_{:,1:k}^T\boldsymbol{\beta}_l$ under the transformed models for both the gene expression ($l = 1$) and the GWAS ($l = 2$) data. By simply replacing $\boldsymbol{X}_l^*$ with $\tilde{\boldsymbol{X}}_l^*$ in the equations presented in the section 2.2, we can estimate all the parameters. Subsequently, for extension towards summary statistics, we follow Zou et al. [4] and express the sufficient statistics in our model using summary statistics based on the following two approximations:

$$(\tilde{\boldsymbol{X}}_{l,j}^*)^T\boldsymbol{y}_l \approx \sqrt{n_l}\hat{\boldsymbol{Q}}_{:,1:k}^T\hat{z}_{l,j}\hat{\sigma}_{l,j}, \tag{61}$$

$$(\tilde{\boldsymbol{X}}_l^*)^T\tilde{\boldsymbol{X}}_l^* \approx n_l\hat{\boldsymbol{Q}}_{:,1:k}^T\left[(1-\lambda_l)\hat{\boldsymbol{R}}_0 + \lambda_l\boldsymbol{I}_p\right]\hat{\boldsymbol{Q}}_{:,1:k}, \tag{62}$$

where the subscript $j = 1, \ldots, k$ denotes the $j$th genotype PC; $\hat{z}_{l,j}$ is the marginal z-score obtained from the single-variant regression analysis; $\hat{\sigma}_{l,j}^2$ is the residual variance estimate obtained from the same single-variant regression analysis and can be calculated from $\hat{z}_{l,j}$ in the form of $\hat{\sigma}_{l,j}^2 = \frac{n_l-1}{n_l-2+\hat{z}_{l,j}^2}$;

$\hat{\boldsymbol{R}}_0$ is a $p \times p$ LD matrix estimated from a reference panel; and $\hat{\boldsymbol{Q}}_{:,1:k}$ is the $p \times k$ matrix of the top $k$ eigenvectors from the eigendecomposition of $\hat{\boldsymbol{R}}_0 = \hat{\boldsymbol{Q}}\hat{\boldsymbol{\Lambda}}_0\hat{\boldsymbol{Q}}^T$. Above, we have approximated $\boldsymbol{R}_l$ in a regularized form: $\hat{\boldsymbol{R}}_l = (1-\lambda_l)\hat{\boldsymbol{R}}_0 + \lambda_l \boldsymbol{I}_p$, where $\lambda_l \in [0,1]$ serves as the degree of regularization. Following Zou et al. [4], we estimate the regularization parameter $\lambda_l$ by maximizing the likelihood under the null ($\boldsymbol{\beta}_l = 0$), in the form of $\hat{\lambda}_l = \underset{\lambda\in[0,1]}{\arg\max}\, N(\tilde{\boldsymbol{z}}_l; \boldsymbol{0}, (1-\lambda)\hat{\boldsymbol{R}}_0 + \lambda \boldsymbol{I}_p)$, where $\tilde{\boldsymbol{z}}_l = \hat{\boldsymbol{z}}_l \hat{\boldsymbol{\sigma}}_l$ represents the PVE (Proportion of phenotypic Variance Explained)-adjusted marginal z-scores. Through regularization, we improve the consistency between the LD matrix and the PVE-adjusted z-scores. Lastly, the quantity in equation (62) can be further simplified to be $n_l(\hat{\boldsymbol{\Lambda}}_l)_{1:k} = n_l\left[(1-\lambda_l)(\hat{\boldsymbol{\Lambda}}_0)_{1:k} + \lambda_l \boldsymbol{I}_k\right]$, where $(\hat{\boldsymbol{\Lambda}}_0)_{1:k}$ is the $k \times k$ diagonal matrix of the top $k$ eigenvalues of $\hat{\boldsymbol{R}}_0$.

###### 2.3.1 Estimation under the full model

Given the above approximations, all the quantities that need to be evaluated in the PX-EM algorithm can be expressed with summary statistics in the following:

1. The conditional expectation and variance of $\tilde{\boldsymbol{\beta}}^*$:

$$\boldsymbol{\Sigma}_{\tilde{\beta}^*}^{(t)} = \left[\begin{pmatrix} n_1(\tilde{\sigma}_1^{-2})^{(t)}(\hat{\boldsymbol{\Lambda}}_1)_{1:k} & -\rho^{(t)}\boldsymbol{I}_k \\ -\rho^{(t)}\boldsymbol{I}_k & n_2(\tilde{\sigma}_2^{-2})^{(t)}(\hat{\boldsymbol{\Lambda}}_2)_{1:k} \end{pmatrix} + (\boldsymbol{D}^{(t)} \otimes \boldsymbol{I}_k)^{-1}\right]^{-1}$$
$$= \begin{pmatrix} n_1(\tilde{\sigma}_1^{-2})^{(t)}(\hat{\boldsymbol{\Lambda}}_1)_{1:k} + |\boldsymbol{D}^{(t)}|^{-1}(\sigma_{\beta_2}^2)^{(t)}\boldsymbol{I}_k & -|\boldsymbol{D}^{(t)}|^{-1}\rho^{(t)}\boldsymbol{I}_k \\ -|\boldsymbol{D}^{(t)}|^{-1}\rho^{(t)}\boldsymbol{I}_k & n_2(\tilde{\sigma}_2^{-2})^{(t)}(\hat{\boldsymbol{\Lambda}}_2)_{1:k} + |\boldsymbol{D}^{(t)}|^{-1}(\sigma_{\beta_1}^2)^{(t)}\boldsymbol{I}_k \end{pmatrix}^{-1}, \tag{63}$$

$$\boldsymbol{\mu}_{\tilde{\beta}^*}^{(t)} = \boldsymbol{\Sigma}_{\tilde{\beta}^*}^{(t)} \begin{pmatrix} \sqrt{n_1}(\tilde{\sigma}_1^{-2})^{(t)}\hat{\boldsymbol{Q}}_{:,1:k}^T\tilde{\boldsymbol{z}}_1 \\ \sqrt{n_2}(\tilde{\sigma}_2^{-2})^{(t)}\hat{\boldsymbol{Q}}_{:,1:k}^T\tilde{\boldsymbol{z}}_2 \end{pmatrix}, \tag{64}$$

where the inversion in equation (63) can be easily evaluated by recognizing that the four blocks in the matrix are all diagonal matrices. Specifically, we have

$$\boldsymbol{\Sigma}_{\tilde{\beta}^*}^{(t)} = \begin{pmatrix} (\boldsymbol{A}_1 - \boldsymbol{A}_2\boldsymbol{A}_4^{-1}\boldsymbol{A}_2)^{-1} & -(\boldsymbol{A}_1 - \boldsymbol{A}_2\boldsymbol{A}_4^{-1}\boldsymbol{A}_2)^{-1}\boldsymbol{A}_2\boldsymbol{A}_4^{-1} \\ -(\boldsymbol{A}_1 - \boldsymbol{A}_2\boldsymbol{A}_4^{-1}\boldsymbol{A}_2)^{-1}\boldsymbol{A}_2\boldsymbol{A}_4^{-1} & (\boldsymbol{A}_4 - \boldsymbol{A}_2\boldsymbol{A}_1^{-1}\boldsymbol{A}_2)^{-1} \end{pmatrix}, \tag{65}$$

where

$$\boldsymbol{A}_1 = n_1(\tilde{\sigma}_1^{-2})^{(t)}(\hat{\boldsymbol{\Lambda}}_1)_{1:k} + |\boldsymbol{D}^{(t)}|^{-1}(\sigma_{\beta_2}^2)^{(t)}\boldsymbol{I}_k, \tag{66}$$
$$\boldsymbol{A}_2 = -|\boldsymbol{D}^{(t)}|^{-1}\rho^{(t)}\boldsymbol{I}_k, \tag{67}$$
$$\boldsymbol{A}_4 = n_2(\tilde{\sigma}_2^{-2})^{(t)}(\hat{\boldsymbol{\Lambda}}_2)_{1:k} + |\boldsymbol{D}^{(t)}|^{-1}(\sigma_{\beta_1}^2)^{(t)}\boldsymbol{I}_k. \tag{68}$$

2. The update of $\tilde{\boldsymbol{D}}$:

$$\tilde{\boldsymbol{D}}^{(t+1)} = \frac{1}{k}\begin{pmatrix} (\boldsymbol{\mu}^{(t)}_{\tilde{\beta}^*_1})^T\boldsymbol{\mu}^{(t)}_{\tilde{\beta}^*_1} + tr(\boldsymbol{\Sigma}^{(t)}_{\tilde{\beta}^*_{1,1}}) & (\boldsymbol{\mu}^{(t)}_{\tilde{\beta}^*_1})^T\boldsymbol{\mu}^{(t)}_{\tilde{\beta}^*_2} + tr(\boldsymbol{\Sigma}^{(t)}_{\tilde{\beta}^*_{1,2}}) \\ (\boldsymbol{\mu}^{(t)}_{\tilde{\beta}^*_2})^T\boldsymbol{\mu}^{(t)}_{\tilde{\beta}^*_1} + tr(\boldsymbol{\Sigma}^{(t)}_{\tilde{\beta}^*_{2,1}}) & (\boldsymbol{\mu}^{(t)}_{\tilde{\beta}^*_2})^T\boldsymbol{\mu}^{(t)}_{\tilde{\beta}^*_2} + tr(\boldsymbol{\Sigma}^{(t)}_{\tilde{\beta}^*_{2,2}}) \end{pmatrix}. \tag{69}$$

3. The update of $\alpha_l$ for $l = 1$ and 2:

$$\alpha_l^{(t+1)} = \frac{(\boldsymbol{\mu}^{(t)}_{\tilde{\beta}^*_l})^T\hat{\boldsymbol{Q}}^T_{:,1:k}\tilde{\boldsymbol{z}}_l}{\sqrt{n_l}(\boldsymbol{\mu}^{(t)}_{\tilde{\beta}^*_l})^T(\hat{\boldsymbol{\Lambda}}_l)_{1:k}\boldsymbol{\mu}^{(t)}_{\tilde{\beta}^*_l} + \sqrt{n_l}tr\left\{\boldsymbol{\Sigma}^{(t)}_{\tilde{\beta}^*_{l,l}}(\hat{\boldsymbol{\Lambda}}_l)_{1:k}\right\}}. \tag{70}$$

4. The update of $\tilde{\sigma}_l^2$ for $l = 1$ and 2:

$$\begin{aligned}(\tilde{\sigma}_l^2)^{(t+1)} =& 1 + (\alpha_l^{(t+1)})^2(\boldsymbol{\mu}^{(t)}_{\tilde{\beta}^*_l})^T(\hat{\boldsymbol{\Lambda}}_l)_{1:k}\boldsymbol{\mu}^{(t)}_{\tilde{\beta}^*_l} - \frac{2}{\sqrt{n_l}}\alpha_l^{(t+1)}(\boldsymbol{\mu}^{(t)}_{\tilde{\beta}^*_l})^T\hat{\boldsymbol{Q}}^T_{:,1:k}\tilde{\boldsymbol{z}}_l \\ &+ (\alpha_l^{(t+1)})^2 tr\left\{\boldsymbol{\Sigma}^{(t)}_{\tilde{\beta}^*_{l,l}}(\hat{\boldsymbol{\Lambda}}_l)_{1:k}\right\}. \end{aligned} \tag{71}$$

5. Observed-data log-likelihood:

$$\begin{aligned}\log L(\boldsymbol{\theta};\boldsymbol{y}) =& -\frac{n_1+n_2}{2}\log 2\pi - \frac{n_1}{2}\log\sigma_1^2 - \frac{n_2}{2}\log\sigma_2^2 - \frac{p}{2}\log|\boldsymbol{D}| \\ &- \frac{n_1}{2}\sigma_1^{-2} - \frac{n_2}{2}\sigma_2^{-2} + \frac{1}{2}\log|\boldsymbol{\Sigma}_{\tilde{\beta}^*}| + \frac{1}{2}\boldsymbol{\mu}^T_{\tilde{\beta}^*}\boldsymbol{\Sigma}^{-1}_{\tilde{\beta}^*}\boldsymbol{\mu}_{\tilde{\beta}^*}, \end{aligned} \tag{72}$$

   where

$$\boldsymbol{\Sigma}_{\tilde{\beta}^*} = \begin{pmatrix} n_1\sigma_1^{-2}(\hat{\boldsymbol{\Lambda}}_1)_{1:k} + |\boldsymbol{D}|^{-1}\sigma^2_{\beta_2}\boldsymbol{I}_k & -|\boldsymbol{D}|^{-1}\rho\boldsymbol{I}_k \\ -|\boldsymbol{D}|^{-1}\rho\boldsymbol{I}_k & n_2\sigma_2^{-2}(\hat{\boldsymbol{\Lambda}}_2)_{1:k} + |\boldsymbol{D}|^{-1}\sigma^2_{\beta_1}\boldsymbol{I}_k \end{pmatrix}^{-1}, \tag{73}$$

$$\boldsymbol{\mu}_{\tilde{\beta}^*} = \boldsymbol{\Sigma}_{\tilde{\beta}^*}\begin{pmatrix} \sqrt{n_1}\sigma_1^{-2}\hat{\boldsymbol{Q}}^T_{:,1:k}\tilde{\boldsymbol{z}}_1 \\ \sqrt{n_2}\sigma_2^{-2}\hat{\boldsymbol{Q}}^T_{:,1:k}\tilde{\boldsymbol{z}}_2 \end{pmatrix}, \tag{74}$$

   which can be similarly evaluated following equations (65)-(68). In particular, the determinant $|\boldsymbol{\Sigma}_{\tilde{\beta}^*}|$ can be evaluated in linear time complexity with respect to $k$ based on the property of a block matrix.

6. In the initialization of $\sigma^2_{\beta_l}$, equation (44) can be expressed with summary statistics as:

$$\begin{aligned}\log f(\boldsymbol{y}_l;\sigma_l^2 = 1, \alpha_l = 1) =& -\frac{n_l}{2}\log 2\pi - \frac{n_l}{2} - \frac{k}{2}\log\sigma^2_{\beta_l} - \frac{1}{2}\log|n_l(\hat{\boldsymbol{\Lambda}}_l)_{1:k} + \sigma^{-2}_{\beta_l}\boldsymbol{I}_k| \\ &+ \frac{1}{2}n_l\tilde{\boldsymbol{z}}_l^T\hat{\boldsymbol{Q}}_{:,1:k}\left(n_l(\hat{\boldsymbol{\Lambda}}_l)_{1:k} + \sigma^{-2}_{\beta_l}\boldsymbol{I}_k\right)^{-1}\hat{\boldsymbol{Q}}^T_{:,1:k}\tilde{\boldsymbol{z}}_l, \end{aligned} \tag{75}$$

Above, $\boldsymbol{\mu}_{\tilde{\beta}_l^*}^{(t)}$ are $k \times 1$ block vectors in $\boldsymbol{\mu}_{\tilde{\beta}^*}^{(t)}$ and $\boldsymbol{\Sigma}_{\tilde{\beta}_{l,l'}^*}^{(t)}$ are $k \times k$ block matrices in $\boldsymbol{\Sigma}_{\tilde{\beta}^*}^{(t)}$ with the following relationship:

$$\boldsymbol{\mu}_{\tilde{\beta}^*}^{(t)} = ((\boldsymbol{\mu}_{\tilde{\beta}_1^*}^{(t)})^T, (\boldsymbol{\mu}_{\tilde{\beta}_2^*}^{(t)})^T)^T, \tag{76}$$

$$\boldsymbol{\Sigma}_{\tilde{\beta}^*}^{(t)} = \begin{pmatrix} \boldsymbol{\Sigma}_{\tilde{\beta}_{1,1}^*}^{(t)} & \boldsymbol{\Sigma}_{\tilde{\beta}_{1,2}^*}^{(t)} \\ \boldsymbol{\Sigma}_{\tilde{\beta}_{2,1}^*}^{(t)} & \boldsymbol{\Sigma}_{\tilde{\beta}_{2,2}^*}^{(t)} \end{pmatrix}. \tag{77}$$

Finally, we provide a summary of the VINTAGE algorithm for parameter estimation under the full model using summary statistics:

**Algorithm 1** VINTAGE Algorithm

```
Initialize:
    θ ← θ^(0)                                              Eq.(41)-(44),(75)
for t = 0 to T do:
    E-step: update Σ^(t)_{β̃*} and μ^(t)_{β̃*}              Eq.(63)-(64)
    M-step: update D̃^(t+1), α_l^(t+1), and (σ̃_l^2)^(t+1)  Eq.(69)-(71)
    Reduction step: transform θ̃^(t+1) to θ^(t+1)           Eq.(35)-(37)
    Evaluate log L(θ; y)^(t+1)                              Eq.(72)
    if |log L(θ; y)^(t+1) − log L(θ; y)^(t)| < tol then
        break
    end if
end for
```

##### 2.3.2 Estimation under $H_0 : \sigma^2_{\beta_2} = \rho = 0$

In this scenario, all the quantities that need to be evaluated in the section (2.2.2) can be similarly expressed with summary statistics in the following:

1. The conditional expectation and variance of $\tilde{\boldsymbol{\beta}}_1^*$:

$$\boldsymbol{\Sigma}_{\tilde{\beta}_1^*}^{(t)} = \left[ n_1 (\tilde{\sigma}_1^{(t)})^{-2} (\hat{\boldsymbol{\Lambda}}_1)_{1:k} + (\tilde{\boldsymbol{\sigma}}_{\beta_1}^{(t)})^{-2} \boldsymbol{I}_k \right]^{-1}, \tag{78}$$

$$\boldsymbol{\mu}_{\tilde{\beta}_1^*}^{(t)} = \sqrt{n_1} (\tilde{\sigma}_1^{(t)})^{-2} \boldsymbol{\Sigma}_{\tilde{\beta}_1^*}^{(t)} \hat{\boldsymbol{Q}}_{:,1:k}^T \tilde{\boldsymbol{z}}_1. \tag{79}$$

2. The update of $\alpha_1$:

$$\alpha_1^{(t+1)} = \frac{(\boldsymbol{\mu}_{\tilde{\beta}_1^*}^{(t)})^T \hat{\boldsymbol{Q}}_{:,1:k}^T \tilde{\boldsymbol{z}}_1}{\sqrt{n_1} (\boldsymbol{\mu}_{\tilde{\beta}_1^*}^{(t)})^T (\hat{\boldsymbol{\Lambda}}_1)_{1:k} \boldsymbol{\mu}_{\tilde{\beta}_1^*}^{(t)} + \sqrt{n_1} tr \left\{ \boldsymbol{\Sigma}_{\tilde{\beta}_1^*}^{(t)} (\hat{\boldsymbol{\Lambda}}_1)_{1:k} \right\}}. \tag{80}$$

3. The update of $\tilde{\sigma}_1^2$ :

$$(\tilde{\sigma}_1^2)^{(t+1)} =1 + (\alpha_1^{(t+1)})^2(\boldsymbol{\mu}_{\tilde{\beta}_1^*}^{(t)})^T(\hat{\boldsymbol{\Lambda}}_1)_{1:k}\boldsymbol{\mu}_{\tilde{\beta}_1^*}^{(t)} - \frac{2}{\sqrt{n_1}}\alpha_1^{(t+1)}(\boldsymbol{\mu}_{\tilde{\beta}_1^*}^{(t)})^T\hat{\boldsymbol{Q}}_{:,1:k}^T\tilde{\boldsymbol{z}}_1 + (\alpha_1^{(t+1)})^2 tr\left\{\boldsymbol{\Sigma}_{\tilde{\beta}_1^*}^{(t)}(\hat{\boldsymbol{\Lambda}}_1)_{1:k}\right\}. \tag{81}$$

4. The update of $\tilde{\sigma}_2^2$ :

$$(\tilde{\sigma}_2^2)^{(t+1)} = 1. \tag{82}$$

5. Observed-data log-likelihood:

$$\log L(\boldsymbol{\theta}_2;\boldsymbol{y}) = -\frac{n_1+n_2}{2}\log 2\pi - \frac{n_1}{2}\log\sigma_1^2 - \frac{n_2}{2}\log\sigma_2^2 - \frac{k}{2}\log\sigma_{\beta_1}^2 - \frac{n_1}{2}\sigma_1^{-2} - \frac{n_2}{2}\sigma_2^{-2} + \frac{1}{2}\log|\boldsymbol{\Sigma}_{\tilde{\beta}_1^*}| + \frac{1}{2}\boldsymbol{\mu}_{\tilde{\beta}_1^*}^T\boldsymbol{\Sigma}_{\tilde{\beta}_1^*}^{-1}\boldsymbol{\mu}_{\tilde{\beta}_1^*}, \tag{83}$$

where

$$\boldsymbol{\Sigma}_{\tilde{\beta}_1^*} = \left(n_1\sigma_1^{-2}(\hat{\boldsymbol{\Lambda}}_1)_{1:k} + \sigma_{\beta_1}^{-2}\boldsymbol{I}_k\right)^{-1}, \tag{84}$$

$$\boldsymbol{\mu}_{\tilde{\beta}_1^*} = \sqrt{n_1}\sigma_1^{-2}\boldsymbol{\Sigma}_{\tilde{\beta}_1^*}\hat{\boldsymbol{Q}}_{:,1:k}^T\tilde{\boldsymbol{z}}_1. \tag{85}$$

##### 2.3.3 Estimation under $H_0 : r = 0$

In this scenario, all the quantities that need to be evaluated in the section (2.2.3) can be similarly expressed with summary statistics by setting the off-diagonal elements in $\boldsymbol{D}$ to be zeros.

#### 3 Gene-wise genetic variance test

We have developed a gene-wise genetic variance test to identify gene associations with the trait. Such association reflects the total effect of SNPs in the genic and adjacent regulatory regions (i.e., cis-SNPs) of a gene on the trait. The total effect may be mediated by various molecular mechanisms such as gene expression regulation, protein regulatory changes, and splicing. The test assesses the null hypothesis $H_0 : \boldsymbol{\beta}_2 = 0$, which represents the situation where none of the cis-SNPs have any effect on the trait. Such null hypothesis is equivalent to:

$$H_0 : \sigma_{\beta_2}^2 = \rho = 0. \tag{86}$$

The alternative hypothesis that corresponds to the null consists of three scenarios depending on the specific value of the local genetic correlation, $r$:

$$H_1 : \sigma^2_{\beta_2} \neq 0 \text{ and } \begin{cases} r = 0 & \textit{(I; SKAT)} \\ r = \pm 1 & \textit{(II; TWAS)}, \\ others & \textit{(III)} \end{cases} \tag{87}$$

where scenario I aligns with the SKAT modeling assumption that there is no local genetic correlation between the gene expression and the trait; scenario II aligns with the TWAS modeling assumption that the genetic effects on gene expression and the trait are perfectly correlated; and scenario III is an intermediate scenario.

We have developed an optimal and unified variance component score test to carry out the gene-wise genetic variance test. The test is optimal in the sense that it maximizes power within the class of tests that is a linear combination of SKAT and TWAS statistics and achieves robust performance to a wide range of local genetic correlation values between the gene expression and the trait. The test is unified as it includes the SKAT and TWAS test statistics as two special cases.

To derive the test statistic, we first write the log-likelihood function with respect to the equation (8):

$$l(\boldsymbol{\theta}) = -\frac{n_1 + n_2}{2} \log 2\pi - \frac{1}{2} \log |\boldsymbol{\Sigma}_y| - \frac{1}{2} \boldsymbol{y}^T \boldsymbol{\Sigma}_y^{-1} \boldsymbol{y}, \tag{88}$$

where $\boldsymbol{\Sigma}_y = \boldsymbol{X}(\boldsymbol{D} \otimes \boldsymbol{I}_p)\boldsymbol{X}^T + \boldsymbol{\Sigma}$ is the marginal variance of $\boldsymbol{y}$. Then, we obtain the scores for the variance component $\sigma^2_{\beta_2}$ and the covariance component $\rho$ under the null hypothesis:

$$U_{\theta_{1,j}}(\boldsymbol{\theta}_2) = \frac{\partial l(\boldsymbol{\theta})}{\partial \theta_{1,j}}\Big|_{\boldsymbol{\theta}_1 = \boldsymbol{0}} = \left[ -\frac{1}{2} tr \left( \boldsymbol{\Sigma}_y^{-1} \frac{\partial \boldsymbol{\Sigma}_y}{\partial \theta_{1,j}} \right) + \frac{1}{2} \boldsymbol{y}^T \boldsymbol{\Sigma}_y^{-1} \frac{\partial \boldsymbol{\Sigma}_y}{\partial \theta_{1,j}} \boldsymbol{\Sigma}_y^{-1} \boldsymbol{y} \right]\Big|_{\boldsymbol{\theta}_1 = \boldsymbol{0}}, \tag{89}$$

where $\boldsymbol{\theta}_1 = (\theta_{1,1}, \theta_{1,2}) = (\sigma^2_{\beta_2}, \rho)$, $\boldsymbol{\theta}_2 = (\sigma^2_{\beta_1}, \sigma^2_1, \sigma^2_2)$ are nuisance parameters, and

$$\boldsymbol{\Sigma}_y^{-1}\Big|_{\boldsymbol{\theta}_1 = \boldsymbol{0}} = \begin{pmatrix} (\sigma^2_{\beta_1} \boldsymbol{X}_1 \boldsymbol{X}_1^T + \sigma_1^2 \boldsymbol{I}_{n_1})^{-1} & \boldsymbol{0}_{n_1,n_2} \\ \boldsymbol{0}_{n_2,n_1} & \sigma_2^{-2} \boldsymbol{I}_{n_2} \end{pmatrix}, \tag{90}$$

$$\frac{\partial \boldsymbol{\Sigma}_y}{\partial \theta_{1,1}} = \frac{\partial \boldsymbol{\Sigma}_y}{\partial \sigma^2_{\beta_2}} = \begin{pmatrix} \boldsymbol{0}_{n_1,n_1} & \boldsymbol{0}_{n_1,n_2} \\ \boldsymbol{0}_{n_2,n_1} & \boldsymbol{X}_2 \boldsymbol{X}_2^2 \end{pmatrix}, \tag{91}$$

$$\frac{\partial \boldsymbol{\Sigma}_y}{\partial \theta_{1,2}} = \frac{\partial \boldsymbol{\Sigma}_y}{\partial \rho} = \begin{pmatrix} \boldsymbol{0}_{n_1,n_1} & \boldsymbol{X}_1 \boldsymbol{X}_2^T \\ \boldsymbol{X}_2 \boldsymbol{X}_1^T & \boldsymbol{0}_{n_2,n_2} \end{pmatrix}. \tag{92}$$

Hence, the two scores take the form:

$$U_{\sigma^2_{\beta_2}}(\boldsymbol{\theta}_2) = \frac{1}{2} \sigma_2^{-4} \boldsymbol{y}_2^T \boldsymbol{X}_2 \boldsymbol{X}_2^T \boldsymbol{y}_2 \left[ -\frac{1}{2} \sigma_2^{-2} tr(\boldsymbol{X}_2^T \boldsymbol{X}_2) \right], \tag{93}$$

$$U_\rho(\boldsymbol{\theta}_2) = \sigma_2^{-2} \boldsymbol{y}_1^T (\sigma^2_{\beta_1} \boldsymbol{X}_1 \boldsymbol{X}_1^T + \sigma_1^2 \boldsymbol{I}_{n_1})^{-1} \boldsymbol{X}_1 \boldsymbol{X}_2^T \boldsymbol{y}_2. \tag{94}$$

Note that the second term of $U_{\sigma^2_{\beta_2}}(\boldsymbol{\theta}_2)$ (i.e., $-\frac{1}{2}\sigma_2^{-2}tr(\boldsymbol{X}_2^T\boldsymbol{X}_2)$) can be omitted because it is not random.

The above two scores serve as backbone for constructing our test statistic. Importantly, these two scores are closely related to the SKAT and TWAS test statistics (detailed proofs are provided in the sections 3.2 and 3.3):

**Remark 1** *(Connection to SKAT) $U_{\sigma^2_{\beta_2}}(\boldsymbol{\theta}_2)$, the score for the variance component $\sigma^2_{\beta_2}$, is equivalent to the SKAT statistic with an unweighted and linear kernel function under the null hypothesis.*

**Remark 2** *(Connection to TWAS) $U_\rho(\boldsymbol{\theta}_2)$, the score for the covariance component $\rho$, is equivalent to the two-stage TWAS test statistic under the null hypothesis, when the following two conditions are satisfied: (1) A BLUP prediction model is used in the expression study to obtain the SNP weights on gene expression; (2) the uncertainty associated with the SNP weights on gene expression is taken into account when constructing the TWAS test statistic.*

Because of the close relationship of the two scores to the SKAT and TWAS test statistics and because SKAT and TWAS represent two extreme cases of the alternative hypothesis in our modeling framework, we propose the final test statistic in the form of a weighted combination of the two scores as follows:

$$T_w = (1-w)\frac{U_{\sigma^2_{\beta_2}}}{\sqrt{\tilde{I}_{1,1}}} + w\frac{|U_\rho|}{\sqrt{\tilde{I}_{2,2}}}, \tag{95}$$

where weight $w$ is a scalar that controls the relative contribution of the two scores; the absolute value of $U_\rho$ keeps the sign of the second term consistent with that of the first term, so that their large positive values both reflect a deviation from the null; and $\tilde{I}_{1,1}$ and $\tilde{I}_{2,2}$ are corresponding elements of $\sigma^2_{\beta_2}$ and $\rho$ in the information matrix that ensure the two terms are comparable in scale.

Above, the information matrix of $\boldsymbol{\theta}_1 = (\sigma^2_{\beta_2}, \rho)$ takes the following form:

$$\tilde{\boldsymbol{I}} = \boldsymbol{I}_{\boldsymbol{\theta}_1\boldsymbol{\theta}_1} - \boldsymbol{I}_{\boldsymbol{\theta}_1\boldsymbol{\theta}_2}\boldsymbol{I}^{-1}_{\boldsymbol{\theta}_2\boldsymbol{\theta}_2}\boldsymbol{I}^T_{\boldsymbol{\theta}_1\boldsymbol{\theta}_2}, \tag{96}$$

where

$$\boldsymbol{I}_{\boldsymbol{\theta}_1\boldsymbol{\theta}_1} = -\mathbb{E}\left[\frac{\partial^2 l(\boldsymbol{\theta})}{\partial\boldsymbol{\theta}_1\boldsymbol{\theta}_1^T}\right], \boldsymbol{I}_{\boldsymbol{\theta}_1\boldsymbol{\theta}_2} = -\mathbb{E}\left[\frac{\partial^2 l(\boldsymbol{\theta})}{\partial\boldsymbol{\theta}_1\boldsymbol{\theta}_2^T}\right], \boldsymbol{I}_{\boldsymbol{\theta}_2\boldsymbol{\theta}_2} = -\mathbb{E}\left[\frac{\partial^2 l(\boldsymbol{\theta})}{\partial\boldsymbol{\theta}_2\boldsymbol{\theta}_2^T}\right], \tag{97}$$

are evaluated under $H_0$ with $\boldsymbol{\theta}_1 = \mathbf{0}$. After some algebra, we obtain:

$$\boldsymbol{I}_{\boldsymbol{\theta}_1\boldsymbol{\theta}_1} = \begin{pmatrix} \frac{1}{2}\sigma_2^{-4}tr[(\boldsymbol{X}_2^T\boldsymbol{X}_2)^2] & 0 \\ 0 & \sigma_2^{-2}tr[\boldsymbol{X}_2\boldsymbol{X}_1^T\boldsymbol{\Sigma}_{y_1}^{-1}\boldsymbol{X}_1\boldsymbol{X}_2^T] \end{pmatrix}, \tag{98}$$

$$\boldsymbol{I}_{\boldsymbol{\theta}_2\boldsymbol{\theta}_2} = \frac{1}{2}\begin{pmatrix} tr\{[\boldsymbol{X}_1^T\boldsymbol{\Sigma}_{y_1}^{-1}\boldsymbol{X}_1]^2\} & tr[\boldsymbol{X}_1^T\boldsymbol{\Sigma}_{y_1}^{-2}\boldsymbol{X}_1] & 0 \\ tr[\boldsymbol{X}_1^T\boldsymbol{\Sigma}_{y_1}^{-2}\boldsymbol{X}_1] & tr[\boldsymbol{\Sigma}_{y_1}^{-2}] & 0 \\ 0 & 0 & n_2\sigma_2^{-4} \end{pmatrix}, \tag{99}$$

$$\boldsymbol{I}_{\boldsymbol{\theta}_1\boldsymbol{\theta}_2} = \begin{pmatrix} 0 & 0 & \frac{1}{2}\sigma_2^{-4} tr(\boldsymbol{X}_2^T \boldsymbol{X}_2) \\ 0 & 0 & 0 \end{pmatrix}, \tag{100}$$

where $\boldsymbol{\Sigma}_{y_1} = \sigma_{\beta_1}^2 \boldsymbol{X}_1 \boldsymbol{X}_1^T + \sigma_1^2 \boldsymbol{I}_{n_1}$ is the marginal variance of $\boldsymbol{y}_1$. Hence,

$$\tilde{\boldsymbol{I}} = \begin{pmatrix} \frac{1}{2}\sigma_2^{-4} tr[(\boldsymbol{X}_2^T \boldsymbol{X}_2)^2] - \frac{1}{2n_2}\sigma_2^{-4} tr^2(\boldsymbol{X}_2^T \boldsymbol{X}_2) & 0 \\ 0 & \sigma_2^{-2} tr(\boldsymbol{X}_2 \boldsymbol{X}_1^T \boldsymbol{\Sigma}_{y_1}^{-1} \boldsymbol{X}_1 \boldsymbol{X}_2^T) \end{pmatrix}, \tag{101}$$

and

$$\tilde{I}_{1,1} = \frac{1}{2}\sigma_2^{-4} tr[(\boldsymbol{X}_2^T \boldsymbol{X}_2)^2] - \frac{1}{2n_2}\sigma_2^{-4} tr^2(\boldsymbol{X}_2^T \boldsymbol{X}_2), \tag{102}$$

$$\tilde{I}_{2,2} = \sigma_2^{-2} tr(\boldsymbol{X}_2 \boldsymbol{X}_1^T \boldsymbol{\Sigma}_{y_1}^{-1} \boldsymbol{X}_1 \boldsymbol{X}_2^T). \tag{103}$$

The test statistic $T_w$ reduces to the SKAT statistic when $w = 0$ and is expected to achieve high power when $r = 0$. Conversely, $T_w$ reduces to the TWAS statistic when $w = 1$ and is expected to achieve high power when $r = \pm 1$. For values of $w \in (0, 1)$, $T_w$ is expected to achieve high power when $r \in (-1, 0) \cup (0, 1)$. In practice, since the true $r$ is unknown, we conduct grid search on a set of $K$ pre-specified weight values ranging from 0 to 1 with equal increments ($K = 11$ by default). For each weight value, we compute the corresponding test statistic $T_w$ and determine the p value $p_w$ through a simulation-based approach (details in the section 3.1). We then use the Cauchy combination rule [5, 6] to combine the p values obtained from different weights into a single p value, denoted as $p_g$. In the last step, the combined Cauchy test statistic is given by:

$$T_g = \frac{1}{K} \sum_{k=1}^{K} \tan\{(0.5 - p_{w_k}\pi)\}, \tag{104}$$

where $p_{w_k}$ is the p value that corresponds to $T_{w_k}$ and tan denotes the tangent function. The final p value, $p_g$, is then derived based on the Cauchy distribution.

Finally, the scores and information can be similarly expressed with summary statistics under the transformed models in the following:

$$U_{\sigma_{\beta_2}^2}(\boldsymbol{\theta}_2) = \frac{1}{2}\sigma_2^{-4} n_2 \tilde{\boldsymbol{z}}_2^T \hat{\boldsymbol{Q}}_{:,1:k} \hat{\boldsymbol{Q}}_{:,1:k}^T \tilde{\boldsymbol{z}}_2, \tag{105}$$

$$U_{\rho}(\boldsymbol{\theta}_2) = \sigma_1^{-2}\sigma_2^{-2}\sigma_{\beta_1}^{-2}\sqrt{n_1 n_2}\, \tilde{\boldsymbol{z}}_1^T \hat{\boldsymbol{Q}}_{:,1:k} \boldsymbol{\Sigma}_{\tilde{\beta}_1^*} \hat{\boldsymbol{Q}}_{:,1:k}^T \tilde{\boldsymbol{z}}_2, \tag{106}$$

$$\tilde{I}_{1,1} = \frac{1}{2}\sigma_2^{-4} n_2^2 tr[(\hat{\boldsymbol{\Lambda}}_2^2)_{1:k}] - \frac{1}{2} n_2 \sigma_2^{-4} k^2, \tag{107}$$

$$\tilde{I}_{2,2} = \sigma_1^{-2}\sigma_2^{-2}\sigma_{\beta_1}^{-2} n_1 n_2 tr\left[\boldsymbol{\Sigma}_{\tilde{\beta}_1^*} (\hat{\boldsymbol{\Lambda}}_1)_{1:k} (\hat{\boldsymbol{\Lambda}}_2)_{1:k}\right], \tag{108}$$

where $\boldsymbol{\Sigma}_{\tilde{\beta}_1^*} = [n_1 \sigma_1^{-2} (\hat{\boldsymbol{\Lambda}}_1)_{1:k} + \sigma_{\beta_1}^{-2} \boldsymbol{I}_k]^{-1}$ is the conditional variance of $\tilde{\boldsymbol{\beta}}_1^*$.

##### 3.1 Simulation-based testing

We have developed a highly scalable simulation-based approach to simulate the test statistics under the null, with which we are able to obtain calibrated p value, $p_w$, from $T_w$. Specifically, we simulate $\tilde{\boldsymbol{z}}_2^{(b)}$ from its null distribution $N(\mathbf{0}, \hat{\boldsymbol{R}}_2)$ $B$ times ($B = 10^6$ by default) and construct the null test statistic $T_w^{(b)}$ with $U_{\sigma^2_{\beta_2}}^{(b)}(\boldsymbol{\theta}_2)$ and $U_\rho^{(b)}(\boldsymbol{\theta}_2)$ in the following form:

$$U_{\sigma^2_{\beta_2}}^{(b)}(\boldsymbol{\theta}_2) = \frac{1}{2}\sigma_2^{-4} n_2 (\tilde{\boldsymbol{z}}_2^{(b)})^T \tilde{\boldsymbol{z}}_2^{(b)}, \tag{109}$$

$$U_\rho^{(b)}(\boldsymbol{\theta}_2) = \sigma_1^{-2}\sigma_2^{-2}\sigma_{\beta_1}^{-2}\sqrt{n_1 n_2}\tilde{\boldsymbol{z}}_1^T \hat{\boldsymbol{Q}}_{:,1:k}\boldsymbol{\Sigma}_{\beta_1^*}\hat{\boldsymbol{Q}}_{:,1:k}^T \tilde{\boldsymbol{z}}_2^{(b)}, \tag{110}$$

where the superscript $b$ denotes the $b$th simulation replicate. With the simulated data, we obtain $p_w$ based on the definition of the p value in the following form:

$$p_w = \frac{1}{B}\sum_{b=1}^{B} I(T_w^{(b)} > T_w), \tag{111}$$

where $I(\cdot)$ is an indicator function that equals one if the expression is true and equals zero otherwise. The simulation-based testing approach is equivalent to a permutation test that breaks the association between the SNPs and trait in GWAS. Importantly, the estimates of the nuisance parameters, $\sigma_1^2$, $\sigma_2^2$, and $\sigma_{\beta_1}^2$, remain unchanged in the permuted dataset. This is because the estimation of these three parameters does not rely on the GWAS data under the null. This way, we do not need to re-estimate the nuisance parameters with each simulation replicate, which makes the procedure computationally efficient.

##### 3.2 Establishing the connection to SKAT

In this section, we provide a proof for *Remark* 1 based on the notations introduced in the section 1.1. We further denote $\boldsymbol{Z}$ as the $n_2 \times m$ matrix of covariates for $n_2$ individuals and $m$ covariates. SKAT [7] models gene variants, a continuous trait, and covariates for adjustment with the following equation:

$$\boldsymbol{y}_2 = \gamma_0 \mathbf{1}_{n_2} + \boldsymbol{Z}\boldsymbol{\gamma} + \boldsymbol{X}_2\boldsymbol{\beta}_2 + \boldsymbol{\epsilon}_2, \tag{112}$$

where $\gamma_0$ is an intercept term and $\boldsymbol{\gamma}$ is a $m$-vector of covariate effects. SKAT assumes that the genetic effect of each variant follows an arbitrary distribution with a mean of zero and a variance of $w_j\tau$, that is:

$$\beta_{2,j} \sim F(0, w_j\tau), \tag{113}$$

where $w_j$ is a pre-specified weight for variant $j$ and $\tau$ is a variance component. SKAT evaluates whether the gene variants influence the trait by testing the null hypothesis: $H_0 : \boldsymbol{\beta}_2 = \mathbf{0}$, which is equivalent to the null hypothesis $H_0 : \tau = 0$. The SKAT statistic takes the form:

$$T_{\text{skat}} = (\boldsymbol{y}_2 - \hat{\boldsymbol{\mu}})^T \boldsymbol{K}(\boldsymbol{y}_2 - \hat{\boldsymbol{\mu}}), \tag{114}$$

where $\hat{\boldsymbol{\mu}} = \hat{\gamma}_0 + \boldsymbol{Z}\boldsymbol{\gamma}$ is the predicted mean of $\boldsymbol{y}_2$ under the null and $\boldsymbol{K}$ is a $n_2 \times n_2$ kernel matrix with each element measuring the genetic similarity between a pair of individuals. Several kernels have been proposed that include the linear kernel, quadratic kernel, and the identity-by-state kernel to capture distinct genetic architectures [7]. Among these kernels, the linear kernel is the most widely used [8, 9] that takes the following form:

$$\boldsymbol{K} = \boldsymbol{X}_2 \boldsymbol{W} \boldsymbol{X}_2^T, \tag{115}$$

where $\boldsymbol{W} = diag(w_1, \ldots, w_p)$ is a diagonal matrix with $w_1, \ldots, w_p$ on the diagonal and zeros elsewhere. If we further assume that the kernel is unweighted (i.e., $w_j = 0$) and consider that $\boldsymbol{y}_2$ has already been sufficiently adjusted for covariates in the GWAS (i.e., $\hat{\boldsymbol{\mu}} = \boldsymbol{0}$), then we have

$$T_{\text{skat}} = (\boldsymbol{y}_2 - \hat{\boldsymbol{\mu}})^T \boldsymbol{K}(\boldsymbol{y}_2 - \hat{\boldsymbol{\mu}}) = \boldsymbol{y}_2^T \boldsymbol{X}_2 \boldsymbol{X}_2 \boldsymbol{y}_2,$$

which is equivalent to $U_{\sigma^2_{\beta_2}}$ in the role of a test statistic.

##### 3.3 Establishing the connection to TWAS

In this section, we provide a proof for *Remark* 2 based on the notations introduced in the section 1.1. Suppose we follow a two-stage model for TWAS analysis [10, 11]. In the first stage, we build the following linear mixed model that uses SNPs to predict gene expression in the gene expression mapping study:

$$\begin{aligned} \boldsymbol{y}_1 &= \boldsymbol{X}_1 \boldsymbol{\beta}_1 + \boldsymbol{\epsilon}_1, \\ \boldsymbol{\beta}_1 &\sim N(\boldsymbol{0}, \sigma^2_{\beta_1} \boldsymbol{I}_p), \\ \boldsymbol{\epsilon}_1 &\sim N(\boldsymbol{0}, \sigma^2_1 \boldsymbol{I}_{n_1}), \end{aligned} \tag{116}$$

with which we obtain the best linear unbiased prediction (BLUP) of $\boldsymbol{\beta}_1$ as gene expression prediction weights and carry these weights forward into the second stage. The BLUP of $\boldsymbol{\beta}_1$ takes the following form:

$$\hat{\boldsymbol{\beta}}_1 = (\sigma_1^{-2} \boldsymbol{X}_1^T \boldsymbol{X}_1 + \sigma^{-2}_{\beta_1} \boldsymbol{I}_p)^{-1} \sigma_1^{-2} \boldsymbol{X}_1^T \boldsymbol{y}_1. \tag{117}$$

In the second stage, we make use of the estimated prediction weights to construct the genetically regulated expression (GReX) in the GWAS and test the association of the GReX with the trait of interest. Specifically, we rely on the following model in the second stage:

$$\begin{aligned} \boldsymbol{y}_2 &= \alpha(\boldsymbol{X}_2\hat{\boldsymbol{\beta}}_1) + \boldsymbol{\epsilon}_2, \\ \boldsymbol{\epsilon}_2 &\sim N(\mathbf{0}, \sigma_2^2\boldsymbol{I}_{n_2}), \end{aligned} \tag{118}$$

where $\boldsymbol{X}_2\hat{\boldsymbol{\beta}}_1$ is the GReX in the GWAS and $\alpha$ is the effect of the GReX on the trait. TWAS evaluates the null hypothesis $H_0 : \alpha = 0$ with a test statistic that takes the following form:

$$T_{\text{twas}} = \frac{\hat{\boldsymbol{\beta}}_1^T\tilde{\boldsymbol{z}}_2}{\sqrt{\hat{\boldsymbol{\beta}}_1^T\hat{\boldsymbol{R}}_2\hat{\boldsymbol{\beta}}_1}}. \tag{119}$$

The connection of the covariance component score, $U_\rho$, with $T_{\text{twas}}$ can be recognized in view of the following equality:

$$\begin{aligned} U_\rho(\boldsymbol{\theta}_2) &= \sigma_2^{-2}\boldsymbol{y}_1^T(\sigma_{\beta_1}^2\boldsymbol{X}_1\boldsymbol{X}_1^T + \sigma_1^2\boldsymbol{I}_{n_1})^{-1}\boldsymbol{X}_1\boldsymbol{X}_2^T\boldsymbol{y}_2 \\ &= \sigma_2^{-2}\sigma_1^{-2}\sigma_{\beta_1}^{-2}\boldsymbol{y}_1^T\boldsymbol{X}_1(\sigma_1^{-2}\boldsymbol{X}_1^T\boldsymbol{X}_1 + \sigma_{\beta_1}^{-2}\boldsymbol{I}_p)^{-1}\boldsymbol{X}_2^T\boldsymbol{y}_2, \\ &= \sqrt{n_2}\sigma_2^{-2}\sigma_{\beta_1}^{-2}\left[(\sigma_1^{-2}\boldsymbol{X}_1^T\boldsymbol{X}_1 + \sigma_{\beta_1}^{-2}\boldsymbol{I}_p)^{-1}\sigma_1^{-2}\boldsymbol{X}_1^T\boldsymbol{y}_1\right]^T\frac{1}{\sqrt{n_2}}\boldsymbol{X}_2^T\boldsymbol{y}_2 \\ &= \sqrt{n_2}\sigma_2^{-2}\sigma_{\beta_1}^{-2}\hat{\boldsymbol{\beta}}_1^T\tilde{\boldsymbol{z}}_2, \end{aligned} \tag{120}$$

where the push-through identity is used to proceed from step 1 to step 2. We observe that the random part of $T_{\text{twas}}$ (i.e., $\hat{\boldsymbol{\beta}}_1^T\tilde{\boldsymbol{z}}_2$) differs from $U_\rho(\boldsymbol{\theta}_2)$ only by a scalar multiplier (i.e., $\sqrt{n_2}\sigma_2^{-2}\sigma_{\beta_1}^{-2}$). However, FUSION evaluates the variance of $\hat{\boldsymbol{\beta}}_1^T\tilde{\boldsymbol{z}}_2$ given that $\hat{\boldsymbol{\beta}}_1$ is a constant vector. In contrast, VINTAGE takes into account the uncertainty in the estimation of $\hat{\boldsymbol{\beta}}_1$ when constructing the gene-wise genetic variance test statistic. This becomes evident in view of the following equalities:

$$\text{Var}[\hat{\boldsymbol{\beta}}_1^T\tilde{\boldsymbol{z}}_2|\hat{\boldsymbol{\beta}}_1] = \hat{\boldsymbol{\beta}}_1^T\hat{\boldsymbol{R}}_2\hat{\boldsymbol{\beta}}_1, \tag{121}$$

$$\begin{aligned} \text{Var}[\hat{\boldsymbol{\beta}}_1^T\tilde{\boldsymbol{z}}_2] &= \mathbb{E}[\text{Var}[\hat{\boldsymbol{\beta}}_1^T\tilde{\boldsymbol{z}}_2|\hat{\boldsymbol{\beta}}_1]] + \text{Var}[\mathbb{E}[\hat{\boldsymbol{\beta}}_1^T\tilde{\boldsymbol{z}}_2|\hat{\boldsymbol{\beta}}_1]] \\ &= \mathbb{E}[\hat{\boldsymbol{\beta}}_1^T\hat{\boldsymbol{R}}_2\hat{\boldsymbol{\beta}}_1] \\ &= \sigma_1^{-4}tr[\boldsymbol{R}_2(\sigma_1^{-2}\boldsymbol{X}_1^T\boldsymbol{X}_1 + \sigma_{\beta_1}^{-2}\boldsymbol{I}_p)^{-1}\boldsymbol{X}_1^T \\ &\quad (\sigma_{\beta_1}^2\boldsymbol{X}_1\boldsymbol{X}_1^T + \sigma_1^2\boldsymbol{I}_{n_1})\boldsymbol{X}_1(\sigma_1^{-2}\boldsymbol{X}_1^T\boldsymbol{X}_1 + \sigma_{\beta_1}^{-2}\boldsymbol{I}_p)^{-1}] \\ &= \sigma_{\beta_1}^4 tr[\boldsymbol{R}_2\boldsymbol{X}_1^T(\sigma_{\beta_1}^2\boldsymbol{X}_1\boldsymbol{X}_1^T + \sigma_1^2\boldsymbol{I}_{n_1})^{-1}\boldsymbol{X}_1] \\ &= n_2^{-1}\sigma_{\beta_1}^4 tr[\boldsymbol{X}_2\boldsymbol{X}_1^T(\sigma_{\beta_1}^2\boldsymbol{X}_1\boldsymbol{X}_1^T + \sigma_1^2\boldsymbol{I}_{n_1})^{-1}\boldsymbol{X}_1\boldsymbol{X}_2^T] \\ &= n_2^{-1}\sigma_2^2\sigma_{\beta_1}^4\tilde{I}_{2,2}. \end{aligned} \tag{122}$$

Therefore, when serving as test statistics, $T_{\text{twas}}$ and $U_\rho(\boldsymbol{\theta}_2)$ can be expressed as follows:

$$T_{\text{twas}} = \frac{\hat{\boldsymbol{\beta}}_1^T \tilde{\boldsymbol{z}}_2}{\sqrt{\text{Var}[\hat{\boldsymbol{\beta}}_1^T \tilde{\boldsymbol{z}}_2 | \hat{\boldsymbol{\beta}}_1]}} \text{ vs. } \frac{U_\rho(\boldsymbol{\theta}_2)}{\sqrt{\tilde{I}_{2,2}}} = \frac{\hat{\boldsymbol{\beta}}_1^T \tilde{\boldsymbol{z}}_2}{\sqrt{\text{Var}[\hat{\boldsymbol{\beta}}_1^T \tilde{\boldsymbol{z}}_2]}}, \tag{123}$$

from which we can find that the TWAS test statistic would be equivalent to $U_\rho(\boldsymbol{\theta}_2)$ in the role of a test statistic if the uncertainty associated with $\hat{\boldsymbol{\beta}}_1$ is taken into account.

#### 3.4 Alternative attempts

The derivation of an effective gene-wise genetic variance test has posed significant technical challenges due to the concurrent presence of variance and covariance parameters under the null hypothesis, compounded by the compositional nature of the alternative hypothesis. Indeed, the test presented above stands out as the sole functional one among the five different tests we have carefully explored. The remaining four tests were unsuccessful for various reasons, and we provide a detailed summary of these failed attempts in the following for the benefit of the community.

1. The first two tests we explored are the standard score test and likelihood ratio test. The two test statistics are given by:

$$T_S = (U_{\sigma^2_{\beta_2}}, U_\rho)^T \tilde{\boldsymbol{I}}^{-1} (U_{\sigma^2_{\beta_2}}, U_\rho) \overset{asy.}{\sim} \frac{1}{2}\chi^2_1 + \frac{1}{2}\chi^2_2 \quad \text{(Score Test)}, \tag{124}$$

$$T_{LR} = -2 \log \frac{\max_{\Theta_0} L(\boldsymbol{\theta})}{\max_{\Theta} L(\boldsymbol{\theta})} \overset{asy.}{\sim} \frac{1}{2}\chi^2_1 + \frac{1}{2}\chi^2_2, \quad \text{(Likelihood Ratio Test)}, \tag{125}$$

where $\Theta_0$ denotes the null parameter space under $H_0$ and $\Theta$ denotes the full parameter space. Note that the second term of $U_{\sigma^2_{\beta_2}}$ in equation (93) cannot be omitted anymore for constructing $T_S$. For both tests, we derive the p value based on a 50:50 mixture of $\chi^2_1$ and $\chi^2_2$ following Stram and Lee [12] and Verbeke and Molenberghs [13]. Unfortunately, both tests produced inflated p values in the null simulations.

2. Besides the standard tests, we also explored the following test statistics that are formulated as a linear combination of the standardized variance and covariance scores:

$$\begin{aligned} T_{LC}^{(+)} &= \frac{U_{\sigma^2_{\beta_2}}}{\sqrt{\tilde{I}_{1,1}}} + \frac{U_\rho}{\sqrt{\tilde{I}_{2,2}}}, \\ T_{LC}^{(-)} &= \frac{U_{\sigma^2_{\beta_2}}}{\sqrt{\tilde{I}_{1,1}}} - \frac{U_\rho}{\sqrt{\tilde{I}_{2,2}}}. \end{aligned} \tag{126}$$

Above, $T_{LC}^{(+)}$ borrows information from the gene expression mapping study under a positive local genetic correlation between gene expression and the trait. $T_{LC}^{(-)}$, on the other hand, borrows information under a negative local genetic correlation. For both test statistics, we derive the p value by matching the first and second moments of the test statistic to a scaled chi-squared distribution, $\kappa\chi^2_\nu$, following the Satterthwaite method [14]. The scale parameter $\nu$ and the degrees of freedom $\kappa$ are given below:

$$\kappa = \frac{2\sigma_2^2\sqrt{\tilde{I}_{1,1}}}{n_2 p}, \nu = \frac{n_2^2 p^2}{4\sigma_2^4 \tilde{I}_{1,1}}. \tag{127}$$

Finally, we use the Cauchy combination rule to combine the two p values such that the test is robust to both positive and negative local genetic correlations. The p values derived based on the Satterthwaite method and the final combined p values are all calibrated with well-controlled type I errors. However, this test is considerably less powerful than SKAT when the local genetic correlation is around zero due to the inconsistent sign of the two scores. We attempted to address this issue by taking the absolute value of $U_\rho$ and used the property of a half-normal distribution to derive its moments. However, the Satterthwaite method produced inflated p values.

3. The last test we explored is to perform SKAT and TWAS separately following their corresponding testing procedures, and then combine the two p values using the Cauchy combination method. However, this test is less powerful than VINTAGE, especially when the true local genetic correlation takes a value other than 0 and $\pm 1$.

### 4 Gene-wise genetic correlation test

We have developed a gene-wise genetic correlation test to evaluate the potential role of gene expression in mediating the genetic variant and trait association. The test assesses the null hypothesis:

$$H_0 : r = 0, \tag{128}$$

which represents the scenario where the gene expression does not mediate any of the genetic effects on the trait. We carry out the gene-wise genetic correlation test only for genes that exhibit significance in the gene-wise genetic variance test because the local genetic correlation is defined only for genes with non-zero genetic variance.

We have developed a standard Rao's score test to carry out the above hypothesis testing. To derive the test statistic, we first obtain the score for the local genetic correlation, $r$, under the null hypothesis:

$$U_r = \frac{\partial l(\boldsymbol{\theta})}{\partial r}\Big|_{r=0} = \sigma_{\beta_1}\sigma_{\beta_2}\boldsymbol{y}_1^T(\sigma_{\beta_1}^2\boldsymbol{X}_1\boldsymbol{X}_1^T + \sigma_1^2\boldsymbol{I}_{n_1})^{-1}\boldsymbol{X}_1\boldsymbol{X}_2^T(\sigma_{\beta_2}^2\boldsymbol{X}_2\boldsymbol{X}_2^T + \sigma_2^2\boldsymbol{I}_{n_2})^{-1}\boldsymbol{y}_2. \tag{129}$$

We then follow similar procedures as described in the section 3 to derive the information for $r$.

Specifically, the information of $r$ takes the following form:

$$\tilde{I}_{rr} = I_{rr} - \boldsymbol{I}_{r\boldsymbol{\theta}_3}^T \boldsymbol{I}_{\boldsymbol{\theta}_3\boldsymbol{\theta}_3}^{-1} \boldsymbol{I}_{r\boldsymbol{\theta}_3}, \tag{130}$$

where $\boldsymbol{\theta}_3 = (\sigma_{\beta_1}^2, \sigma_{\beta_2}^2, \sigma_1^2, \sigma_2^2)$ are nuisance parameters and

$$I_{rr} = -\mathbb{E}\left[\frac{\partial^2 l(\boldsymbol{\theta})}{\partial r^2}\right], \boldsymbol{I}_{r\boldsymbol{\theta}_3} = -\mathbb{E}\left[\frac{\partial^2 l(\boldsymbol{\theta})}{\partial r \boldsymbol{\theta}_3^T}\right], \boldsymbol{I}_{\boldsymbol{\theta}_3\boldsymbol{\theta}_3} = -\mathbb{E}\left[\frac{\partial^2 l(\boldsymbol{\theta})}{\partial \boldsymbol{\theta}_3 \boldsymbol{\theta}_3^T}\right], \tag{131}$$

are evaluated under $H_0$ with $r = 0$. After some algebra, we obtain:

$$I_{rr} = \sigma_{\beta_1}^2 \sigma_{\beta_2}^2 tr(\boldsymbol{\Sigma}_{y_1}^{-1} \boldsymbol{X}_1 \boldsymbol{X}_2^T \boldsymbol{\Sigma}_{y_2}^{-1} \boldsymbol{X}_2 \boldsymbol{X}_1^T), \tag{132}$$

$$\boldsymbol{I}_{r\boldsymbol{\theta}_3} = (I_{r\sigma_{\beta_1}^2}, I_{r\sigma_{\beta_2}^2}, I_{r\sigma_1^2}, I_{r\sigma_2^2})^T = (0,0,0,0)^T, \tag{133}$$

$$\boldsymbol{I}_{\boldsymbol{\theta}_3\boldsymbol{\theta}_3} = \frac{1}{2}\begin{pmatrix} tr[(\boldsymbol{\Sigma}_{y_1}^{-1}\boldsymbol{X}_1\boldsymbol{X}_1^T)^2] & 0 & tr(\boldsymbol{\Sigma}_{y_1}^{-2}\boldsymbol{X}_1\boldsymbol{X}_1^T) & 0 \\ 0 & tr[(\boldsymbol{\Sigma}_{y_2}^{-1}\boldsymbol{X}_2\boldsymbol{X}_2^T)^2] & 0 & tr(\boldsymbol{\Sigma}_{y_2}^{-2}\boldsymbol{X}_2\boldsymbol{X}_2^T) \\ tr(\boldsymbol{\Sigma}_{y_1}^{-2}\boldsymbol{X}_1\boldsymbol{X}_1^T) & 0 & tr(\boldsymbol{\Sigma}_{y_1}^{-2}) & 0 \\ 0 & tr(\boldsymbol{\Sigma}_{y_2}^{-2}\boldsymbol{X}_2\boldsymbol{X}_2^T) & 0 & tr(\boldsymbol{\Sigma}_{y_2}^{-2}) \end{pmatrix}, \tag{134}$$

where $\boldsymbol{\Sigma}_{y_1} = \sigma_{\beta_1}^2 \boldsymbol{X}_1 \boldsymbol{X}_1^T + \sigma_1^2 \boldsymbol{I}_{n_1}$ and $\boldsymbol{\Sigma}_{y_2} = \sigma_{\beta_2}^2 \boldsymbol{X}_2 \boldsymbol{X}_2^T + \sigma_2^2 \boldsymbol{I}_{n_2}$ are the marginal variance of $\boldsymbol{y}_1$ and $\boldsymbol{y}_2$, respectively. Hence,

$$\tilde{I}_{rr} = I_{rr} = \sigma_{\beta_1}^2 \sigma_{\beta_2}^2 tr(\boldsymbol{\Sigma}_{y_1}^{-1} \boldsymbol{X}_1 \boldsymbol{X}_2^T \boldsymbol{\Sigma}_{y_2}^{-1} \boldsymbol{X}_2 \boldsymbol{X}_1^T). \tag{135}$$

Finally, the score test statistic is given by:

$$T_r = \frac{U_r^2}{\tilde{I}_{rr}} \sim \chi_1^2, \tag{136}$$

and we can calculate its associated p value, $p_r$, based on a chi-squared distribution with the degrees of freedom equal to one.

Similarly, the score, information, and test statistic can be expressed with summary statistics under the transformed models in the following:

$$U_r = \sigma_1^{-2} \sigma_2^{-2} \sigma_{\beta_1}^{-1} \sigma_{\beta_2}^{-1} \sqrt{n_1 n_2} \tilde{\boldsymbol{z}}_1^T \hat{\boldsymbol{Q}}_{:,1:k} \boldsymbol{\Sigma}_{\tilde{\beta}_1^*} \boldsymbol{\Sigma}_{\tilde{\beta}_2^*} \hat{\boldsymbol{Q}}_{:,1:k}^T \tilde{\boldsymbol{z}}_2, \tag{137}$$

$$\tilde{I}_{rr} = \sigma_1^{-2} \sigma_2^{-2} n_1 n_2 tr[\boldsymbol{\Sigma}_{\tilde{\beta}_1^*} \boldsymbol{\Sigma}_{\tilde{\beta}_2^*} (\hat{\boldsymbol{\Lambda}}_1)_{1:k} (\hat{\boldsymbol{\Lambda}}_2)_{1:k}], \tag{138}$$

$$T_r = \sigma_1^{-2} \sigma_2^{-2} \sigma_{\beta_1}^{-2} \sigma_{\beta_2}^{-2} \frac{(\tilde{\boldsymbol{z}}_1^T \hat{\boldsymbol{Q}}_{:,1:k} \boldsymbol{\Sigma}_{\tilde{\beta}_1^*} \boldsymbol{\Sigma}_{\tilde{\beta}_2^*} \hat{\boldsymbol{Q}}_{:,1:k}^T \tilde{\boldsymbol{z}}_2)^2}{tr[\boldsymbol{\Sigma}_{\tilde{\beta}_1^*} \boldsymbol{\Sigma}_{\tilde{\beta}_2^*} (\hat{\boldsymbol{\Lambda}}_1)_{1:k} (\hat{\boldsymbol{\Lambda}}_2)_{1:k}]}, \tag{139}$$

where $\boldsymbol{\Sigma}_{\tilde{\beta}_1^*} = [n_1 \sigma_1^{-2} (\hat{\boldsymbol{\Lambda}}_1)_{1:k} + \sigma_{\beta_1}^{-2} \boldsymbol{I}_k]^{-1}$ and $\boldsymbol{\Sigma}_{\tilde{\beta}_2^*} = [n_2 \sigma_2^{-2} (\hat{\boldsymbol{\Lambda}}_2)_{1:k} + \sigma_{\beta_2}^{-2} \boldsymbol{I}_k]^{-1}$ are the conditional variance of $\tilde{\boldsymbol{\beta}}_1^*$ and $\tilde{\boldsymbol{\beta}}_2^*$, respectively.

### 5 Additional simulation and real data analyses

#### 5.1 Evaluation of COJO and MESuSiE for estimating the number of non-zero effect SNPs

We conducted simulations to evaluate two existing methods, COJO [15] and MESuSiE [16], for their ability to accurately estimate the number of SNPs with non-zero effects on gene expression and the trait. Simulation details are provided in the Materials and methods in the main text. In the simulations, for each gene of focus, we randomly selected one shared cis-SNP to have a non-zero effect on both gene expression and the trait, and selected two additional cis-SNPs – one with a non-zero effect on gene expression and the other with a non-zero effect on the trait. We found that while both methods tended to underestimate the number of SNPs with non-zero effects, likely due to low power, MESuSiE provided more accurate estimates than COJO (Figure S34). Specifically, MESuSiE estimated an average of 1.2 and 1.0 SNPs to have non-zero effects on gene expression and the trait, respectively, whereas COJO estimated only 0.74 and 0.49 SNPs. Notably, MESuSiE explicitly models shared and study-specific SNPs, allowing it to provide more accurate estimates of the number of shared SNPs compared to COJO. On average, MESuSiE estimated 0.71 shared SNPs, while COJO only estimated 0.042 shared SNPs. Therefore, we decided to use MESuSiE to estimate the number of SNPs with non-zero effects on gene expression and the trait in the real data.

#### 5.2 A complication for evaluating type I error and power under a sparse genetic architecture

For simulations under a sparse genetic architecture, there is an important complication for evaluating the type I error and power. To understand this complication, let us first consider a simple example with two distinct sparse scenarios. In this example, we consider $q = 1$ for simplicity. In the first sparse scenario, a single SNP in the genomic region is causal, affecting both gene expression and the trait. This scenario clearly represents an alternative setting of non-zero local genetic correlation with true $r = \pm 1$, indicating that the SNP's effect on the trait is potentially mediated through gene expression. We evaluated this scenario in Figures S20-S22, where VINTAGE's local genetic correlation test demonstrated clear power. The complexity, however, arises from the simulation of this setting. When a bivariate normal distribution is used to simulate causal SNP effects on both gene expression and the trait, the correlation parameter in the bivariate normal distribution, denoted as $\tilde{r}$, can take any value within $[-1, 1]$. Regardless of the value of $\tilde{r}$, the resulting scenario remains unchanged: the causal SNP affects both gene expression and the trait, leading to the true $r = \pm 1$. Therefore, in this sparse scenario, $\tilde{r}$ becomes non-informative, as any value of $\tilde{r}$ produces the same outcome of $r = \pm 1$.

In the second sparse scenario, a single SNP in the genomic region is causal for gene expression, while a different SNP is causal for the trait. This scenario clearly represents a null setting of zero local genetic correlation, where true $r = 0$ and the SNP's effect on the trait is not mediated through gene expression. We evaluated this scenario in Figures S12-S14, where VINTAGE's local genetic correlation test achieved calibrated type I error control.

As you can see, the complexity here arises from the discrepancy between the simulation parameter ($\tilde{r}$) and the true underlying local genetic correlation ($r$), a situation that does not occur in non-sparse settings where $\tilde{r}$ is always consistent with $r$. Certainly, the situation becomes even more complicated in the first sparse setting when $q > 1$; in our study, we also explored $q = 2$ and $q = 10$. In these scenarios, the true $r$ is no longer $+1$ or $-1$, but rather is enriched around these values with variability across simulation replicates. To illustrate this, since we know the true effect sizes in simulations, we can compute in each simulation replicate the true $r$ by using the Pearson correlation coefficient between the simulated genetic effects on gene expression and on trait (Figures S20-S22). Consistent with our explanation, $r$ was either $+1$ or $-1$ when $q = 1$, while the distribution of $r$ varied between $-1$ and $+1$ with enrichment around these two values when $q = 2$ or $q = 10$. In addition, in some simulation replicates, $r$ was very close to zero, and these replicates allowed us to directly demonstrate that VINTAGE's local genetic correlation test has calibrated type I error control under the first sparse scenario (Figures S20-S22).

##### 5.3 Exploration of expression pleiotropy

We carefully examined the 61 genes detected by the gene-wise genetic correlation test of VINTAGE and identified four sets of genes that might be associated with the phenotype due to expression pleiotropy. The genes within each of these sets reside in proximity on the genome and share the same top eQTLs. The four gene sets and their associated phenotypes include 1) CDC123 and CAMK1D with FEV1; 2) CIAO3, HAGHL, and ANTKMT with SH; 3) ITGA4 and CERKL with WBC; and 4) TNFRSF10B and TNFRSF10C with WBC (Figure S43). In these cases, it is possible that the SNP effects on the trait are mediated by one of the genes rather than all genes within the set. This observation highlights the need of developing fine-mapping methods based on the VINTAGE framework in the future. In addition to the four sets of genes, we also identified some additional genes that are located near each other on the genome but do not appear to be associated with the trait due to expression pleiotropy. For example, CREB5 and JAZF1 are located nearby and display significant local genetic correlation with WBC, but do not share any top eQTLs (Figure S44).

[9] Zhiwen Jiang, Haoyu Zhang, Thomas U Ahearn, Montserrat Garcia-Closas, Nilanjan Chatterjee, Hongtu Zhu, Xiang Zhan, and Ni Zhao. The sequence kernel association test for multicategorical outcomes. *Genetic Epidemiology*, 2023.
